# Phylogenetic, ecological and intraindividual variability patterns in grass phytolith shape

**DOI:** 10.1101/2021.08.12.456097

**Authors:** Kristýna Hošková, Jiří Neustupa, Petr Pokorný, Adéla Pokorná

## Abstract

- Grass silica short cell (GSSC) phytoliths appear to be the most reliable source of fossil evidence for tracking the evolutionary history and paleoecology of grasses. In recent years, modern techniques have been used to quantitatively assess phytolith shape variation. This progress has widened opportunities with respect to the classification of grass fossil phytoliths. However, phylogenetic, ecological and intraindividual variability patterns in phytolith shape remain largely unexplored.
- The full range of intraindividual phytolith shape variation (3650 2D outlines) from 73 extant grass species, 48 genera, 18 tribes, and 8 subfamilies (with special attention paid to Pooideae) was analysed using the geometric morphometric analysis based on the semilandmarks spanning phytolith outlines.
- Although we showed that 2D phytolith shape is mainly driven by deep-time diversification of grass subfamilies, a closer look uncovered distinct phytolith shape variation in early-diverging lineages of Pooideae.
- The phylogenetic pattern in phytolith shape was successfully revealed by applying geometric morphometrics to 2D phytolith shape outlines. This finding strengthens the potential of phytoliths to track the evolutionary history and paleoecology of grasses. Moreover, geometric morphometrics of 2D phytolith shape proved to be an excellent tool for analysis requiring large sums of phytolith outlines, making it useful for quantitative palaeoecological reconstruction.

## Introduction

As the plant silica microfossils, phytoliths have the great potential to track the evolutionary history and paleoecology of grasses (Poaceae) (Strömberg, 2005; Prasad *et al*., 2011; Strömberg, 2011; Strömberg, 2013). Due to their composition of biogenic opaline silica (SiO2.nH2O), phytoliths are preserved in various sedimentary environments, even when other grass fossils are not (Piperno, 2006; Strömberg *et al*., 2018). Grass phytoliths, and grass silica short cell (GSSC) phytoliths, in particular, are known to vary in shape depending on multiple taxonomic levels (subfamilies, tribes, sometimes even genera; Metcalfe, 1960; Mulholland & Rapp, 1992; Piperno & Pearsall, 1998; Rudall *et al*., 2014), whereas other grass fossil remains such as pollen, leaves, or seeds are either rare or not taxonomically informative below the family level (Jacobs *et al*., 1999; but for the case of pollen see Mander *et al*., 2013). The recent work of Gallaher *et al*. (2020) quantitatively documented that GSSC phytolith 3D shape carries a strong phylogenetic signal, one which can distinguish grass subfamilies and tribes. They investigated 70 species of early-diverging grasses, plus Oryzoideae and Bambusoideae. However, phylogenetic, ecological, and intraindividual variability patterns in phytolith shape in other grass taxonomic groups remain unexplored.

The greatest amount of grass diversity falls in either of two clades, the BOP (Bambusoideae, Oryzoideae, Pooideae) and PACMAD clade (Panicoideae, Arundinoideae, Chloridoideae, Micrairoideae, Aristidoideae, and Danthonioideae) (Strömberg, 2011; Soreng *et al*., 2015, 2017). Twiss *et al*. (1969) proposed three major divisions of GSSC phytoliths corresponding to the three dominant grass subfamilies native to the Great Plains of the United States: bi-/poly-lobate/cross in Panicoideae, saddle in Chloridoideae, and circular/oblong/rectangular in Pooideae. These divisions are mainly useful for phytolith research in regions where grasses are abundant and diverse on this (sub-familiar) taxonomic level [(for example, in grasslands of the North American Great Plains (Fredlund & Tiezsen, 1994) and Sub-Saharan Africa (Barboni *et al*., 2007; Bremond *et al*., 2005)]. However, they are less informative in regions where only one subfamily dominates (and has dominated throughout long-term history), as, for example, in large areas of temperate and boreal regions of Eurasia, where subfamily Pooideae prevails (Gibson, 2009).

Pooideae is the largest Poaceae subfamily, with almost 4000 species, most of them adapted to open and cold environments (Bouchenak-Khelladdi *et al*. 2010; Edwards & Smith, 2010; Soreng *et al*., 2017; Schubert *et al*., 2019a, b). While considerable effort has been made to refine the categorization of phytolith shape variation within subfamilies like Panicoideae, Chloridoideae, Bambusoideae and Oryzoideae (Fahmy, 2008; Novello *et al*., 2012; Lu & Liu 2003; Neumann *et al*., 2017; Cai & Ge 2017; Gallaher *et al*., 2020), phytolith shape variation in Pooideae is largely unexplored, with only a handful of studies touching this area (for example in studies where *Stipa* type phytoliths are recognised, Gallego & Distel 2004; Silantyeva *et al*., 2018; Mullholand, 1989).

Individual grass subfamilies are mostly adapted to certain environmental conditions and tend to prevail in specific vegetation zones (Gibson, 2009). In phytolith analysis, this association is used to define indices that can be applied as proxies of past environments. For example, the so-called aridity index (Iph; Diester-Haass *et al*. 1973; Alexandre *et al*., 1997) presents the proportion of Chloridoideae prevailing in arid conditions (saddle-shaped morphotypes) and Panicoideae prevailing in more humid conditions (bilobates, crosses), and can be used as a proxy for aridity in past ecosystems. Similarly, the climatic index (Ic; Twiss, 1992) can be used to reconstruct past climates on the basis of the proportion of morphotypes characteristic of Pooideae adapted to open and cold environments (rondels) versus the proportion of morphotypes characteristic of Chloridoideae and Panicoideae, which are adapted to higher temperatures. However, it is untenable to use phytolith spectra *per se* for the indication of past habitat conditions without a knowledge of the distribution of phytolith variation across the phylogenetic tree, including the proportion of intraspecific variation.

Our recent study (Hošková *et al*., 2021) observed very low GSSC phytolith shape plasticity between populations of the same species, compared to interspecific variation, according to a hierarchically designed study of two grass species with restricted ecological niches (*Brachypodium pinnatum, B. sylvaticum*). Thus, we considered phytolith shape variation due to environmental conditions to be negligible. However, significantly high residual variation (51% of the total variation, which was not captured by the defined levels: species, populations, individuals, leaves and parts of leaves) was related to the intraindividual variation of phytolith shape within the individual sample. This type of variation (which is intrinsic to phytoliths and demarcates it from variation in other fossils) causes species to carry more than one phytolith morphotype, resulting in an overlap of phytolith shapes between taxa (e. g. Rovner & Russ, 1992). Piperno & Pearsall (1998), in their seminal work, proposed that intraindividual variation in phytolith shape may be lineage-specific; thus, species could vary in the amount of intraindividual phytolith shape variation on the basis of their position in the phylogenetic tree. Thus, knowledge of the full range of intraindividual variation could complete our image of the phylogenetic pattern in phytolith shape.

In order to quantify the proportion of intraindividual variation in relation to the phylogenetic pattern in phytolith shape, we used methods of geometric morphometrics (Hošková *et al*., 2021). We explored whether GSSC phytolith shapes changed in response to diversification during grass evolution and how closely phytolith shapes reflect phylogenetic relationships in the Poaceae family. As the degree of intraindividual variation in phytolith shape in most grasses has never been explored, we aimed to separate the intraindividual variation from the variation due to phylogeny. We paid particular attention to the understudied grass subfamily Pooideae, adapted to open and cold conditions, focusing, on the potential affinity of ecological adaptation of species to phytolith shape variation within this group.

Geometric morphometrics is undoubtedly one of the most frequently applied techniques of the biological shape analysis today (Polly & Motz, 2016). It enables size to be effectively removed and thus focuses purely on the analysis of shape, allowing the simple collection of coordinate data and easy visualisation of results as transformations of shapes themselves rather than as tables of numbers (Bookstein, 1989; Klingenberg, 2013). Hence, unlike the traditional description of predefined phytolith morphotypes {(International Code for Phytolith Nomenclature 2.0 [International Committee for Phytolith Taxonomy (ICPT), 2019]}, geometric morphometrics allows the quantification and visualisation of a continuous phytolith shape variation. Therefore, different phytolith morphotypes are analysed together within a single multivariate space (morphospace), and, as a result, the major trends in shape variation are explored. The Procrustes superimposition method, which is the core of the geometric morphometric analysis, relies on the point-to-point correspondence of individual landmarks among the analysed specimens. In case of phytolith outlines, these points are represented by series of equidistant semilandmarks. In those phytoliths having symmetric 2D shapes, there may be two or more fixed points delimiting individual symmetric curves. These points are typically derived from the orientation of phytoliths within plant tissue (Hošková *et al*., 2021).

## Materials and Methods

### Plant material processing

3650 modern grass phytoliths from 73 species, 48 genera, 18 tribes, 8 subfamilies were analysed (Table S1).

Plant material was processed following the *in situ* charring method of Kumar *et al*. (2017). This method preserves the original phytolith position within the plant epidermis. One leave per plant per species was sampled. Leaves were cleaned in an ultrasonic cleaner (Digital Ultrasonic Cleaner CE-7200A). The segment of leave was laid on a glass slide. Small pieces of folded aluminium foil were placed near the two shorter sides of the slide. Another glass slide was placed on top of the slide, holding the sample in place. The slides were put into a muffle furnace at 550 °C for 5 h. The aluminium foil between the slides prevented the slides from sticking together. The slides containing burnt material were washed with 1 _N_ HCl and distilled water (using a pipette). After the slides dried, plant material placed on the bottom slide was covered with one drop of a 15 % solution of glycerol and the cover slide. The slides were then analysed using transmission light microscopy (Leica DM 1000 LED).

### Data acquisition

Sequential microphotographs of rows of GSSC phytoliths in the charred epidermis were acquired under ×400 magnification (Leica camera ICC50 W). The planar view of GSSC costal (over veins) phytolith morphotypes with a long axis parallel with the long axis of the leaf was chosen for the analysis. First, two fixed landmarks were placed at the phytolith edges perpendicular to the longest axis of the leaf (Fig. 1; Methods S1). Then, 48 equidistant points were placed along both outline halves, resulting in 96 points which were treated as semilandmarks in the subsequent geometric morphometric analysis. For each individual phytolith image (3650 in total), 98 two-dimensional points were digitised. This approach was only applied to phytolith morphotypes observed from the planar view called bilobates, polylobates, saddles, crenates and trapezoids {(International Code for Phytolith Nomenclature 2.0 [International Committee for Phytolith Taxonomy (ICPT), 2019]}, whereas rondels, positioned from the planar view in leaf epidermis, had no identifiable landmarks and were not analysed. During the plant material processing, segments of leaves were laid on a glass slide with random orientation regarding abaxial-adaxial leaf sides; however, the outer periclinal surface of phytoliths was chosen for phytolith outline analysis since under light microscopy it has clear, distinct edges that are well defined. Digitisation was carried out using the semiautomated background curves tool in TpsDig, ver. 2.31 (Rohlf, 2015). Equidistant positions of semilandmarks along the outlines relative to the positions of the fixed landmarks were obtained using the ‘*digit.curves*’ function in geomorph, v. 3.3.2 (Dryen & Mardia, 2016; Adams *et al*., 2021), in R, v. 3.6.3 (R Core Team, 2020) (for summarized methodological workflow see Supplemental files Methods S2).

**Fig. 1.**
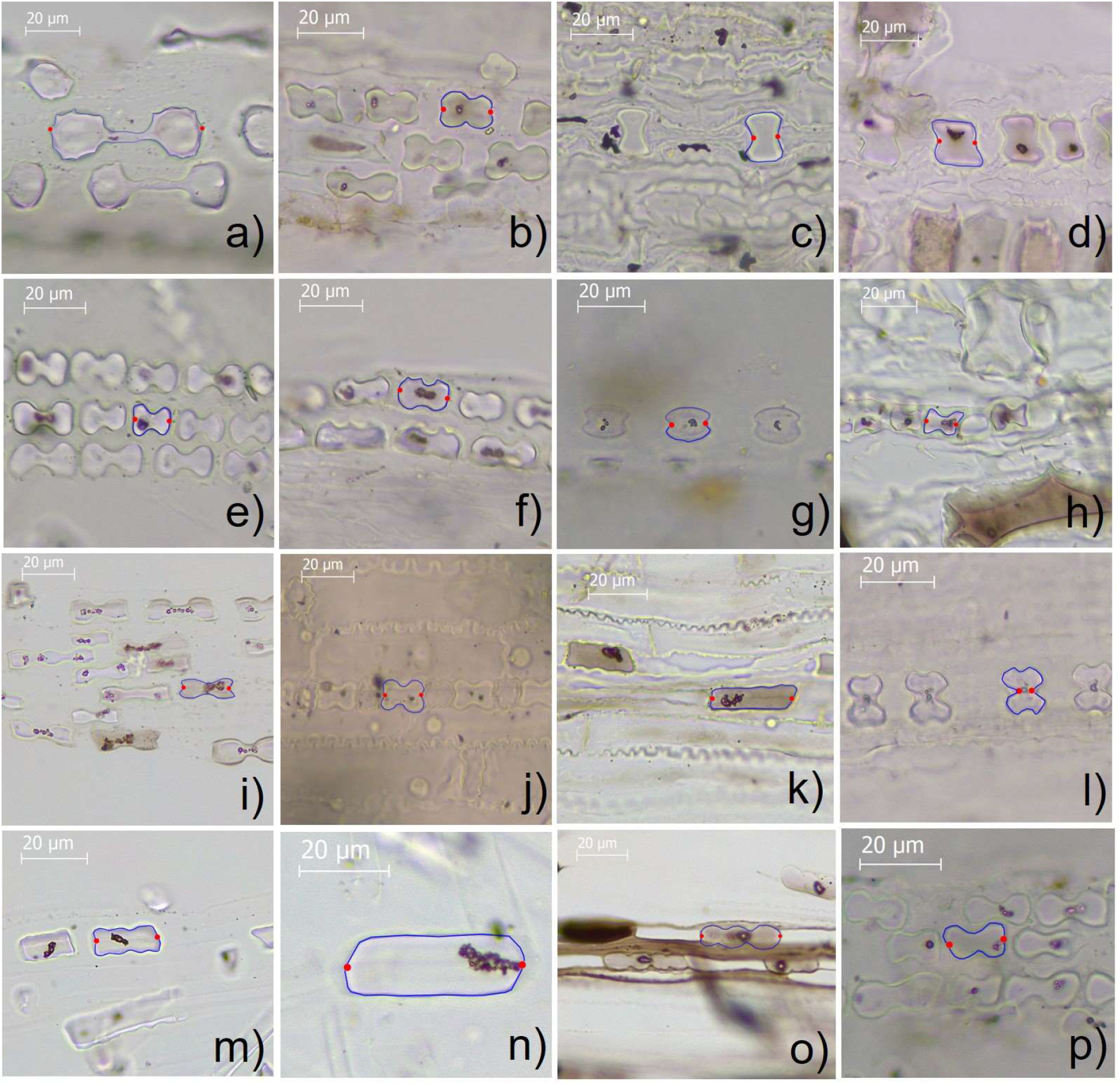
Microphotographs of charred grass epidermises with GSSC phytoliths. Red dots indicate two fixed landmarks; blue lines indicate phytolith outline with 96 equidistant semilandmarks. (a) *Aristida rhiniochloa*, Aristidoideae; (b) *Hakonechloa macra*, Arundinoideae; (c) *Bambusa tuldoides*, Baumbusoideae; (d) *Arundinaria gigantea*, Bambusoideae; (e) *Danthonia alpina*, Danthonioideae; (f) *Schismus arabicus*, Danthonioideae; (g) *Cynodon dactylon;* (h) *Brachyelytrum erectum*, Pooideae; (i) *Bothriochloa ischaemum*, Panicoideae; (j) *Coix lacrymajobi*, Panicoideae; (k) *Alopecurus pratensis*, Pooideae; (l) *Zizania latifolia*, Oryzoideae; (m) *Bromus erectus*, Pooideae; (n) *Holcus lanatus*, Pooideae; (o) *Melica picta*, Pooideae; (p) *Stipa sibirica*, Pooideae.

### Data analysis

#### Generalised Procrustes Analysis of phytoliths with biradial symmetry

Geometric morphometrics was performed on a data set of 3650 phytolith configurations, each consisting of 98 landmark coordinates (following Hošková *et al*., 2021). The 2D shape of phytoliths from planar view (long axis of phytolith parallel with the long axis of the leaf) is typical by its biradial symmetry, meaning that left-right and upper-bottom parts of the 2D outline are not differentiated. Thus, to achieve correspondence of all phytolith configurations under study, we applied geometric morphometrics for the analyses of biradial symmetry (Savriama *et al*., 2010; Savriama & Klingenberg, 2011; Savriama *et al*. 2012; Neustupa, 2013) (Methods S1). A Generalised Procrustes Analysis, which minimises the sum of squared distances between corresponding landmarks to extract shape data by removing the extraneous information of size, location and orientation, was applied (e. g. Zelditch *et al*., 2012; Dryden & Mardia, 2016). The semilandmark position was optimised by their iterative sliding along the curve tangents to achieve the lowest bending energy yielding the smoothest possible deformation between each configuration and the mean shape (Bookstein, 1997; Pérez *et al*., 2006; Gunz & Mitteroecker, 2013). Original phytolith configurations were transformed and re-labelled and then subjected to a Generalised Procrustes Analysis. The resulting multiplied dataset consisted of Procrustes coordinates of original configurations and transformed and relabelled copies (a reflected copy about the horizontal adaxial-abaxial axis; a reflected copy about the vertical left-right axis; a copy reflected about both axes) (Savriama & Klingenberg, 2011; Klingenberg, 2015; Savriama, 2018; Hošková *et al*. 2021). By averaging the original configuration and transformed copies of each specimen, symmetrised phytolith configurations were obtained. These are symmetric and thus invariant under all transformations. A Generalised Procrustes Analysis was conducted using the *‘procGPA’* function in shapes, v. 1.2.5 in R, v. 3.6.3.

#### Quantification of symmetric and asymmetric components of shape variation

Principal component analysis (PCA) was conducted with the superimposed Procrustes coordinates consisting of all the original configurations and their transformed copies. This PCA separated components of symmetric shape variation (variation between symmetrised configurations) from three components of asymmetry (asymmetry under reflection in the adaxial-abaxial direction, asymmetry in the left-right direction and asymmetry regarding both these axes) (Savriama *et al*., 2010; Klingenberg, 2015). The proportion of variation in the subspaces of biradial symmetry and three asymmetric patterns were quantified by summing up the percentages of variance explained by PCs belonging to a given subspace using scores from PCs obtained by *‘procGPA’* function in shapes, v. 1.2.5 in R, v. 3.6.3.

#### Quantification of different sources of shape variation

Different sources of the shape variation among phytoliths were quantified by multivariate Procrustes analysis of variance (ANOVA) of the symmetrised configurations of individual phytoliths (e. g. Klingenberg, 2015). Data were analysed in a nested structure that was reflected by the Procrustes ANOVA models. The analysis decomposed the matrix of Procrustes distances among individual configurations into different sources specified by the independent factors. Besides quantifying the Procrustes sum of squares (SS) spanned by each factor and their proportion on the total variation (η^2^), the significance of the effects were evaluated by comparing their original Procrustes SS values with their random distribution yielded by 999 permutations (Schaefer *et al*., 2006; Neustupa & Woodard, 2021). The randomisation design reflected the nested structure of the independent factors. The main effect evaluating the differentiation between the phytoliths of the BOP and PACMAD lineages was tested against the random distribution based on the repeated reshuffling of individual subfamilies between BOP and PACMAD. Likewise, the SS spanned by the ‘subfamily’ effect nested within ‘BOP vs. PACMAD’ were evaluated by comparison with the random distribution yielded by reshuffling of tribes among subfamilies within the BOP and PACMAD groups. Then, the effect of tribes was tested against the random SS distribution based on the reshuffling of genera among different tribes, and the effect of genus identity on phytolith shapes was tested against the random SS distribution based on a reshuffling of species among genera. Finally, the differentiation of phytoliths among the species was evaluated by randomisation of individual specimens. The function *‘procD.lm’* in geomorph, v. 3.3.2 (Dryen & Mardia, 2016; Adams *et al*., 2021), in R, v. 3.6.3 (R Core Team, 2020) was used for the decomposition of sources of phytolith shape variation. Pairwise randomised residual permutation procedure*posthoc* tests were performed using *‘pairwise’* function in rrpp v. 0.5.2 in R, v. 3.6.3.

To visualise the discrimination of grass subfamilies by their phytolith shape, canonical variates analysis (CVA) was performed on symmetrised phytolith shape configurations in MorphoJ (Klingenberg, 2011).

#### Amount of intraindividual variation in phytolith shape in individual grass species

The amount of intraindividual variation in phytolith shape in individual grass species, represented by average Procrustes distances of individual phytoliths to species centroids, was compared using the function *‘betadisper’* of in vegan (Oksanen *et al*. 2019) in R v. 3.6.3.

#### Phytolith shape and grass phylogeny

Grass phylogeny was generated (using S3 scenario) with function *‘phylo.maker’* in v.phylomaker, v.0.1.0 in R, v. 3.6.3 (Jin & Qian 2019) using ‘backbone’ tree based on molecular data from seed plant phylogeny (mega-tree ‘GBOTB.extended.tre’; Smith & Brown, 2018). Out of the 73 species examined, 54 species were in the Smith & Brown (2018) backbone tree with the rest added using the S3 scenario of Jin & Qian (2019).

To visualise the phylogenetic history of phytolith shape change, grass species positions along PC1 were mapped on the grass phylogeny tree. Likewise, PCs spanning different components of asymmetric variation were mapped on this tree. Intraindividual shape variation, represented by averaged distances from species group centroid in multivariate space, was also mapped on the grass phylogeny. We used function *‘phylosig’* in R/phytools v. 0.7.47 (Revell, 2012) to determine Pagel’s lambda as a measure of phylogenetic signal in individual components of the shape data (individual PCA axes, amount of intraindividual variation). We used function *‘pgls’* in R/caper, v. 1.0.1 to calculate confidence intervals (Orme *et al*., 2018). We used function *‘contMap’* to visualise the phylogenetic history of individual components of the shape data, using the function *‘fastAnc’* in R/phytools v. 0.7.47 to reconstruct maximum likelihood values at tree nodes (Revell, 2012).

The phylogenetic tree of the studied taxa was projected onto the shape tangent space by squared-change parsimony in MorphoJ (Klingenberg, 2011). The resulting tree was plotted in the plane of the PC1 vs PC2 ordination plot of the species-level averaged shapes.

## Results

### Decomposition of symmetric and asymmetric variation in phytolith shape

The first group of PCs, associated with entirely symmetric shape variation, accounted for 89.5 % of the total variation of the data set. The second group of PCs, associated with three subspaces of asymmetric shape variation, accounted for 11.0 % of the total variation. Thus, the PCA results indicated that phytolith shape variation consisted mainly of symmetric variation between individual phytoliths and relatively little asymmetric shape variation within phytoliths (e. g. between their left and right sides). Therefore, in the following analyses of phytolith shape, we considered only the data set of the symmetrised phytolith configurations.

### Sources of variation in phytolith shape

Individual taxonomic levels accounted for 81.9 % of the total variation in symmetric phytolith shapes (Fig. 2, Table 1). Residual variation consisting of phytolith shape variation within individuals was considerably lower (18.2 %). Variation between subfamilies was the most pronounced (42.7 %, P = 0.001), followed by variation between tribes (14.0 %, P=0.001), species (13.3 %, P=0.001), and genera (10.4 %, P=0.9). The phytolith shape variation between PACMAD and BOP clades accounted for only 1.4 % of the total variation (P=0.7).

**Fig. 2.**
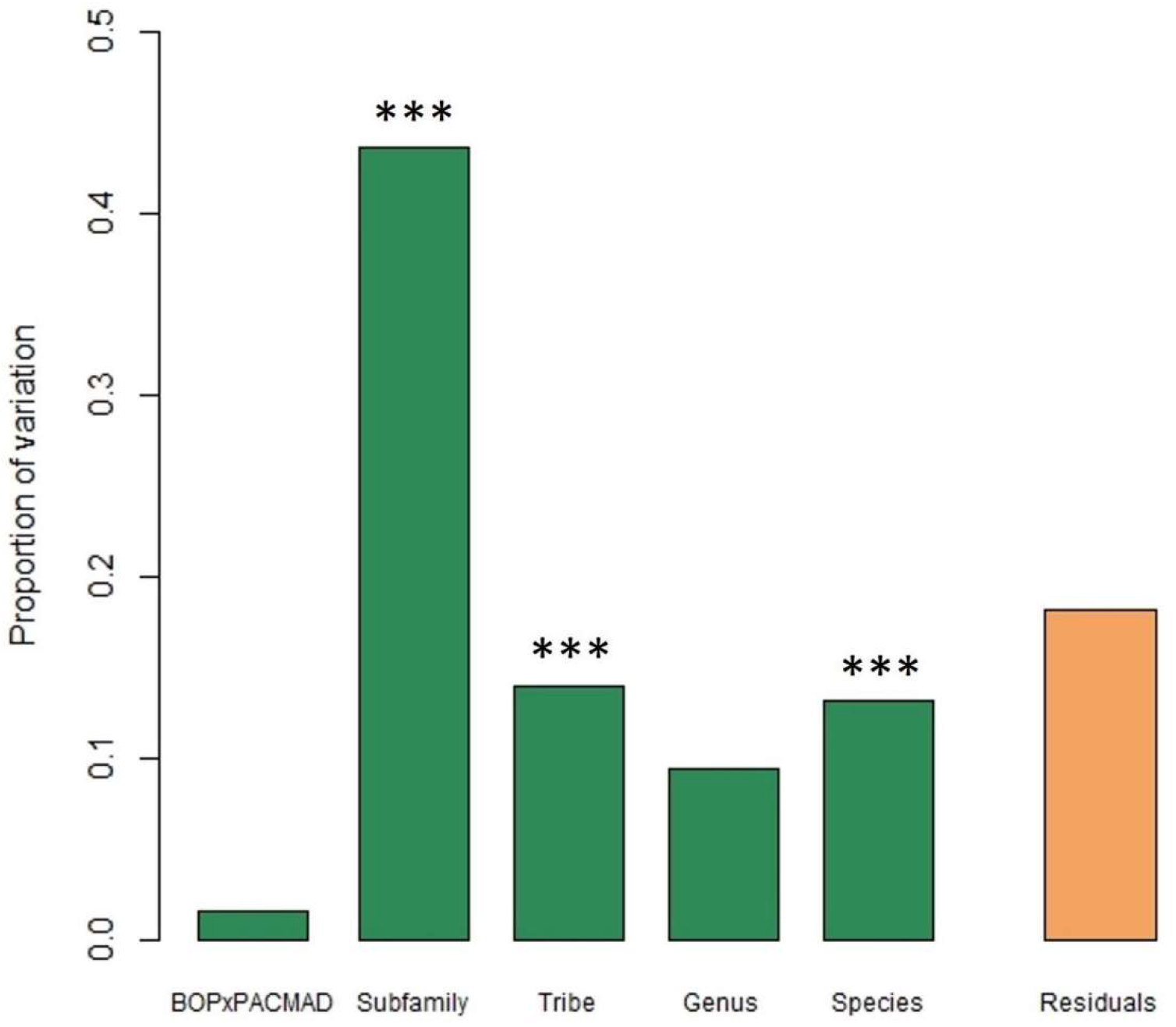
Coloured bars represent the proportion of variation in phytolith shape attributed to the various taxonomic levels and residual intraindividual variation. Stars indicate statistically significant differences in phytolith shape between the subfamilies, the tribes and the species. ***, P < 0.001

**Table 1.**
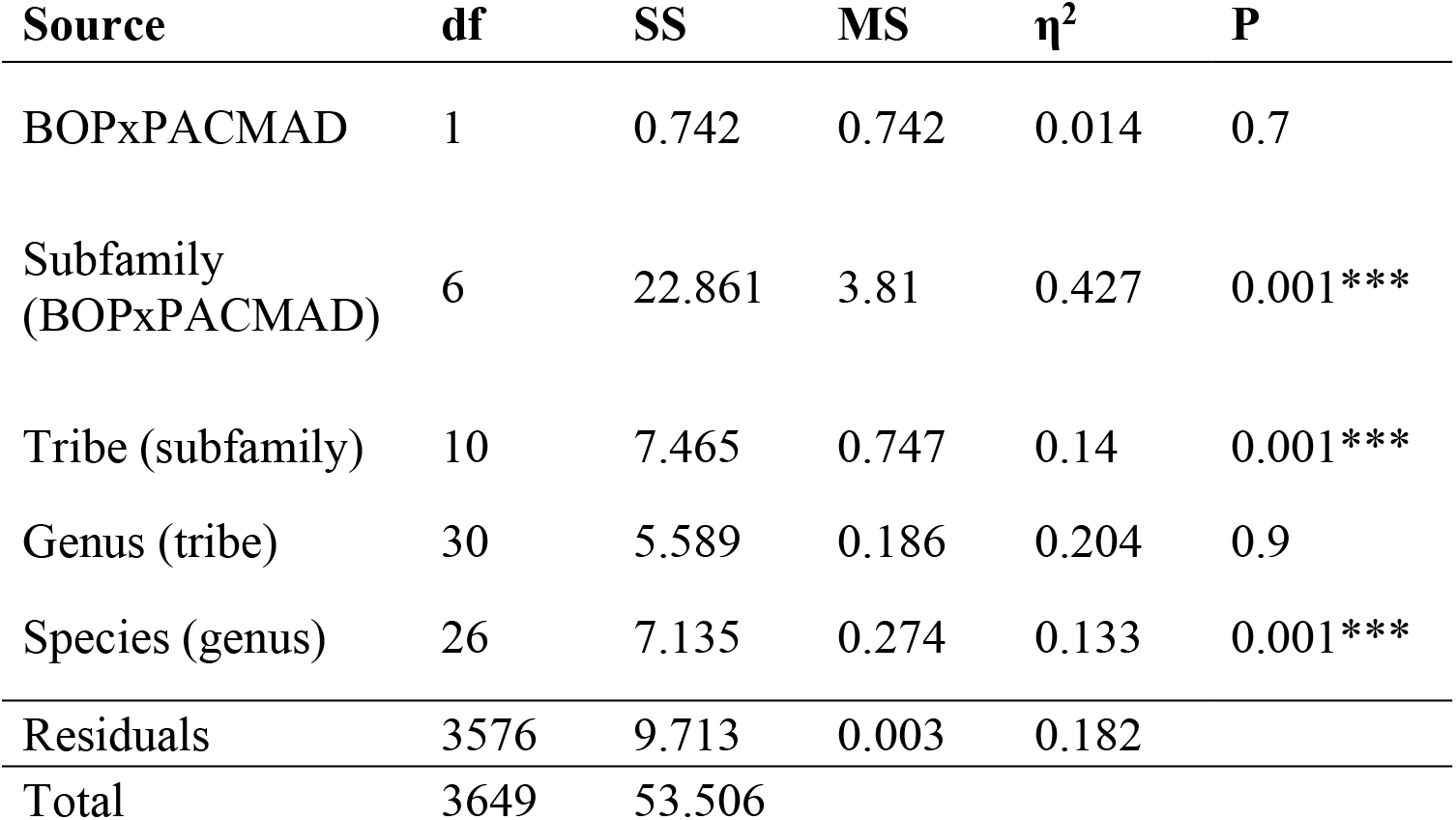
Results of Procrustes ANOVA evaluating different sources of variation in symmetrised phytolith shape. df: degrees of freedom; SS: sum of squares; MS: means squares; η^2^: proportion on the total variation; P: probability of the null hypothesis. ***, P < 0.001

Significant differences in phytolith shape were found for all the 28 subfamily pairs in the *posthoc* pairwise tests (at the significance level of 0.01) (Fig. 3; Table S2).

**Fig. 3.**
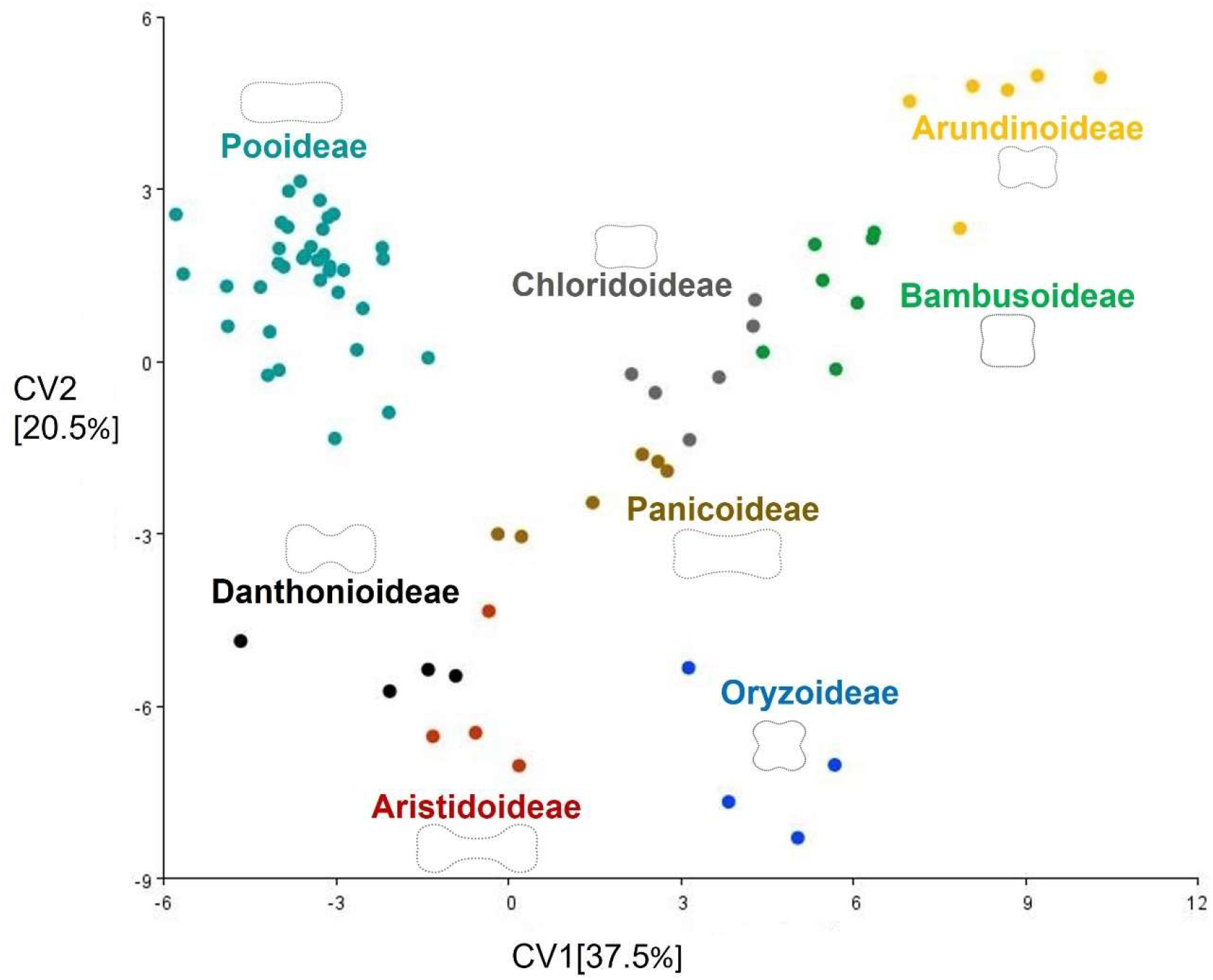
Canonical variates analysis used to discriminate between grass subfamilies by symmetrised grass silica short cell phytoliths shape. Dots represent averaged phytolith shape of individual grass species. Colours of dots indicate grass subfamilies. Phylogenetic signal in phytolith shape is apparent from the distinctive position of individual grass subfamilies in the morphospace.

### Intraindividual phytolith shape variation across species

The amount of intraindividual GSSC phytolith shape variation differed across species (Fig. 4; Notes S1). The phytolith shape of some species varied to such an extent that two different morphotypes, as traditionally defined, occurred – we reported variation from bilobate-to saddle-shaped phytoliths in *Eragrostis minor*, and variation from bilobate-to polylobate-shaped phytoliths in *Brachypodium dystachion, B. pinnatum, Bromus inermis, Calamagrostis epigejos, Bothriochloa ischaemum, Coix lacryma-jobi*, and *Digitaria sanguinalis*. Other species, like *Bromus erectus* (tribe Bromeae), *Melica picta* (tribe Meliceae), *Festuca gigantea, Festuca arundinacea, Helictotrichon pubescens, Milium effusum, Phleum pratense, Poa nemoralis*, and *Trisetum flavescens* (tribe Poeae) significantly varied intraindividually in the length and width of GSSC phytoliths (the so-called crenate morphotype). *Ehrharta erecta* (tribe Ehrharteae) varied intraindividually in the length of the polylobate phytolith shape, whereas *Brachyelytrum erectum* (tribe Brachyelytreae), *Dichanthium annulatum* (tribe Andropogoneae), and *Echinochloa crus-galli* (tribe Paniceae) varied intraindividually in the length of the bilobate shape.). In contrast, some species were very conservative in their phytolith shape, like, for example, *Bambusa tuldoides*, *Phragmites australis*, and *Aristida rhinochloa*.

**Fig. 4.**
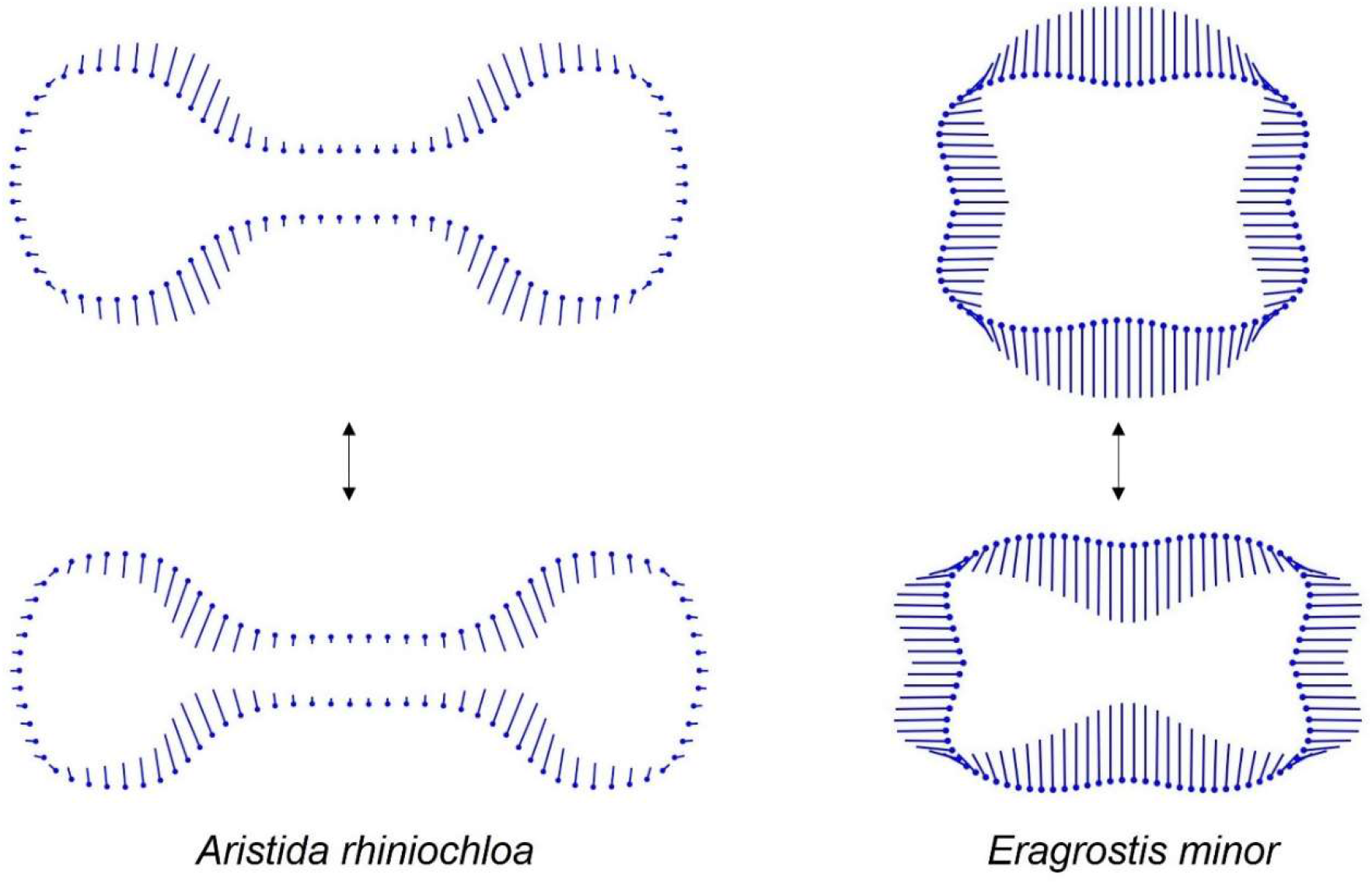
The amount of intraindividual variation in phytolith shape (represented by the average distance of individual phytolith to species centroid) is visualised by ‘lollipop’ graphs where the stick of the lollipop indicates the direction of the variation, whereas the lollipops (filled dots) represent together mean phytolith shape outline. *Aristida rhinochloa* represent species with less variable phytolith shape, whereas *Eragrostis minor* has highly variable phytolith shape. (For the phytolith outlines of more species see Supplemental files Notes S1)

### Mapping phytolith shape data onto the phylogeny

According to the PCA performed on the symmetrised phytolith configurations, the first two PCs explained >96 % of the total variation (Fig. 5). Along PC1 (explaining 89.5 % of the total variation), phytoliths varied between elongated shapes with two deeply incised lobes and shorter and taller shapes on opposite sides. Along PC2 (explaining 7.0 % of the total variation), phytolith shape varied between shorter outlines with two deeply incised lobes and elongated shapes with a pronounced middle part on the opposite sides.

**Fig. 5.**
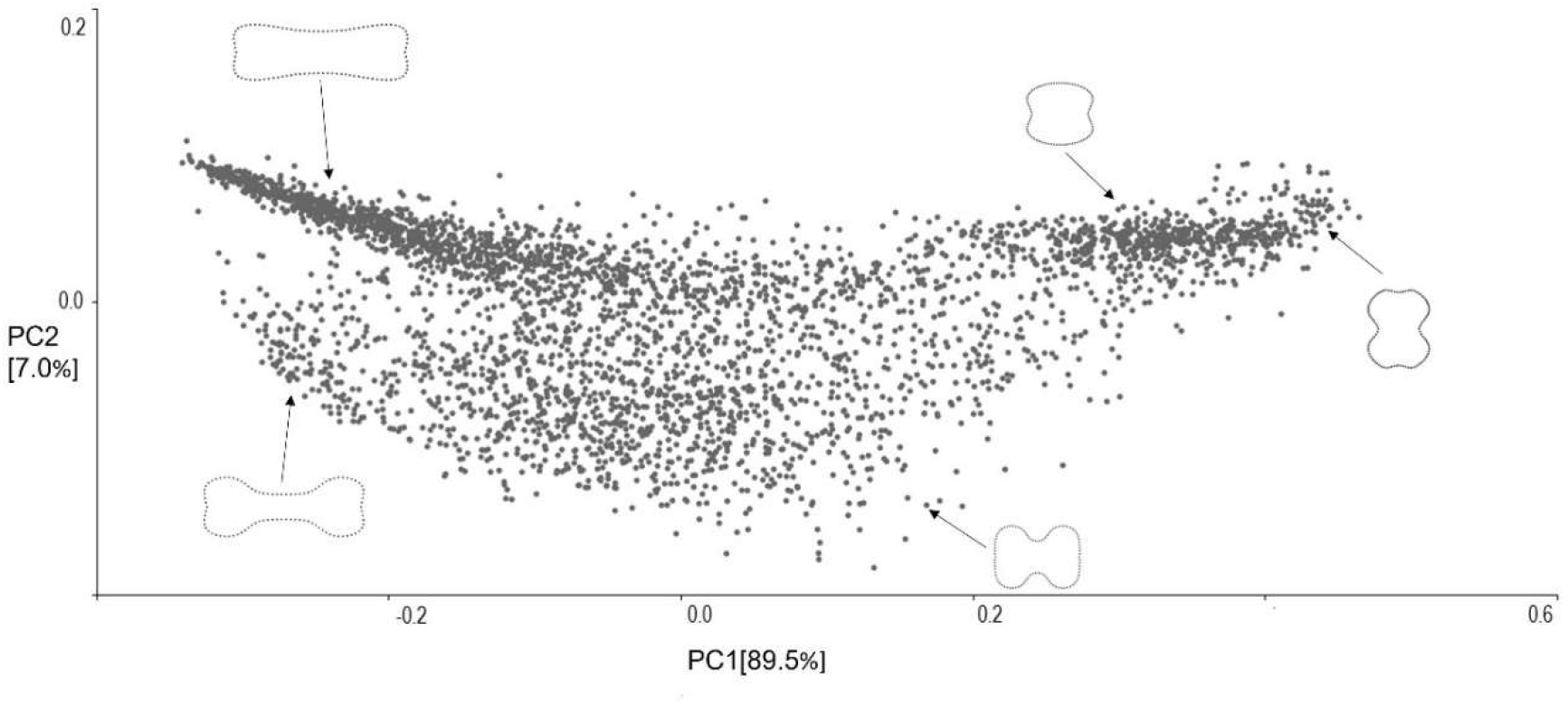
Principal component analysis of 3650 symmetrised grass silica short cell phytolith shape configurations. Dots indicate individual phytoliths. The first two PCs are associated with the most pronounced shape variation within the dataset. Phytolith outlines represent shape variation along both axes.

To visualise the phylogenetic history of symmetric phytolith shape change, we mapped grass species positions along PC1 on the grass phylogeny tree (Fig. 6). The most prominent shape variation in the data — an elongated shape with two deeply incised lobes vs shorter and taller shapes — was strongly phylogenetically conserved at the subfamily level. Thus, individual monophyletic subfamilies substantially differed in their phytolith shapes. On the other hand, similar phytolith shapes occurred in two major grass subclades: BOP and PACMAD. The measure of phylogenetic conservatism for individual PCs describing symmetrised shape variation, along with the measure of phylogenetic conservatism in asymmetric components of shape variation and in the amount of intraindividual variation (Fig. S1), are summarised in Table 2.

**Fig. 6.**
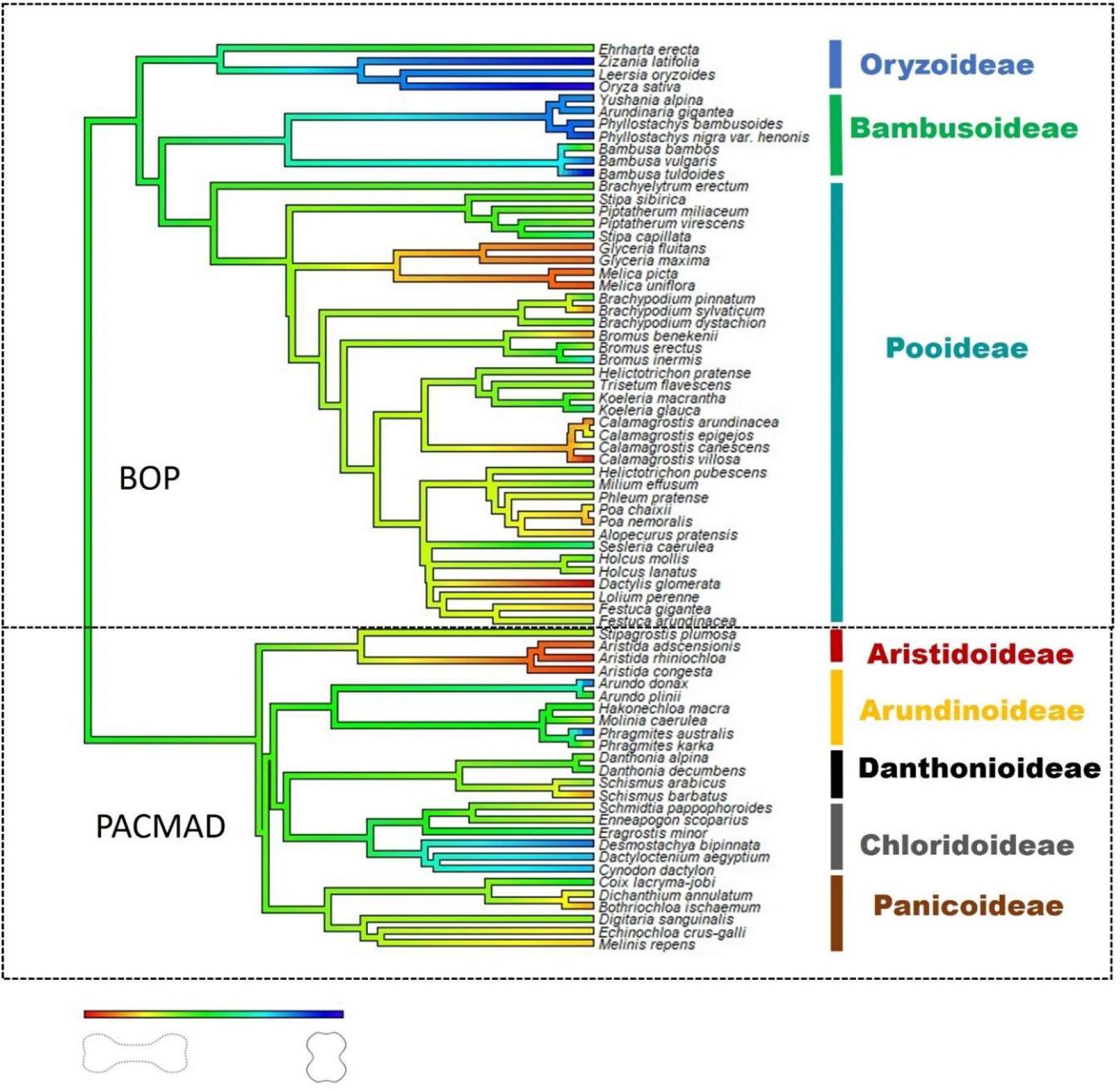
Phytolith shape variation pattern across grass phylogeny. Grass phylogeny was constructed with V.PhyloMaker package in R (Jin & Qian, 2019), ‘backbone’ tree based on molecular data from seed plant phylogeny (mega-tree ‘GBOTB.extended.tre’; Smith & Brown, 2018). Grass species positions along the first principal component axis (PC1) were mapped on the tree. Colour scale indicates grass silica short cell phytolith shape variation along PC1.

**Table 2.**
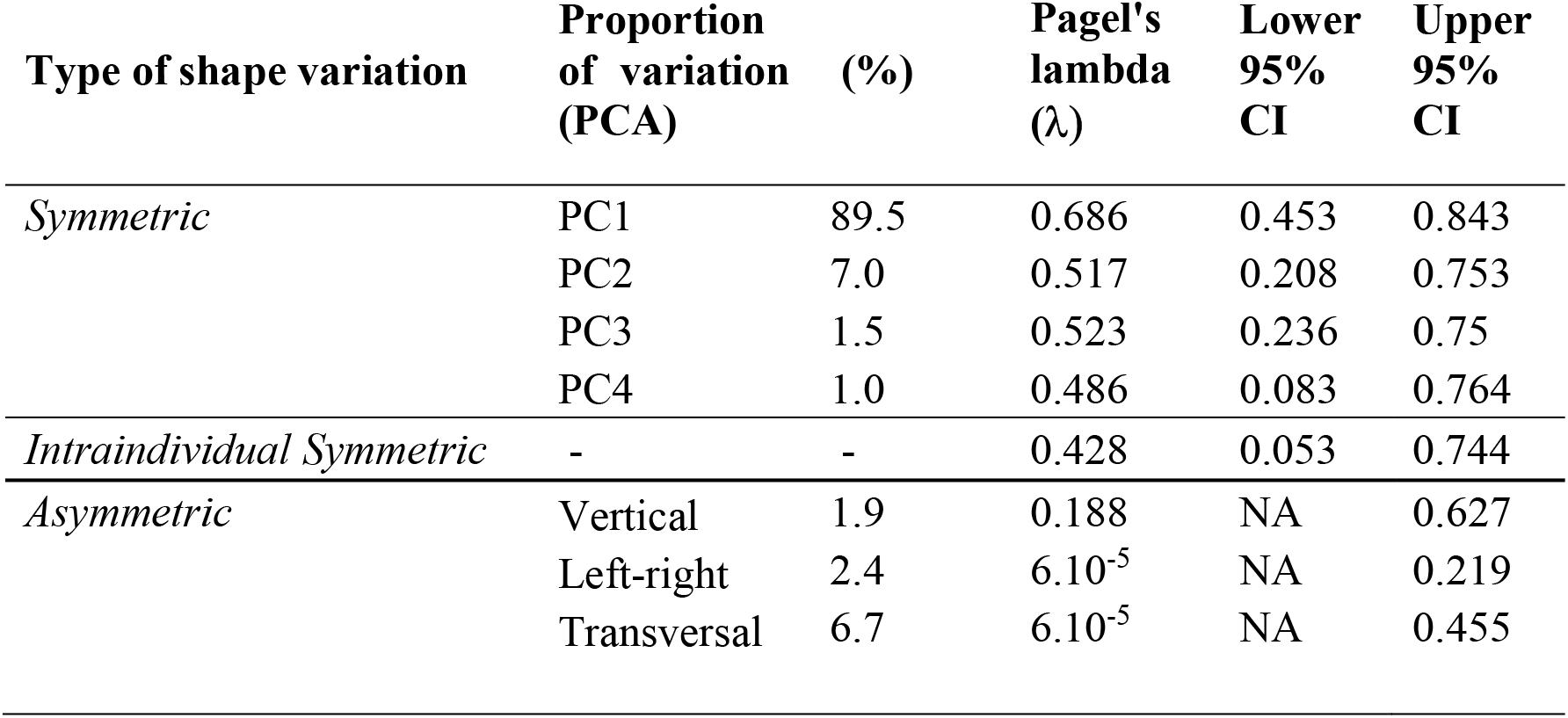
Measure of phylogenetic conservatism in averaged phytolith shape represented by principal components of symmetric (PC1-PC4) and asymmetric shape variation (vertical, left-right, transversal). Phylogenetic conservatism of the amount of intraindividual variation in symmetrised phytolith shape was also measured (see also Fig. S1). The amount of intraindividual variation in phytolith shape in individual grass species is represented by average distance of individual phytolith to species centroid. N=73

| Type of shape variation | Proportion of variation (PCA) | (%) | Pagel's lambda ($\lambda$) | Lower 95% CI | Upper 95% CI |
| --- | --- | --- | --- | --- | --- |
| *Symmetric* | PC1 | 89.5 | 0.686 | 0.453 | 0.843 |
|  | PC2 | 7.0 | 0.517 | 0.208 | 0.753 |
|  | PC3 | 1.5 | 0.523 | 0.236 | 0.75 |
|  | PC4 | 1.0 | 0.486 | 0.083 | 0.764 |
| *Intraindividual Symmetric* | - | - | 0.428 | 0.053 | 0.744 |
| *Asymmetric* | Vertical | 1.9 | 0.188 | NA | 0.627 |
| | Left-right | 2.4 | $6.10^{-5}$ | NA | 0.219 |
| | Transversal | 6.7 | $6.10^{-5}$ | NA | 0.455 |

The phylogenetic tree of the subfamily Pooideae was also projected onto the shape tangent space by squared-change parsimony. The resulting tree was plotted in the plane of the PC1 vs PC2 ordination plot of the species-level mean shapes (Fig. 7). The phylogenetically closely-related taxa were not necessarily close to each other in the morphospace, so there are long branches that criss-cross the plot. On the other hand, a number of other closely-related taxa, especially *Stipa capillata, S. sibirica, Piptatherum miliaceum, P. virescens* in the tribe Stipeae and *Melica picta, M. uniflora, Glyceria fluitans, G. maxima* in the tribe Meliceae showed close correspondence between phylogeny and phytolith shape variation described by the first two PCs.

**Fig. 7.**
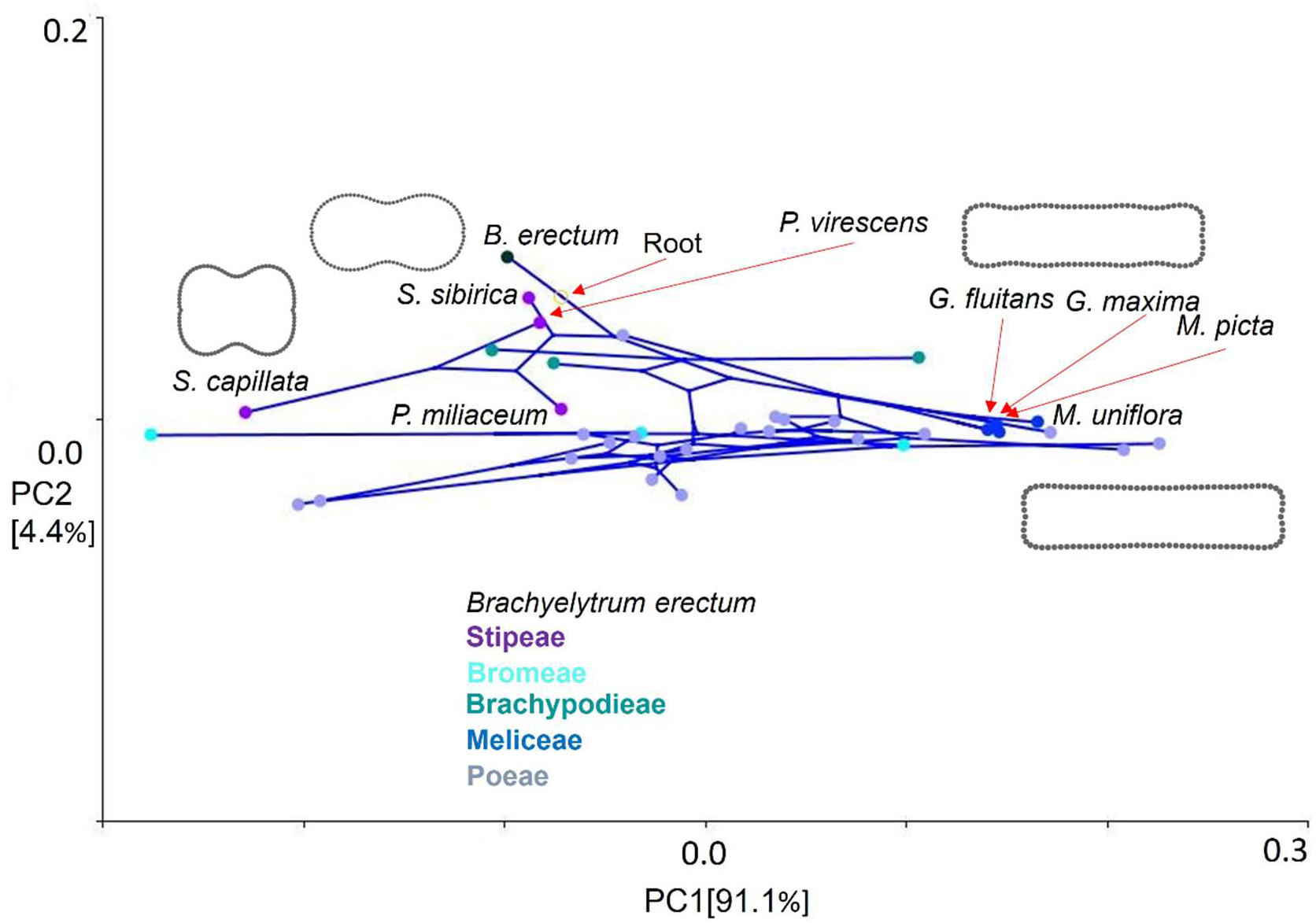
Grass phylogeny mapped onto morphospace represented by the PC1 vs PC2 of the subfamily Pooideae. Coloured dots indicate analysed species from the various tribes; lines represent phylogenetic tree diversification. Closely related species (*Stipa capillata, Stipa sibirica, Piptatherum miliaceum, Piptatherum virescens, Melica picta, Melica uniflora, Glyceria fluitans, Glyceria maxima*) with phytolith shape variation well correlated with relationships in phylogenetic trees and their phytolith shape outlines are indicated.

## Discussion

### The observed evolutionary pattern in phytolith shape

In general, the results of this study concur with previous reports that GSSC phytolith shapes closely reflect phylogenetic relationships between taxa (Piperno & Pearsall, 1998; Gallaher *et al*., 2020). Thus, in the case of Poaceae, this means that phytolith shape changed along with other anatomical and micromorphological leaf traits in response to their diversification in the course of evolutionary history (Thomasson, 1978; Romaschenko, 2011; Rudall *et al*,. 2014). The deep-time diversification of grass subfamilies mainly drives the basic pattern of phytolith shape variation. The ancestors of recent grass species probably carried phytoliths of similar shapes to those we see in extant species occurring in modern ecosystems (Thomasson, 1987; Strömberg, 2005; Prasad *et al*., 2005, 2011).

We found differences between subfamilies known for their distinct phytolith shape (Chloridoideae, Oryzoideae, Bambusoideae, Panicoideae, Pooideae; Twiss *et al*., 1969; Gallaher *et al*., 2020) but also between smaller subfamilies like Aristidoideae, Arundinoideae and Danthonioideae. We also revealed diversification in phytolith shape in some tribes. Axially-oriented bilobate-shaped phytoliths of *Ehrharta erecta* (Ehrharteae) differed from those in Oryzoideae, while *Oryza sativa, Zizania latifolia*, and *Leersia oryzoides* (Oryzeae) formed a separate cluster within this subfamily. Similarly, bilobate-shaped phytoliths of *Eragrostis minor* and *Enneapogon scoparius* (Eragrostideae) differed from those in Chloridoideae. Also, *Stipa* and *Piptatherum* (Stipeae) and *Melica* and *Glyceria* (Meliceae) differed from Pooideae. From this, it is clear that further analysis of phytolith shape employing a more comprehensive range of grass tribes would be useful.

### The intraindividual variation in relation to the phylogenetic pattern in phytolith shape

The amount of intraindividual variation was variable across species. However, the phylogenetic signal in intraindividual variation was generally low. Nevertheless, we detected high intraindividual variation in some early-diverging lineages (as was previously shown by Piperno & Pearsall, 1998). The phytolith shapes of *Eragrostis minor* were the most variable-it carried saddle-shaped phytoliths characteristic of its subfamily Chloridoideae and bilobate-shaped phytoliths generally attributed to Panicoideae. Similarly, polylobate-shaped phytoliths occurred in *Ehrharta*, the lineage diverging early from Oryzoideae, itself characterised by tall, transversally-oriented, bilobate-shaped phytoliths. In Pooideae, the early-diverging lineages (Brachyelytrum, Brachypodieae, Bromeae, and Meliceae) also significantly varied in phytolith shape. However, we also found some highly variable phytolith shape in ‘crown’ groups (e. g. Poeae, Andropogoneae) and very conservative phytolith shape in early-diverging lineages (e. g. Aristideae). Further studies of phylogenetic signals in intraindividual phytolith shape variation are needed to clarify these patterns observed in phytolith shape.

Whether there is a phylogenetic signal in intraindividual phytolith shape variation or not, we still need to consider this variation while classifying fossil phytoliths. In other words, we must consider whether to classify fossil taxa on the basis of comparison with averaged phytolith shape (representing whole intraindividual shape variation by a single shape) or compare them with whole intraindividual phytolith shape variation within the species of our reference collection. According to the findings of this study, the second option seems to be the better approach since the averaged phytolith shape of some species does not reflect the natural variation in phytolith shape (as seen in the extreme case of *Eragrostis minor*). A reference collection based on the whole intraindividual phytolith shape variation of the studied species is necessary.

### The ecological component in phytolith shape variation

Although we found a clear phylogenetic signal in phytolith shape, we consider the high contrast in terms of the phytolith shape (namely the distinction between elongated and shorter shapes) of some closely related taxa as striking. As shown before, grass long cell phytoliths, like crenate and polylobate GSSC phytoliths in some species, are consistently bigger and proportionally longer in reduced light conditions (Dunn *et al*., 2015). It was suggested that the addition of trichomes and stomata to the epidermis might shorten phytoliths, and that the densities of both stomata and trichomes increase with enhanced light conditions in grasses (Allard *et al*., 1991; Knapp & Gilliam, 1985). Focusing on the Pooideae subfamily, we found such a contrast between the Stipeae and Meliceae tribes. Stipeae comprises species that generally occupy drier open grasslands and steppe communities (e. g. Romaschenko *et al*., 2011, 2012). Meliceae is mostly found in shady woodlands or wet environments. More studies focusing on phytolith shape variation along environmental gradients and on comparison with other anatomical traits (e. g. stomata and trichome densities) would be needed in these groups; however, on the basis of what we have learned from this dataset, we expect this shape variation to be more an outcome of the long-term ecological adaptation of species than a short-term plastic response of phytolith shape to the environment. Further testing is crucial in this direction.

### Limitations of the current study

Geometric morphometrics enables the quantitative assessment of phytolith shape variation and the exploration of trends in phytolith shape variation within one morphospace. However, the Procrustes superimposition of phytolith shapes may not be universally applicable to all known morphotypes occurring in real samples. As shown in this study, in phytoliths with symmetric 2D shapes, there may be two or more fixed points delimiting individual symmetric curves. These points are typically derived from the orientation of phytoliths within plant tissue. This applies to wide range of phytolith morphotypes; however, phytoliths that do not bear any such corresponding points, such as those classified into rondel or spheroid groups, would be unsuitable for a clear-cut morphometric analysis by the generalized Procrustes analysis. This is the major limitation of our approach, because it prevents us from including all grass species that carry only such morphotypes, such as the genera *Nardus*, which represents an early-diverging lineage of Pooideae, or *Festuca* (in particular, species with narrow leaves).

Landmark-based geometric morphometrics applied to a 3D phytolith surface model to quantify the overall GSSC phytolith shape partially overcomes this problem (Gallaher *et al*., 2020). 3D shape assessment helps to establish the positional homology of all GSSC phytolith morphotypes, allowing them to be analysed within a common framework. However, the 3D morphometrics of phytoliths is based on confocal microscopy (Evett & Cuthrell, 2016), which makes it more expensive and time-consuming than analysis based on light microscopy. The fact that 2D geometric morphometrics is more accessible allows robust studies based on large datasets. This particularly applies to the use of geometric morphometrics in paleoecology, where high numbers of phytoliths are typically used for the reconstruction of vegetation dynamics (Piperno, 2006; Strömberg, 2009).

### Conclusions

In this study, we demonstrated that the 2D shape of GSSC phytoliths is highly relevant for the reconstruction of the evolution and paleoecology of Poaceae. Although we showed that phytolith shape is mainly driven by the deep-time diversification of grass subfamilies, a closer look also uncovered distinct phytolith shape variation in early-diverging lineages of the subfamily Pooideae. Geometric morphometrics proved to be particularly helpful in this regard. It enables the quantitative assessment of the entire phytolith shape and visualises its variation. Moreover, the geometric morphometrics of 2D phytolith shape is cheap and relatively non-laborious, making it an excellent tool for achieving goals requiring large sums of phytolith outlines, such as in palaeoecological reconstruction.

## Acknowledgements

We thank Tomáš Herben at the Charles University in Prague for his help in every stage of conducting this study, including the study planning, analyses and fruitful comments on the drafts of the paper. We also thank the Herbarium Collections of the Charles University in Prague for providing the grass specimens; we thank Pavel Zdvořák, in particular, who kindly searched for us every grass species we wished for. We thank Matthew Nicholls for his diligent proofreading of the final manuscript.

## Author contributions

KH conceived the study, conducted imagining and data analysis. KH and AP collected plant material in the field. JN developed the scripts for automated analysis of phytolith shape with biradial symmetry and Procrustes ANOVA. KH wrote the text with contribution of JN, PP, and AP. All authors approved the final version of the manuscript.

## Supplemental files

Table S1 List of grass species used for analysis of grass silica short cell phytoliths within *in situ* charred epidermis.

Table S2 Tests of significant difference in phytoliths shape between the subfamilies.

Fig. S1 Pattern in the amount of intraindividual phytolith shape variation across grass phylogeny.

Methods S1 Workflow sequence of landmark-based geometric morphometrics performed on phytoliths with biradial symmetry.

Methods S2 Workflow sequence of landmark-based geometric morphometrics and other methods used in the current study.

Notes S1 Zip file of the ‘lollipop’ graphs visualising the amount of intraindividual variation in phytolith shape for all grass species under study.

