## Supplemental files Tables S1-S2; Fig S1; Methods S1-S2; for "Phylogenetic, ecological and intraindividual variability patterns in grass phytolith shape"

Table S1	List of grass species used for analysis of grass silica short cell phytoliths within *in situ* charred epidermis.

| **Subfamily** | **Tribe** | **Species** | **Provenience** |
| --- | --- | --- | --- |
| **B**ambusoideae | Arundinarieae | *Arundinaria gigantea* (Walter) Muhl. | Herbarium Collections of the Charles University, Prague (CZ) |
|  | Arundinarieae | *Phyllostachys bambusoides* Siebold & Zucc. | Herbarium Collections of the Charles University, Prague (CZ) |
|  | Arundinarieae | *Phyllostachys nigra var. henonis* (Mitford) Rendle | Herbarium Collections of the Charles University, Prague (CZ) |
|  | Arundinarieae | *Yushania alpine* (K.Schum.) W.C.Lin | Herbarium Collections of the Charles University, Prague (CZ) |
|  | Bambuseae | *Bambusa bambos* (L.) Voss | Herbarium Collections of the Charles University, Prague (CZ) |
|  | Bambuseae | *Bambusa tuldoides* Munro | Herbarium Collections of the Charles University, Prague (CZ) |
|  | Bambuseae | *Bambusa vulgaris* Nees | Herbarium Collections of the Charles University, Prague (CZ) |
| **O**ryzoideae | Ehrharteae | *Ehrharta erecta* Lam. | Herbarium Collections of the Charles University, Prague (CZ) |
|  | Oryzeae | *Leersia oryzoides* (L.) Sw. | Herbarium Collections of the Charles University, Prague (CZ) |
|  | Oryzeae | *Oryza sativa* L. | Herbarium Collections of the Charles University, Prague (CZ) |
|  | Oryzeae | *Zizania latifolia* (Griseb.) Turcz. ex Stapf | Herbarium Collections of the Charles University, Prague (CZ) |
| **P**ooideae | Brachyelytreae | *Brachyelytrum erectum* (Schreb.) P.Beauv. | Herbarium Collections of the Charles University, Prague (CZ) |
|  | Brachypodieae | *Brachypodium pinnatum* (L.) P.Beauv. | Český kras (CZ) |
|  | Brachypodieae | *Brachypodium dystachion* (Pers.) P.Beauv. | Herbarium Collections of the Charles University, Prague (CZ) |
|  | Brachypodieae | *Brachypodium sylvaticum* (Huds.) P.Beauv. | Kersko (CZ) |
|  | Bromeae | *Bromus benekenii* Beck | České středohoří (CZ) |
|  | Bromeae | *Bromus erectus* Ledeb. | Český kras (CZ) |
|  | Bromeae | *Bromus inermis* Steven | Vysočina (CZ) |
|  | Meliceae | *Glyceria fluitans* (L.) R.Br. | České Švýcarsko (CZ) |
|  | Meliceae | *Glyceria maxima* (Hartm.) Holmb. | České Švýcarsko (CZ) |
|  | Meliceae | *Melica picta* K.Koch | Český kras (CZ) |
|  | Meliceae | *Melica uniflora* Retz. | České Švýcarsko (CZ) |
|  | Poeae | *Alopecurus pratensis* Bourg. ex Lange | Hrubý Jeseník (CZ) |
|  | Poeae | *Calamagrostis arundinacea* Wibel | České Švýcarsko (CZ) |
|  | Poeae | *Calamagrostis canescens* (Weber) Roth | České Švýcarsko (CZ) |
|  | Poeae | *Calamagrostis epigejos* (L.) Roth | České středohoří (CZ) |
|  | Poeae | *Calamagrostis villosa* St.-Lag. | České Švýcarsko (CZ) |
|  | Poeae | *Dactylis glomerata* L. | České Švýcarsko (CZ) |
|  | Poeae | *Festuca arundinacea* Lilj. | České Švýcarsko (CZ) |
|  | Poeae | *Festuca gigantea* Krock. | Poodří (CZ) |
|  | Poeae | *Helictotrichon pratense* (L.) Pilg. | Botanical Garden of the Charles University, Prague (CZ) |
|  | Poeae | *Helictotrichon pubescens* (Huds.) Schult. & Schult.f. | Botanical Garden of the Charles University, Prague (CZ) |
|  | Poeae | *Holcus lanatus* L. | České Švýcarsko (CZ) |
|  | Poeae | *Holcus mollis* L. | Botanical Garden of the Charles University, Prague (CZ) |
|  | Poeae | *Koeleria glauca* (Spreng.) DC. | Botanical Garden of the Charles University, Prague (CZ) |
|  | Poeae | *Koeleria macrantha* (Ledeb.) Schult. | České středohoří (CZ) |
|  | Poeae | *Lolium perenne* L. | Botanical Garden of the Charles University, Prague (CZ) |
|  | Poeae | *Milium effusum* Lour. | Botanical Garden of the Charles University, Prague (CZ) |
|  | Poeae | *Phleum pratense* L. | Džbán (CZ) |
|  | Poeae | *Poa chaixii* Vill. | Hrubý Jeseník (CZ) |
|  | Poeae | *Poa nemoralis* L. | České středohoří (CZ) |
|  | Poeae | *Sesleria caerulea* (L.) Ard. | České středohoří (CZ) |
|  | Poeae | *Trisetum flavescens* (L.) P.Beauv. | Český kras (CZ) |
|  | Stipeae | *Piptatherum miliaceum* (L.) Coss. | Botanical Garden of the Charles University, Prague (CZ) |
|  | Stipeae | *Piptatherum virescens* (Trin.) Boiss. | Botanical Garden of the Charles University, Prague (CZ) |
|  | Stipeae | *Stipa capillata* L. | České středohoří (CZ) |
|  | Stipeae | *Stipa sibirica* (L.) Lam. | Herbarium Collections of the Charles University, Prague (CZ) |
| **P**anicoideae | Andropogoneae | *Bothriochloa ischaemum* (L.) Keng | Český kras (CZ) |
|  | Andropogoneae | *Coix lacryma-jobi* L. | Botanical Garden of the Charles University, Prague (CZ) |
|  | Andropogoneae | *Dichanthium annulatum* (Forssk.) Stapf | Jebel Sabaloka (SU) |
|  | Paniceae | *Digitaria sanguinalis* (L.) Scop. | Botanical Garden of the Charles University, Prague (CZ) |
|  | Paniceae | *Echinochloa crus-galli* (L.) P.Beauv. | Botanical Garden of the Charles University, Prague (CZ) |
|  | Paniceae | *Melinis repens* (Willd.) Zizka | Botanical Garden of the Charles University, Prague (CZ) |
| **A**ristidoideae | Aristideae | *Aristida adscensionis* L. | Jebel Sabaloka (SU) |
|  | Aristideae | *Aristida congesta* Roem. & Schult. | Makvekve (BW) |
|  | Aristideae | *Aristida rhiniochloa* Hochst. | Sijarira (ZW) |
|  | Aristideae | *Stipagrostis plumosa* Munro ex T.Anderson | Jebel Sabaloka (SU) |
| **C**hloridoideae | Cynodonteae | *Cynodon dactylon* (L.) Pers. | Jebel Sabaloka (SU) |
|  | Cynodonteae | *Dactyloctenium aegyptium* (L.) Willd. | Jebel Sabaloka (SU) |
|  | Cynodonteae | *Desmostachya bipinnata* (L.) Stapf | Jebel Sabaloka (SU) |
|  | Eragrostideae | *Enneapogon scoparius* Stapf | Tsodillo hills (BW) |
|  | Eragrostideae | *Eragrostis minor* Host | Herbarium Collections of the Charles University, Prague (CZ) |
|  | Eragrostideae | *Schmidtia pappophoroides* Steud. ex J.A.Schmidt | Tsodillo hills (BW) |
| **A**rundinoideae | Arundineae | *Arundo donax* Forssk. | Herbarium Collections of the Charles University, Prague (CZ) |
|  | Arundineae | *Arundo plinii* Turra | Herbarium Collections of the Charles University, Prague (CZ) |
|  | Molinieae | *Hakonechloa macra* (Munro) Honda | Herbarium Collections of the Charles University, Prague (CZ) |
|  | Molinieae | *Molinia caerulea* (L.) Moench | Herbarium Collections of the Charles University, Prague (CZ) |
|  | Molinieae | *Phragmites australis* (Cav.) Trin. ex Steud. | Hrubý Jeseník (CZ) |
|  | Molinieae | *Phragmites karka* (Retz.) Trin. ex Steud. | Herbarium Collections of the Charles University, Prague (CZ) |
| **D**anthonioideae | Danthonieae | *Danthonia alpina* Buchanan | Herbarium Collections of the Charles University, Prague (CZ) |
|  | Danthonieae | *Danthonia decumbens* (L.) DC. | České Švýcarsko (CZ) |
|  | Danthonieae | *Schismus arabicus* Nees | Herbarium Collections of the Charles University, Prague (CZ) |
|  | Danthonieae | *Schismus barbatus* (L.) Thell. | Jebel Sabaloka (SU) |

Table S2 Tests of significant difference in phytoliths shape between subfamilies. d = distance between group means; UCL (95%) = pairwise 95% upper confidence limits between means; Z= pairwise effect sizes between means; Pr > d = pairwise P values between means.

| **Pairwise distances between means plus statistics** | **d** | **UCL-95%** | **Z** | **Pr>d** |
| --- | --- | --- | --- | --- |
| Aristidoideae:Arundinoideae | 0.330 | 0.041 | 21.059 | 0.001 |
| Aristidoideae:Bambusoideae | 0.514 | 0.039 | 26.220 | 0.001 |
| Aristidoideae:Danthonioideae | 0.168 | 0.044 | 11.532 | 0.001 |
| Aristidoideae:Chloridoideae | 0.397 | 0.040 | 23.405 | 0.001 |
| Aristidoideae:Oryzoideae | 0.488 | 0.045 | 24.616 | 0.001 |
| Aristidoideae:Panicoideae | 0.148 | 0.039 | 11.177 | 0.001 |
| Aristidoideae:Pooideae | 0.151 | 0.033 | 13.424 | 0.001 |
| Arundinoideae:Bambusoideae | 0.199 | 0.034 | 16.551 | 0.001 |
| Arundinoideae:Danthonioideae | 0.188 | 0.039 | 14.058 | 0.001 |
| Arundinoideae:Chloridoideae | 0.089 | 0.037 | 7.025 | 0.001 |
| Arundinoideae:Oryzoideae | 0.191 | 0.041 | 13.948 | 0.001 |
| Arundinoideae:Panicoideae | 0.189 | 0.036 | 15.490 | 0.001 |
| Arundinoideae:Pooideae | 0.218 | 0.028 | 20.857 | 0.001 |
| Bambusoideae:Danthonioideae | 0.385 | 0.040 | 23.053 | 0.001 |
| Bambusoideae:Chloridoideae | 0.132 | 0.036 | 10.969 | 0.001 |
| Bambusoideae:Oryzoideae | 0.063 | 0.039 | 4.231 | 0.001 |
| Bambusoideae:Panicoideae | 0.379 | 0.035 | 24.881 | 0.001 |
| Bambusoideae:Pooideae | 0.393 | 0.025 | 27.689 | 0.001 |
| Danthonioideae:Chloridoideae | 0.267 | 0.043 | 17.688 | 0.001 |
| Danthonioideae:Oryzoideae | 0.371 | 0.044 | 21.314 | 0.001 |
| Danthonioideae:Panicoideae | 0.065 | 0.042 | 4.147 | 0.002 |
| Danthonioideae:Pooideae | 0.123 | 0.034 | 10.932 | 0.001 |
| Chloridoideae:Oryzoideae | 0.135 | 0.040 | 10.149 | 0.001 |
| Chloridoideae:Panicoideae | 0.256 | 0.036 | 19.414 | 0.001 |
| Chloridoideae:Pooideae | 0.269 | 0.028 | 23.132 | 0.001 |
| Oryzoideae:Panicoideae | 0.361 | 0.041 | 22.232 | 0.001 |
| Oryzoideae:Pooideae | 0.374 | 0.034 | 24.720 | 0.001 |
| Panicoideae:Pooideae | 0.069 | 0.027 | 7.500 | 0.001 |

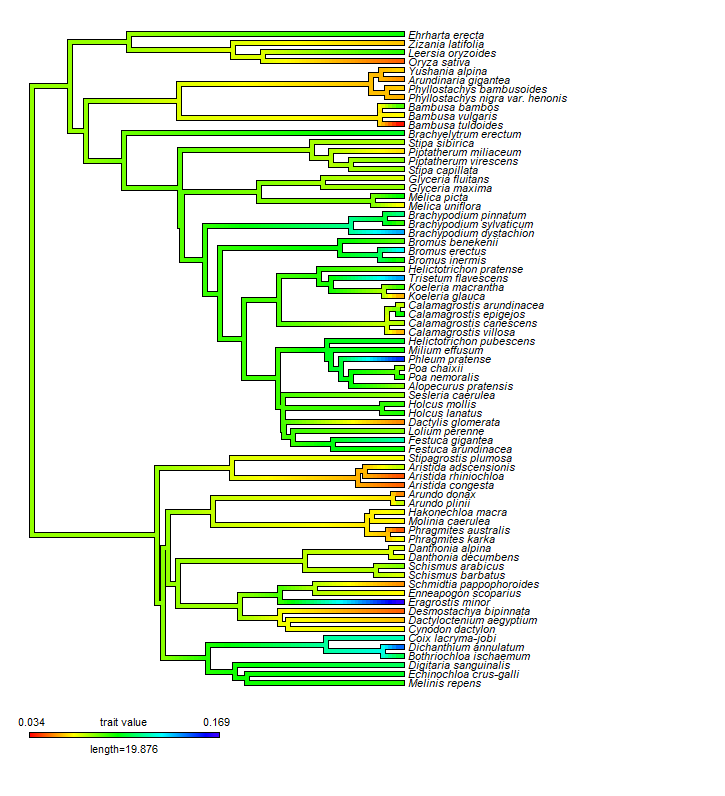

Fig. S1 Pattern in the amount of intraindividual phytolith shape variation across grass phylogeny. Grass phylogeny was constructed with the V.PhyloMaker package in R (Jin & Qian 2019); ‘backbone’ tree based on molecular data from seed plant phylogeny (mega-tree ‘GBOTB.extended.tre’; Smith & Brown, 2018). The amount of intraindividual variation in phytolith shape in individual grass species is represented by average distance of individual phytolith to species centroid. Colour scale visualises this variation.

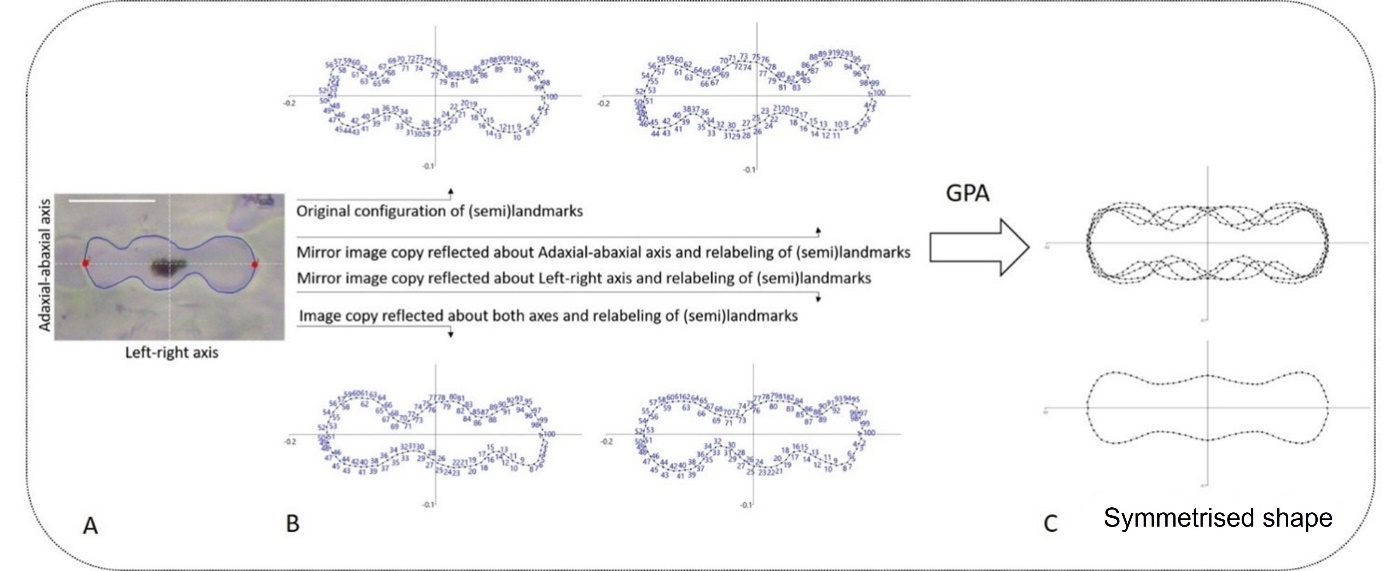

Methods S1 Workflow sequence of landmark-based geometric morphometrics performed on phytoliths with biradial symmetry (modified after Hošková *et al*. 2021). (A) Digitation of landmarks. (B) Reflection and relabelling of semilandmarks. Relabelling is required to ensure that original configuration and transformed copies match. (C) Generalized Procrustes Analysis (GPA) performed

on the multiplied data set. The symmetrised configurations (obtained by averaging of original and transformed configurations) are invariant under all transformations. Scale bar = 20 micron

Methods S2	Workflow sequence of landmark-based geometric morphometric and other methods used in the current study.

| **Processes** | **Software (hardware)** | **References** | **Output** |
| --- | --- | --- | --- |
| Microphotograph acquisition (Fig. 1) | Leica microscope DM1000LED  Leica camera ICC50 W |  | sequential images of rows of GSSC phytoliths |
| Digitation – the placement of 2 fixed landmarks and 98 semilandmarks per phytolith (Fig. 1, Methods S1) | semi-automated background curves tool in TpsDig, ver. 2.31 | Rohlf (2015)  <https://life.bio.sunysb.edu/morph/> | phytolith semilandmarks coordinates (in .TPS format) |
| Semilandmarks equidistantization – the one halve of the phytolith outline first, and then the another; merging of the phytolith outline halves | *digit.curves* function in the *geomorph* package, ver. 3.3.2 in R, ver. 3.6.3., RStudio, ver. 1.2.5033 | Bookstein (1997); Gunz & Mitteroecker (2013); Adams *et al*. (2021) | matrix of equidistant positions of semilandmarks along the outlines relative to the positions of the fixed landmarks |
| Generalised Procrustes analysis (GPA) performed on equidistant semilandmarks. GPA minimises the sum of squared distances between corresponding semilandmarks to extract shape data by removing the extraneous information of size, location and orientation | *arrayspec* and *gpagen* function in the *geomorph* package, ver. 3.3.2. in R, ver. 3.6.3., RStudio, ver. 1.2.5033 | e. g., Zelditch *et al*. (2012); Dryden &Mardia (2016); Adams et al. (2021) | matrix of superimposed Procrustes coordinates of semilandmarks |
| Reflection and relabelling of semilandmarks. Relabelling is required to ensure that original configuration and transformed copies match. (Methods S1) | R, ver. 3.6.3., RStudio, ver. 1.2.5033 | Savriama (2018) | multiplied dataset of Procrustes coordinates  – 4 transformed re-labelled copies per configuration: original configuration of semilandmarks, a reflected copy about the horizontal adaxial-abaxial axis; a reflected copy about the vertical left-right axis and copy reflected about both axes |
| GPA performed on multiplied dataset (Methods S1) | *procGPA* function in *shapes* package, ver. 1.2.5 in R, ver. 3.6.3., RStudio, ver. 1.2.5033 | Dryden (2019) | matrix of superimposed Procrustes coordinates of semilandmarks from multiplied dataset  – the scores of configurations on principal components (PCs)  – the symmetrised phytolith shape configurations (obtained by averaging of original and transformed configurations) which are invariant under all transformation |
| Principal component analysis (PCA) performed on multiplied dataset to separate phytolith shape variation into component of symmetric variation and 3 components of asymmetry | R, ver. 3.6.3., RStudio, ver. 1.2.5033 | Savriama & Klingenberg (2011); Savriama *et al*. (2012); Savriama (2018) | PCs associated with shape change which corresponds with type of symmetry or asymmetry=quantification of proportion of symmetric and asymmetric variation within dataset |
| Multivariate *Procrustes* ANOVA – separation and quantification of sources of phytolith shape variation performed on consensus phytolith shapes | R, ver. 3.6.3., RStudio, ver. 1.2.5033 | Klingenberg et al. (2002); Savriama *et al*. (2012); Neustupa (2013); Neustupa & Woodard (2021) | proportion of variation in symmetrised phytolith shape configurations explained by different sources of variation: relation to ‘BOP vs. PACMAD’ clades, subfamilies, tribes, genera, species (Fig. 2) |
| Pairwise randomised residual permutation procedure *posthoc* tests | *‘pairwise’* function in RRPP package in  R, ver. 3.6.3., RStudio, ver. 1.2.5033 | Collyer & Adams (2018) |  |
| Canonical variates analysis (CVA) performed on symmetrised phytolith configurations | MorphoJ | Klingenberg (2011) | scores of phytoliths on CVs  scatterplot used to discriminate between symmetrised phytolith shape of grass subfamilies (Fig. 3) |
| Comparison of Procrustes distances of individual phytoliths to species centroids for all species under study | ‘*betadisper*’ function in package VEGAN in R, ver. 3.6.3., RStudio, ver. 1.2.5033 | Oksanen *et al*. (2019) | amount of intraindividual variation in phytolith shape for individual grass species (Fig. 4, Notes S1) |
| Principal component analysis (PCA) performed on symmetrised phytolith shape configurations | MorphoJ | Klingenberg (2011) | PCA scatterplot of variation in symmetrised phytolith shape configurations and visualisation of this shape variation (Fig. 5) |
| Construction of grass phylogeny (one phylogeny generated using S3 scenario) of species under study using ‘backbone’ tree based on molecular data from seed plant phylogeny (mega-tree ‘GBOTB.extended.tre’) | ‘*phylo.maker*’ function in V.PhyloMaker package in R, ver. 3.6.3., RStudio, ver. 1.2.5033 | Jin and Qian (2019); Smith & Brown (2018) |  |
| Mapping of symmetrised phytolith shape variation represented by PCs, PCs of asymmetrical component of shape variation, intraindividual shape variation represented by represented by averaged distances from species group centroid in multivariate space onto phylogenetic tree; measuring of phylogenetic conservatism of above mentioned traits, calculation of confidence intervals | ‘pgls’ in CAPER package, functions ‘contmap’, ‘phylosig’, ‘fastAnc’ in PHYTOOLS package in R, ver. 3.6.3., RStudio, ver. 1.2.5033 | Orme *et al*. (2018); Revell (2012) | symmetrised phytolith shape variation, phytolith shape asymmetrical component, intraindividual symmetrised phytolith shape variation pattern across the grass phylogeny and measure of phylogenetic conservatism (Fig. 6; Fig. S1; Table 2) |
| Projection of phylogenetic tree of species under study onto the shape tangent space by squared-change parsimony | MorphoJ | Klingenberg (2011) | phylogenetic tree of species under study plotted in the plane of the PC1 vs. PC2 and PC3 vs. PC4 of symmetrised phytolith shape variation (Fig. 7) |
