## Supplementary figures and images for "Phylogenetic, ecological and intraindividual variability patterns in grass phytolith shape"

### AruGig-0.1.jpg

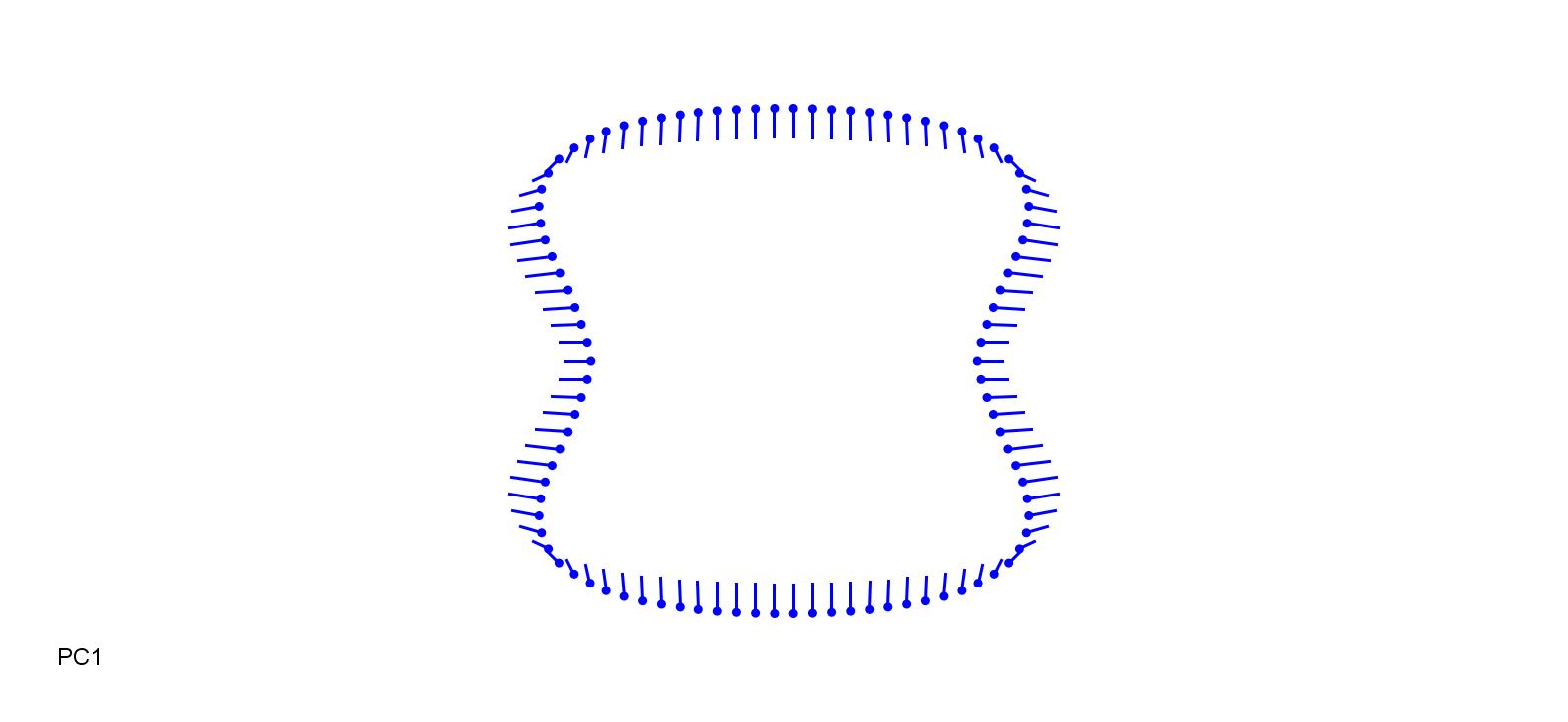

### AruGig_0.1.jpg

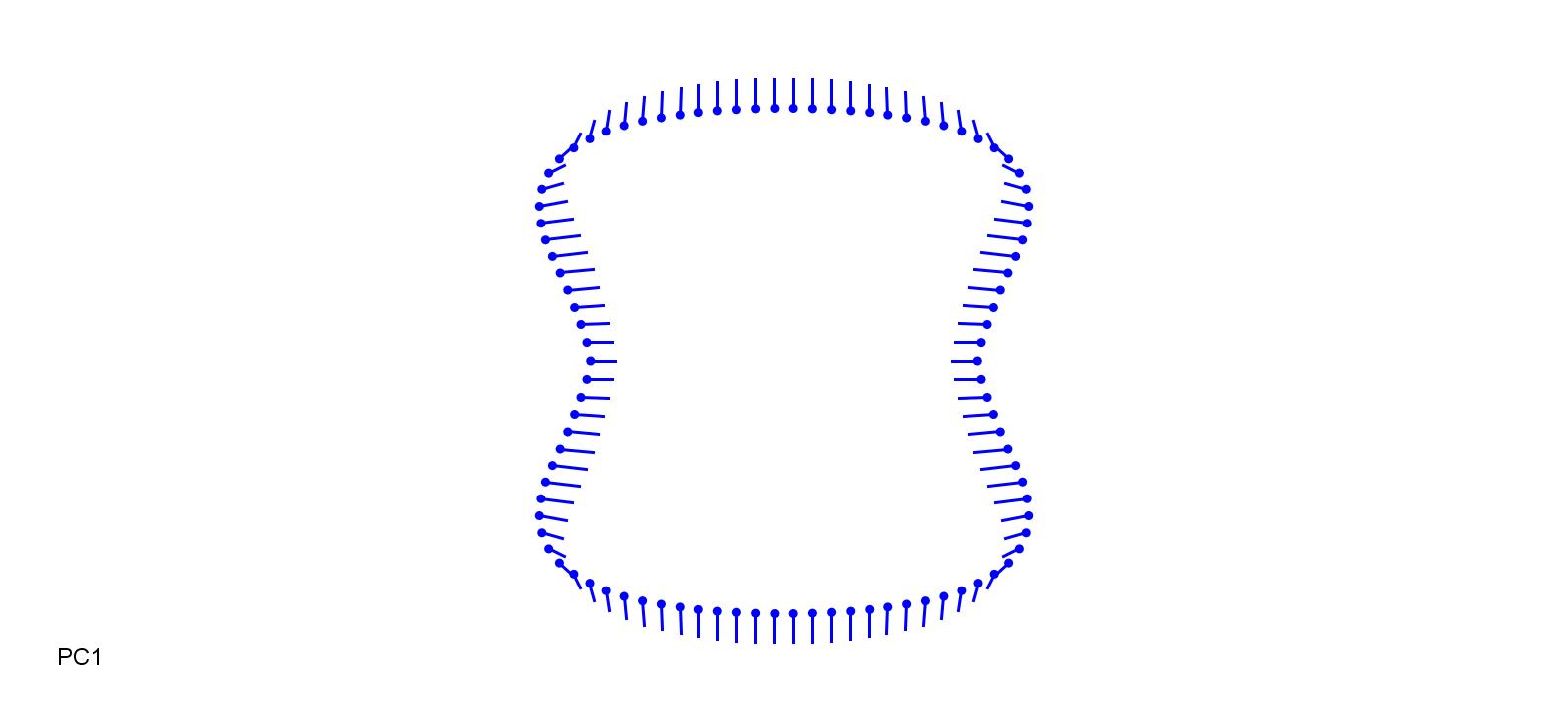

### BamBam-0.15.jpg

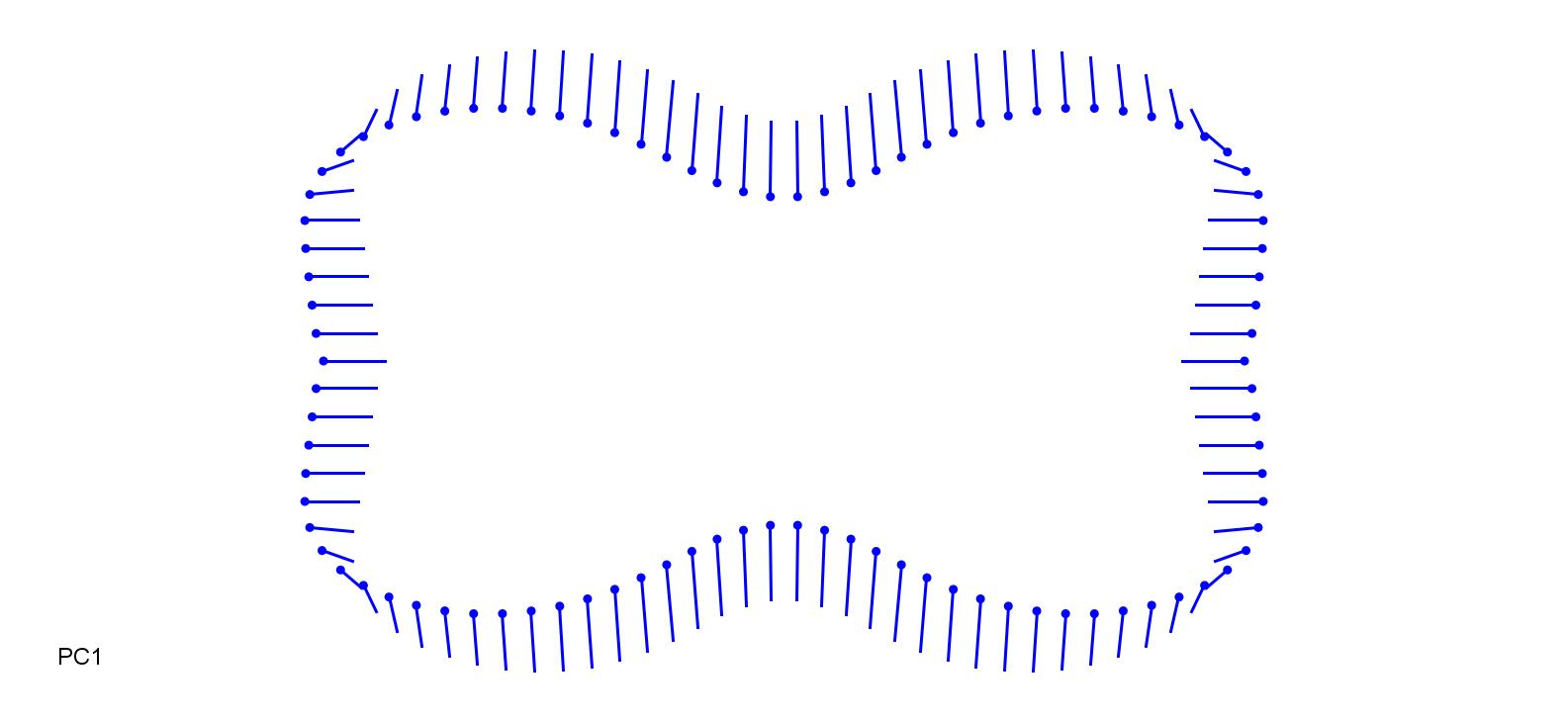

### BamBam_0.15.jpg

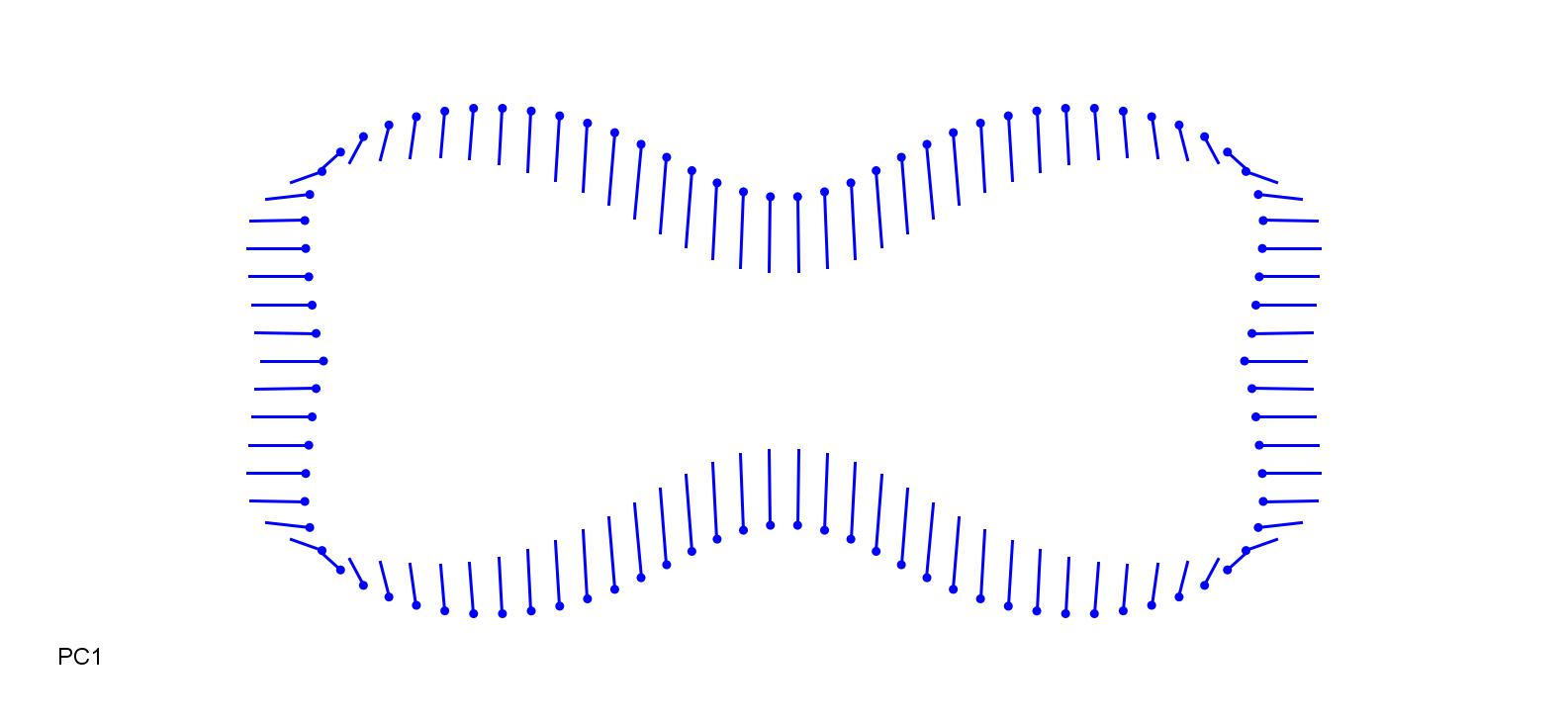

### BamTul-0.05.jpg

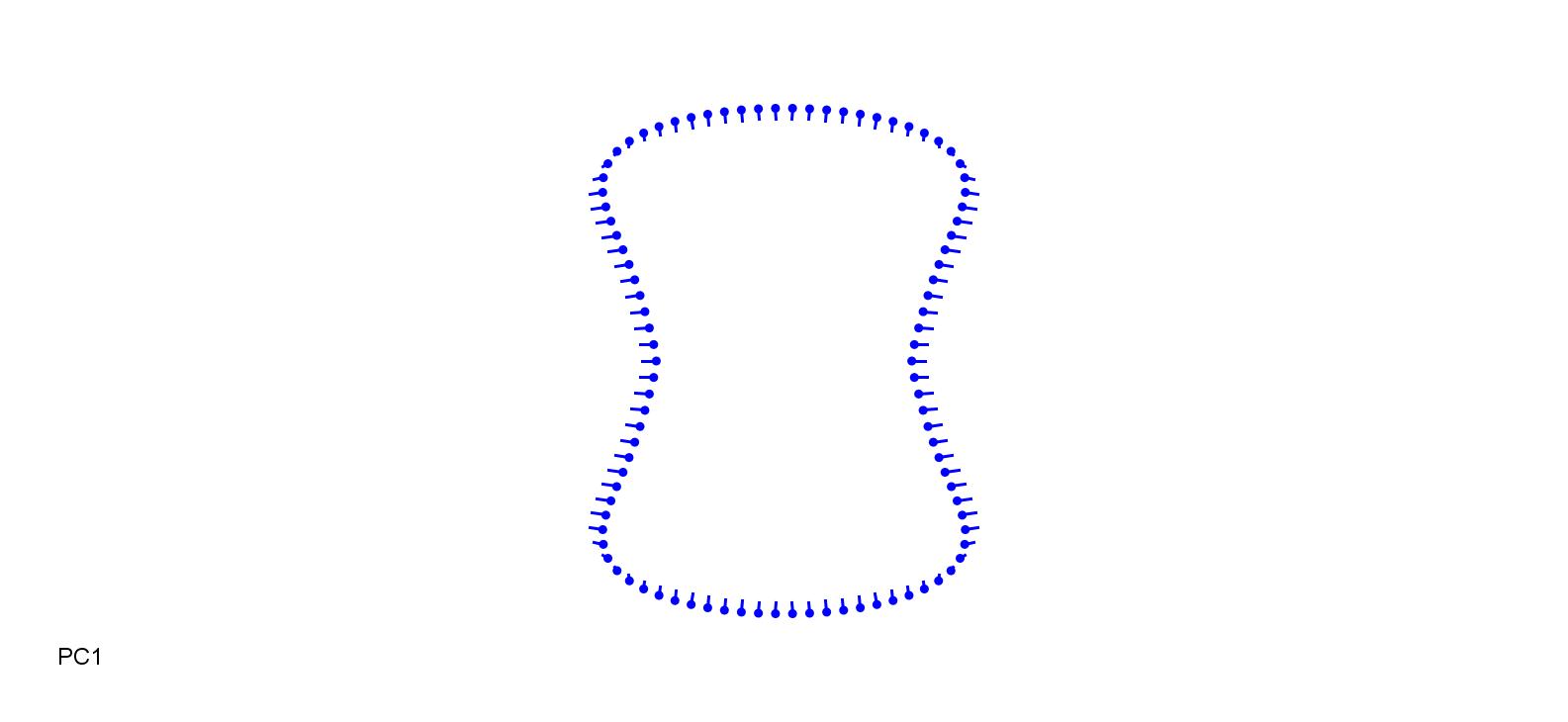

### BamTul_0.1.jpg

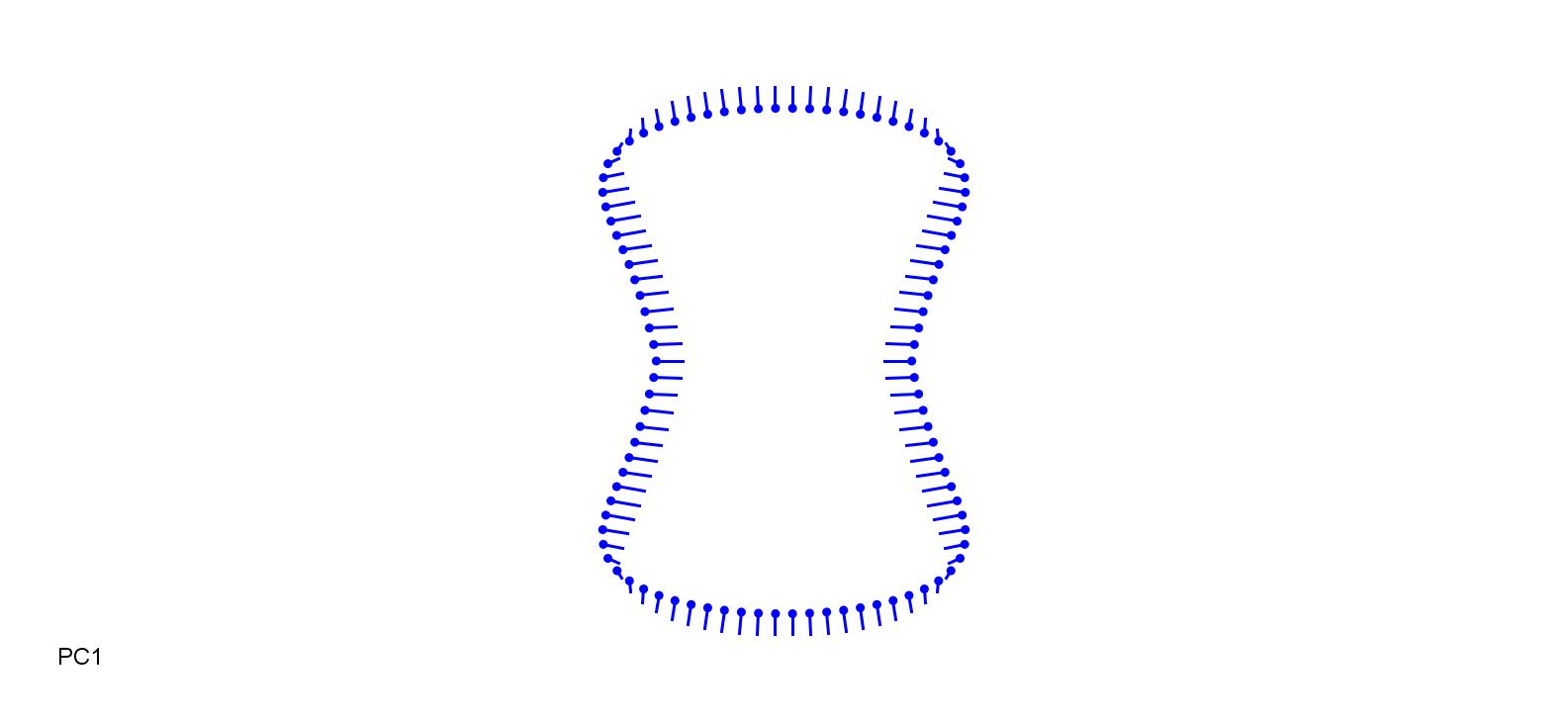

### BamVul-0.18.jpg

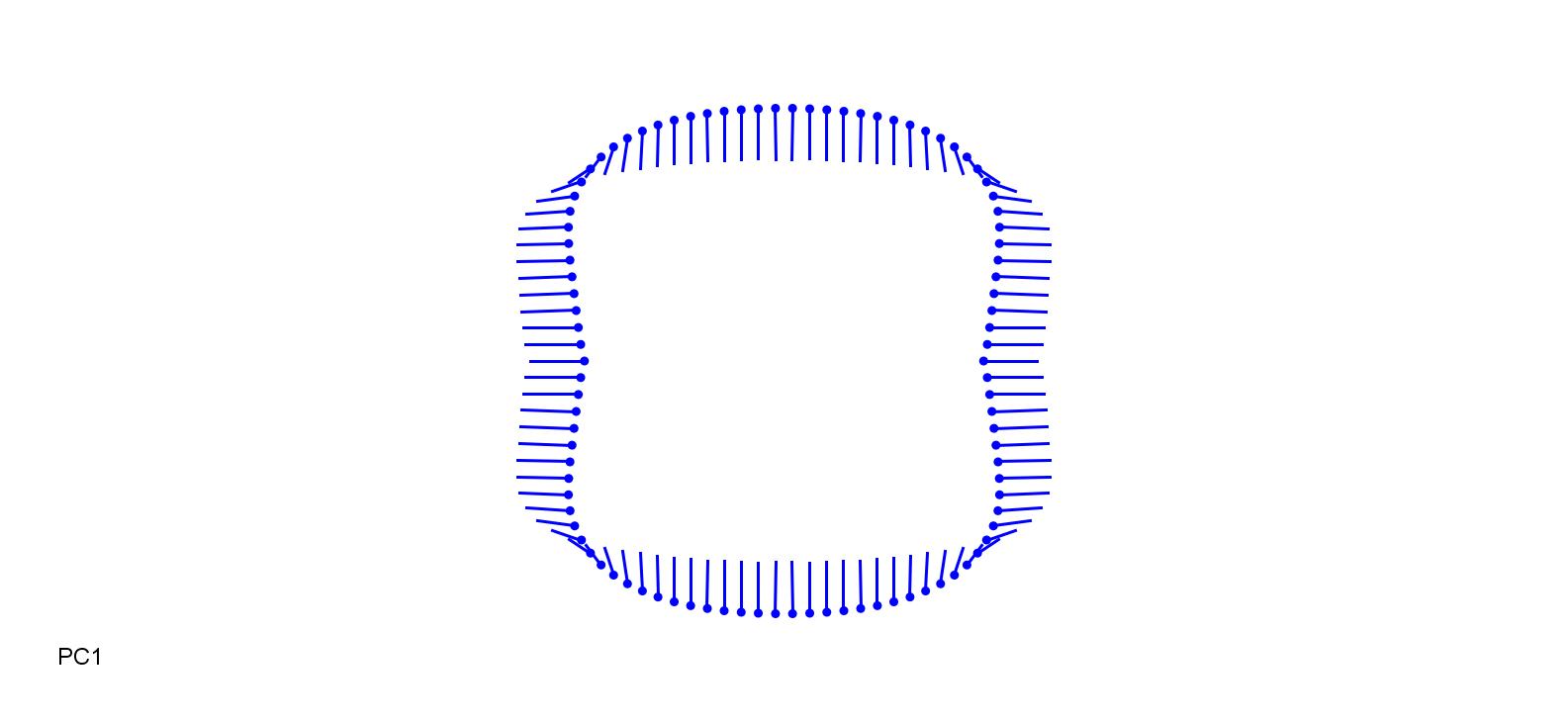

### BamVul_0.18.jpg

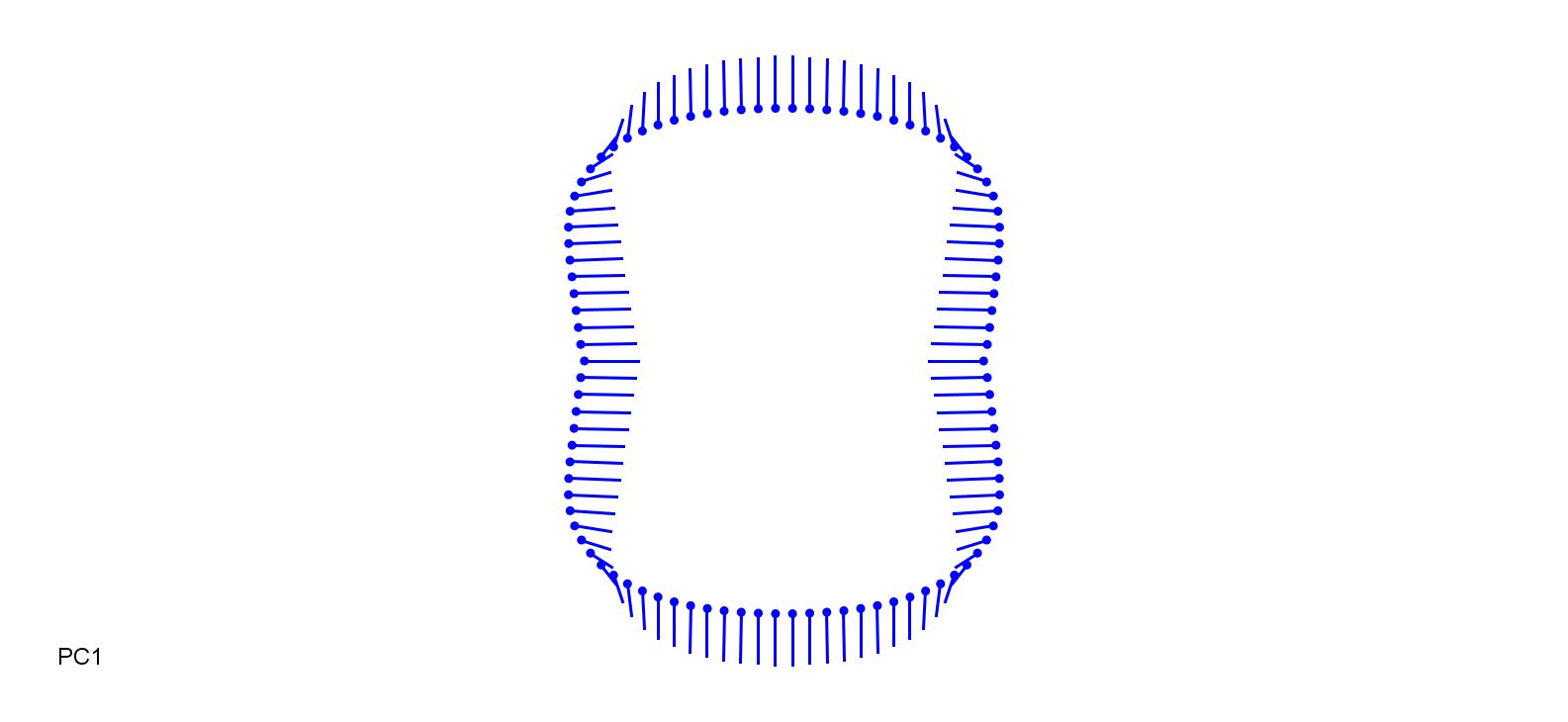

### BraDys-0.3.jpg

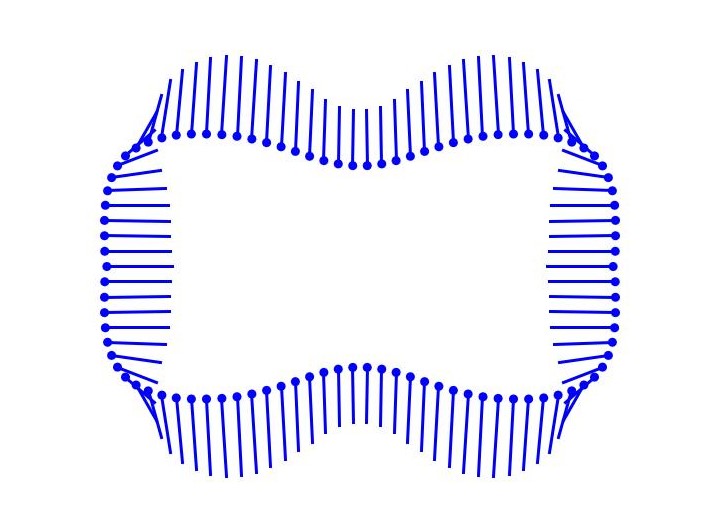

### BraDys_0.3.jpg

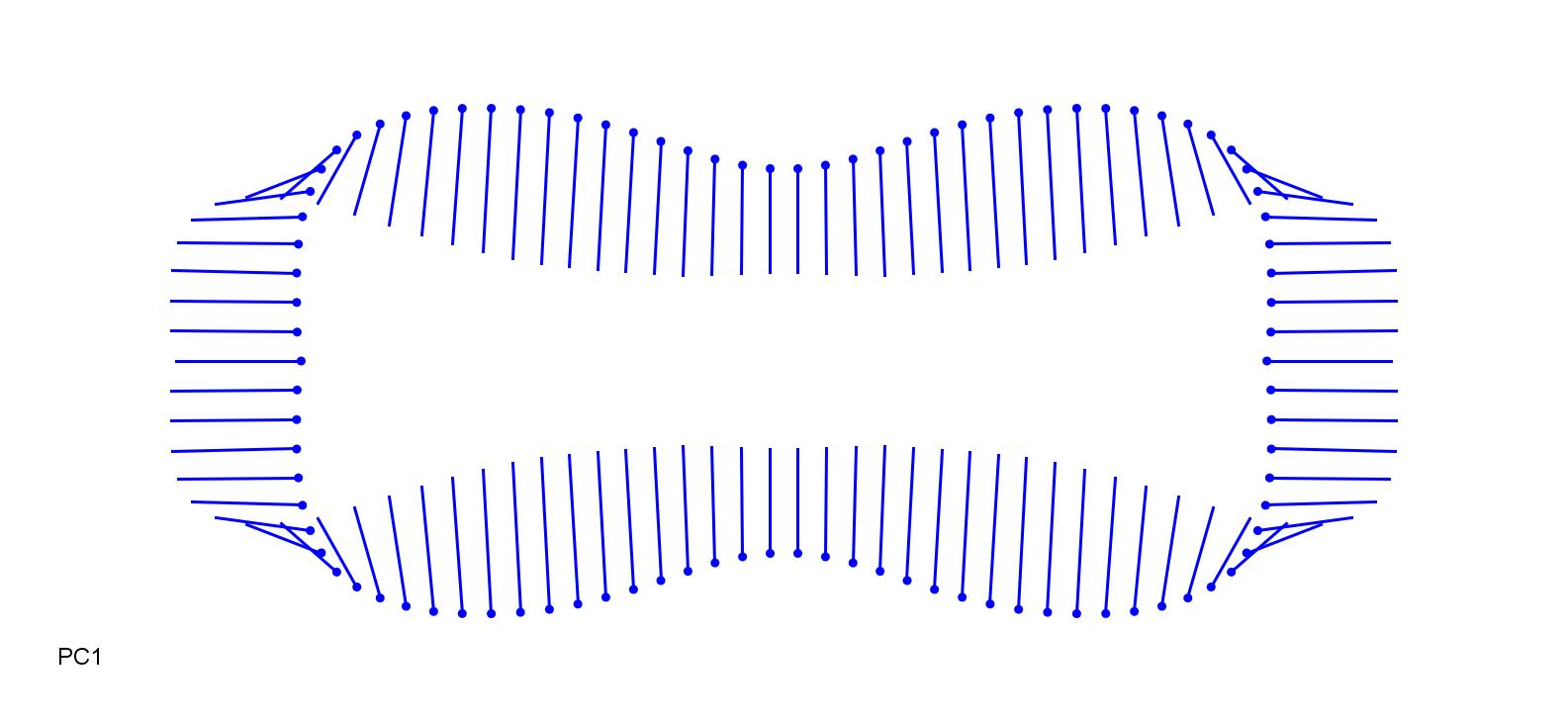

### BraEre-0.25.jpg

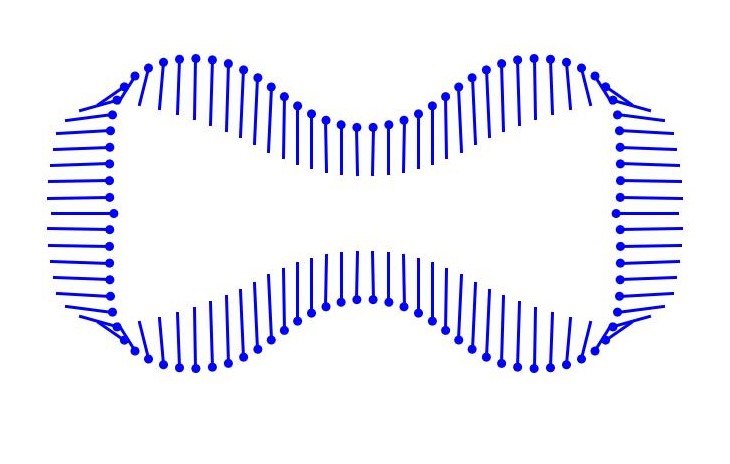

### BraEre_0.3.jpg

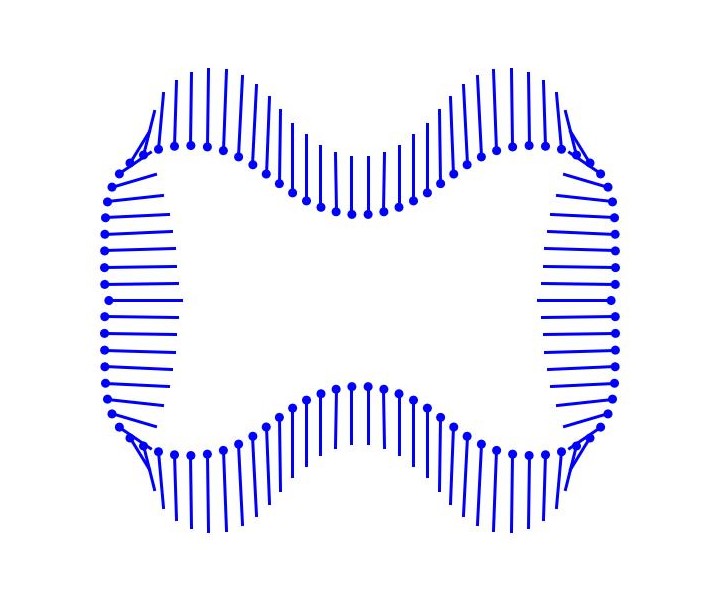

### BraPinn-0.2.jpg

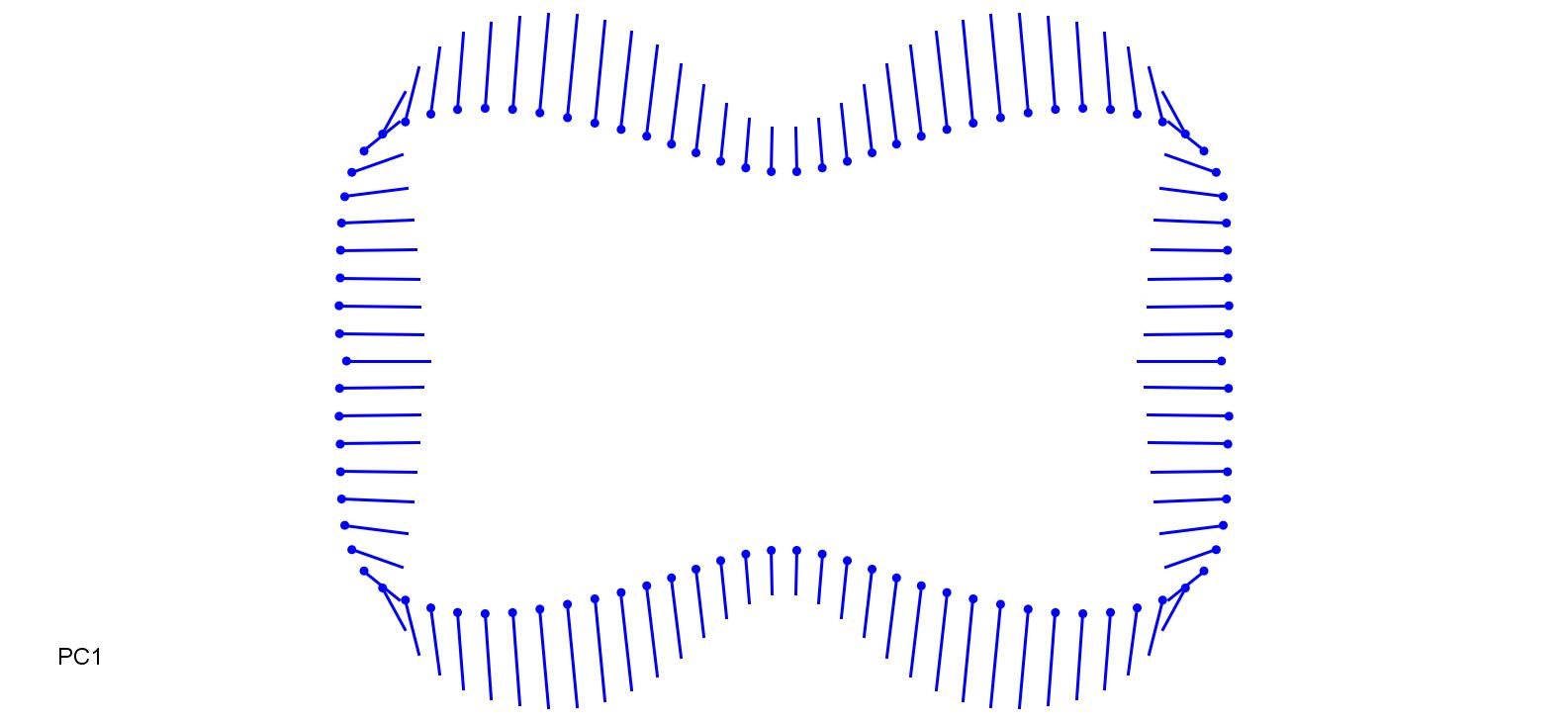

### BraPinn_0.3.jpg

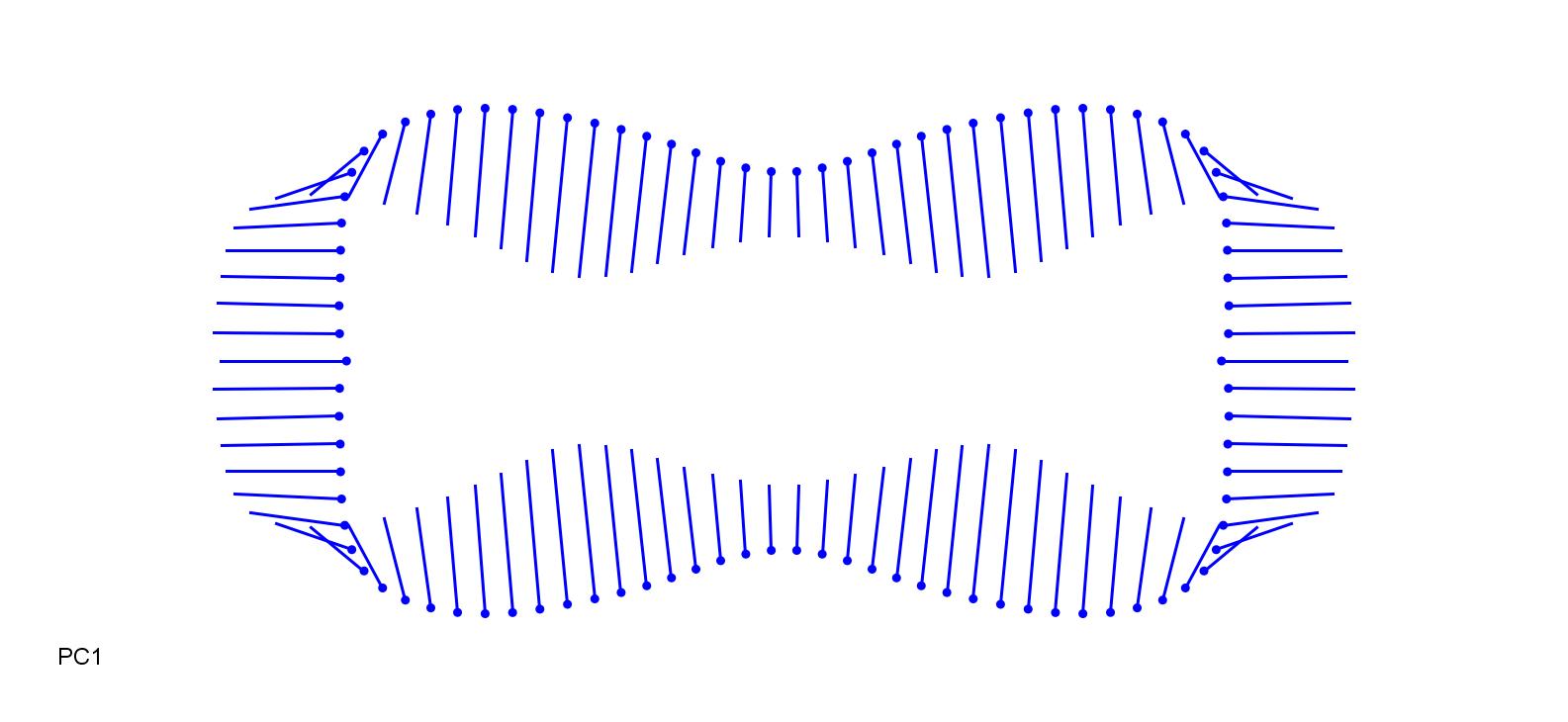

### BraSyl-0.15.jpg

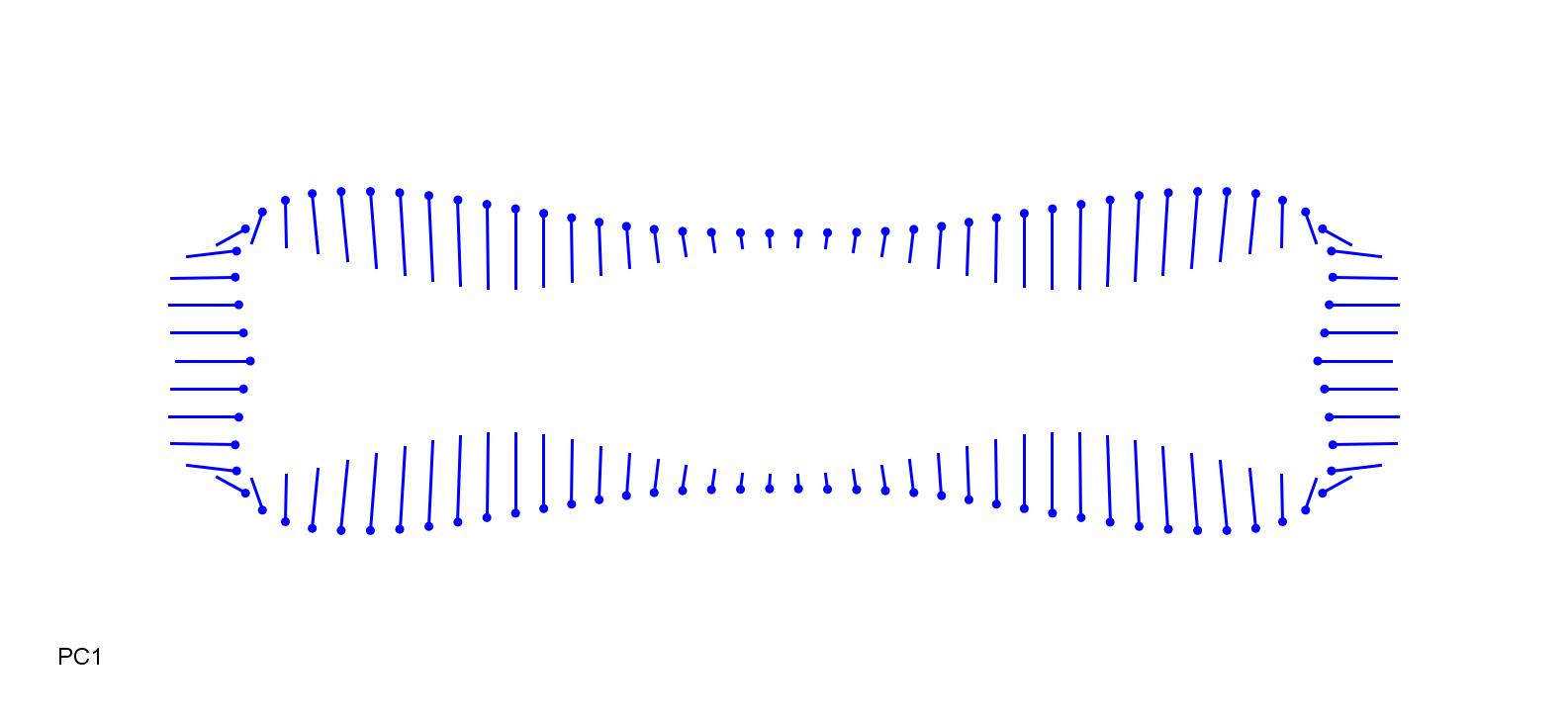

### BraSyl_0.15.jpg

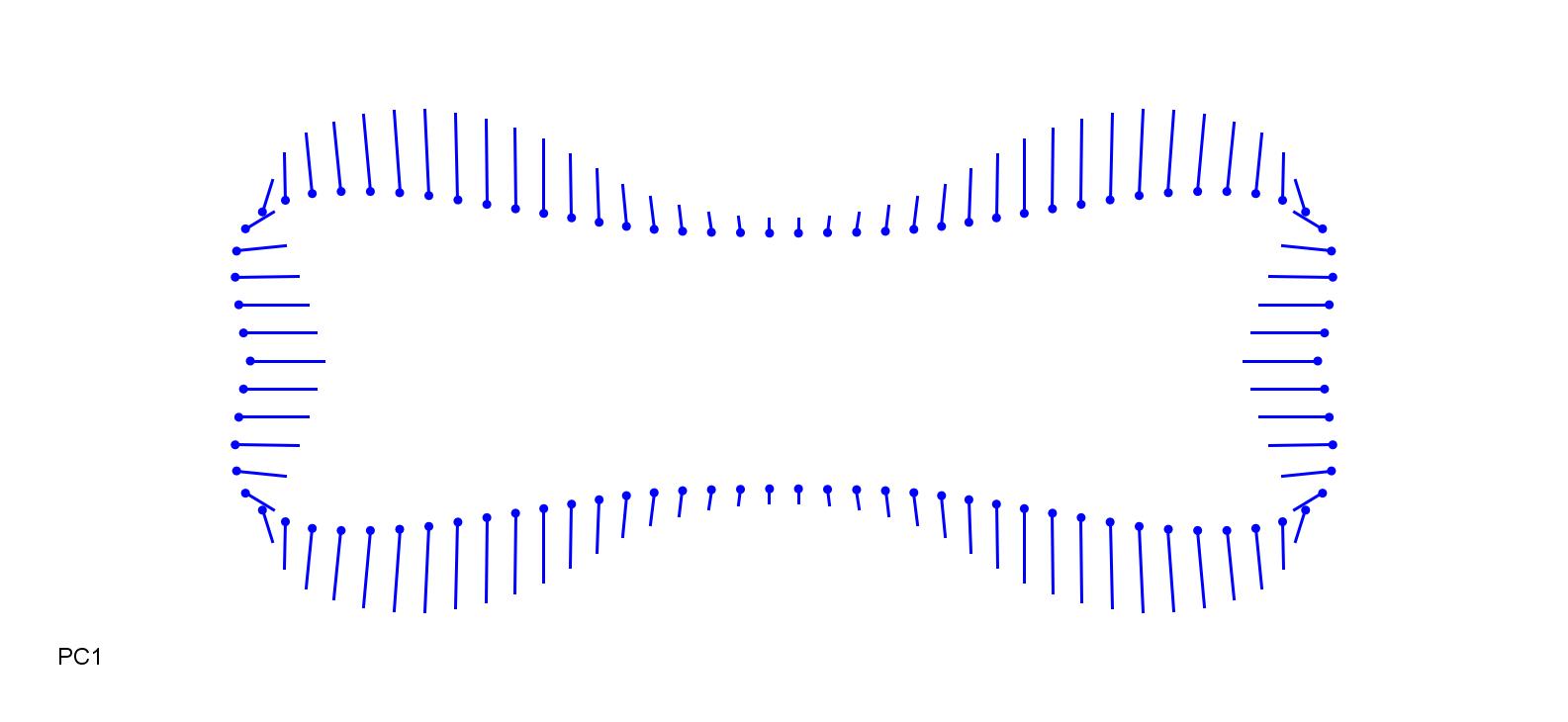

### BroBen-0.15.jpg

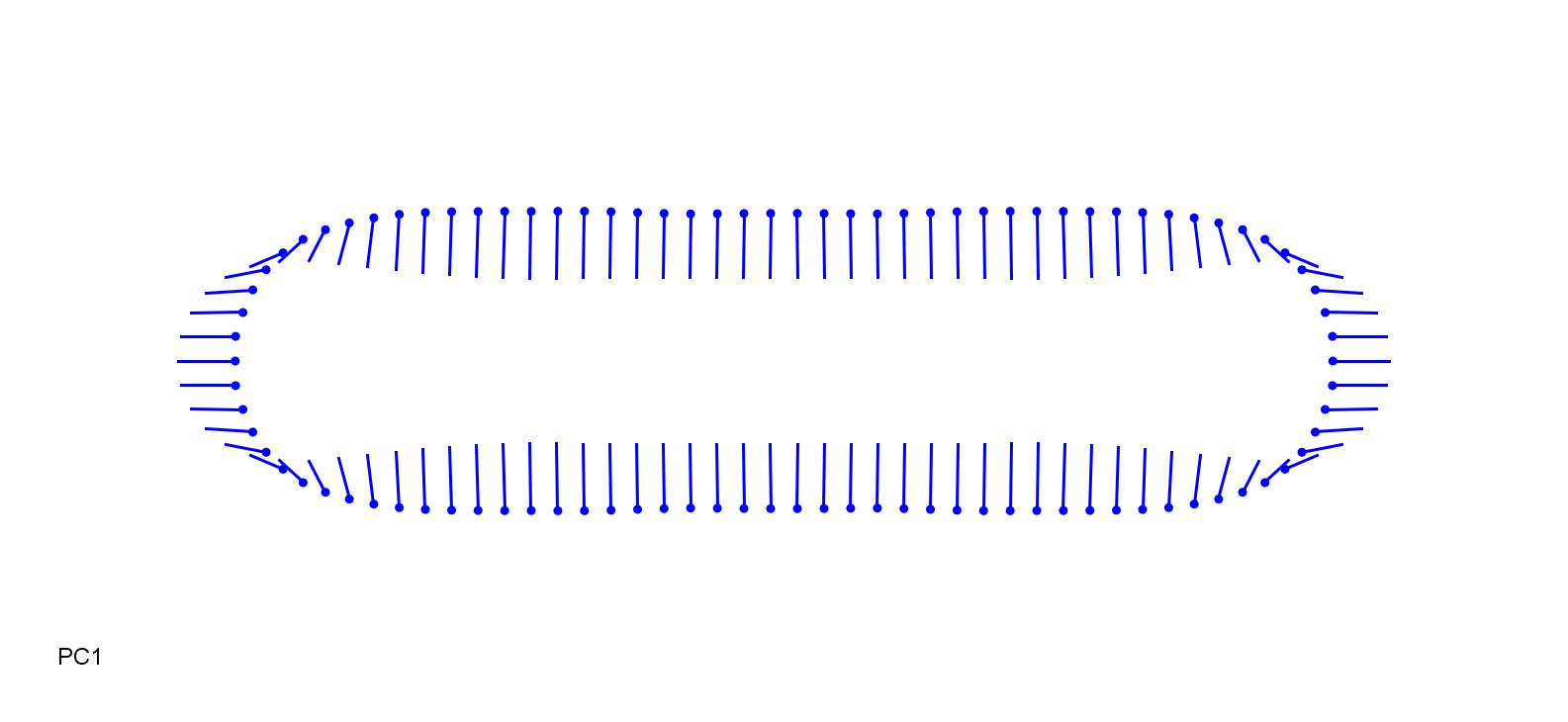

### BroBen_0.3.jpg

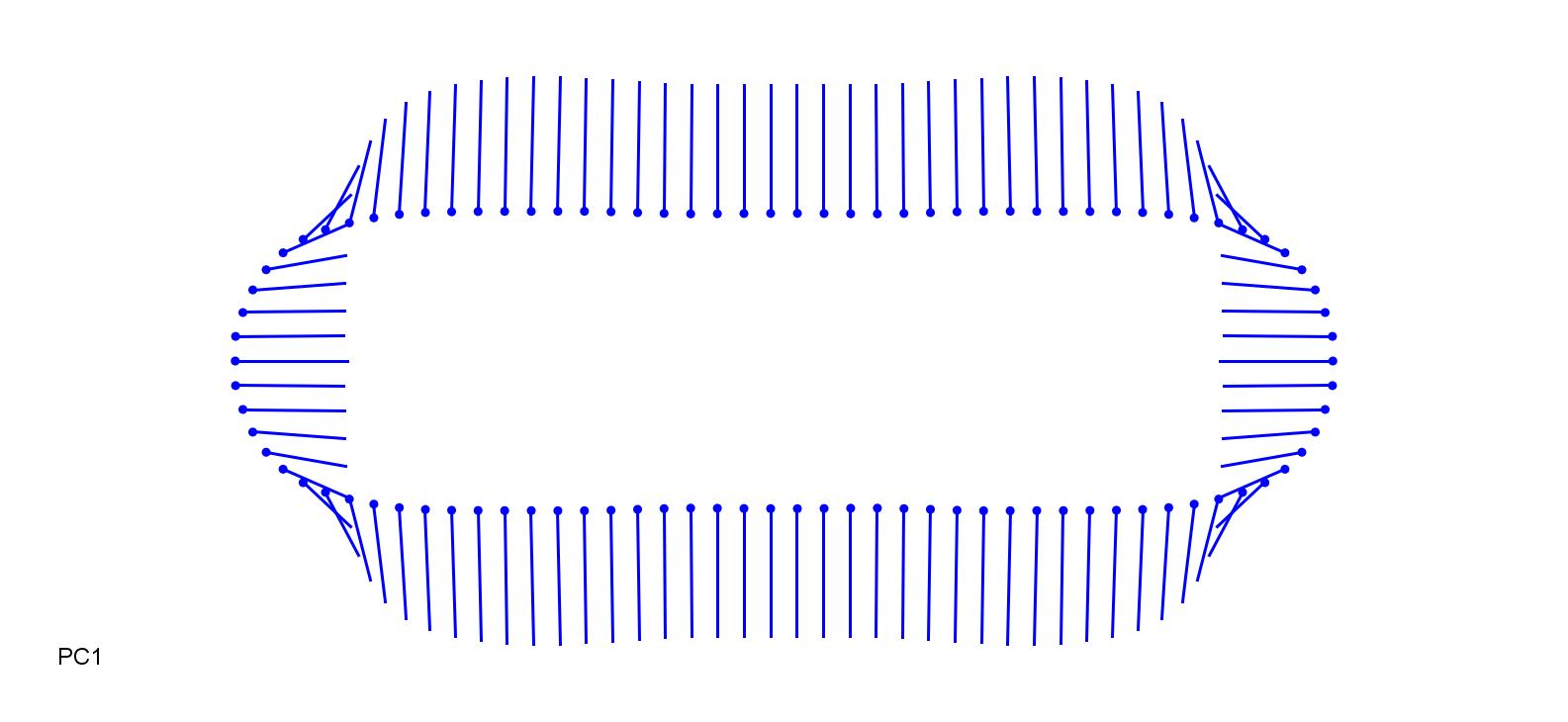

### BroEre-0.25.jpg

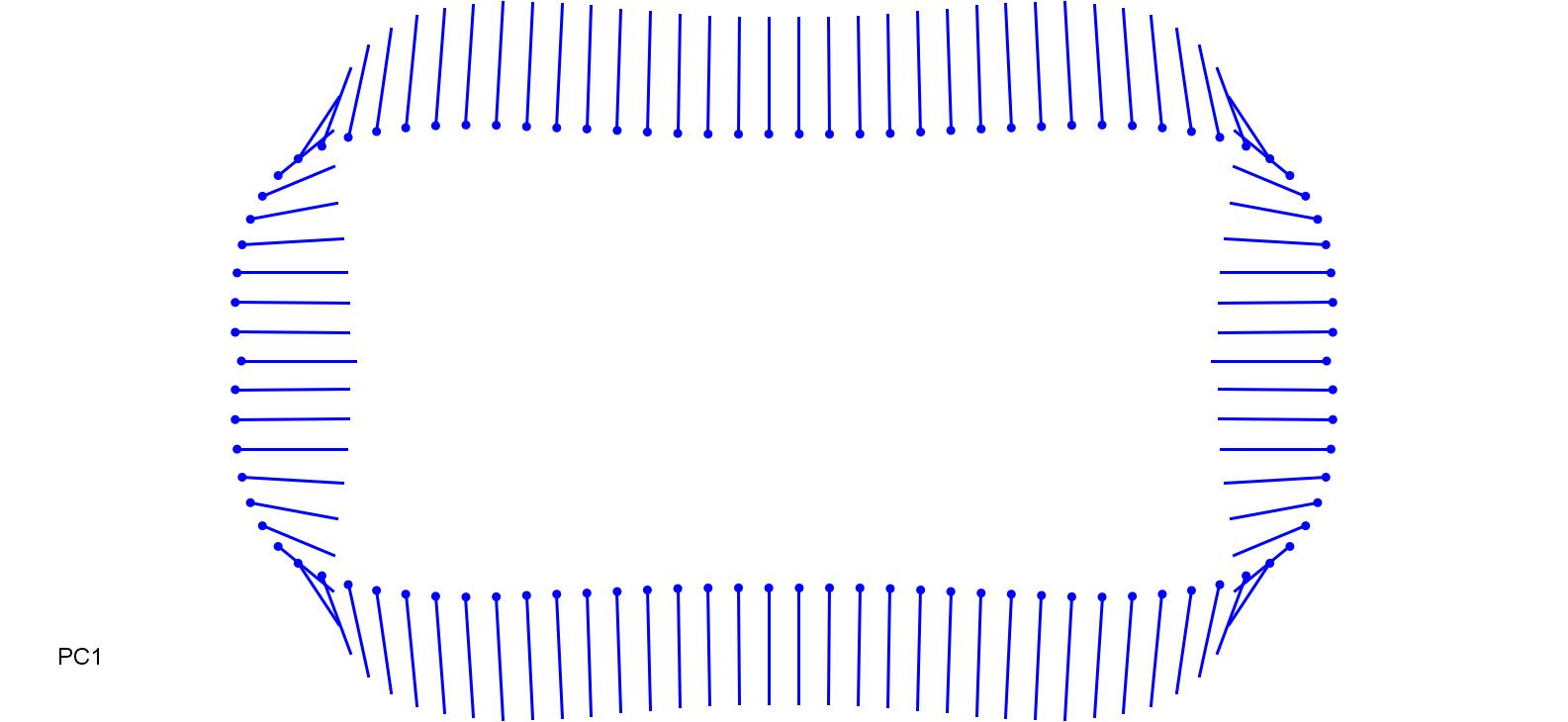

### BroEre_0.3.jpg

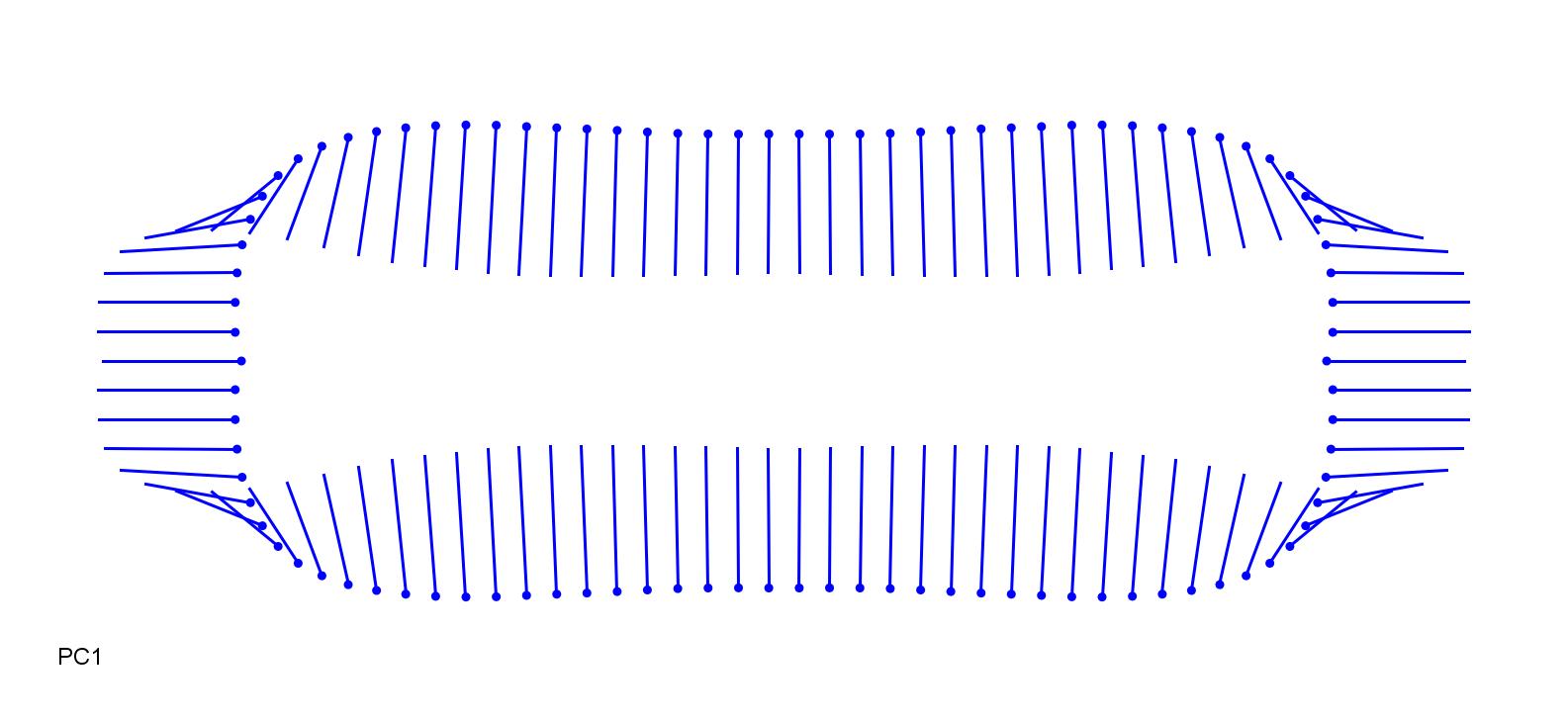

### BroIne-0.1.jpg

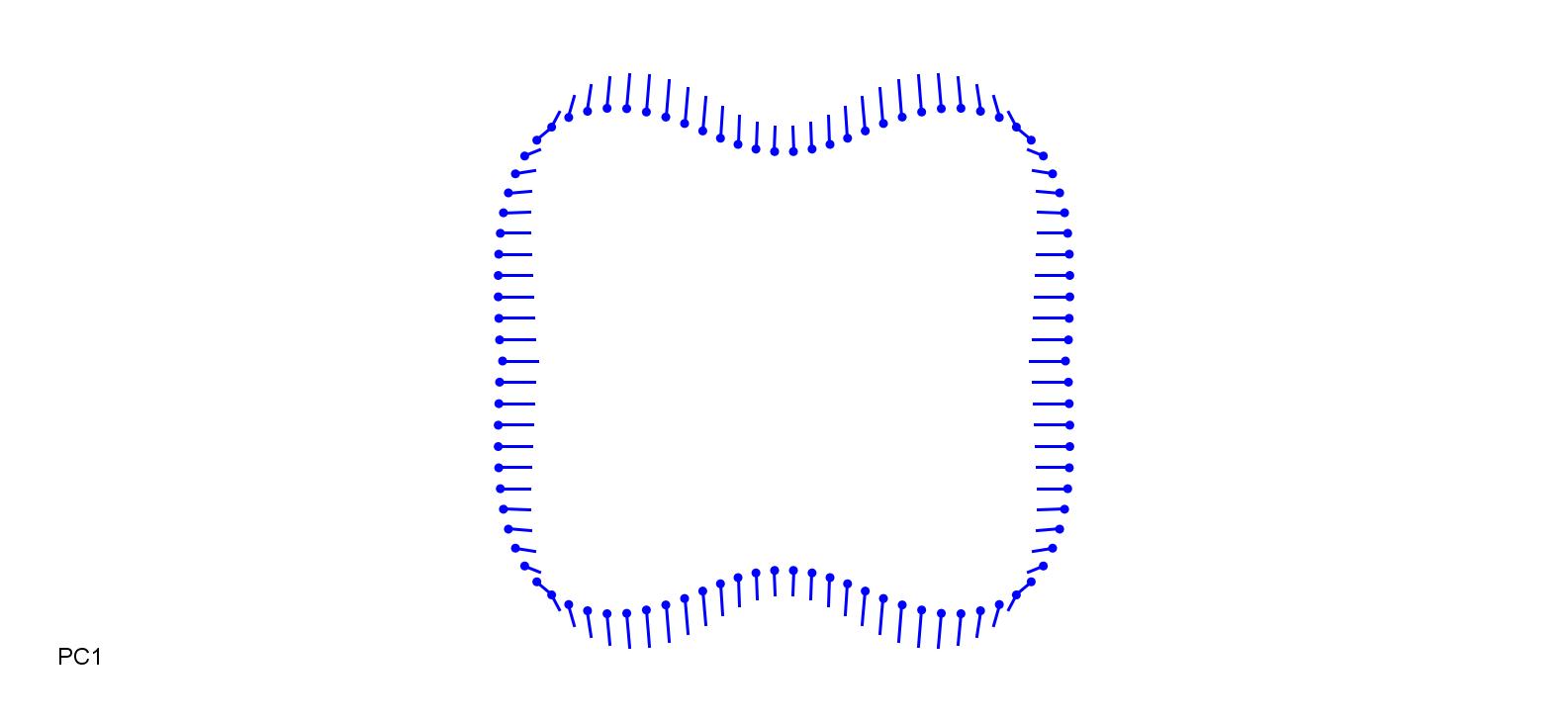

### BroIne_0.3.jpg

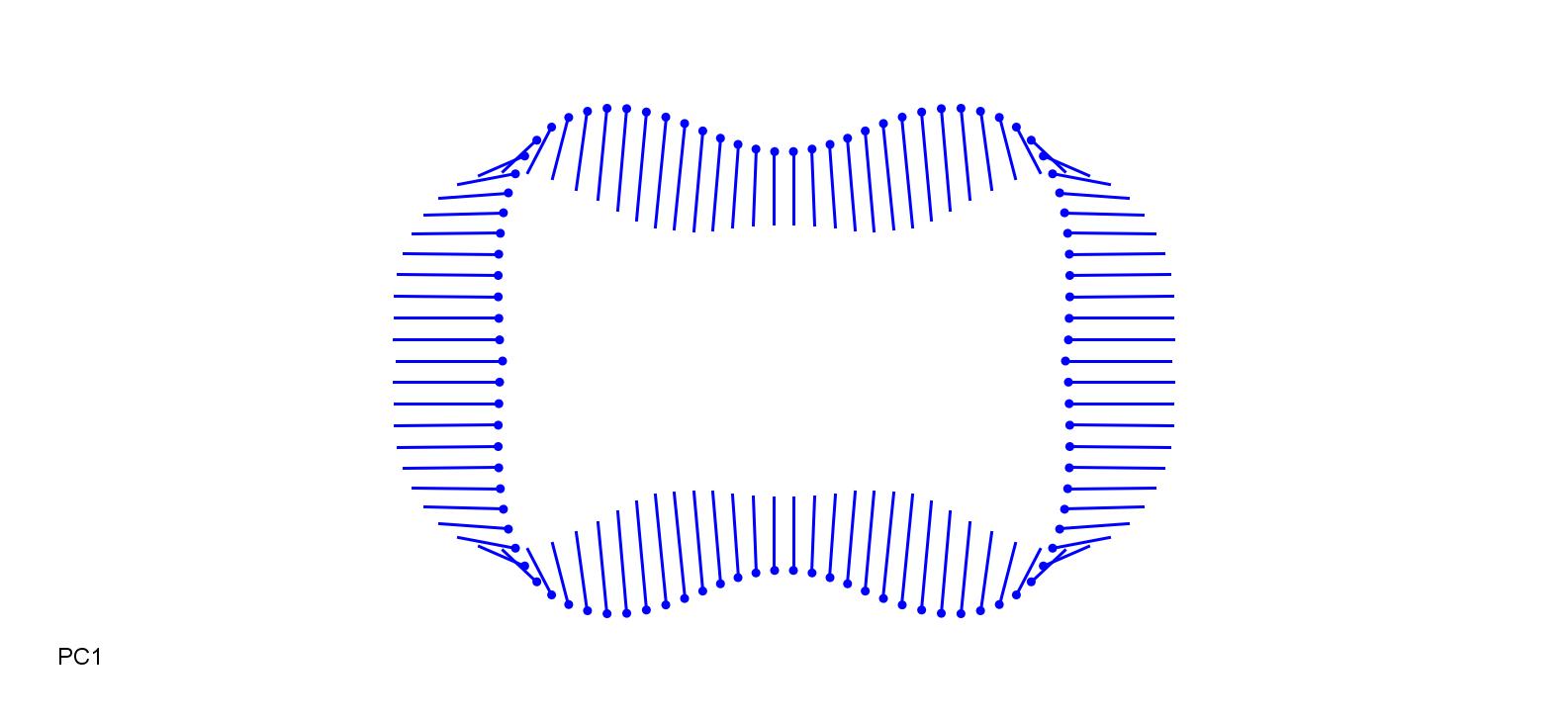

### GlyFlu-0.13.jpg

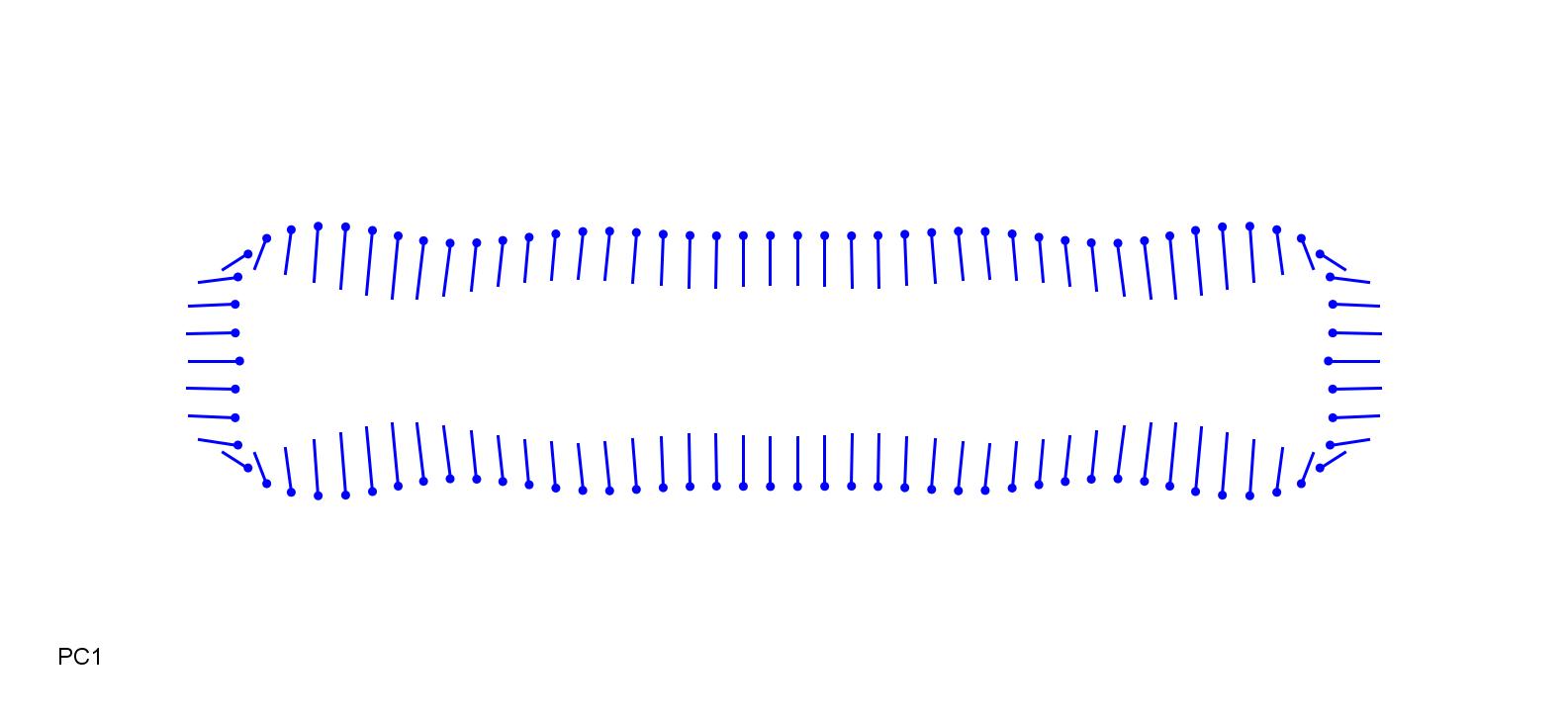

### GlyFlu_0.21.jpg

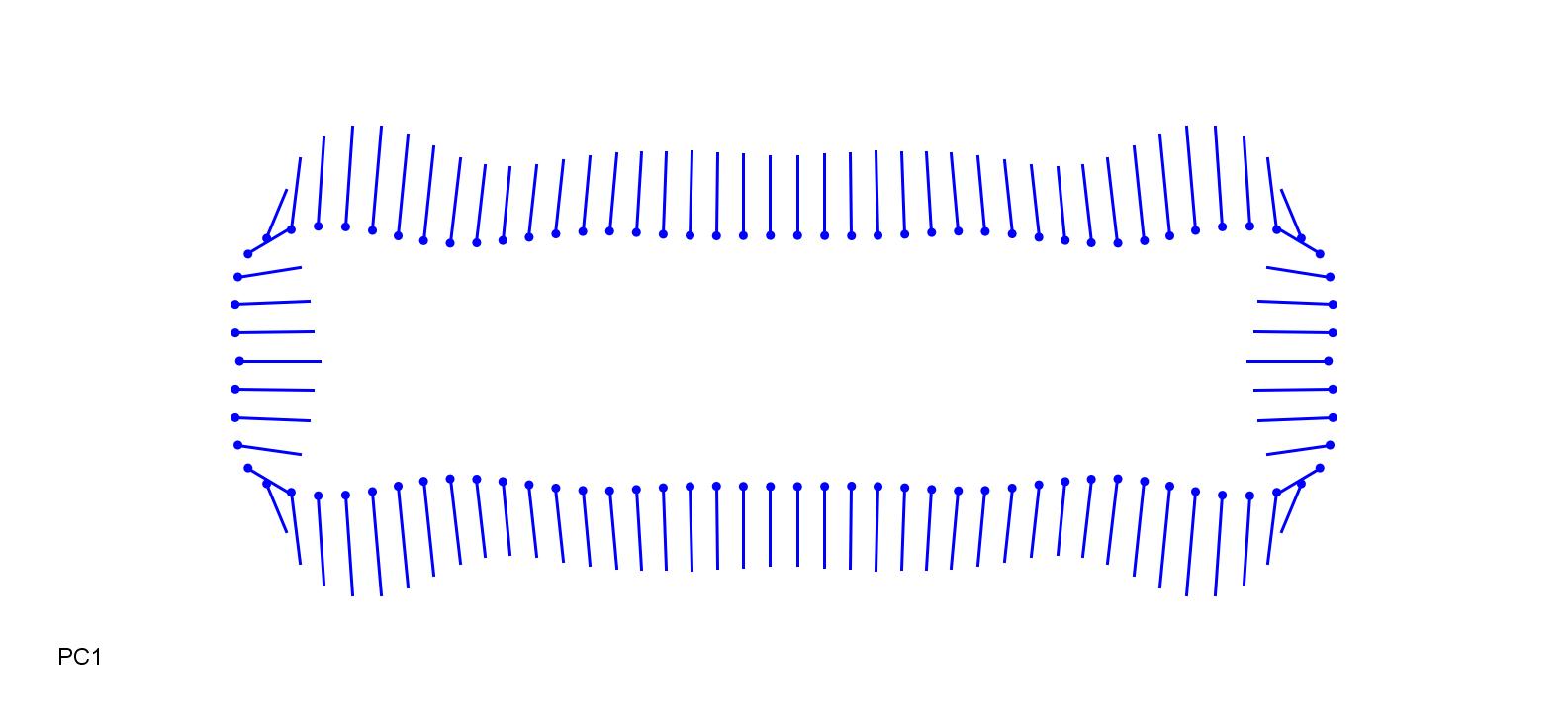

### GlyMax-0.15.jpg

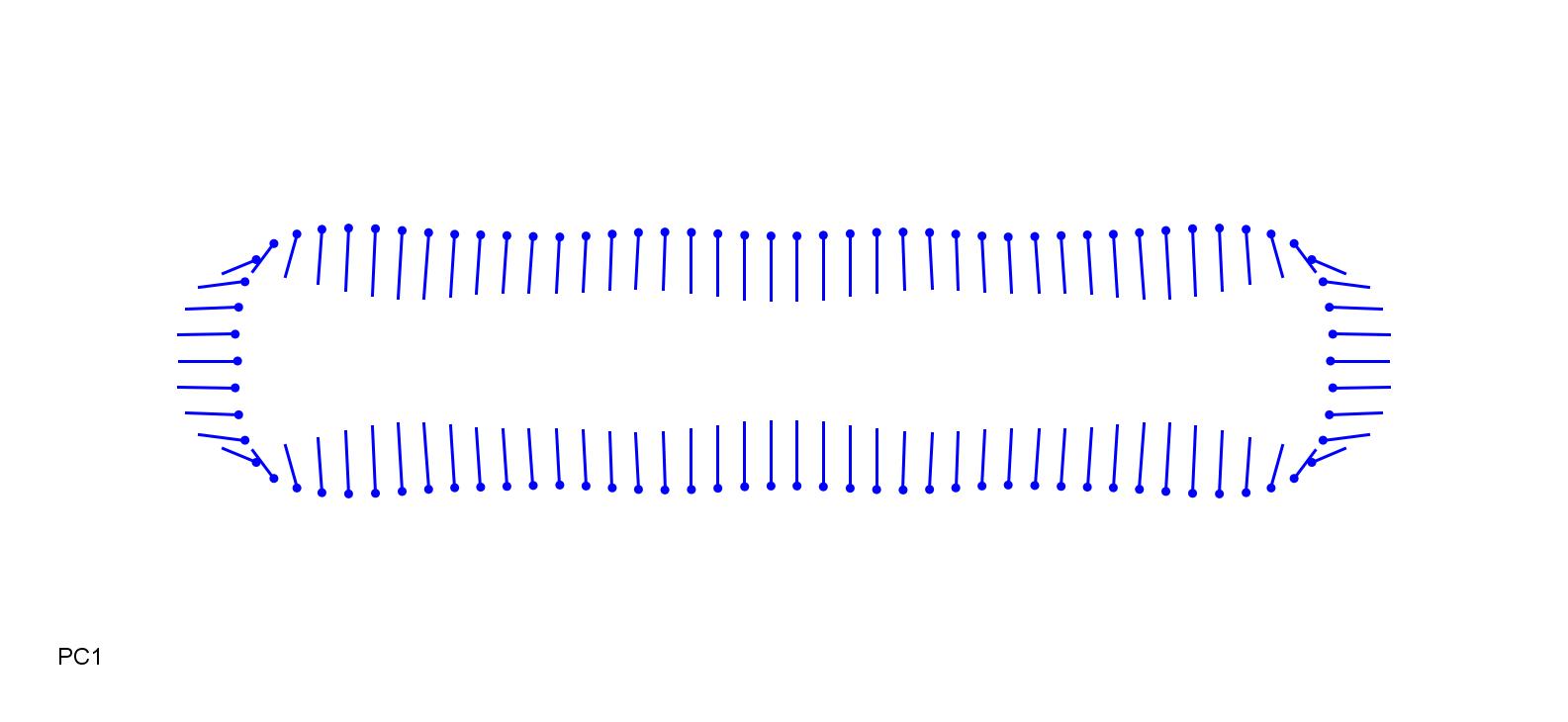

### GlyMax_0.25.jpg

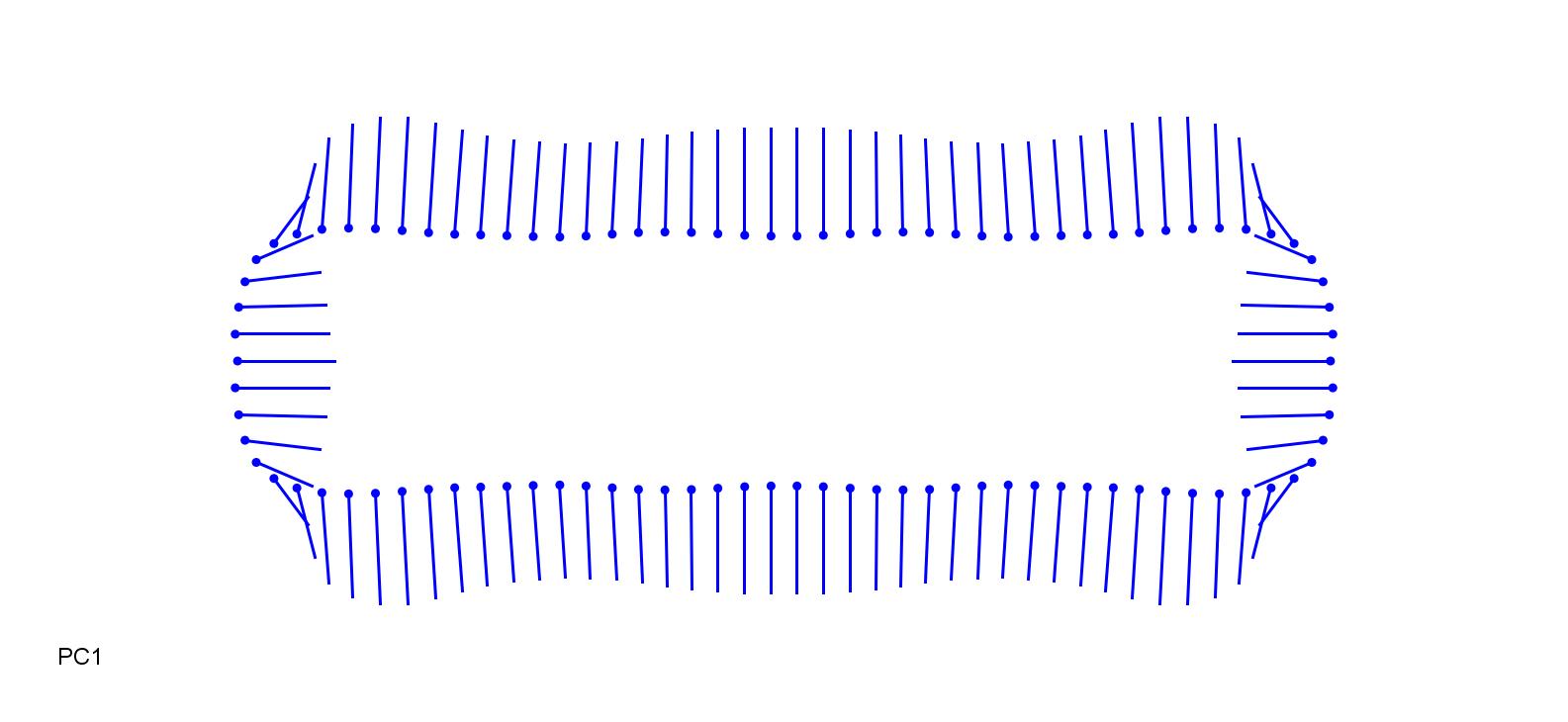

### MelPic-0.15.jpg

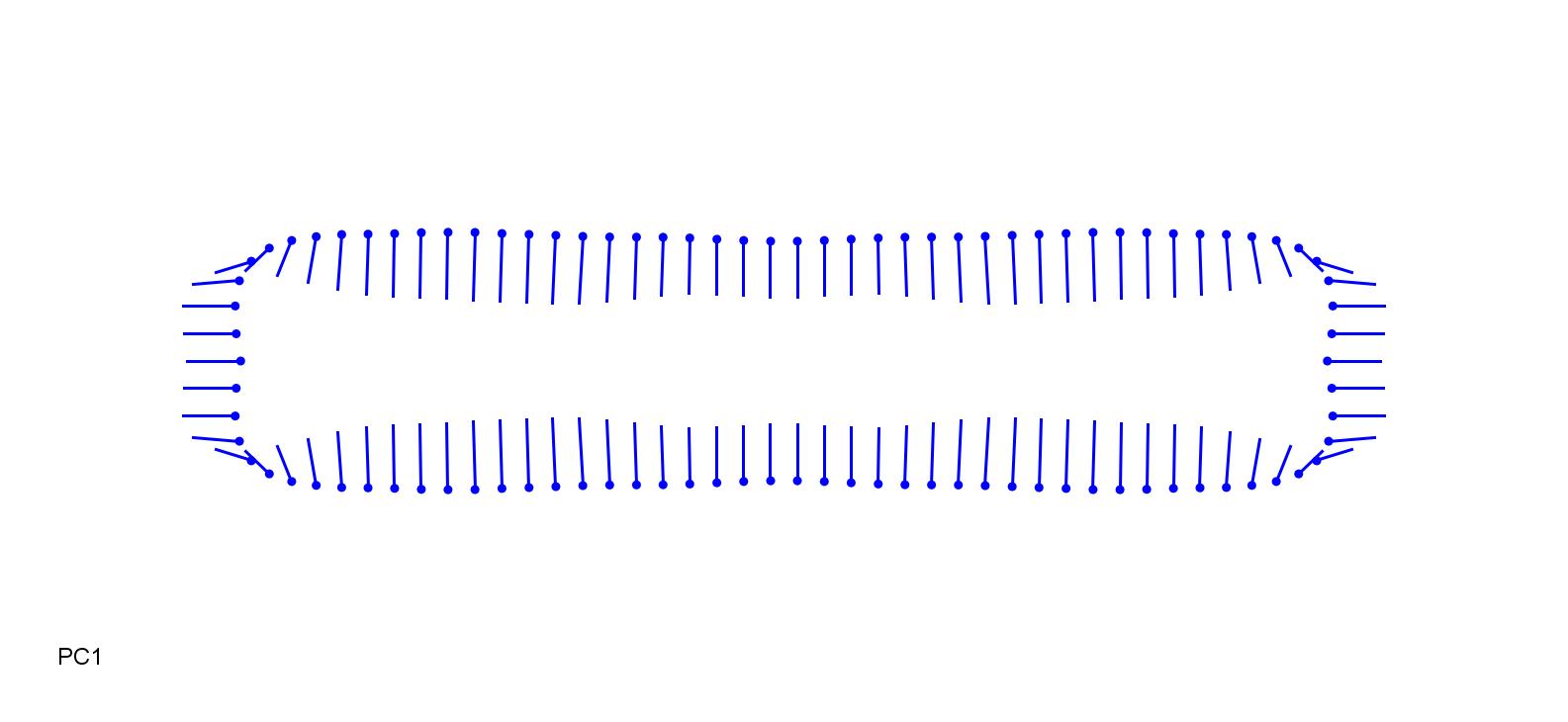

### MelPic_0.3.jpg

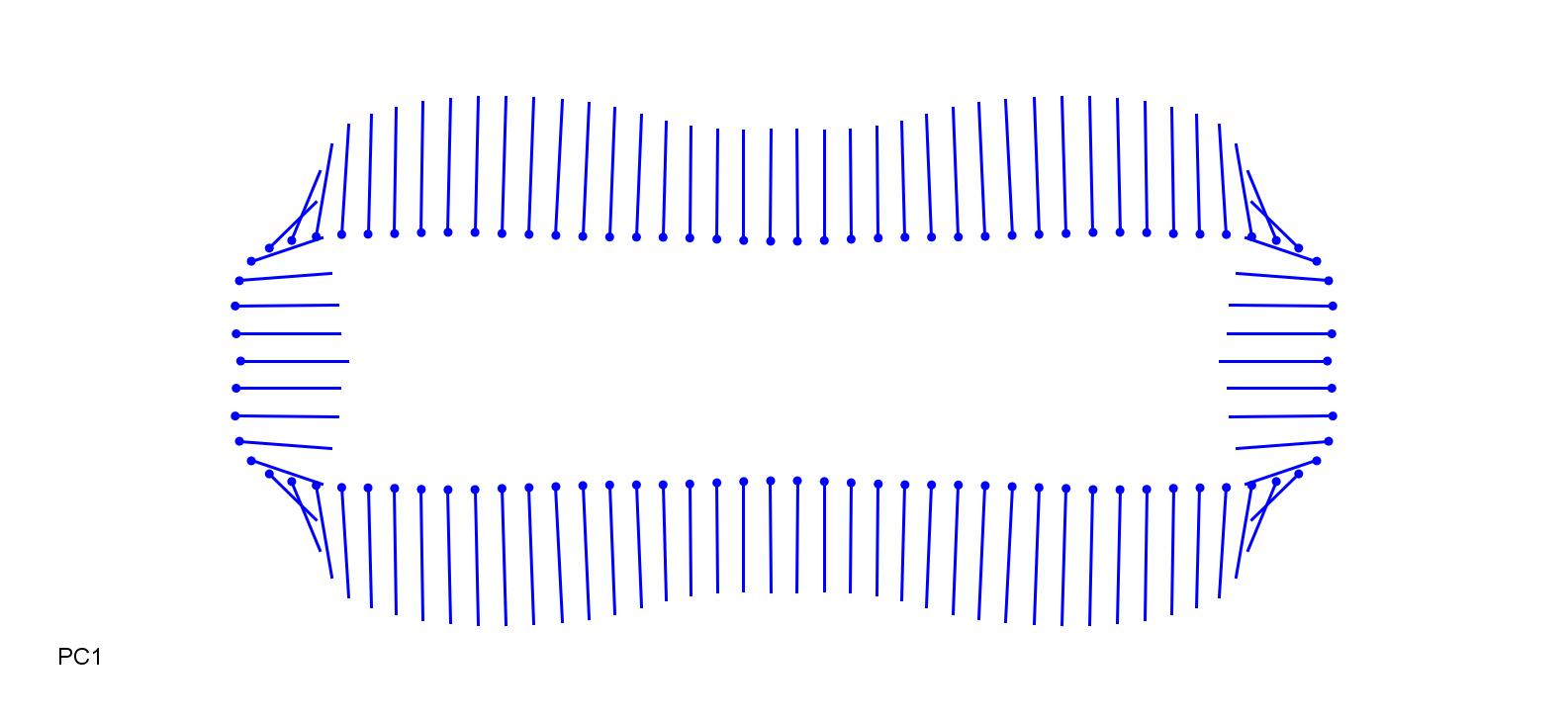

### MelUni-0.13.jpg

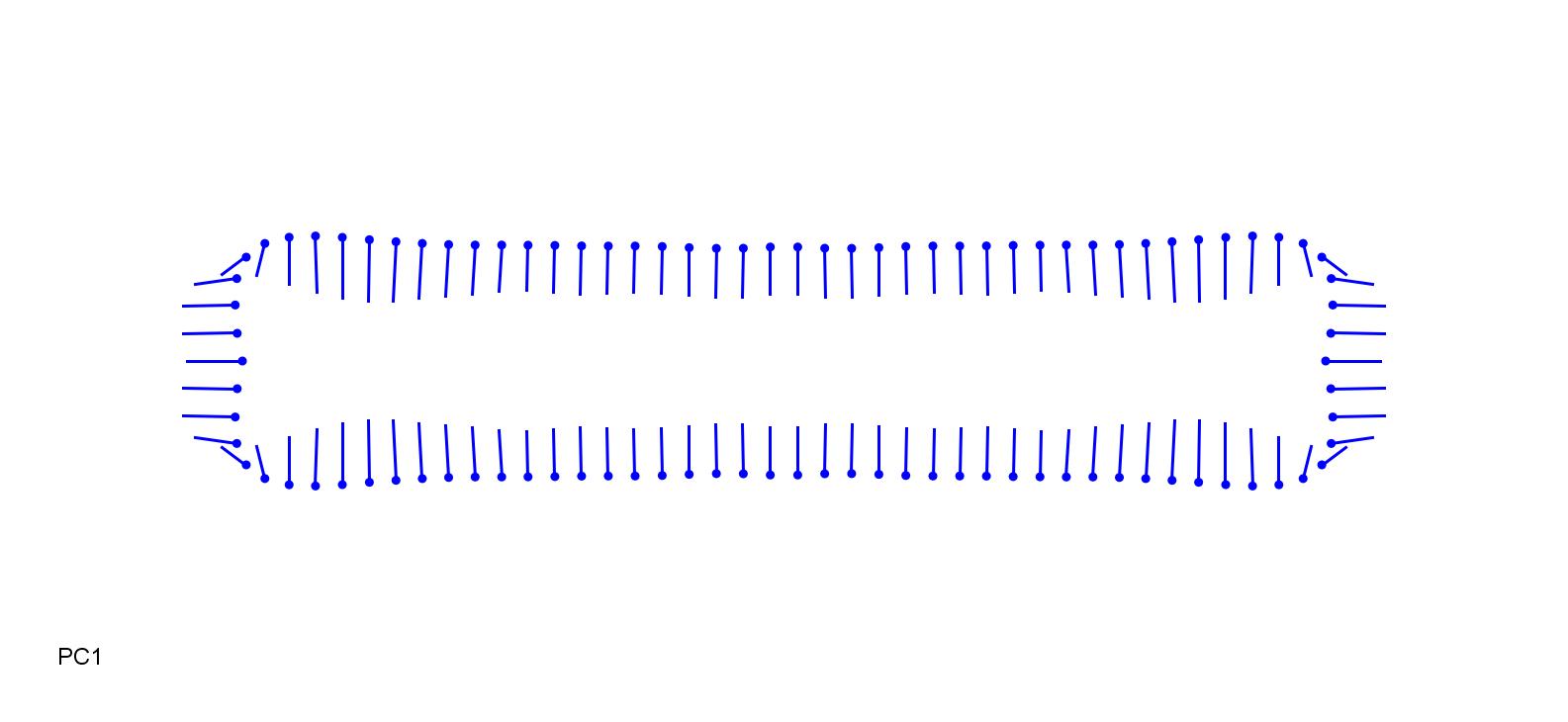

### MelUni_0.18.jpg

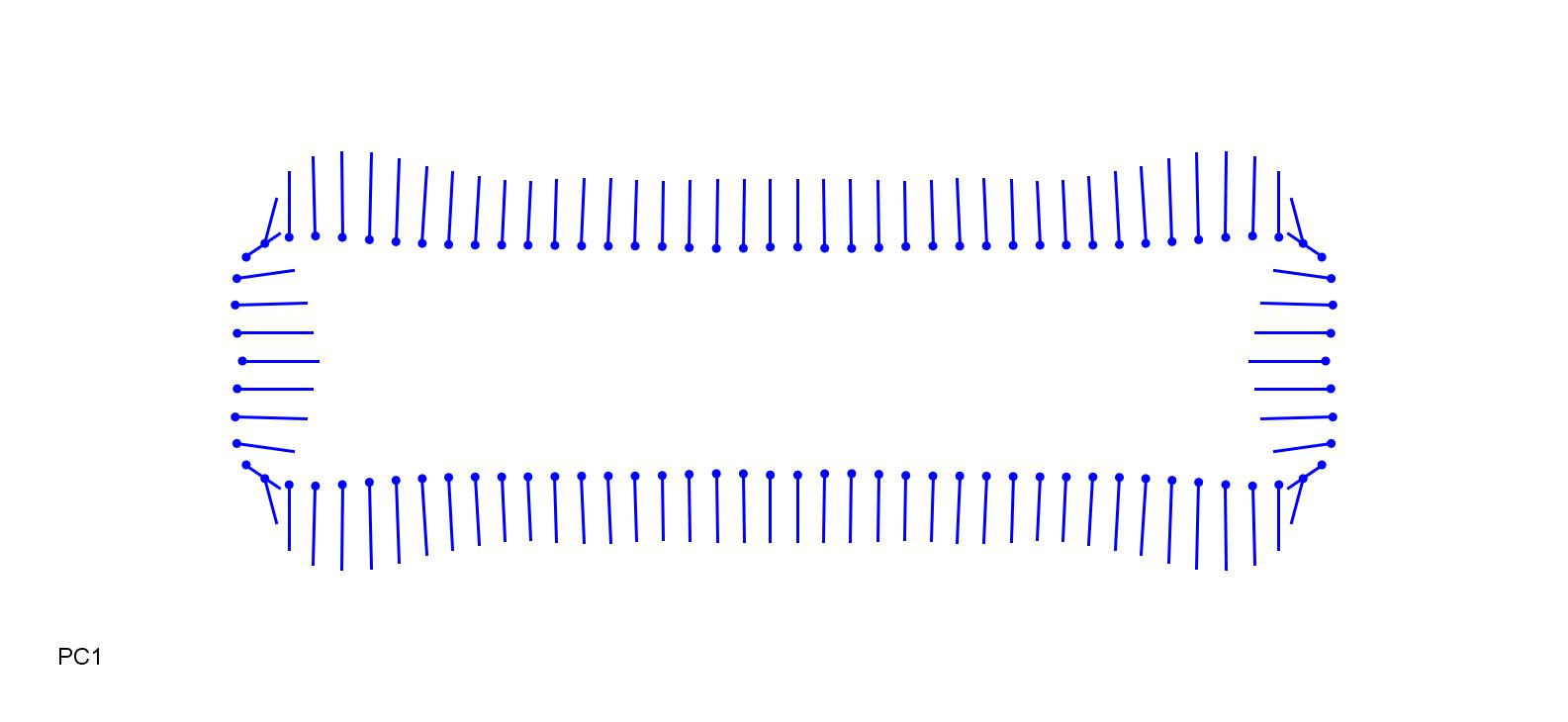

### PhyBam-0.18.jpg

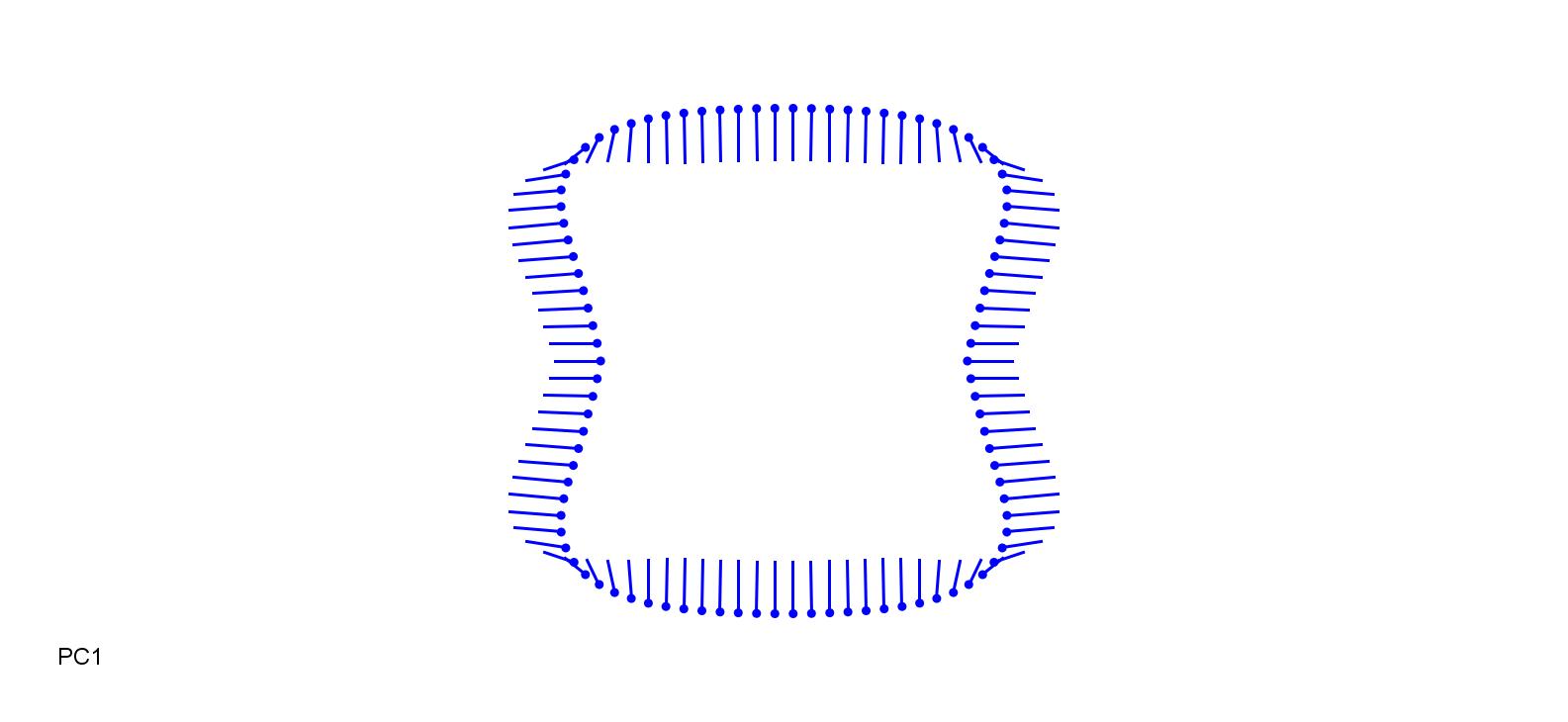

### PhyBam_0.18.jpg

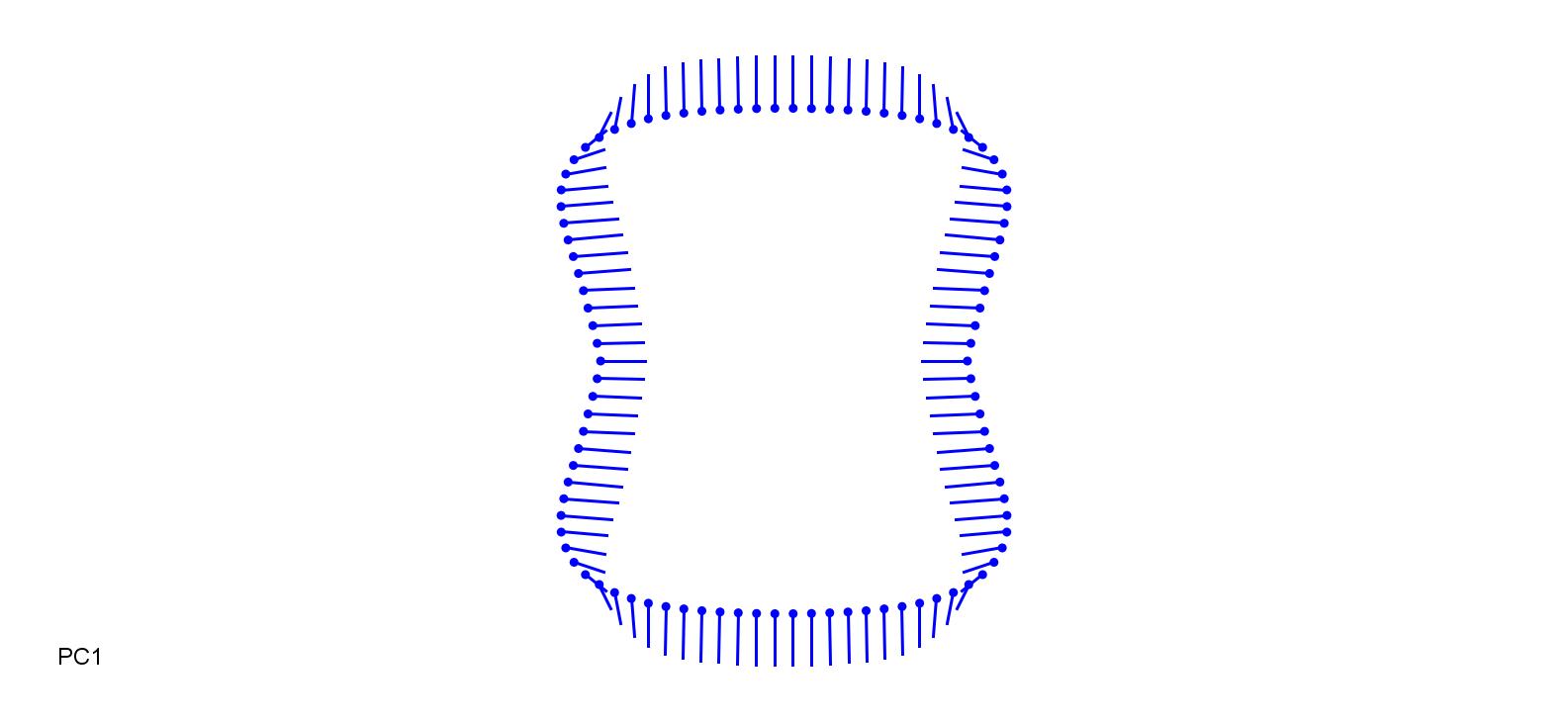

### PhyNig-0.13.jpg

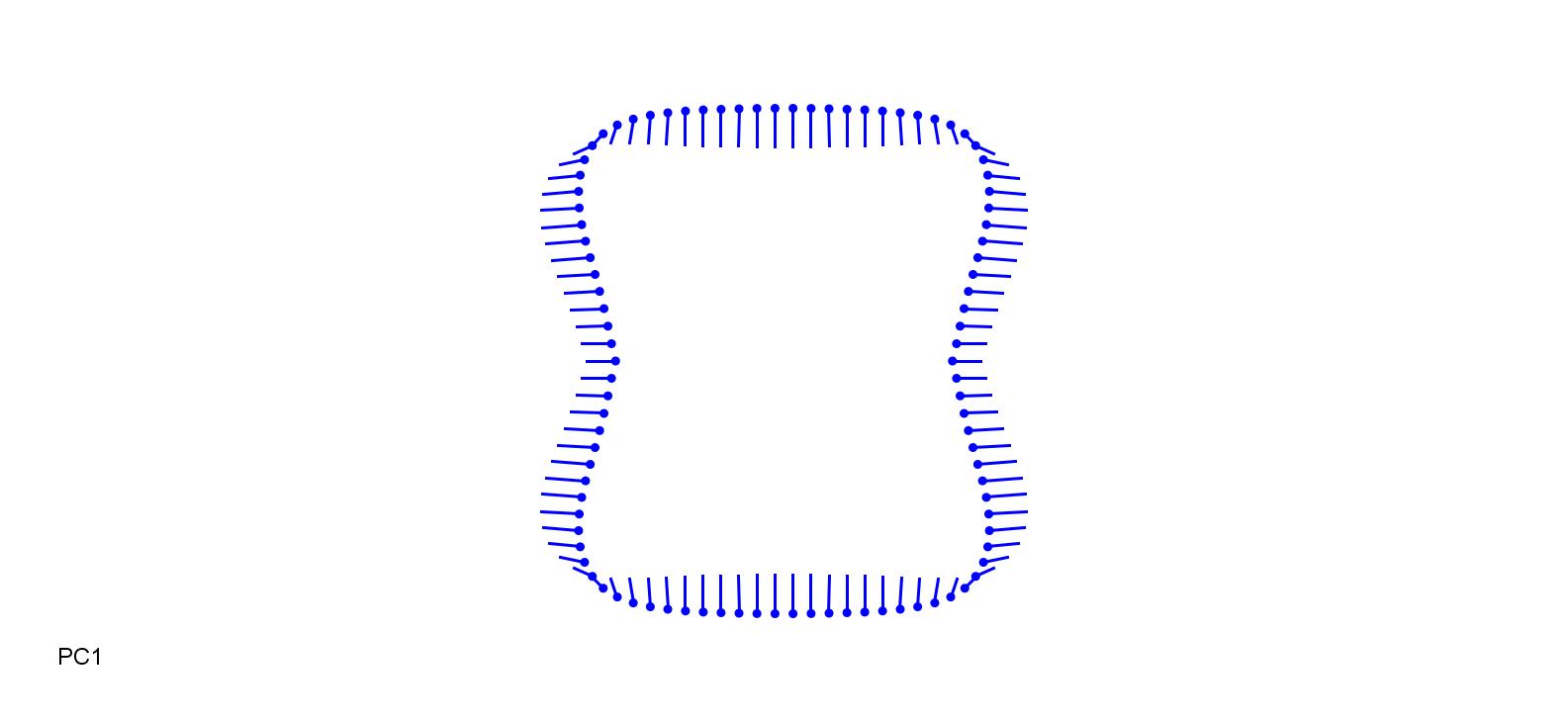

### PhyNig_0.13.jpg

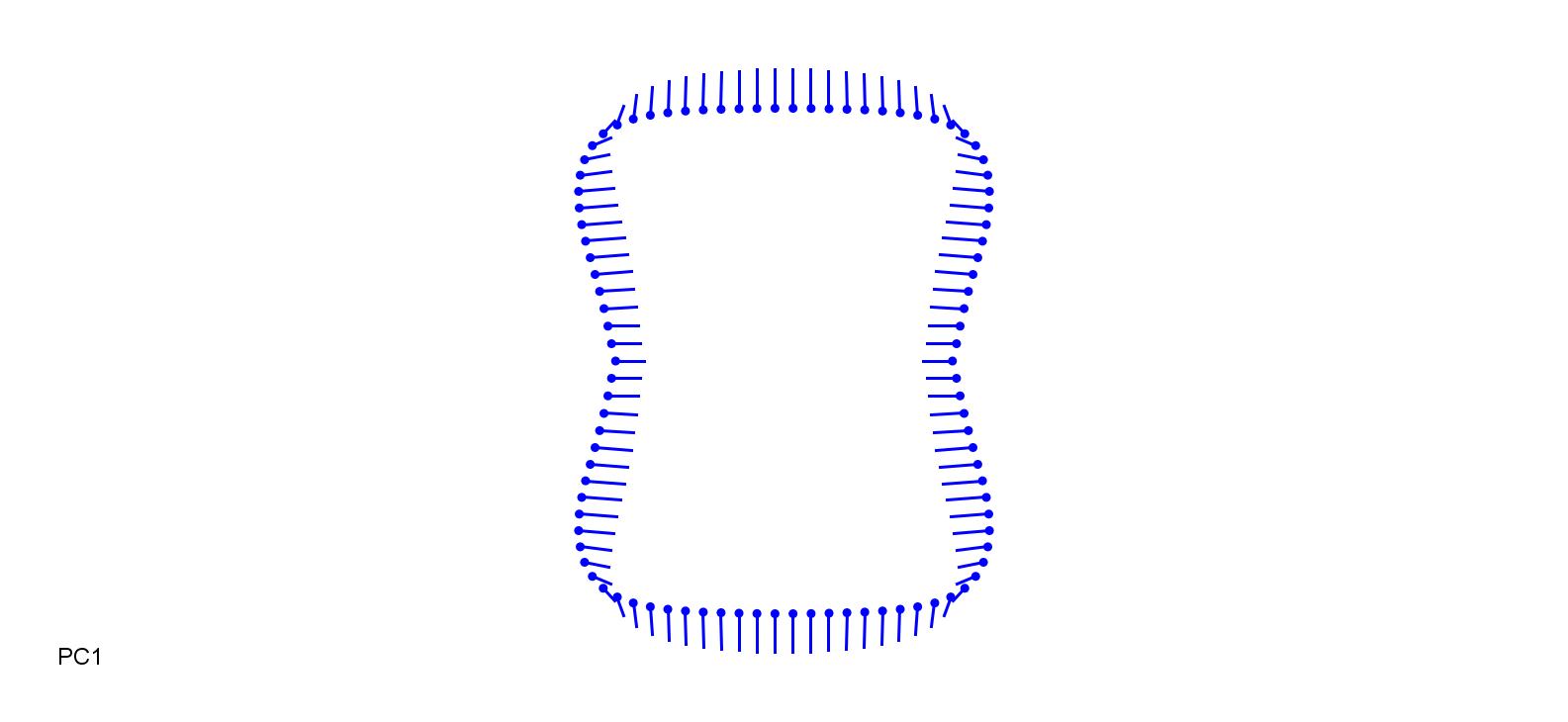

### PhyNig_0.15.jpg

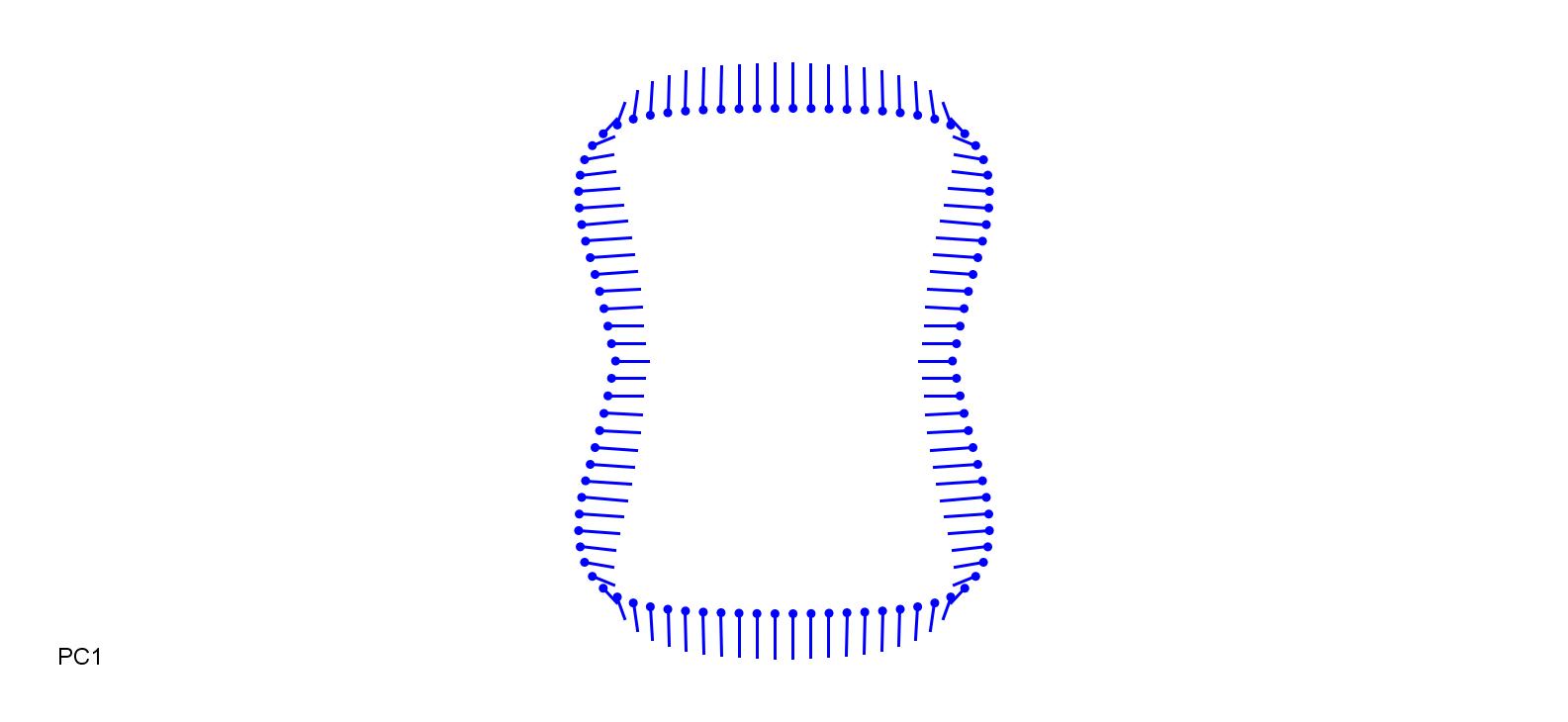

### PipMil+0.15.jpg

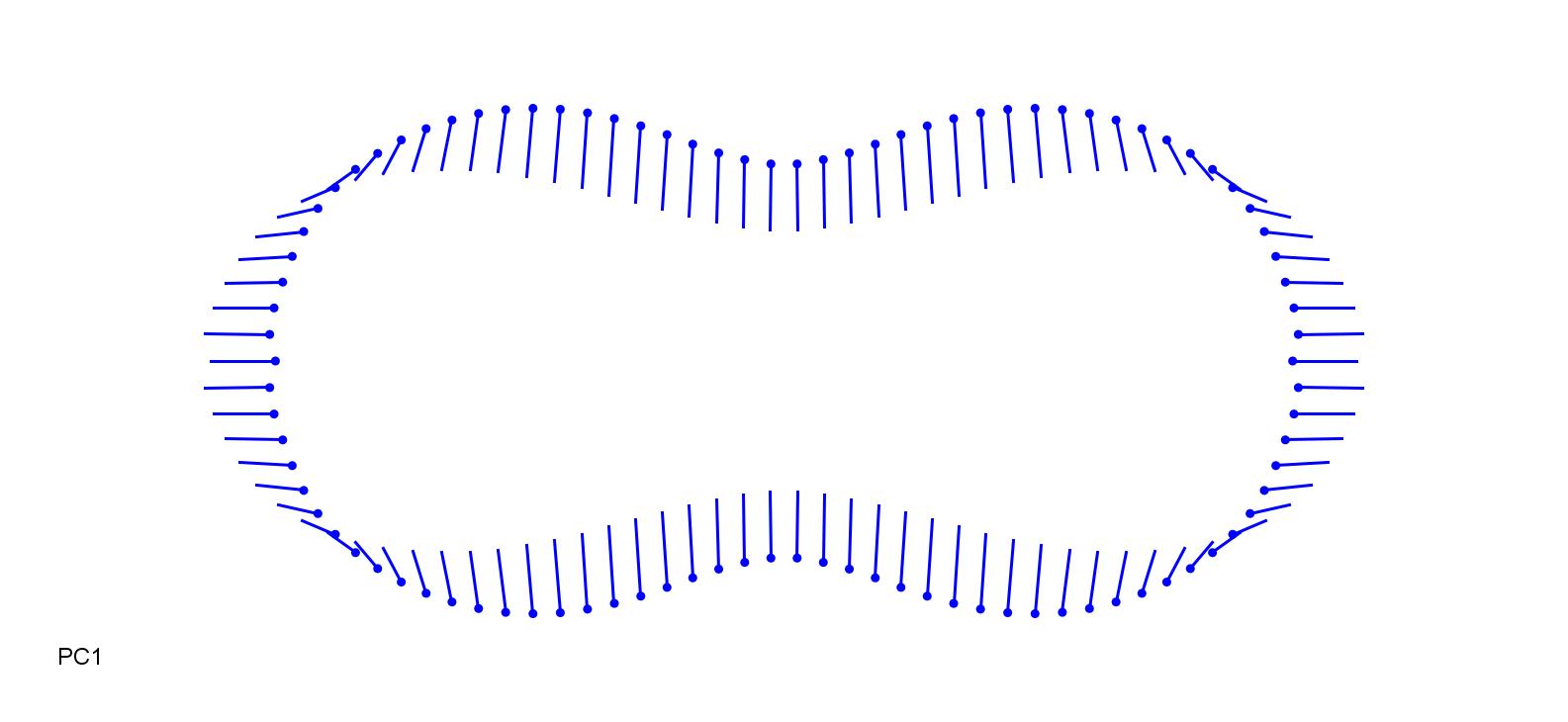

### PipMil-0.15.jpg

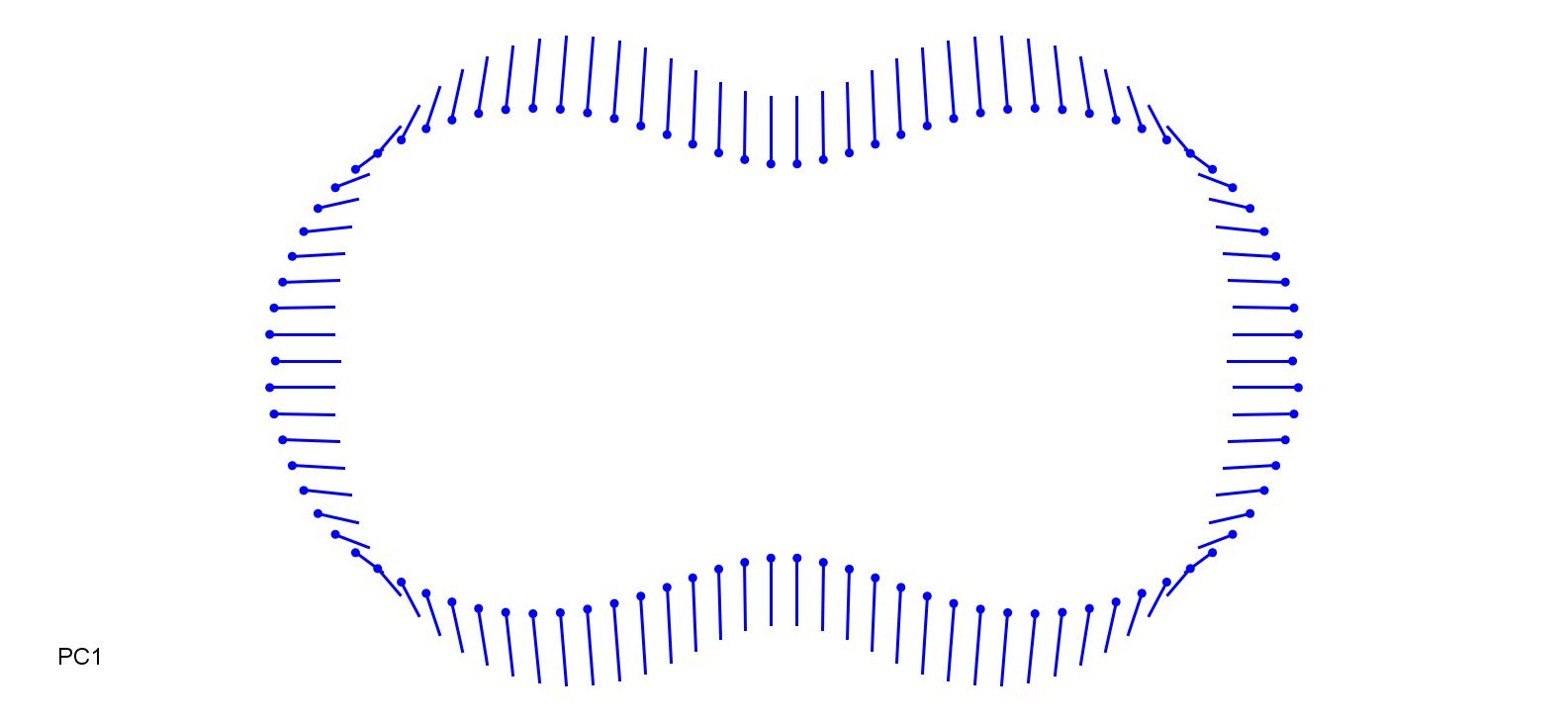

### PipVir-0.1.jpg

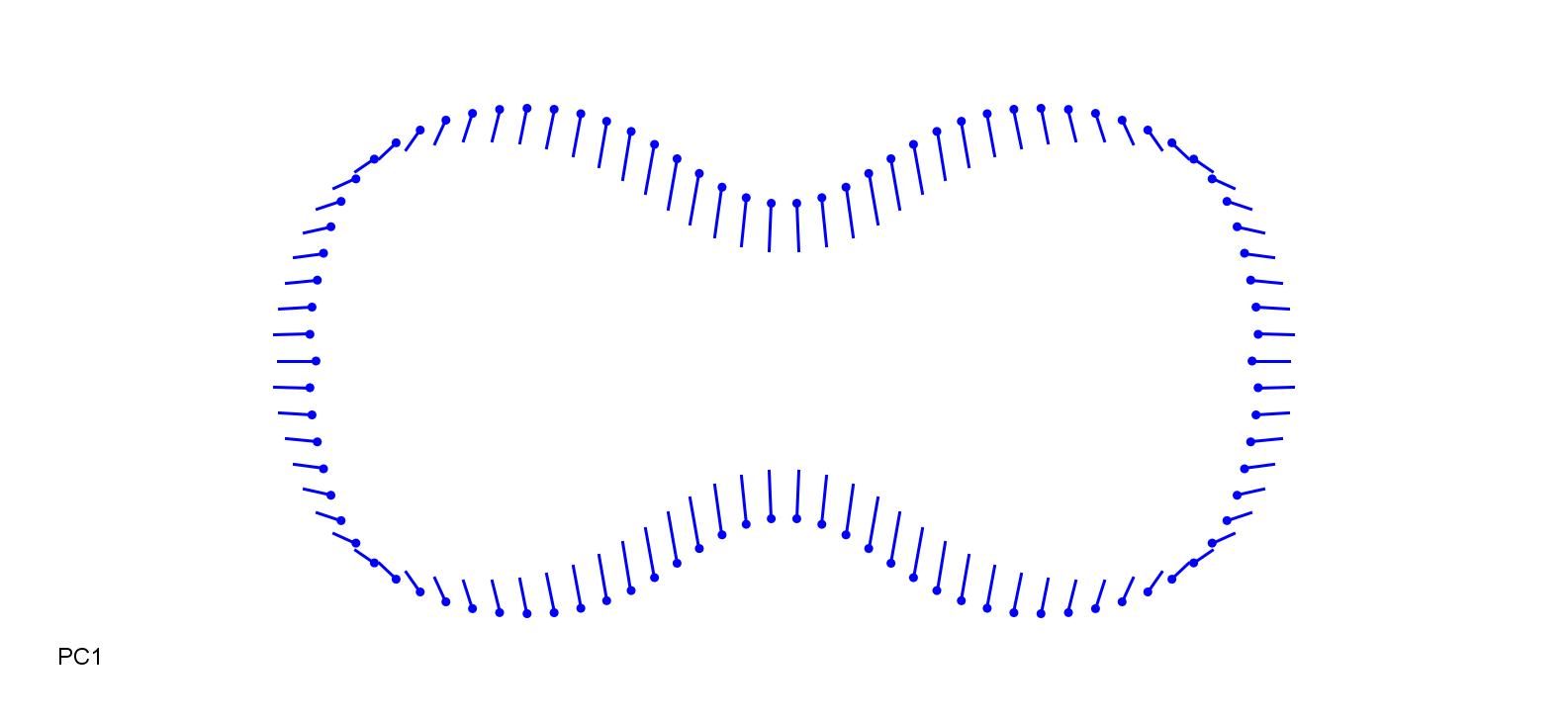

### PipVir.jpg

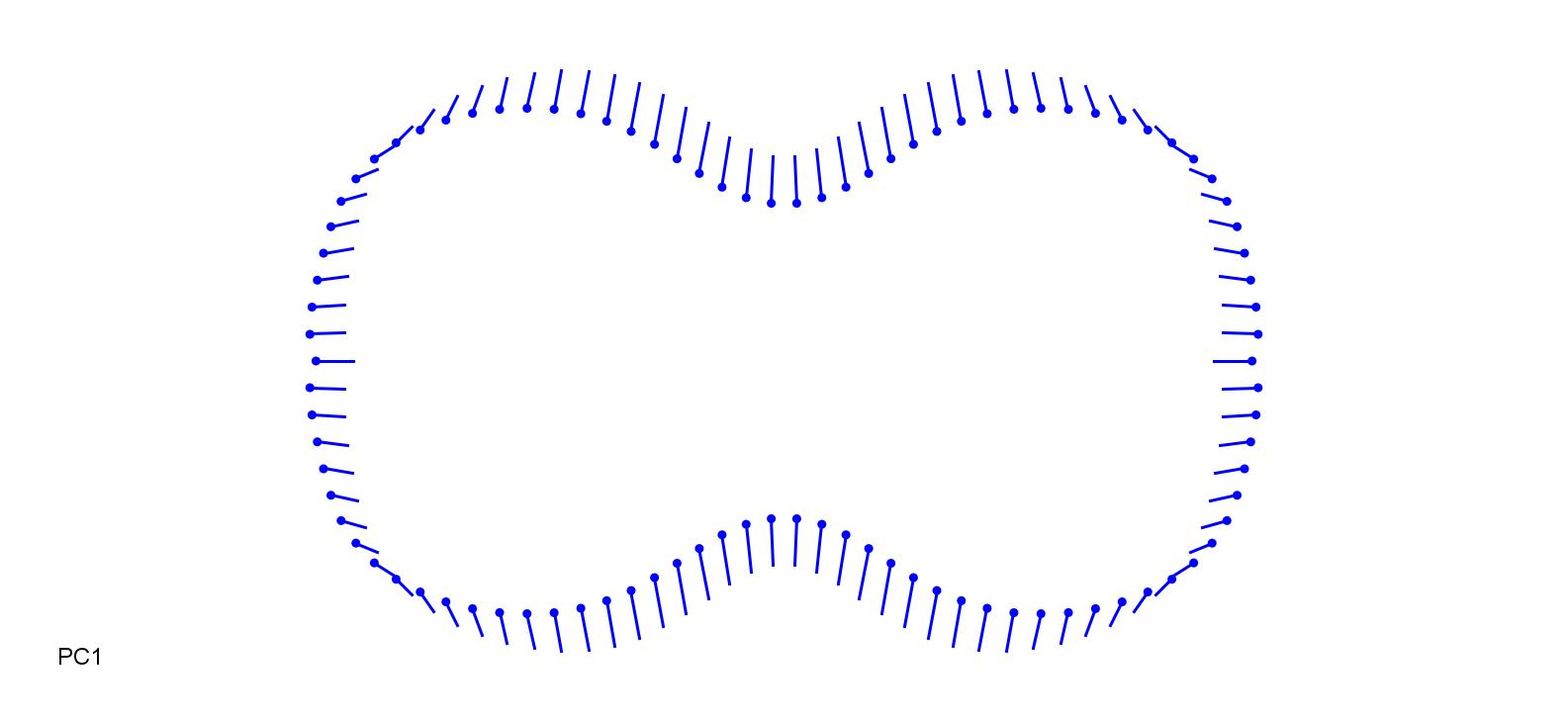

### StiCap-0.18.jpg

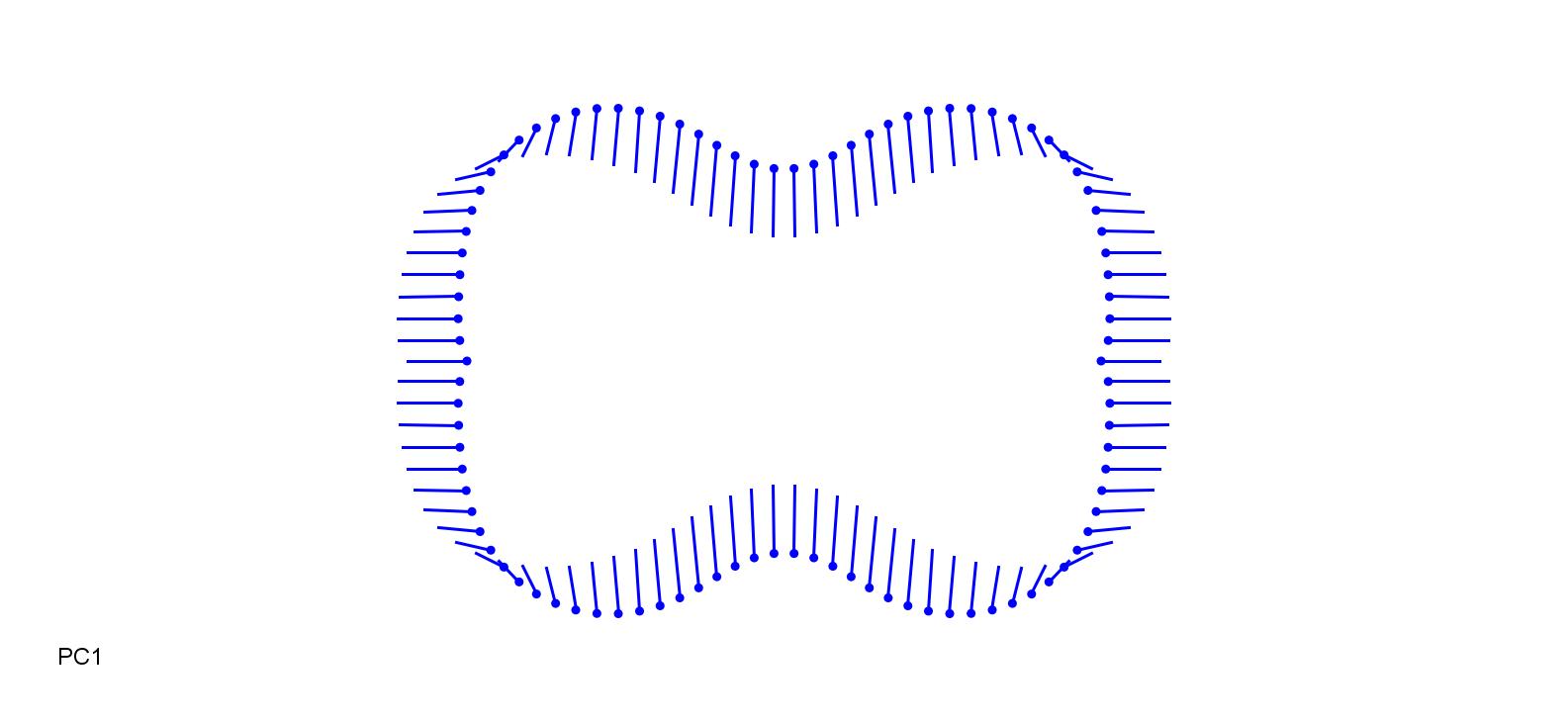

### StiCap_0.18.jpg

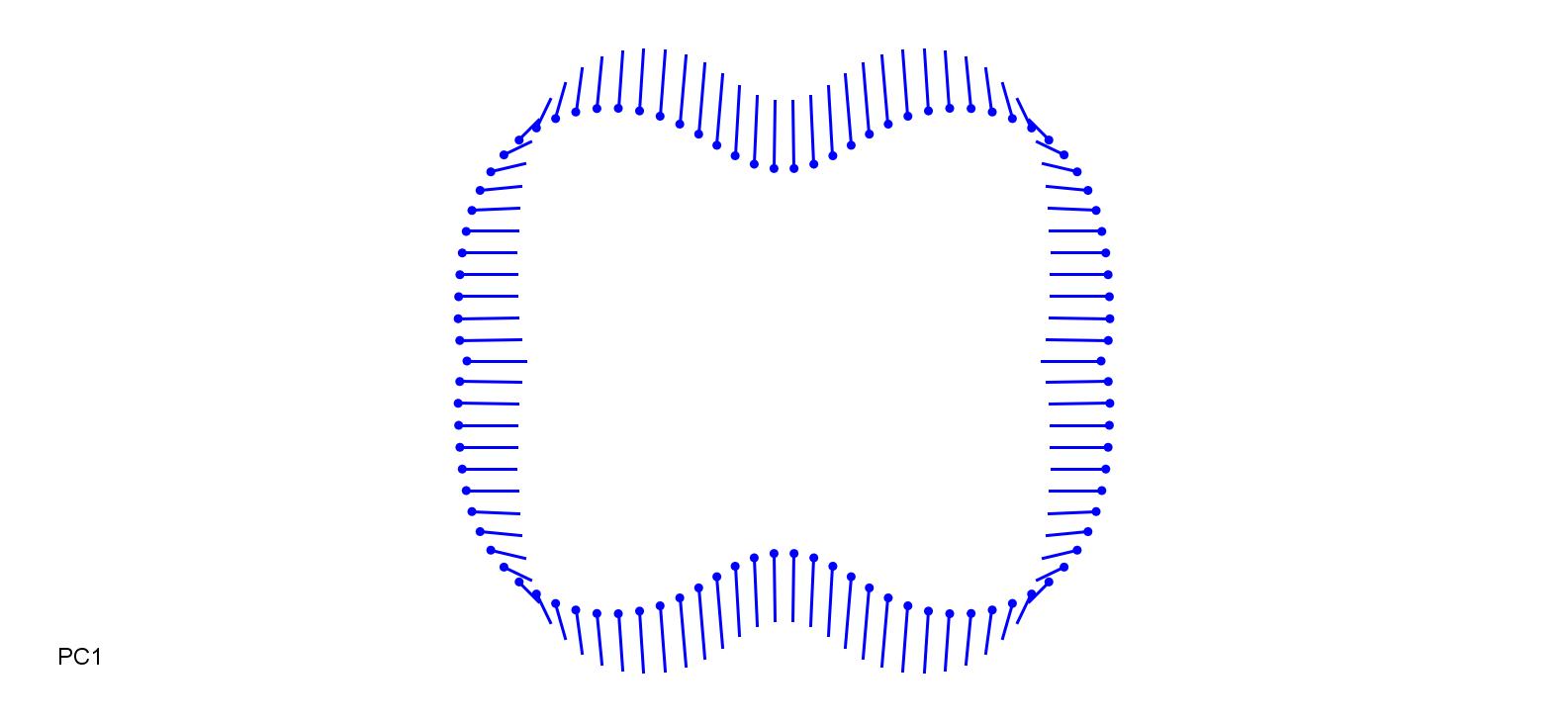

### StiSib-0.15.jpg

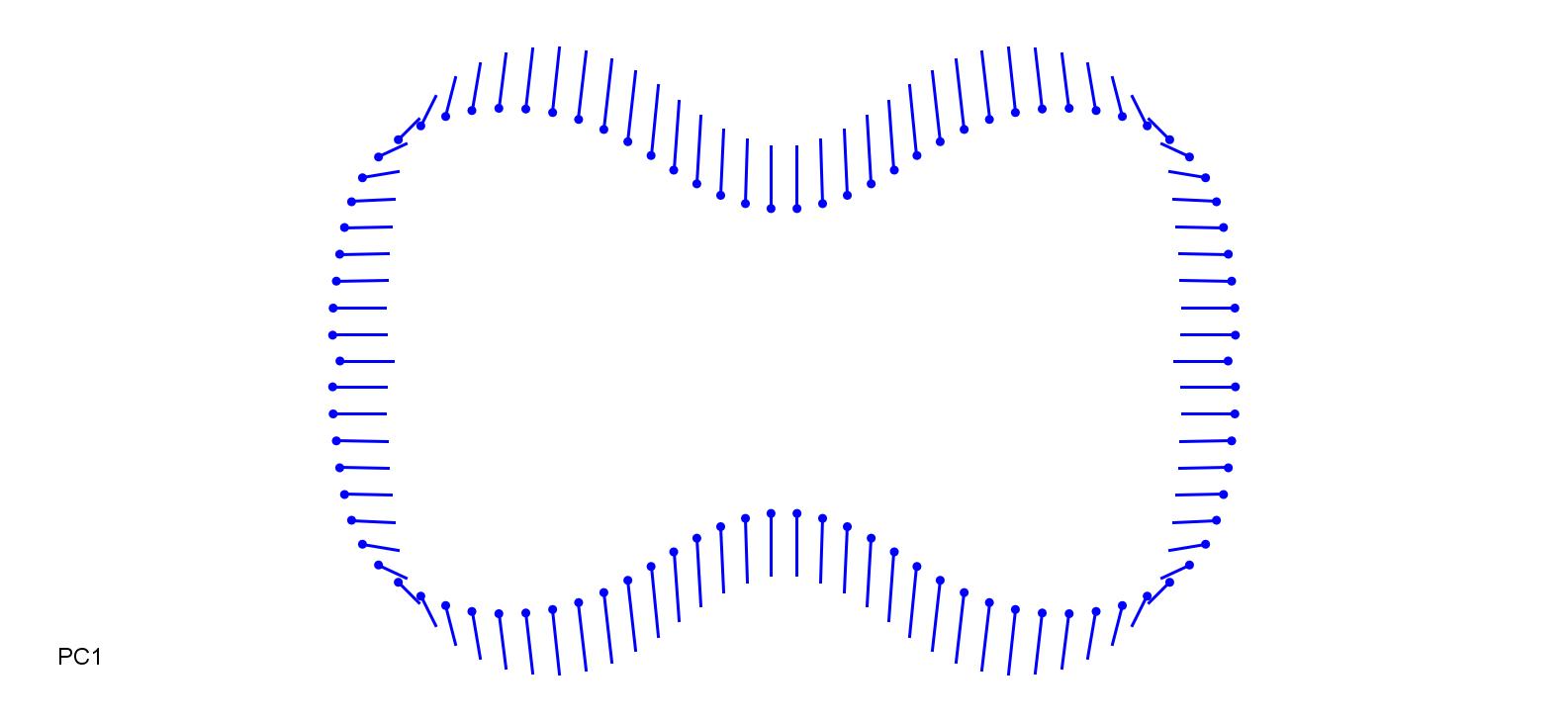

### StiSib_0.15.jpg

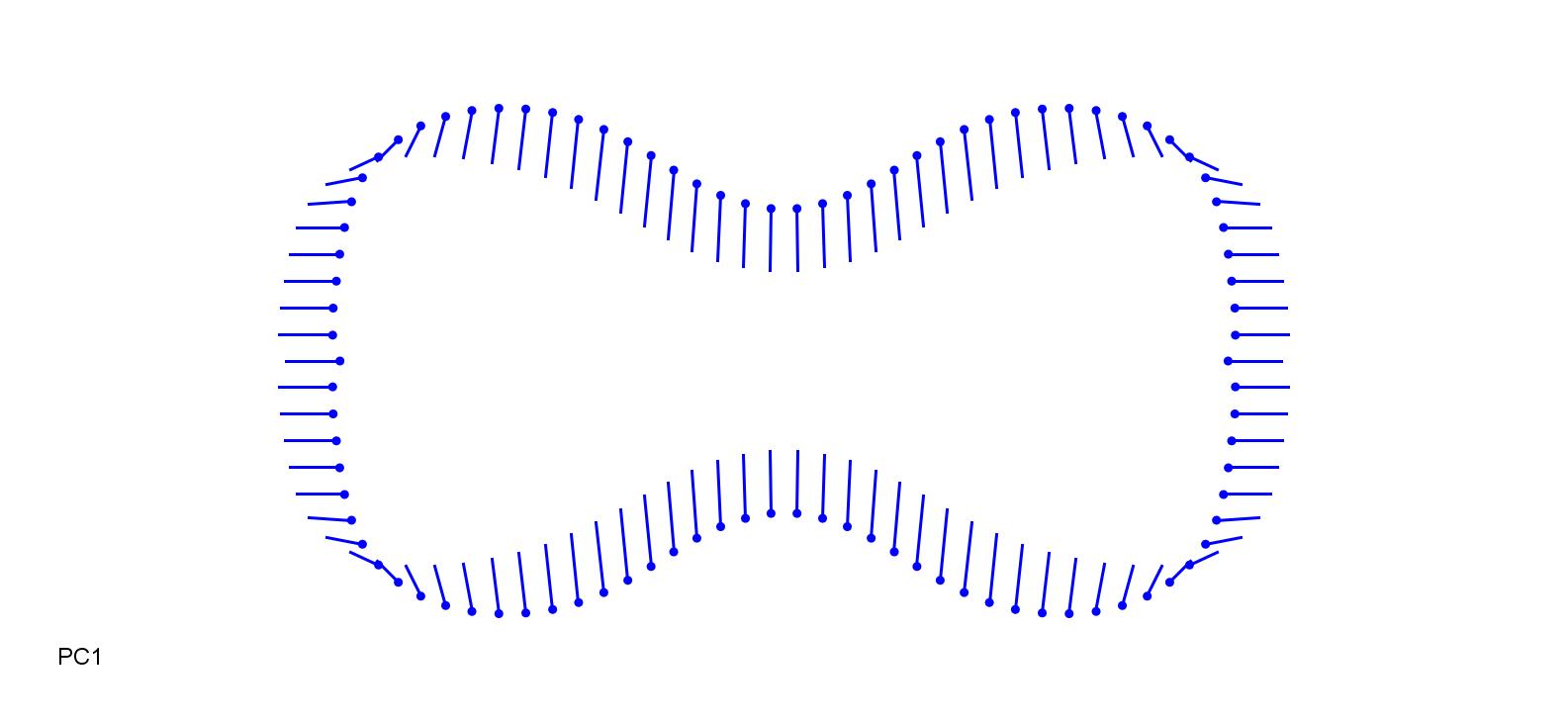

### YuaAlp-0.2.jpg

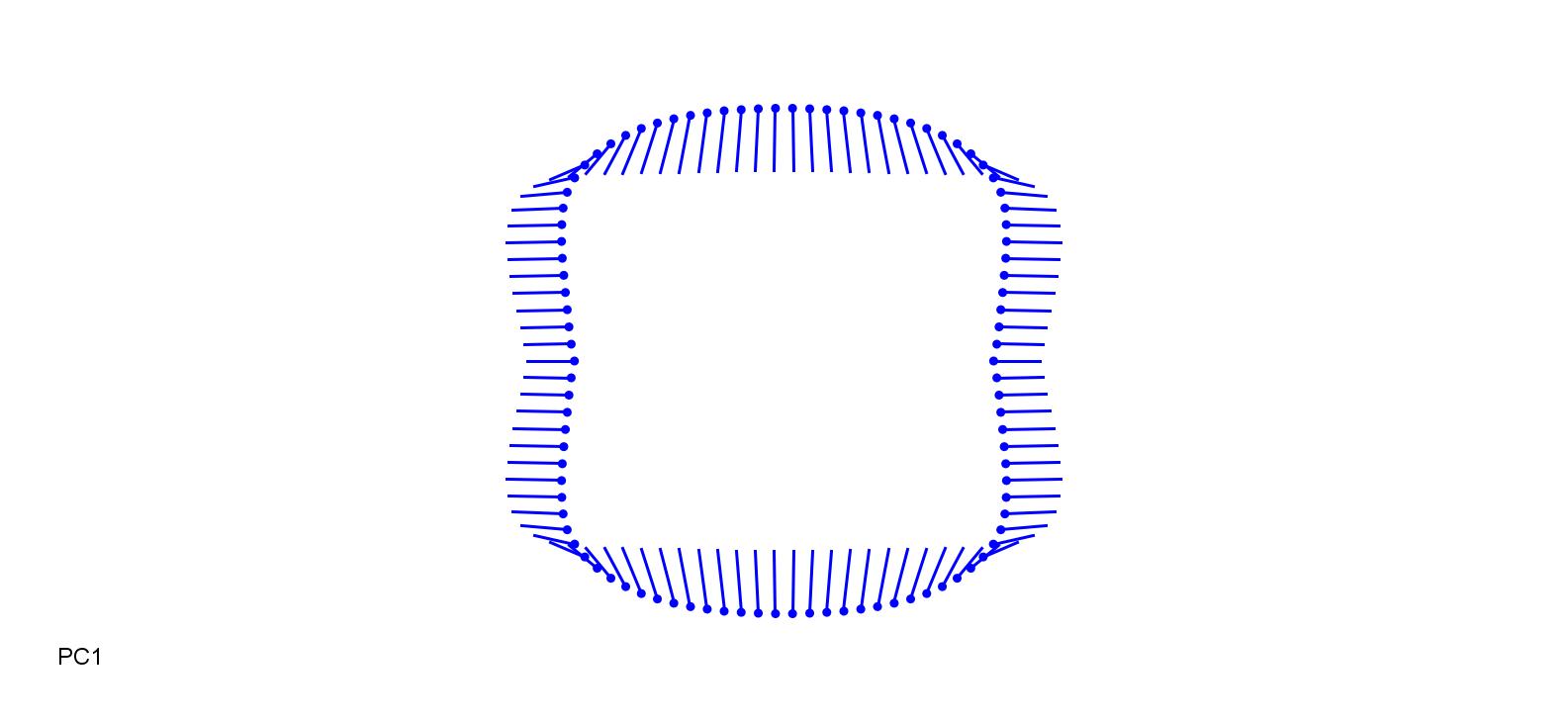

### YuaAlp_0.15.jpg

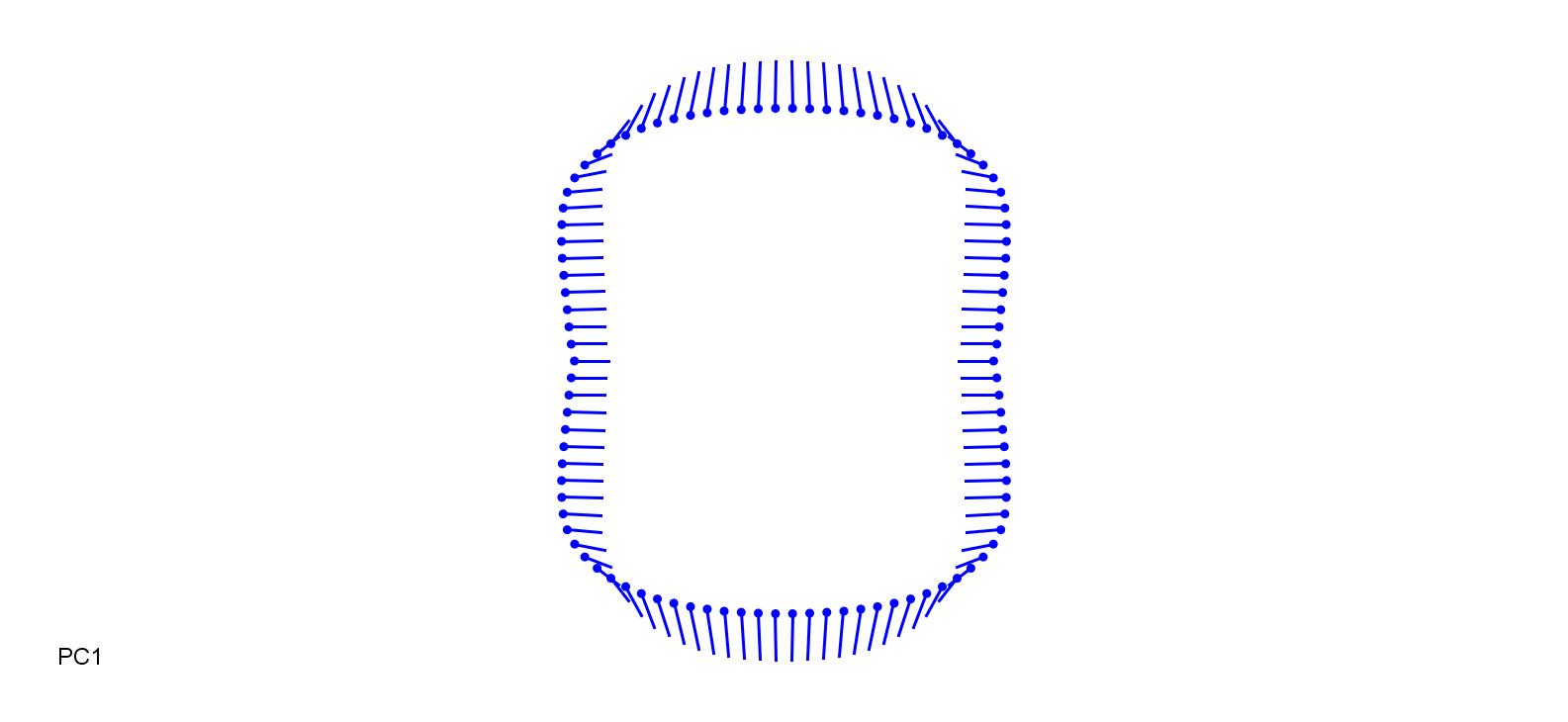
