## Supplementary figures and images for "Phylogenetic, ecological and intraindividual variability patterns in grass phytolith shape"

### AloPrat-0.15.jpg

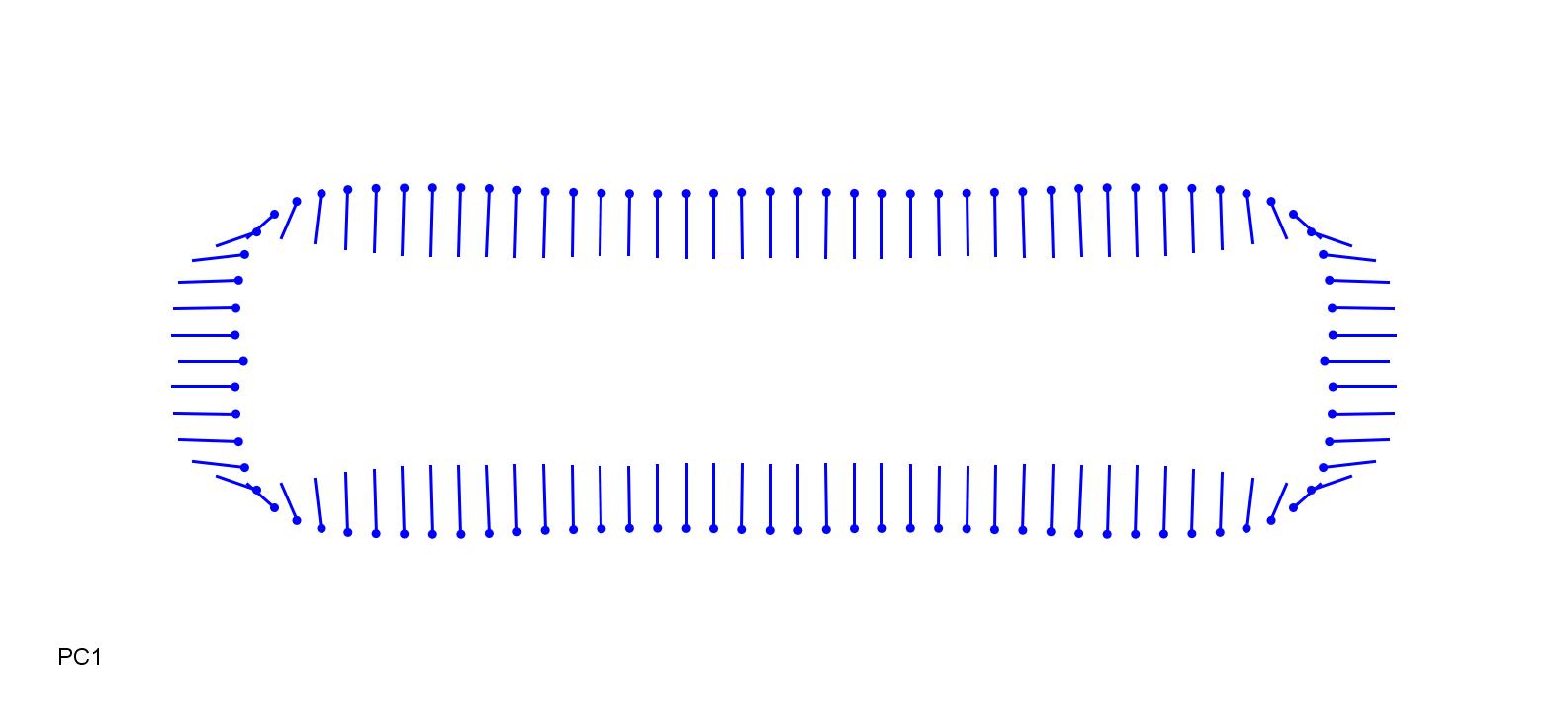

### AloPrat_0.25.jpg

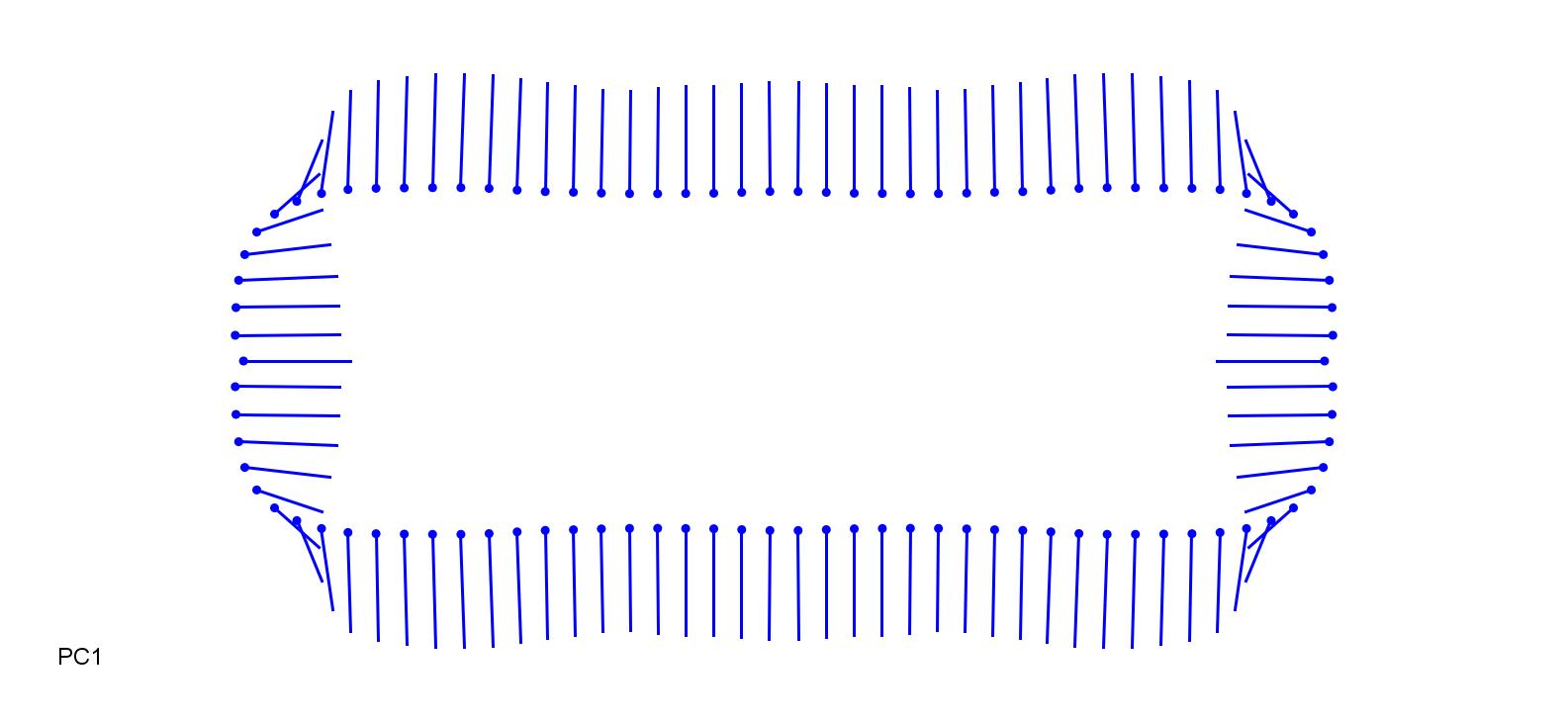

### AriAds-0.15.jpg

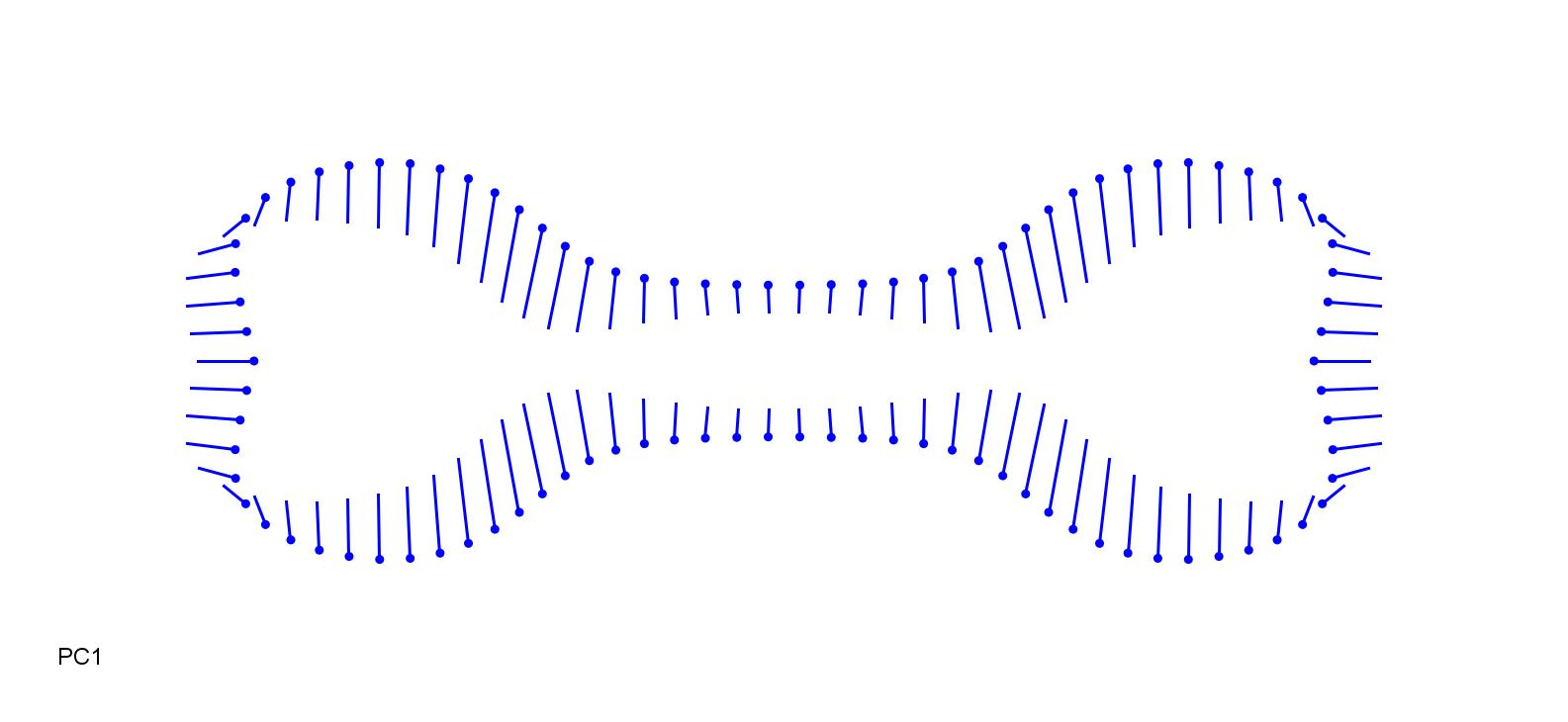

### AriAds_0.2.jpg

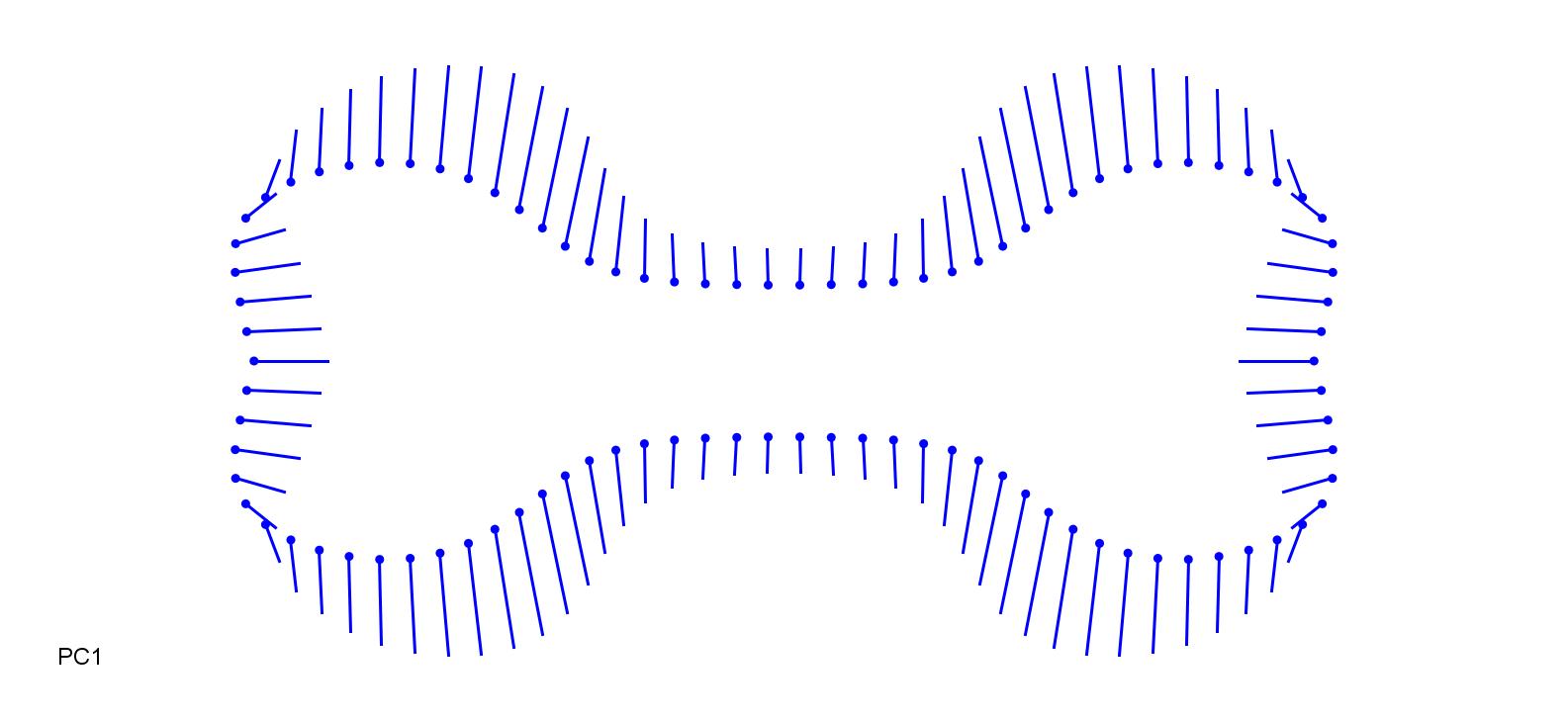

### AriCon-0.05.jpg

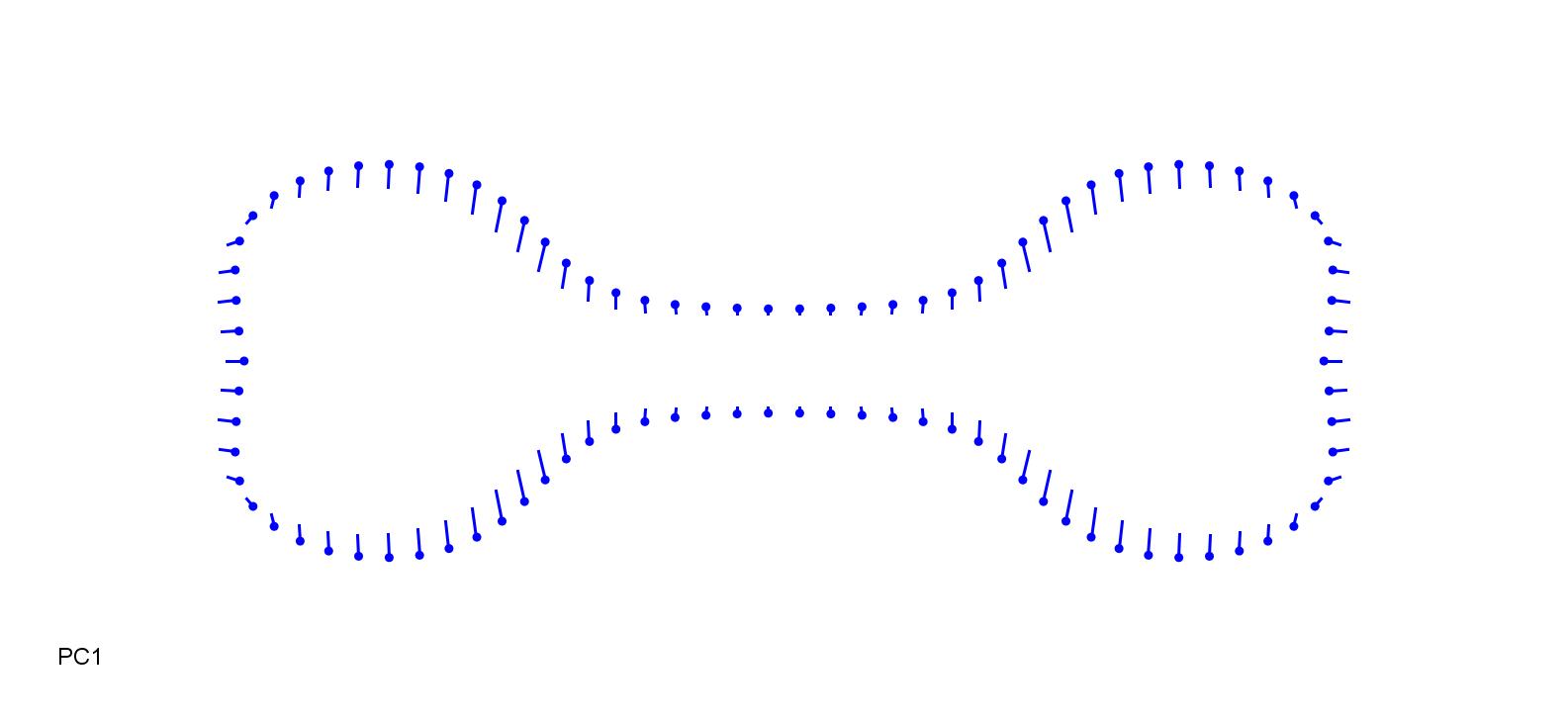

### AriCon_0.1.jpg

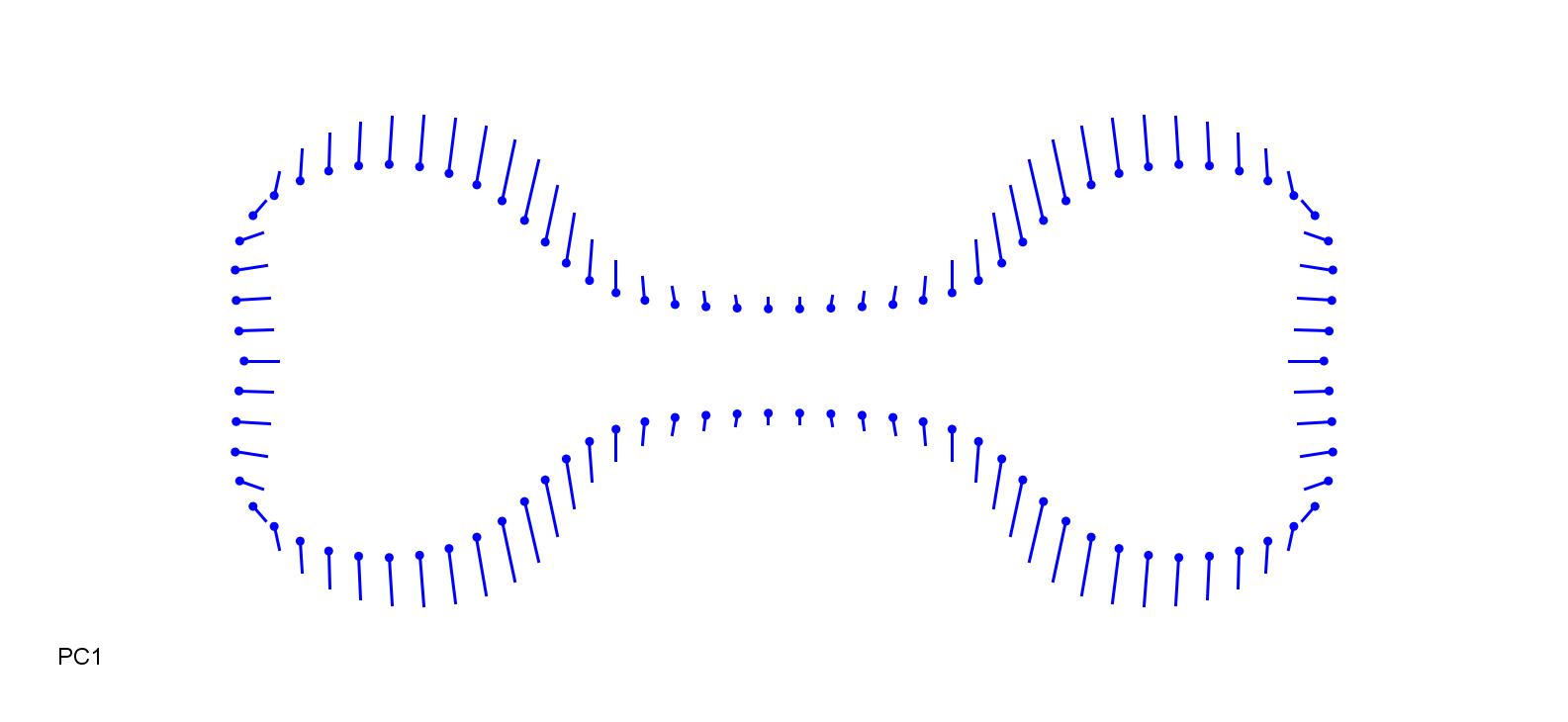

### AriRhi-0.1.jpg

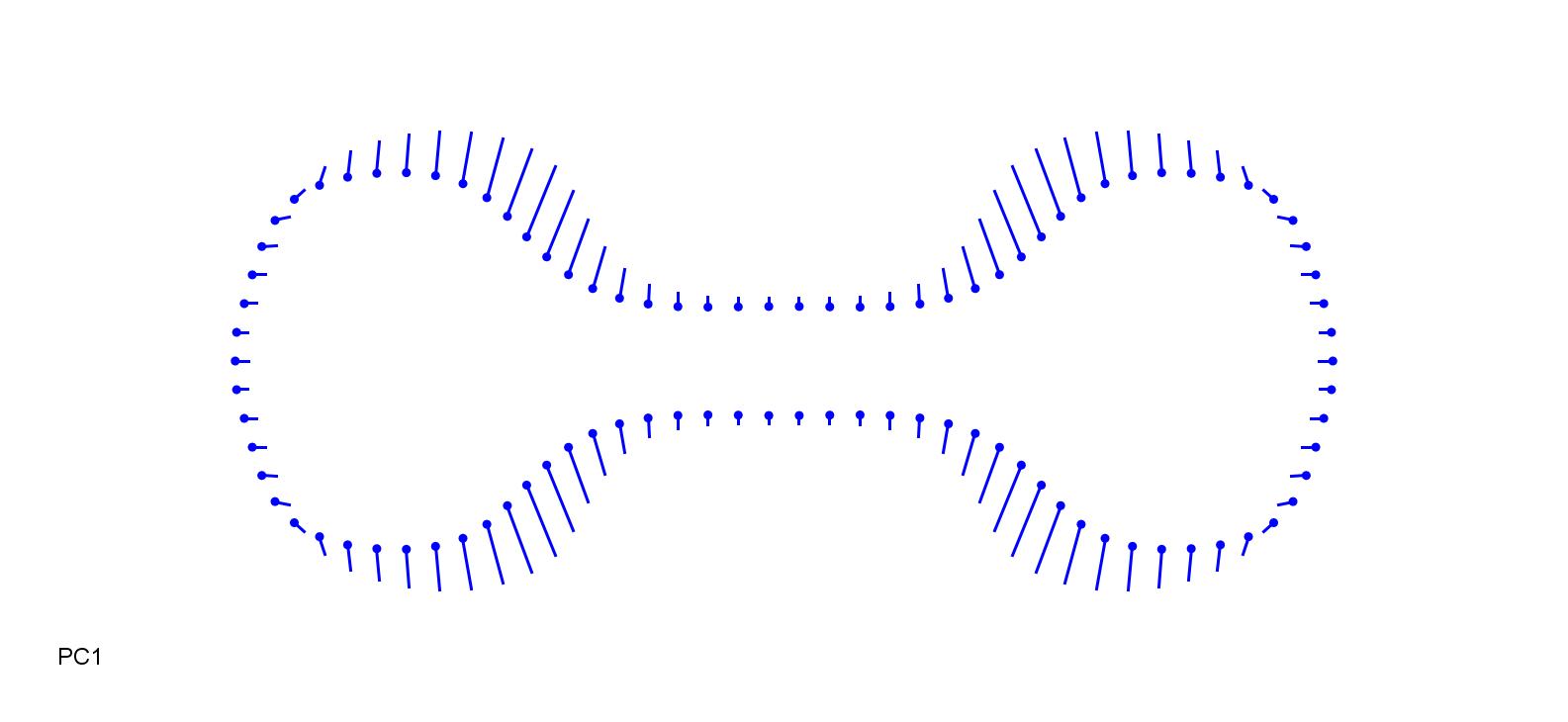

### AriRhi_0.1.jpg

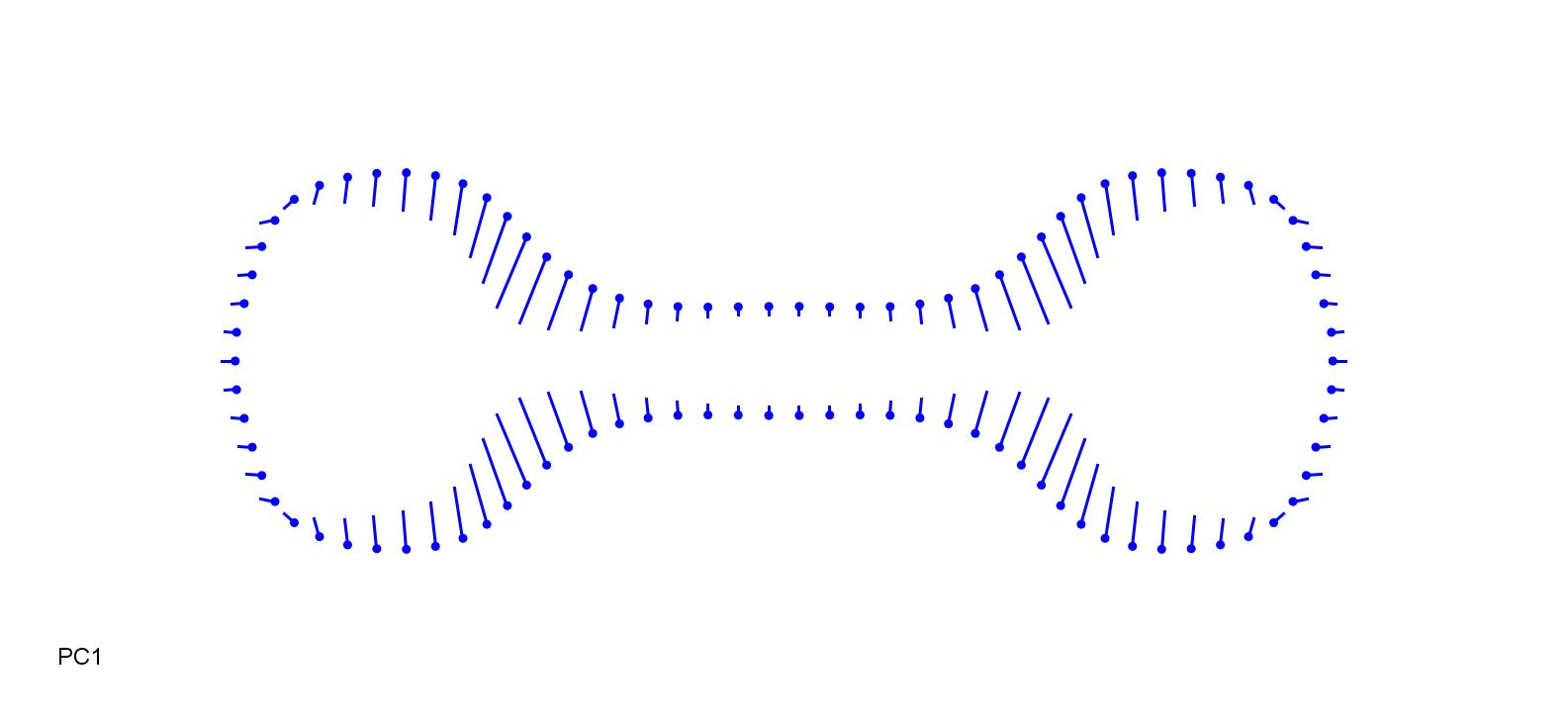

### AruDon-0.1.jpg

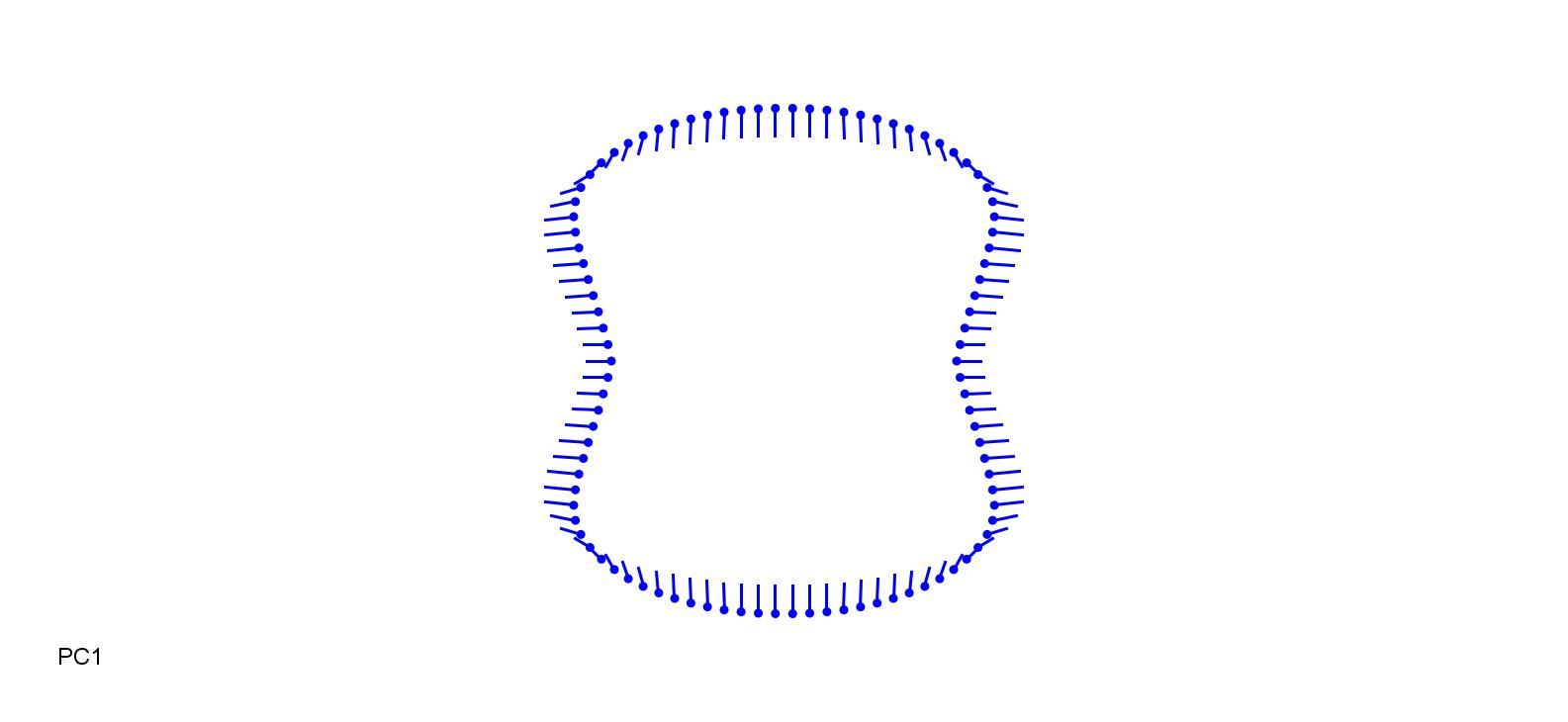

### AruDon_0.1.jpg

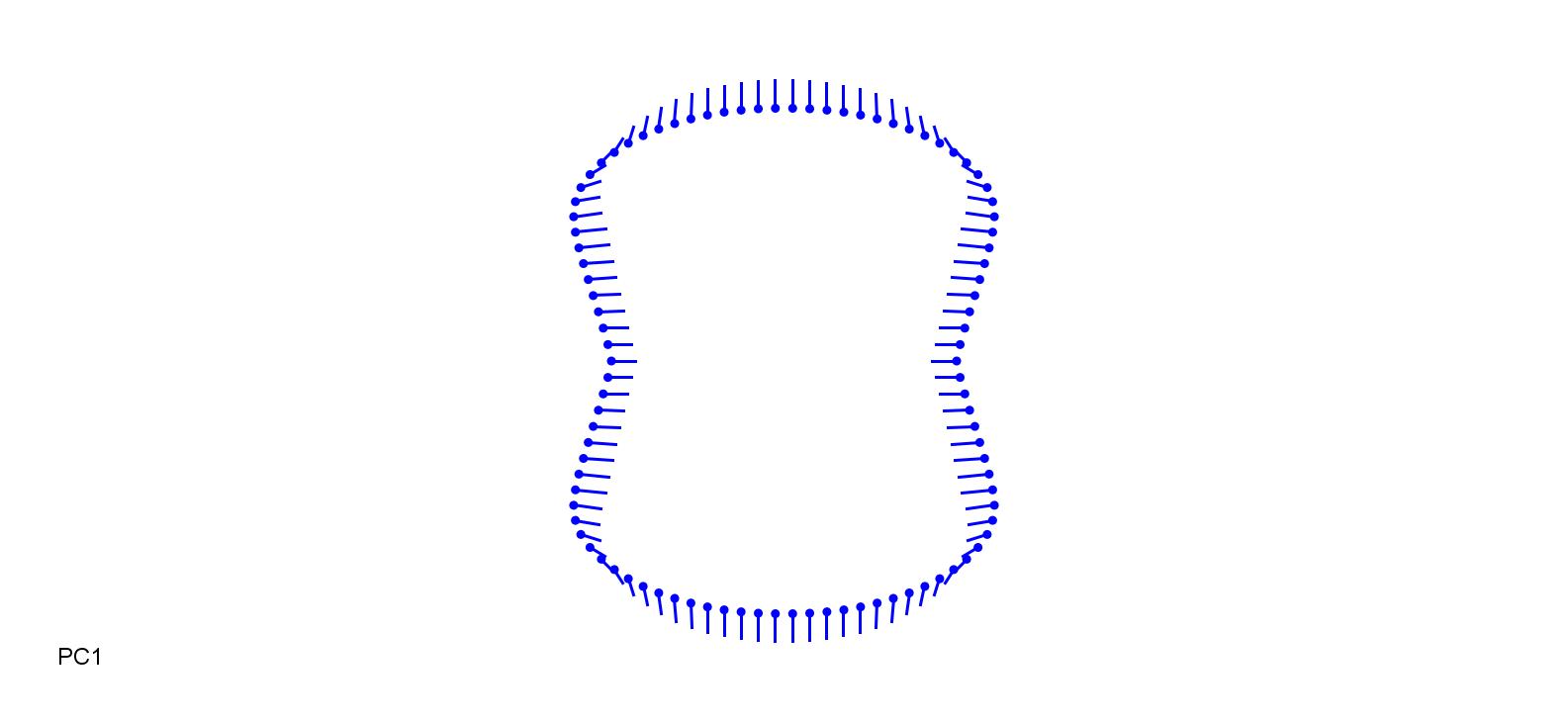

### AruPli-0.2.jpg

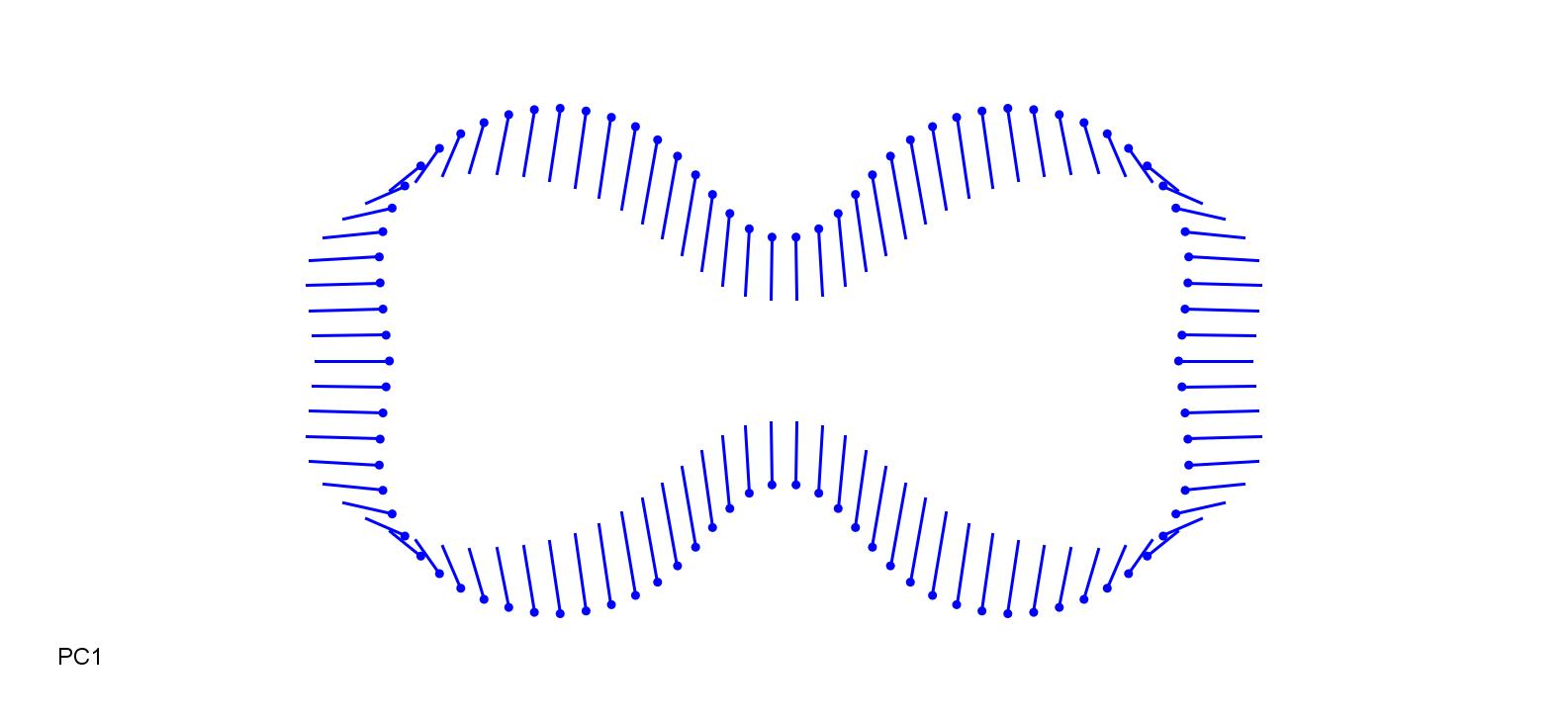

### AruPli_0.2.jpg

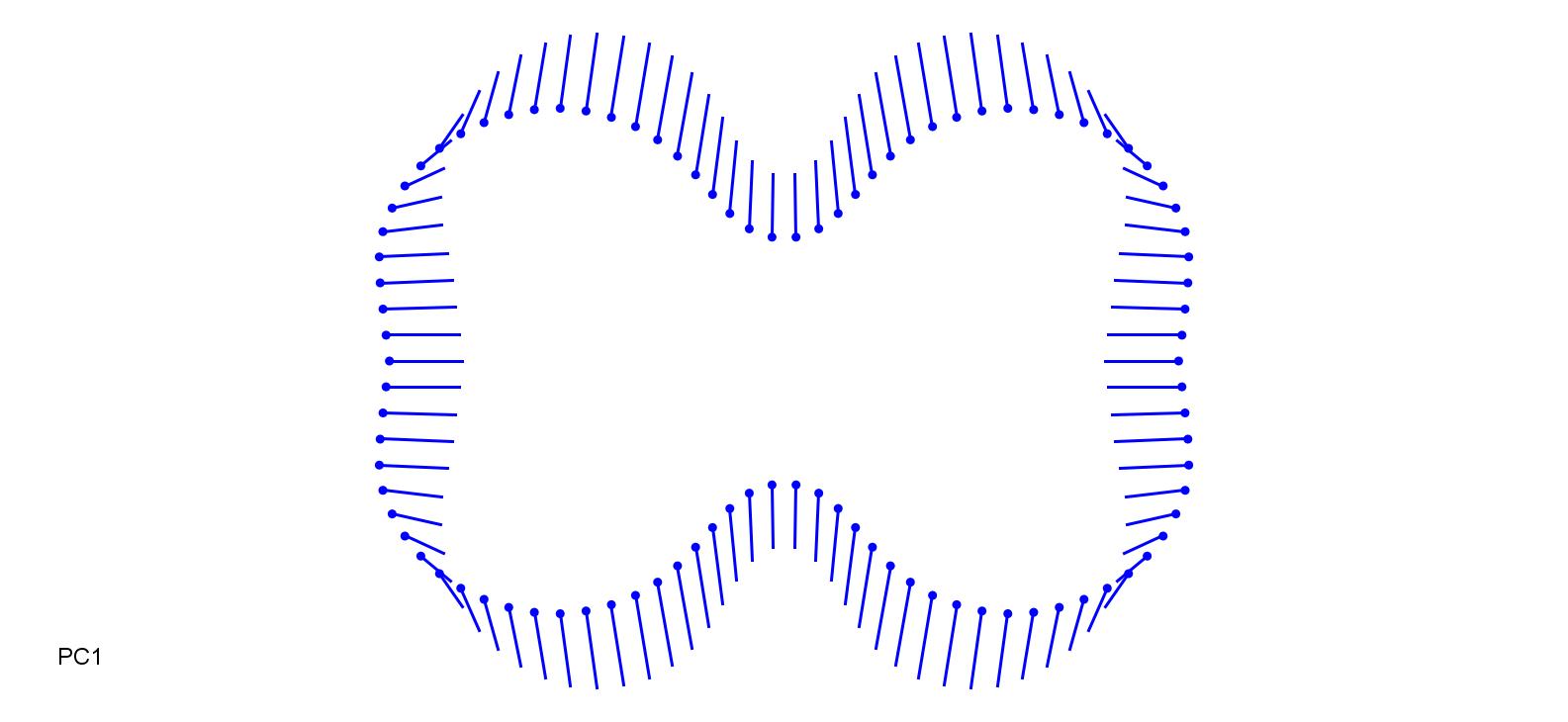

### BotIsc-0.2.jpg

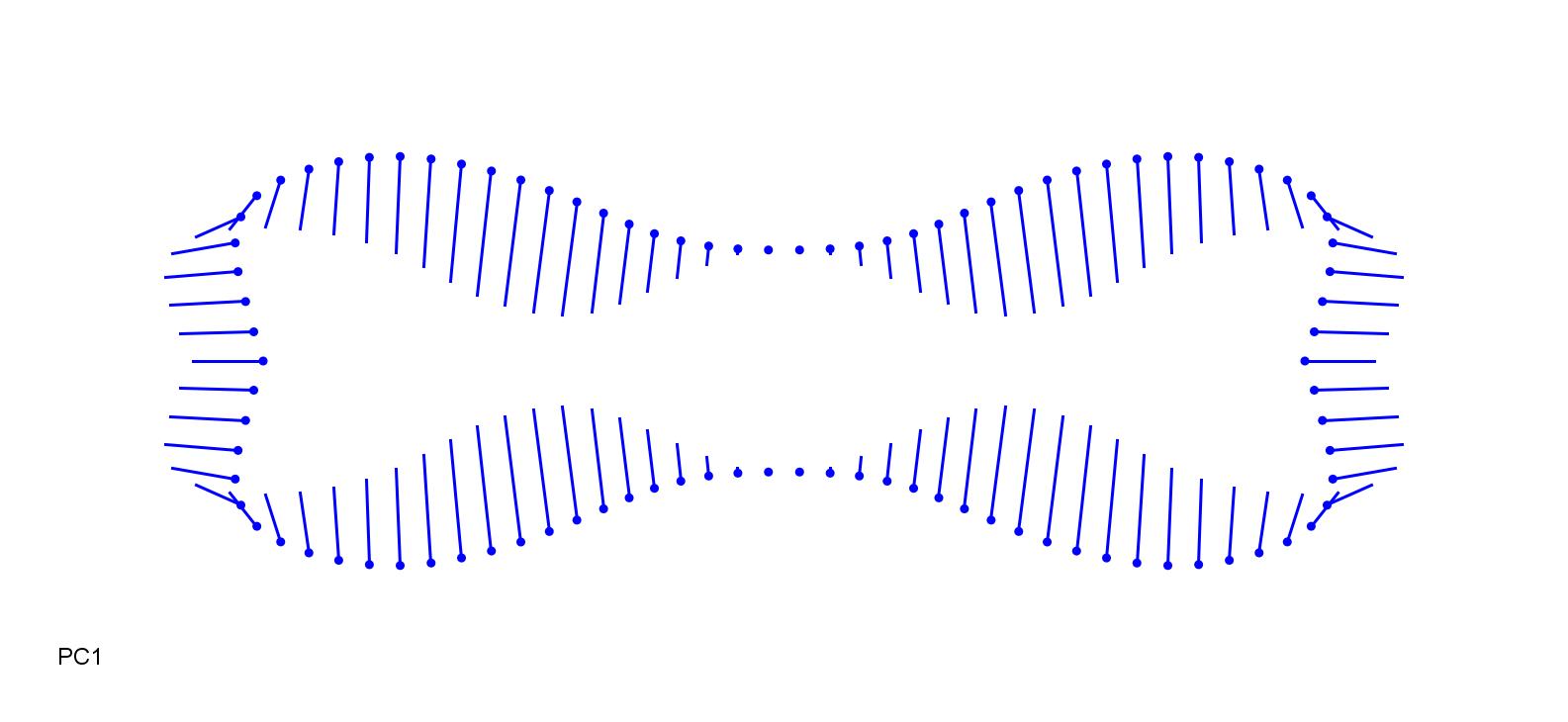

### BotIsc_0.25.jpg

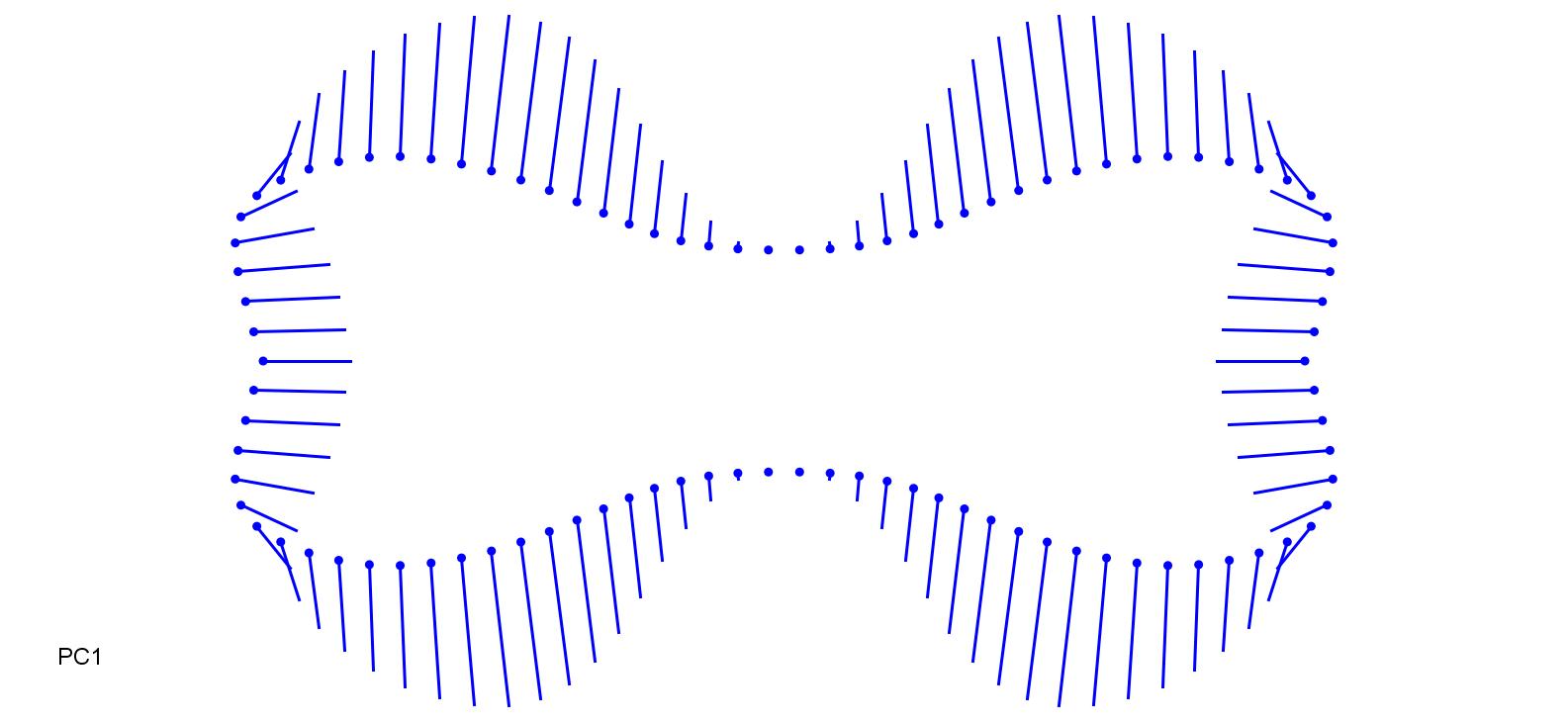

### CalAru-0.1.jpg

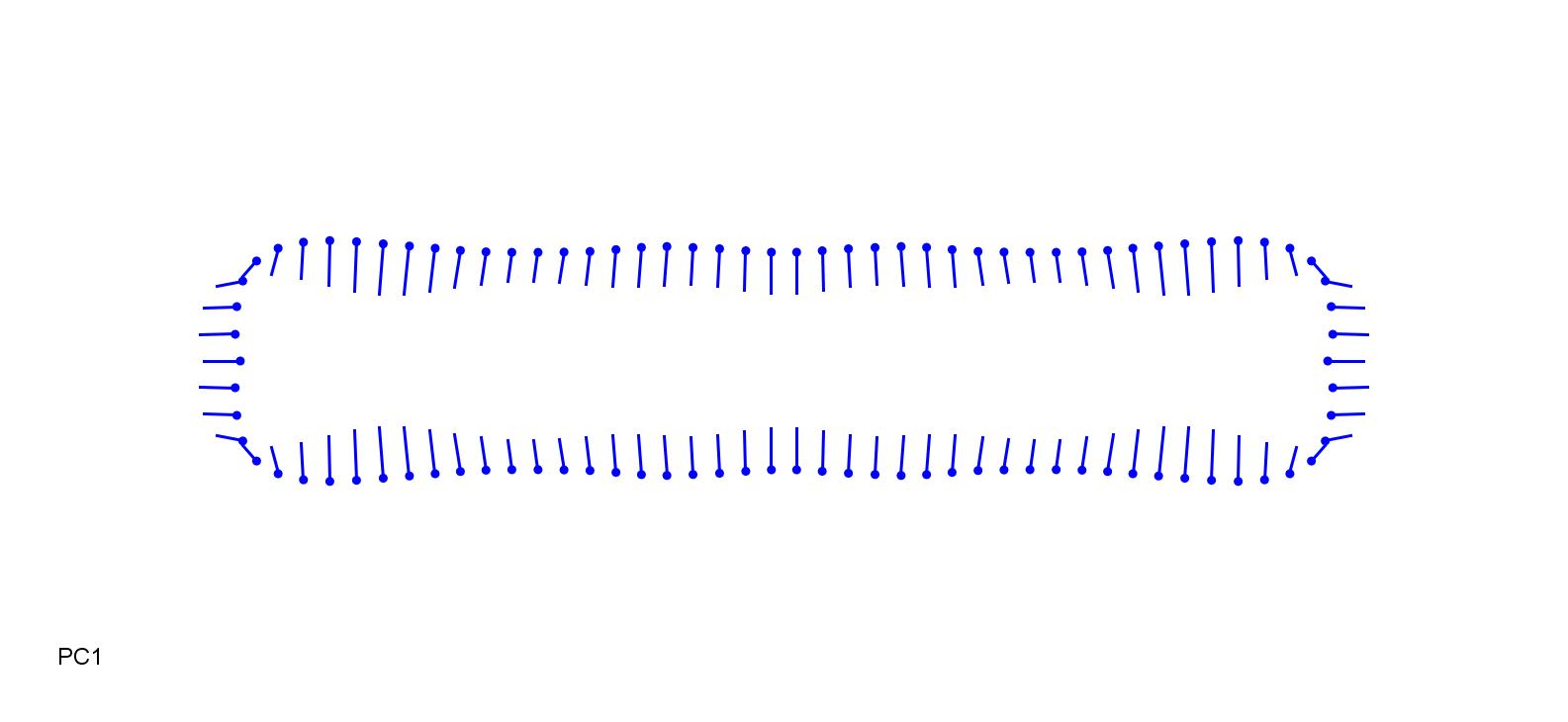

### CalAru_0.1.jpg

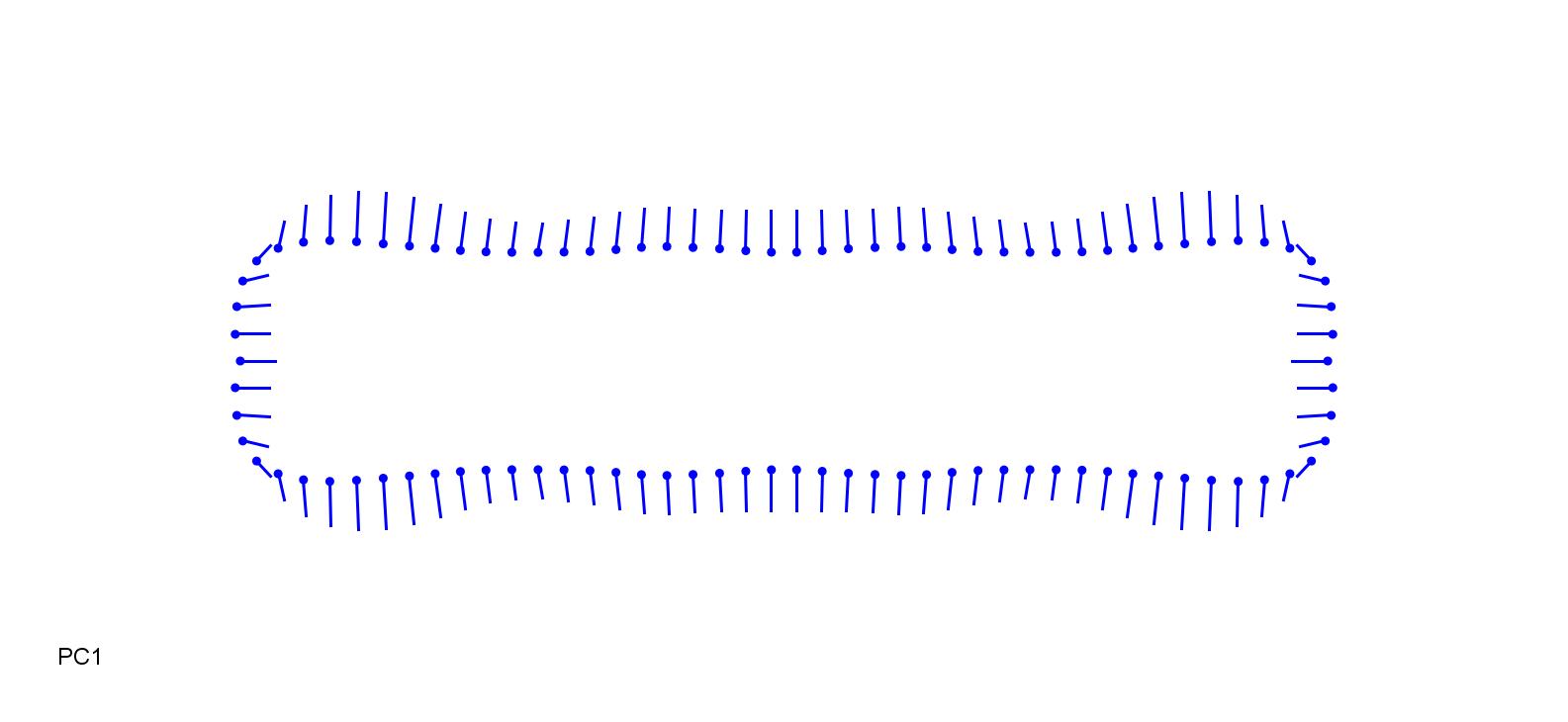

### CalCan-0.15.jpg

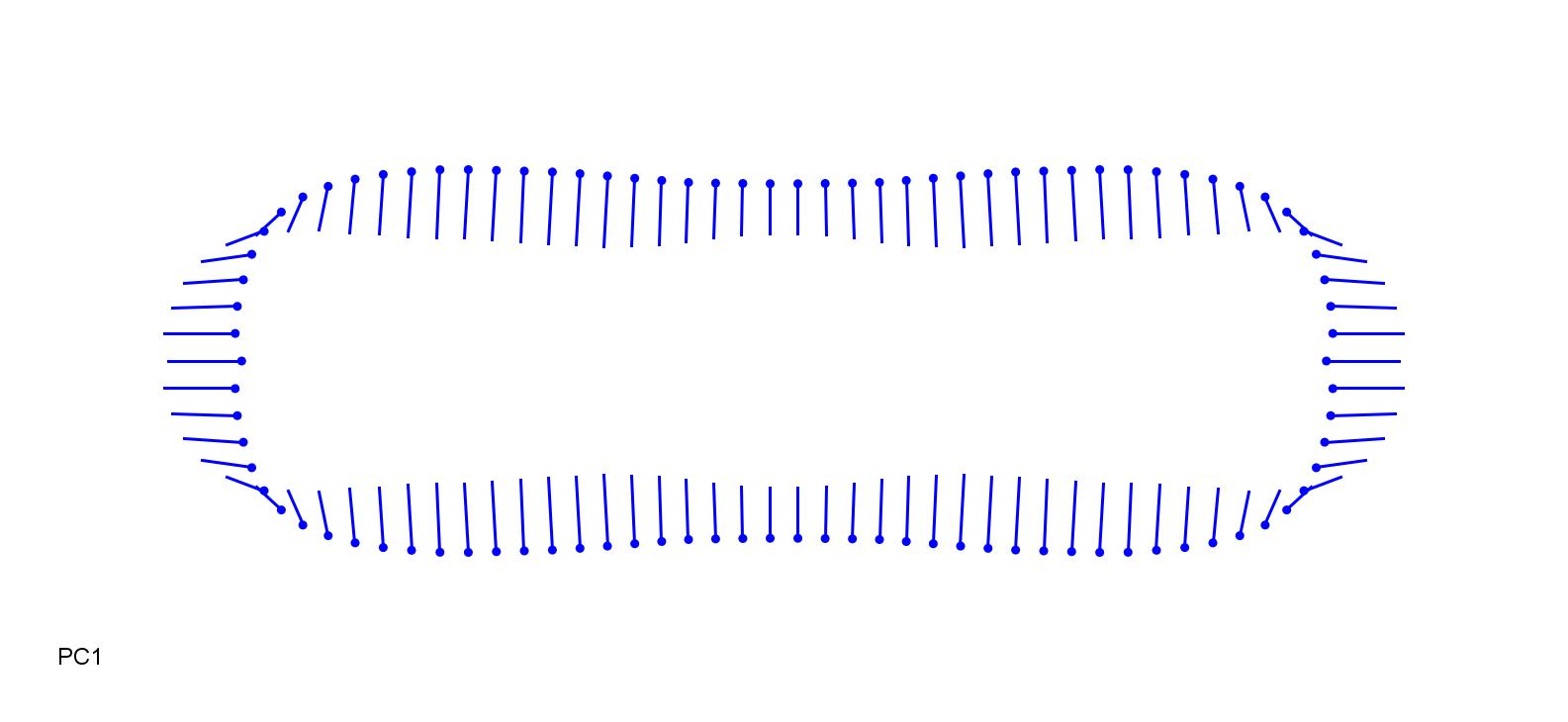

### CalCan_0.15.jpg

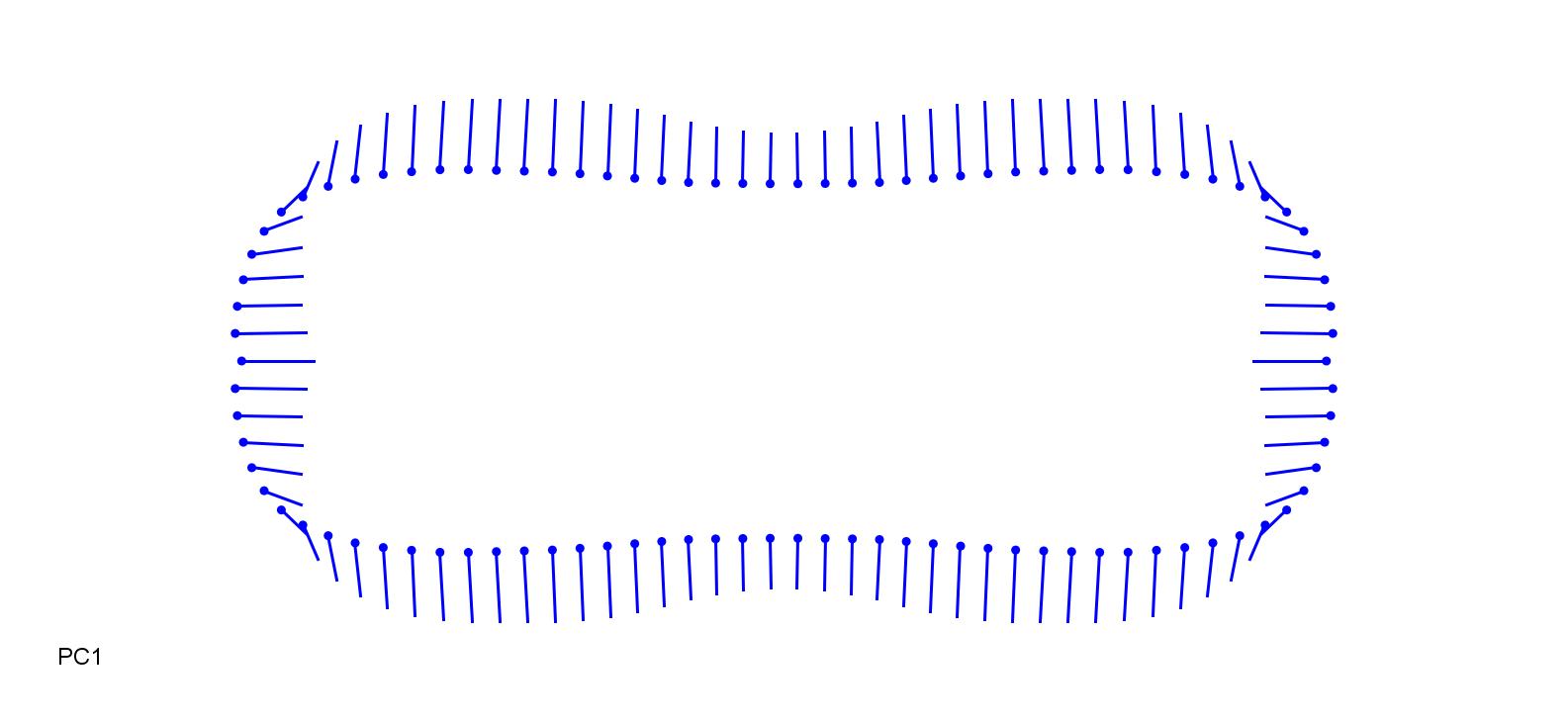

### CalEpi-0.2.jpg

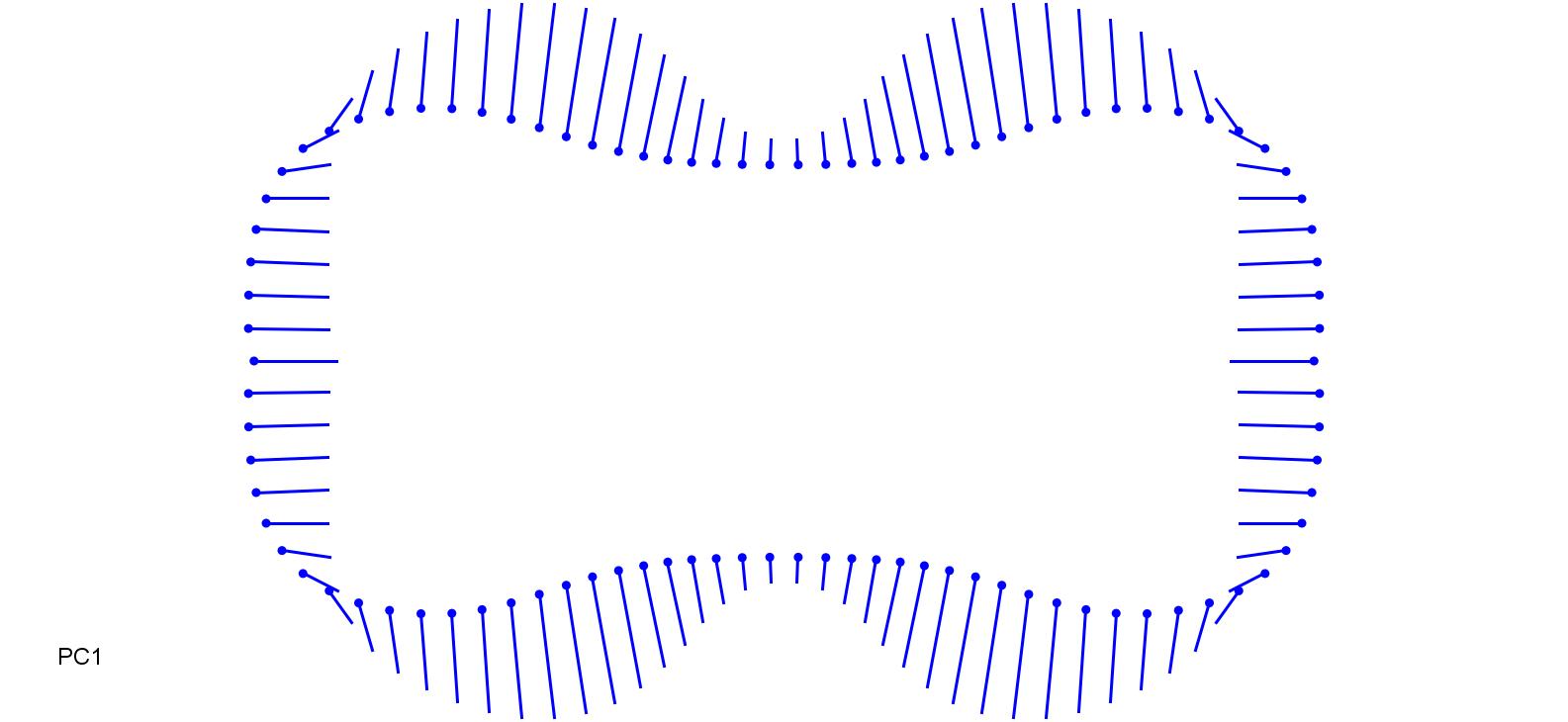

### CalEpi_0.2.jpg

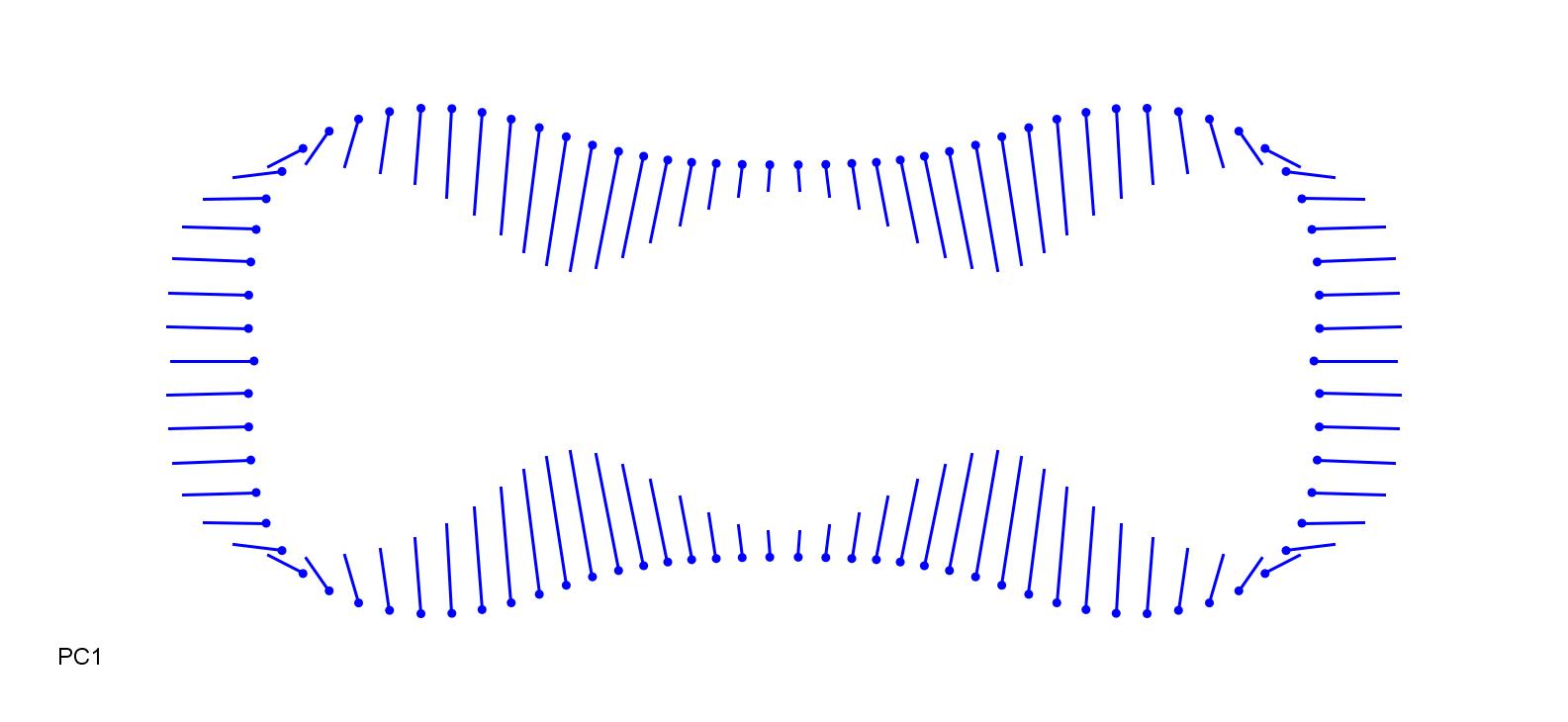

### CalVil-0.15.jpg

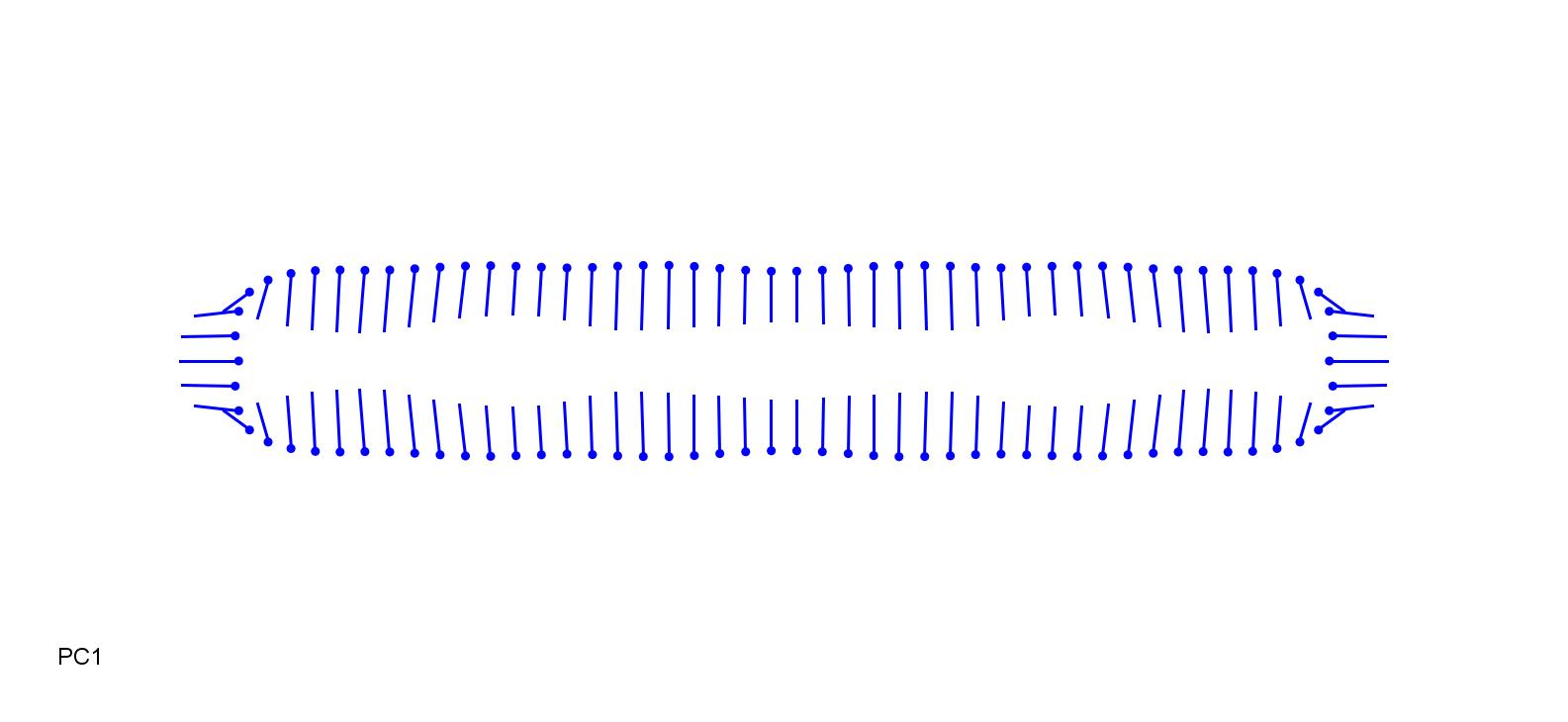

### CalVil_0.15.jpg

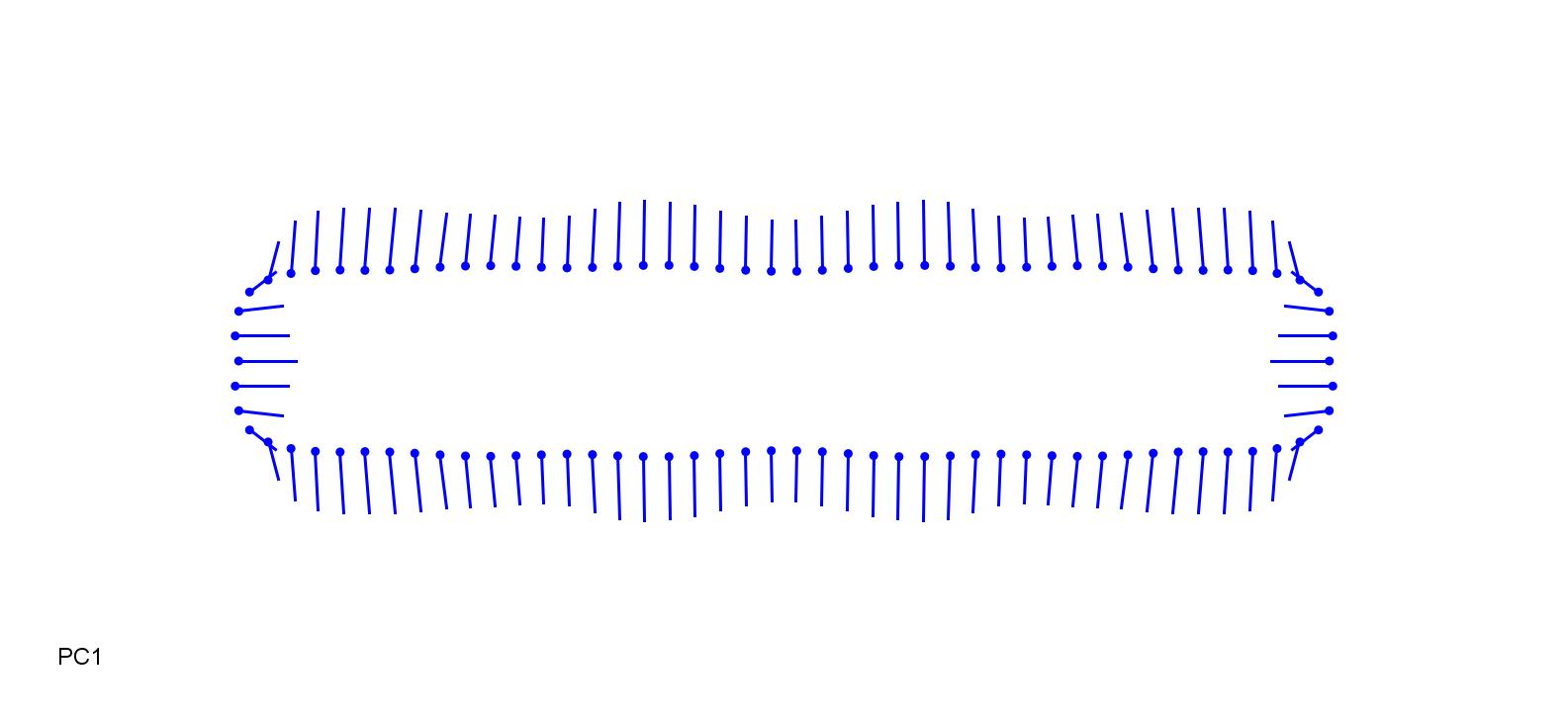

### CoiLac-0.2.jpg

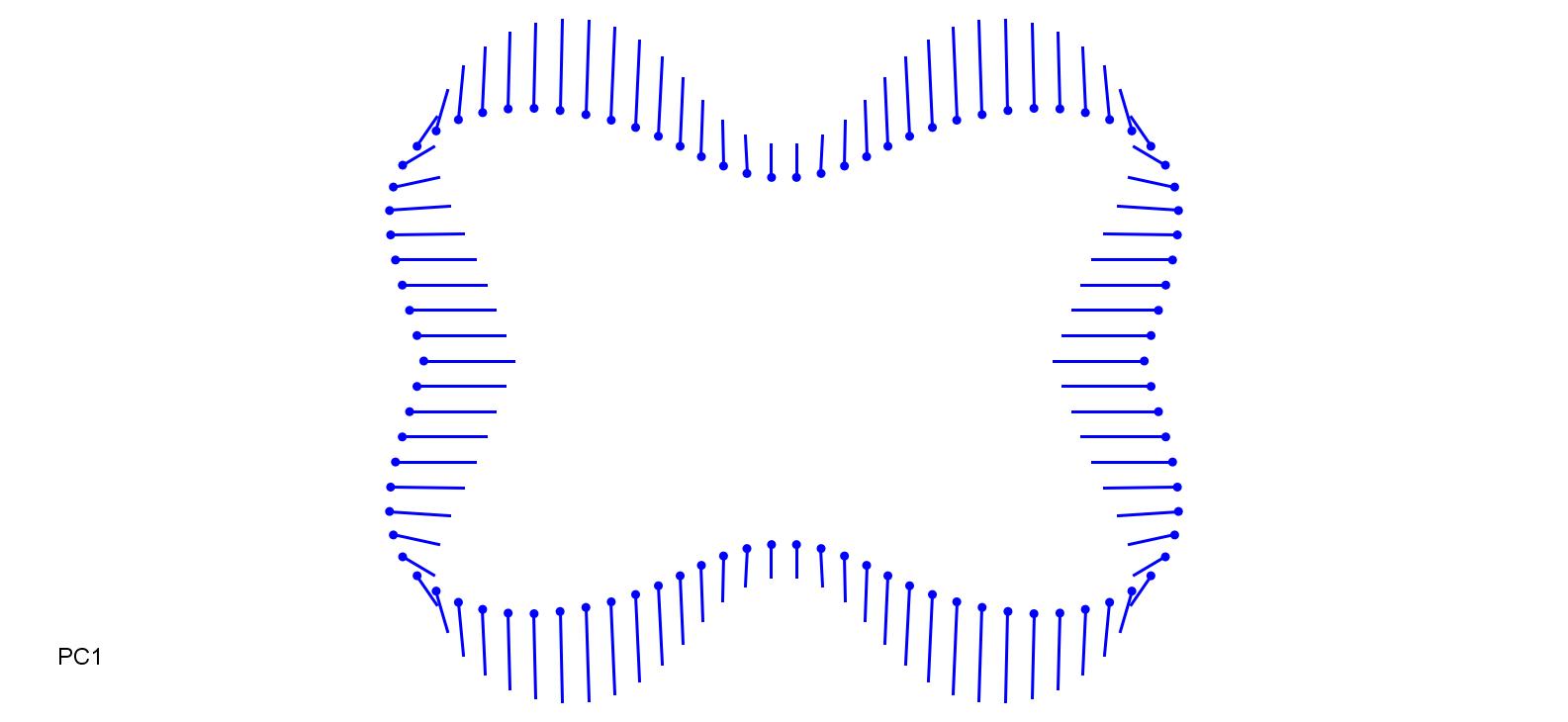

### CoiLac_0.25.jpg

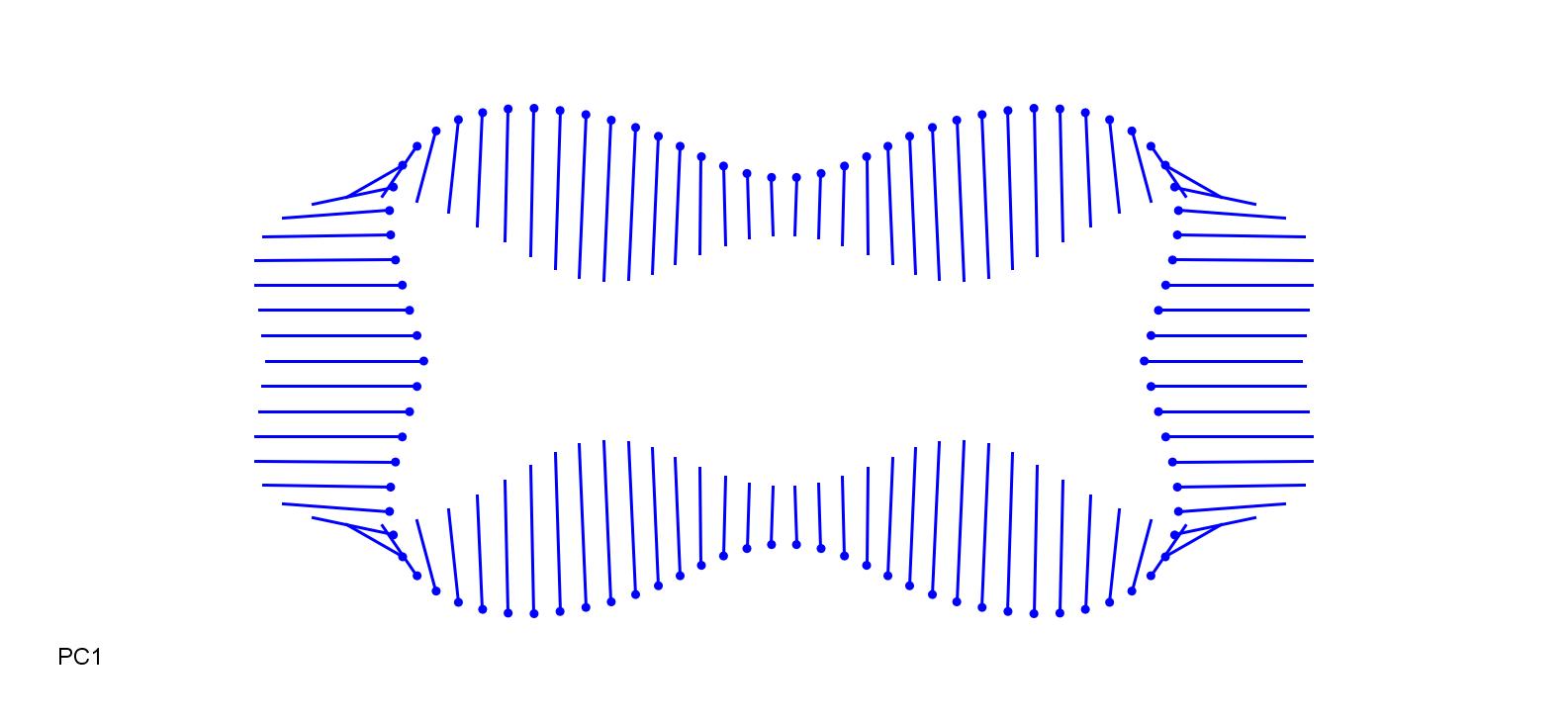

### CynDac-0.1.jpg

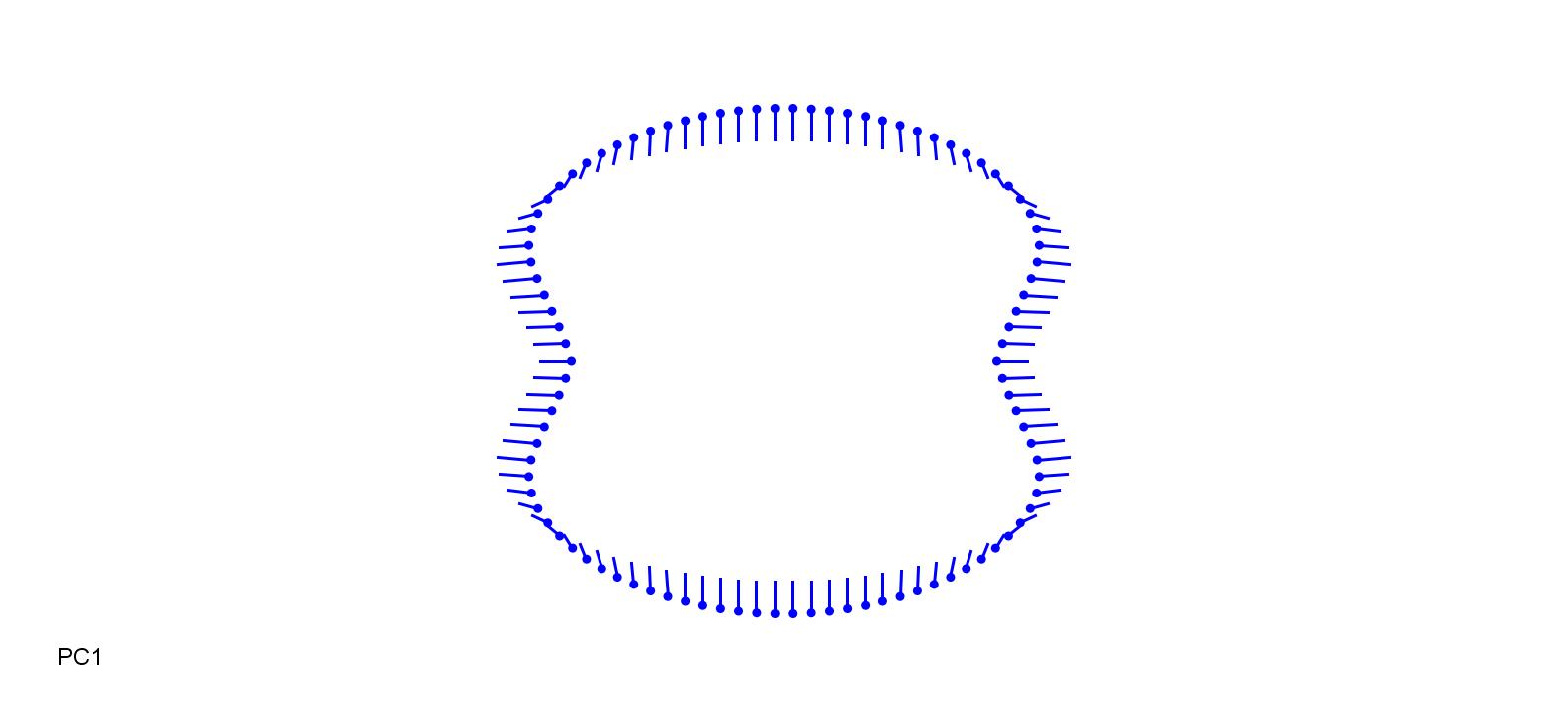

### CynDac_0.15.jpg

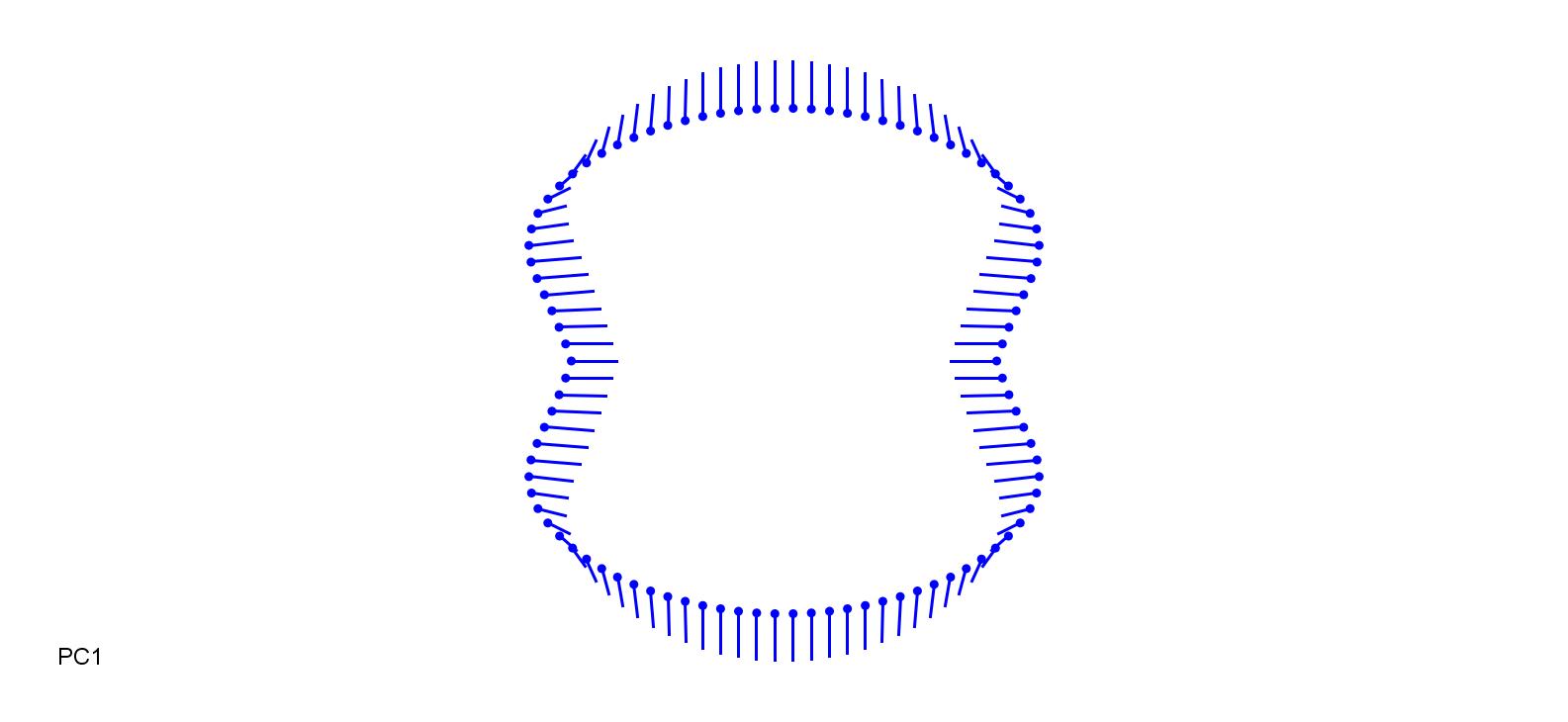

### DacAeg-0.13.jpg

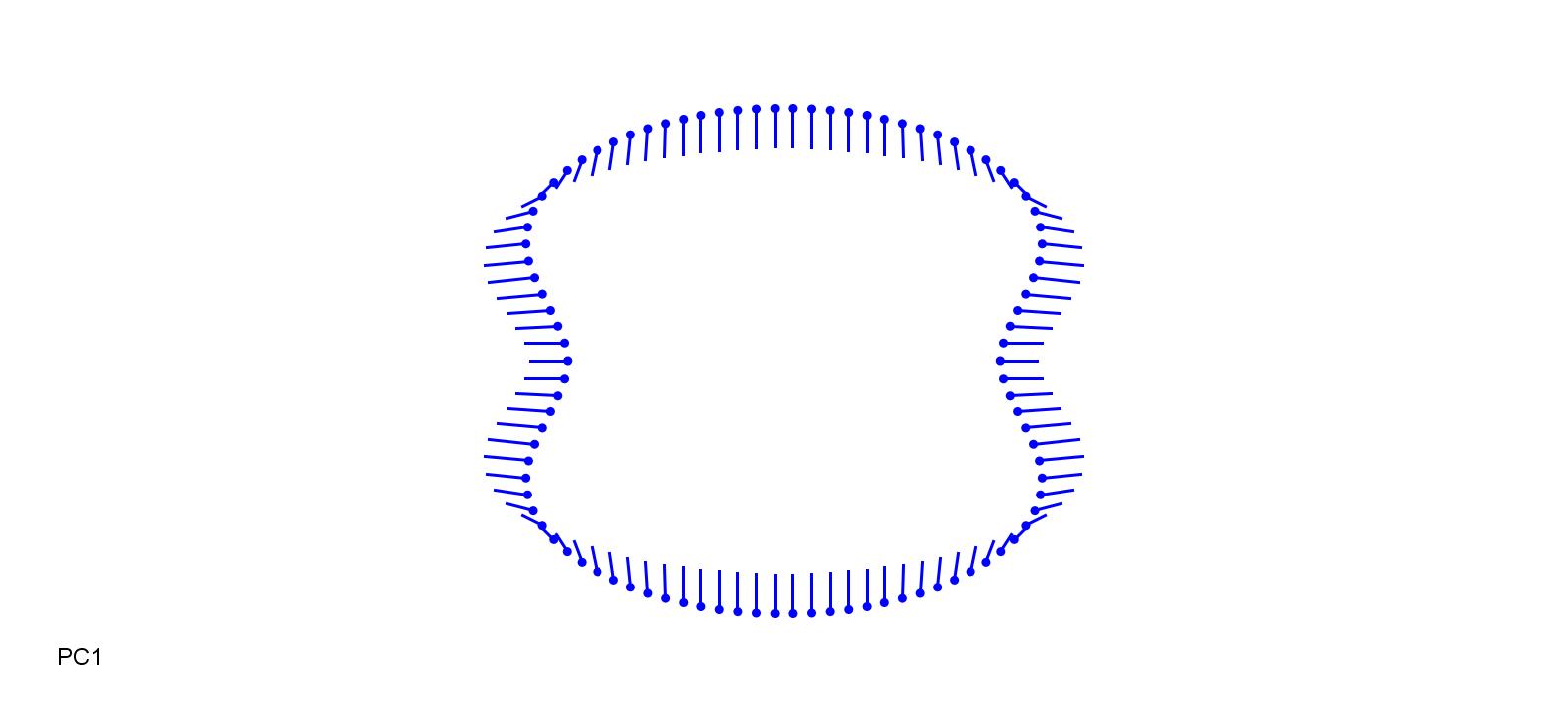

### DacAeg_0.13.jpg

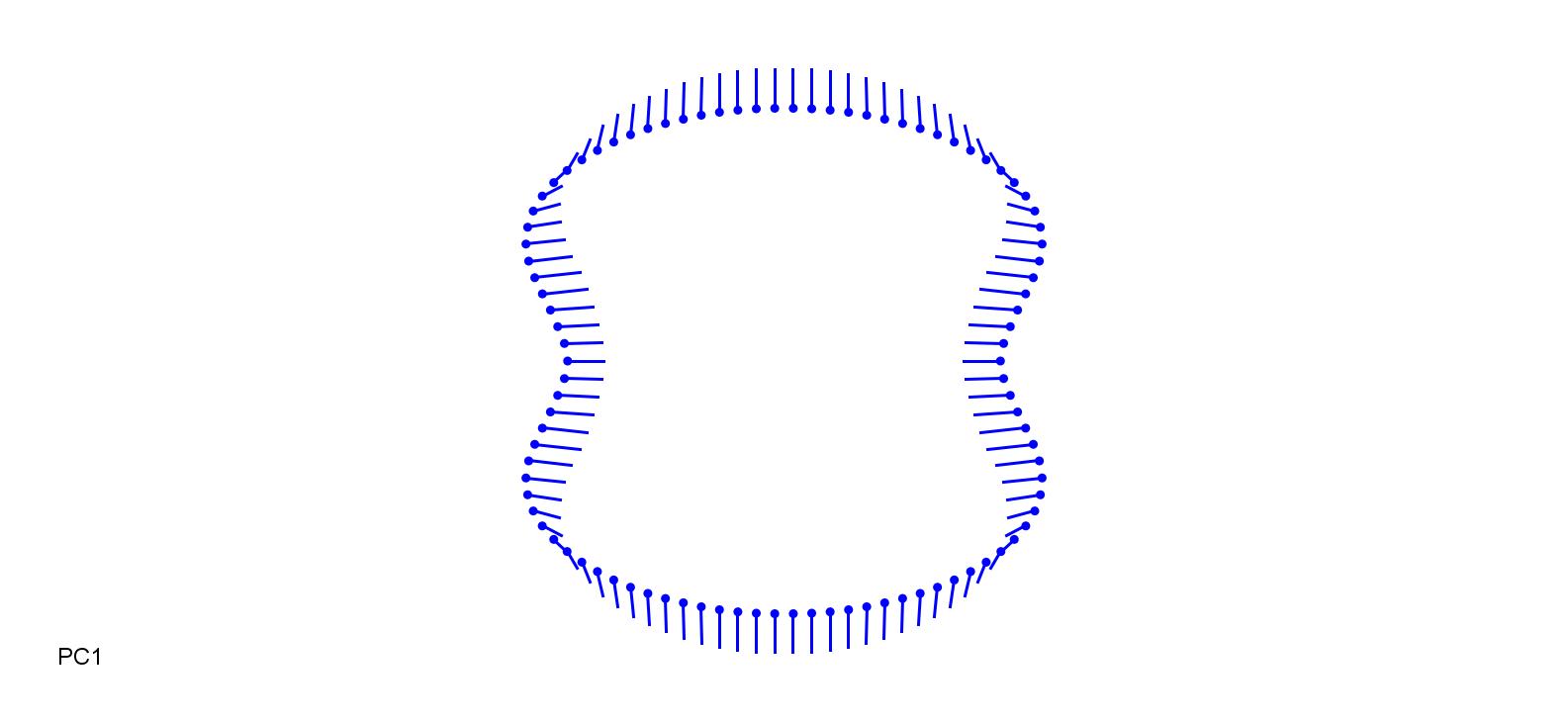

### DacGlo_PC2-0.1.jpg

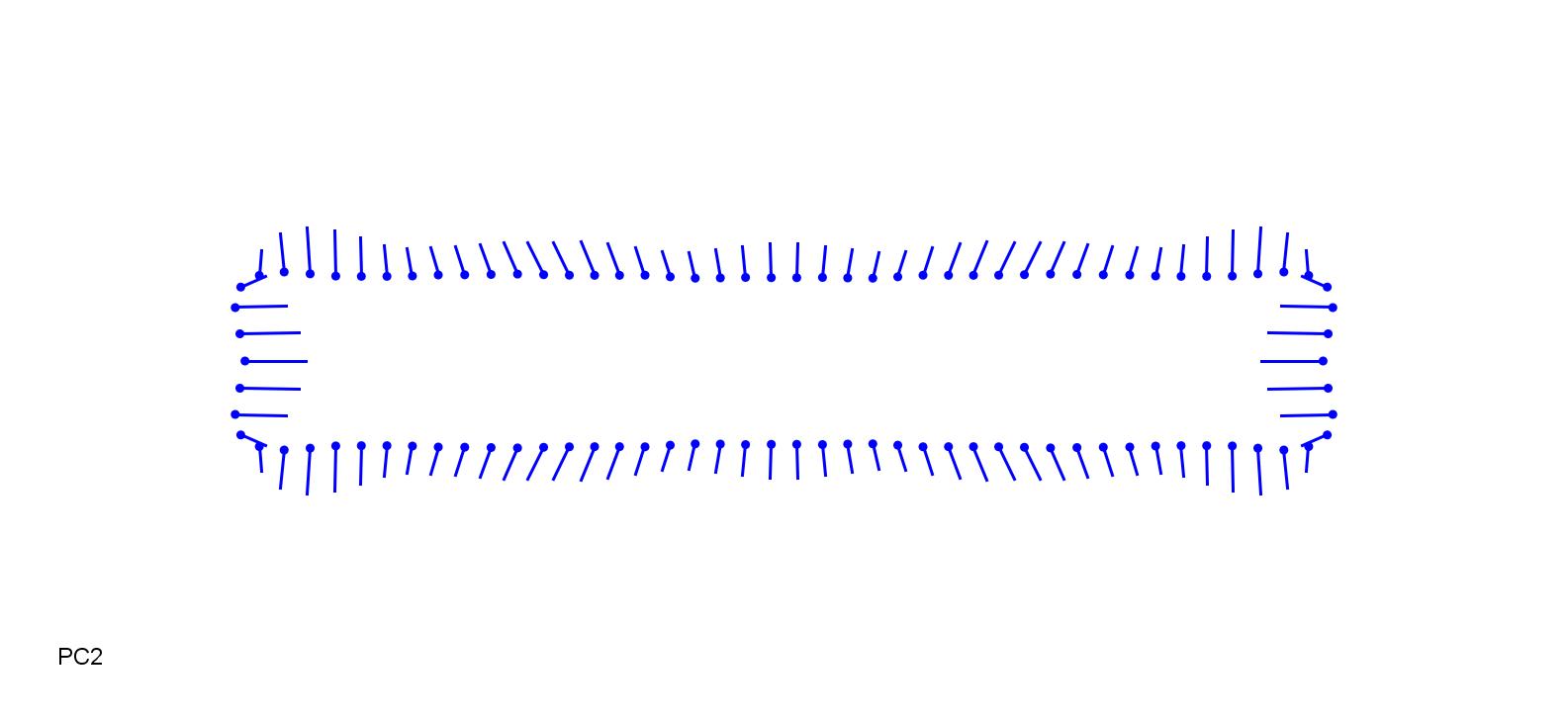

### DacGlo_PC2_0.1.jpg

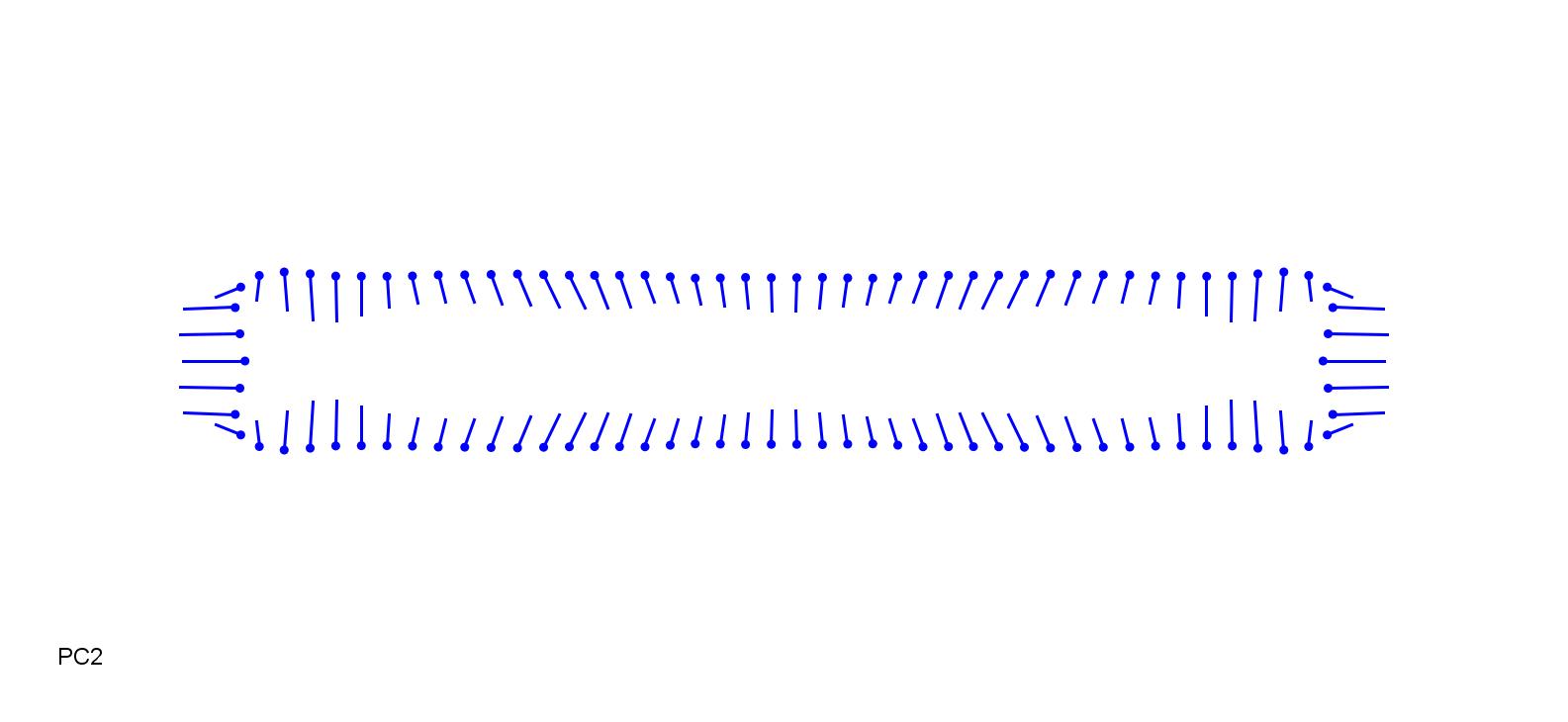

### DanAlp-0.13.jpg

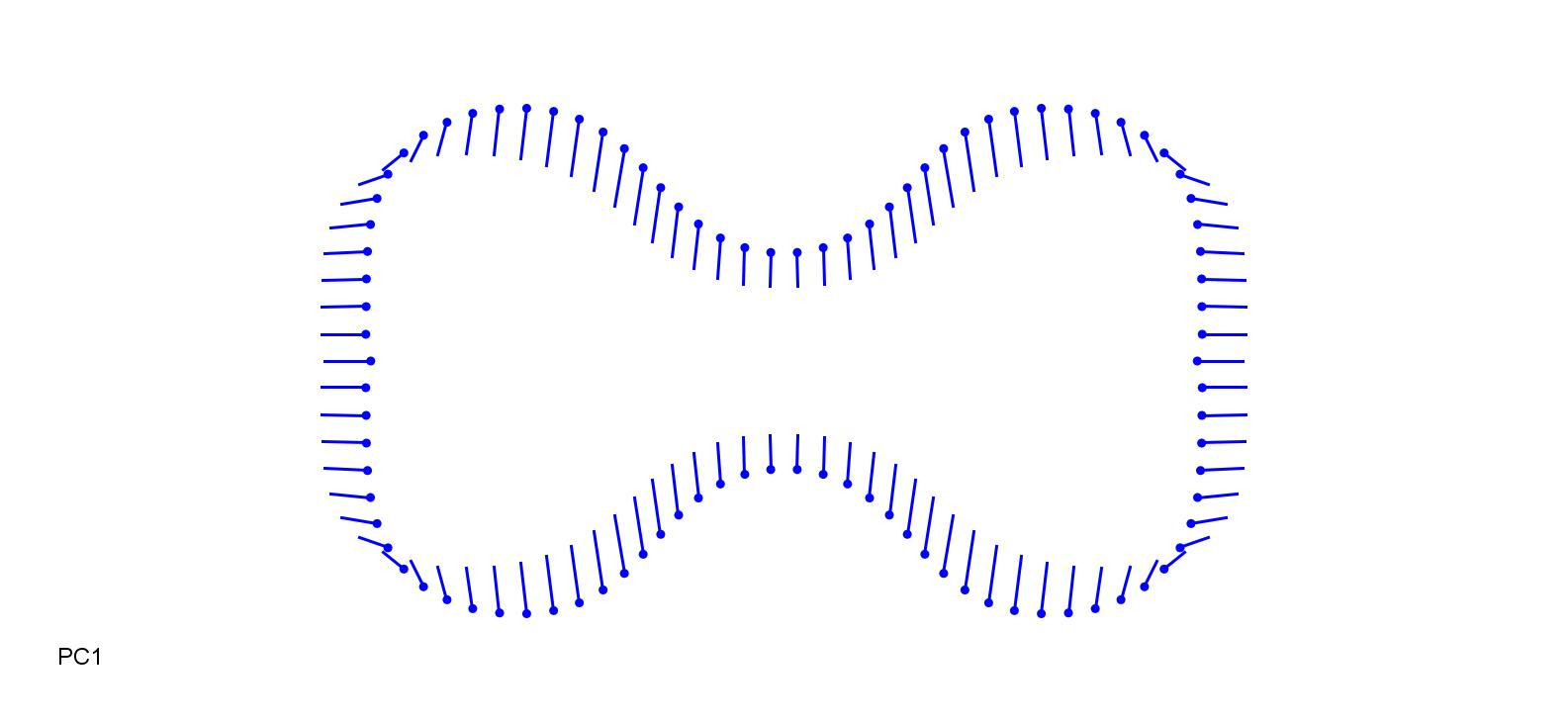

### DanAlp_0.13.jpg

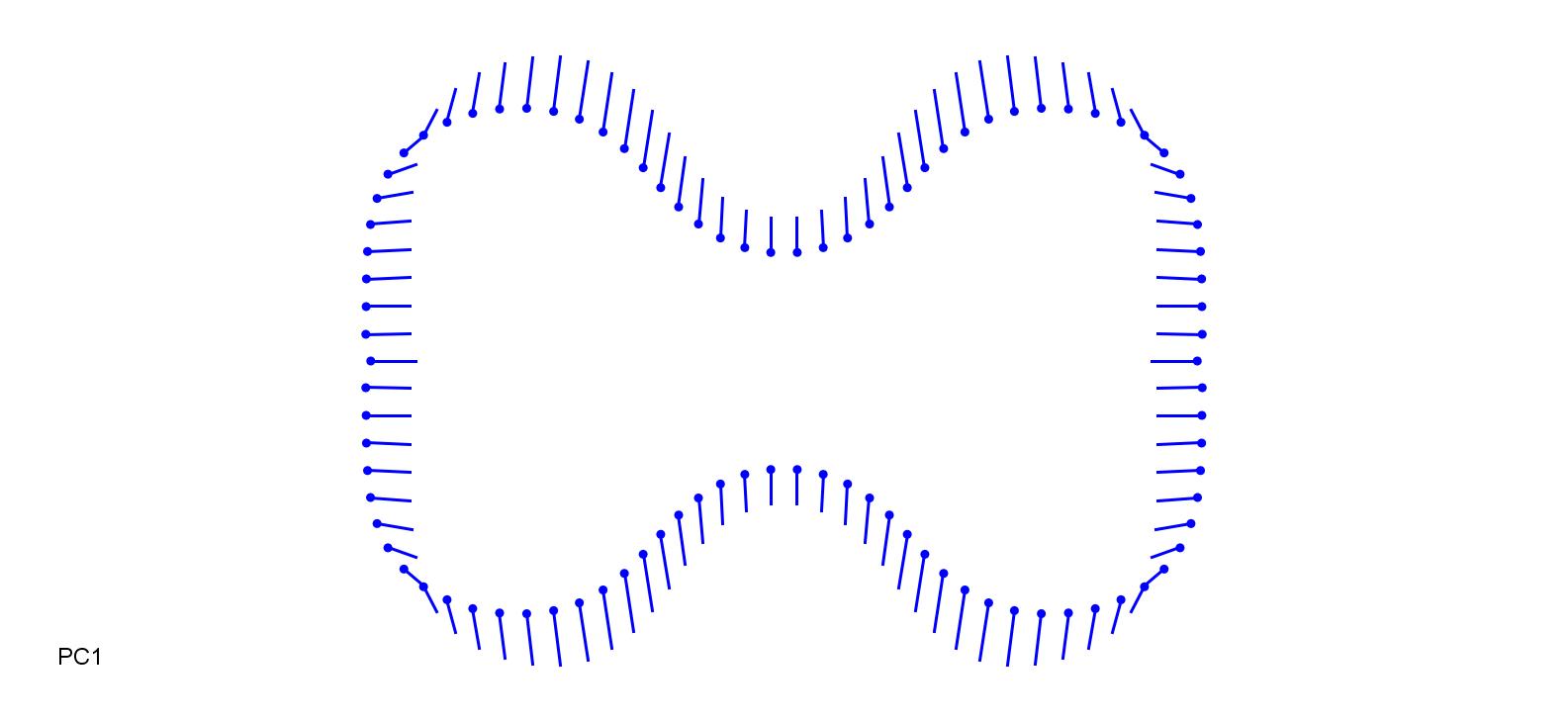

### DanDec-0.15.jpg

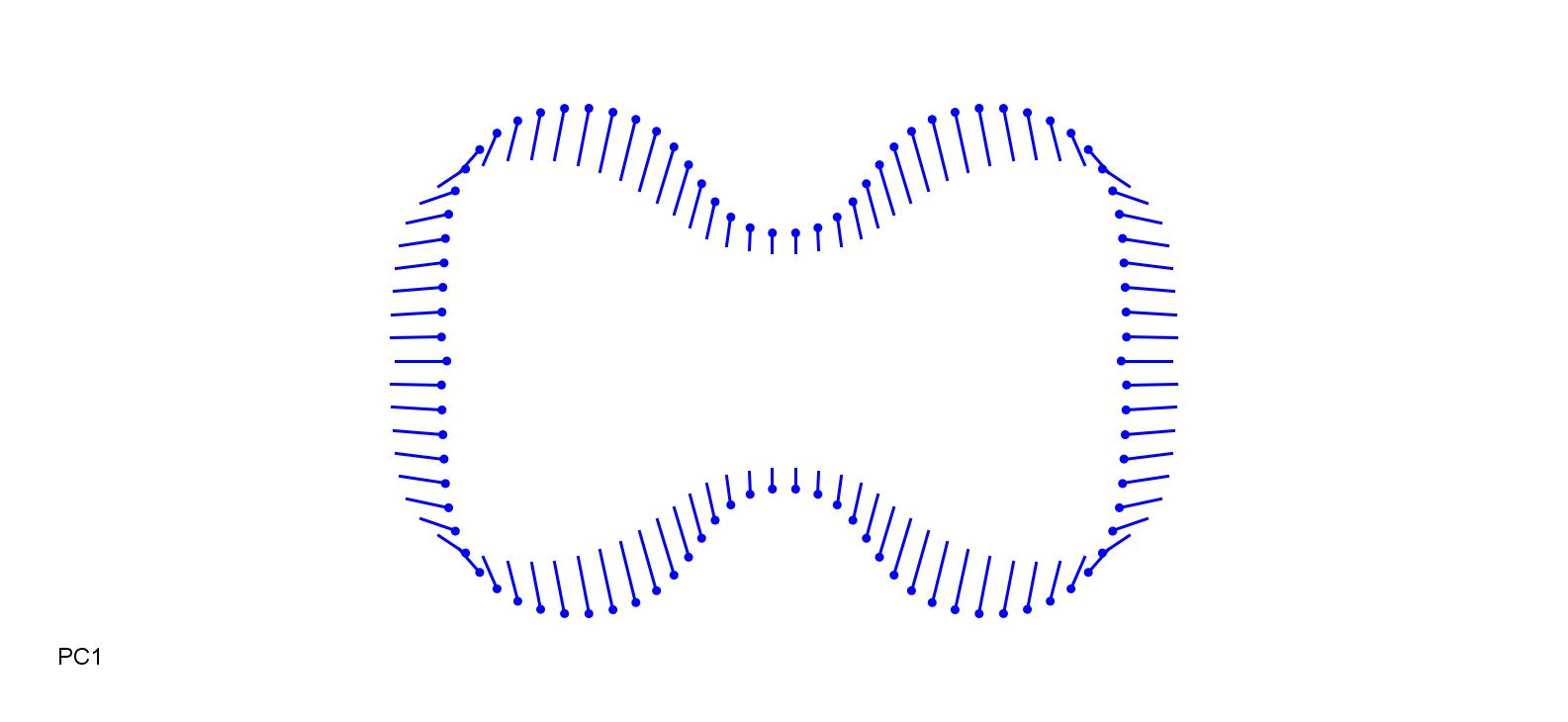

### DanDec_0.15.jpg

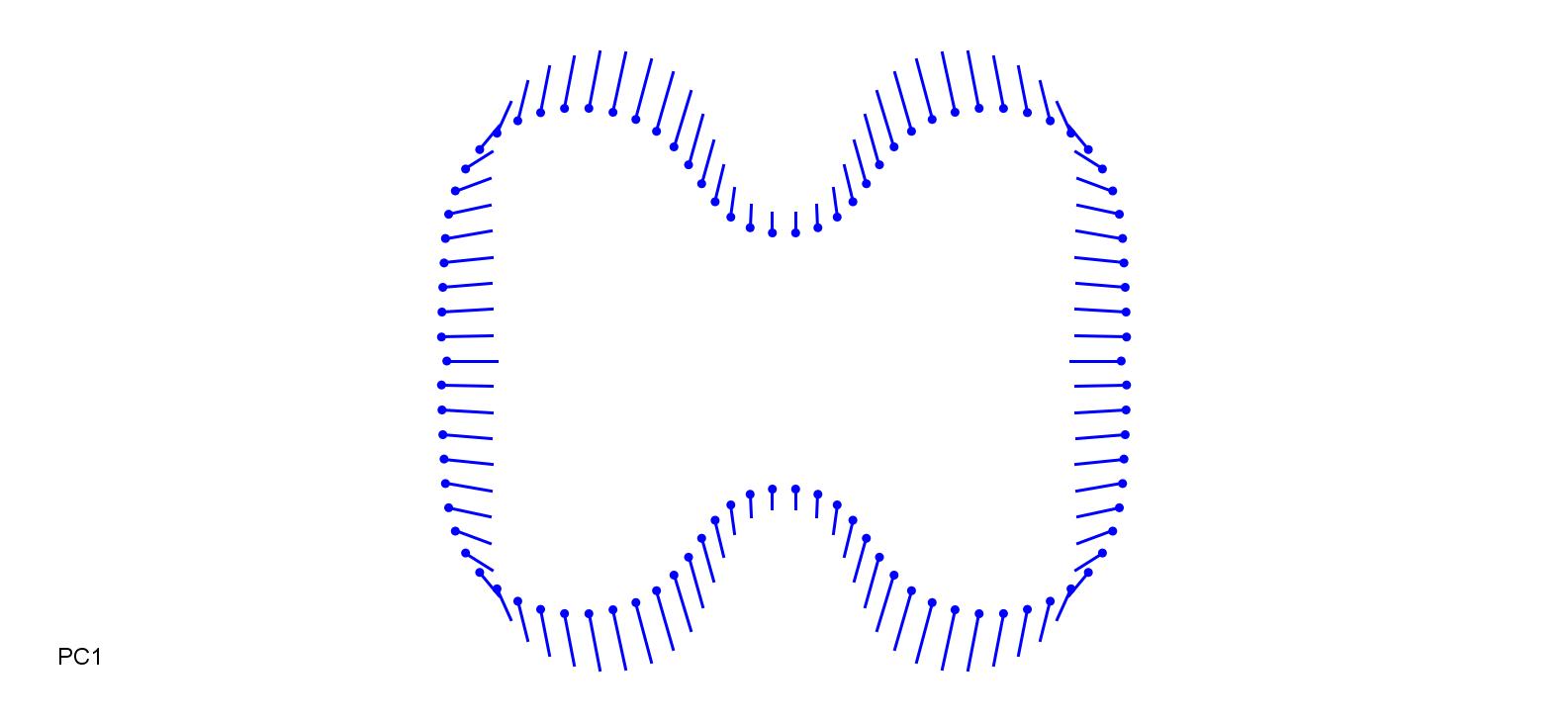

### DesBip-0.13.jpg

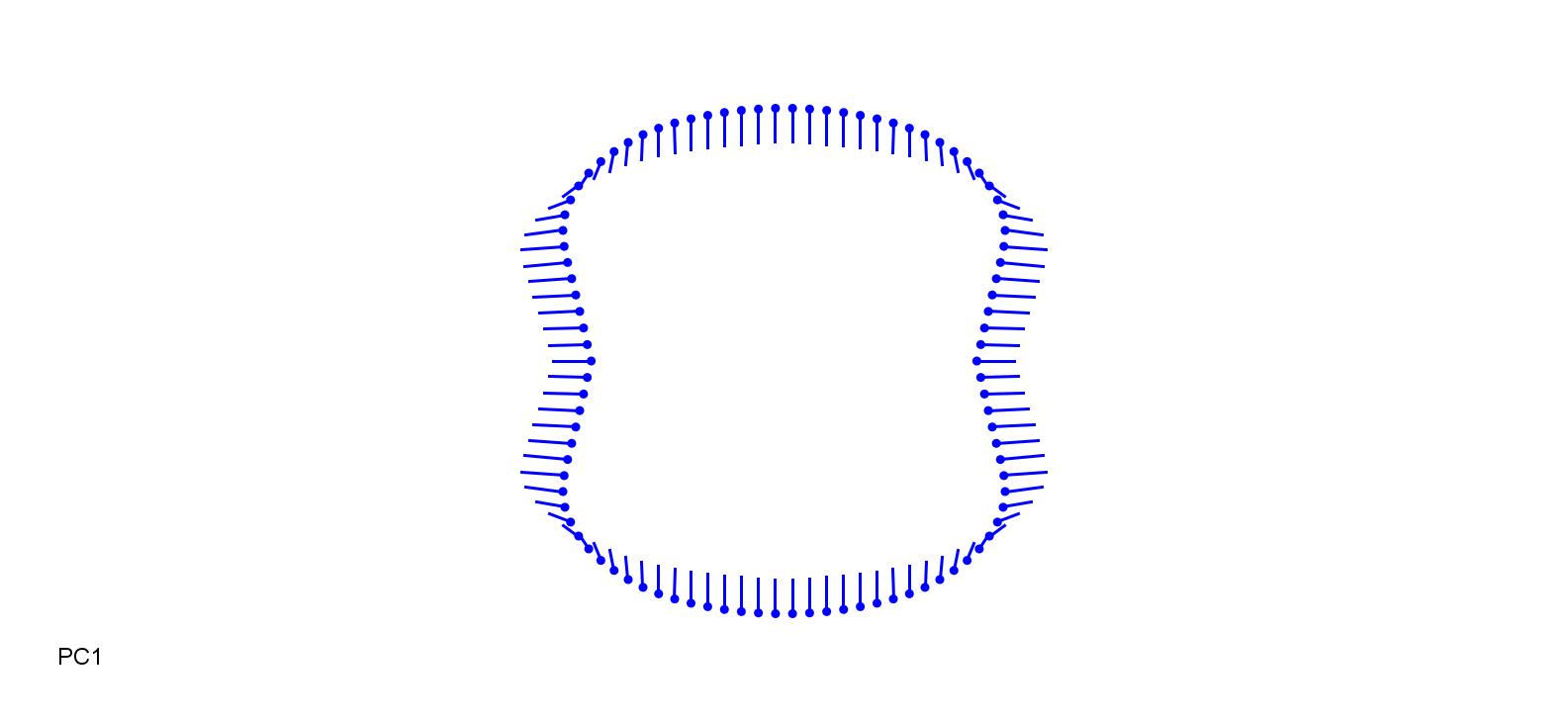

### DesBip_0.13.jpg

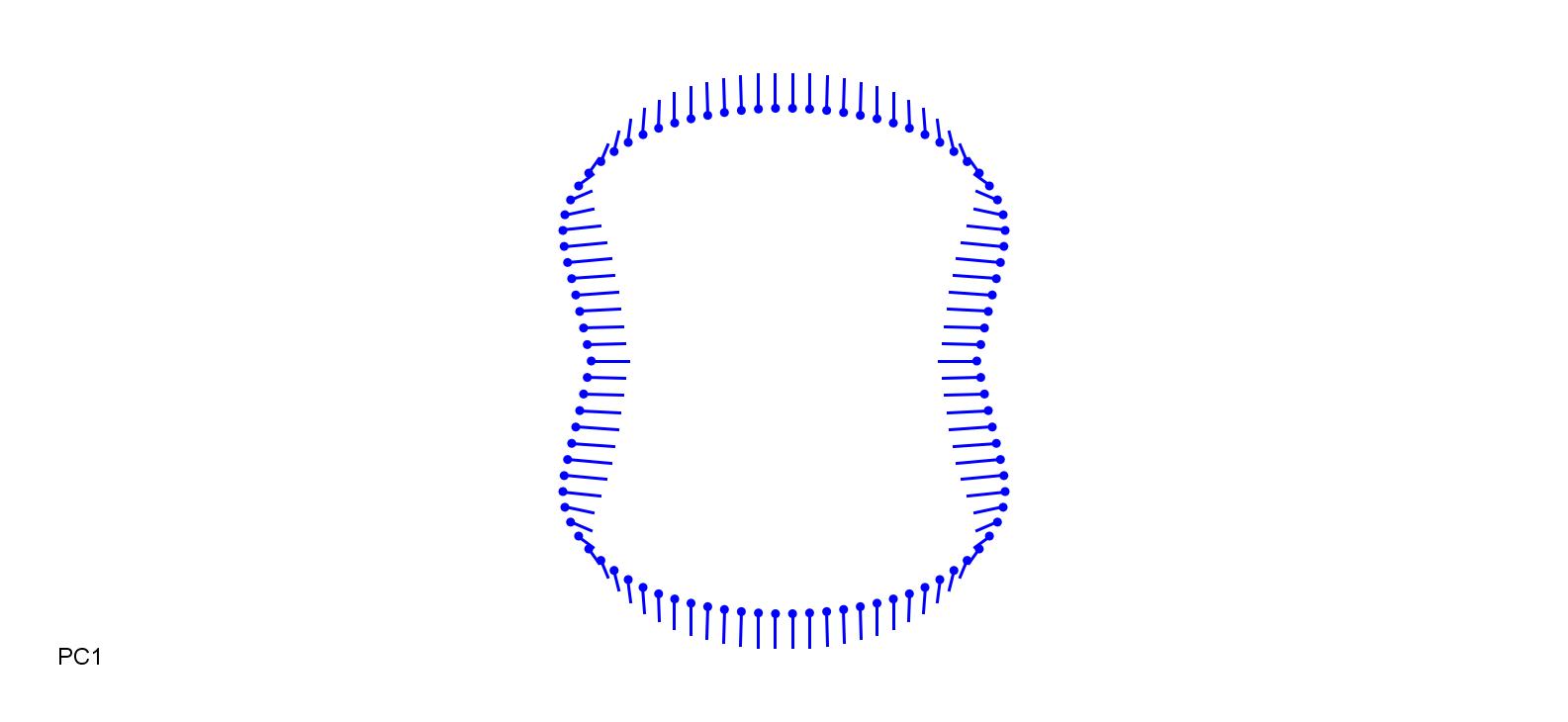

### DichAnn-0.15.jpg

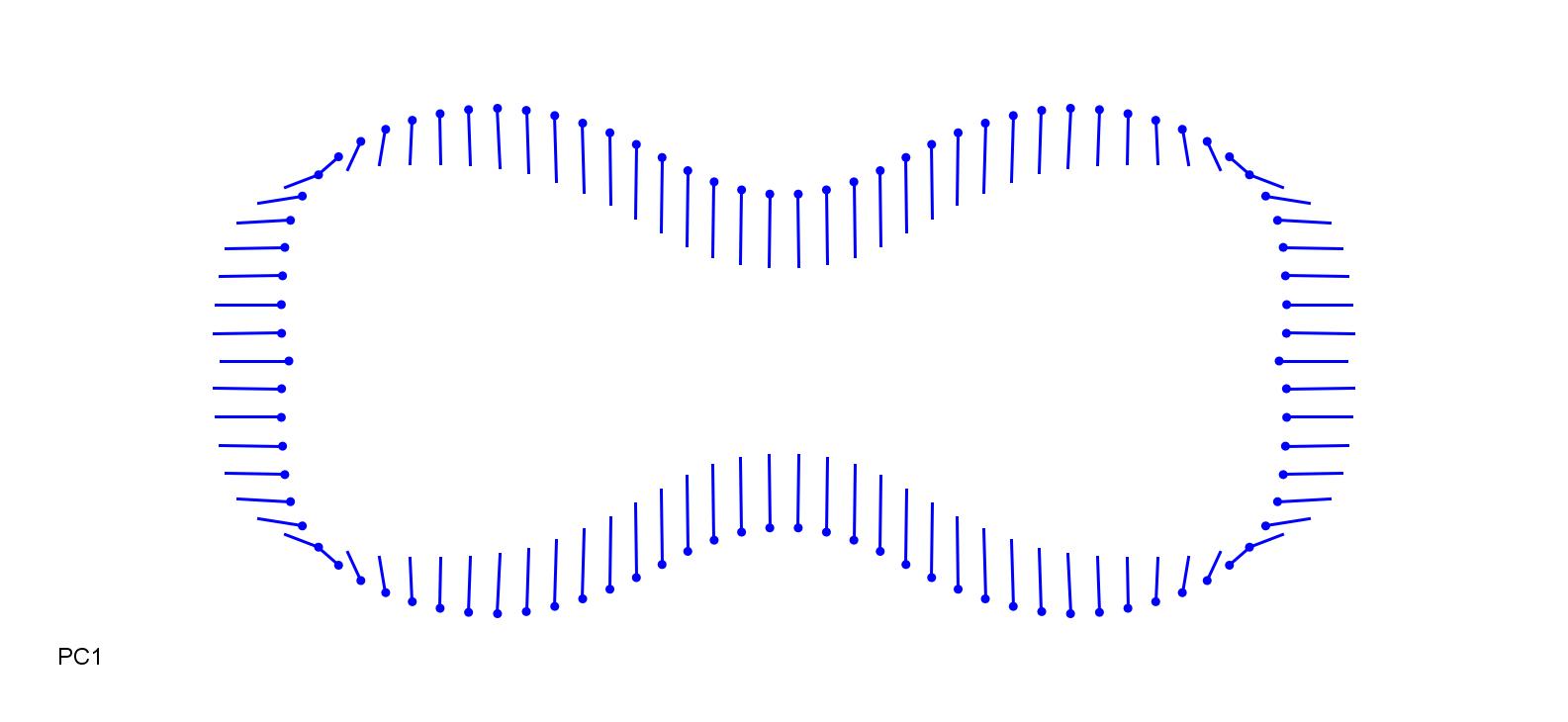

### DichAnn_0.25.jpg

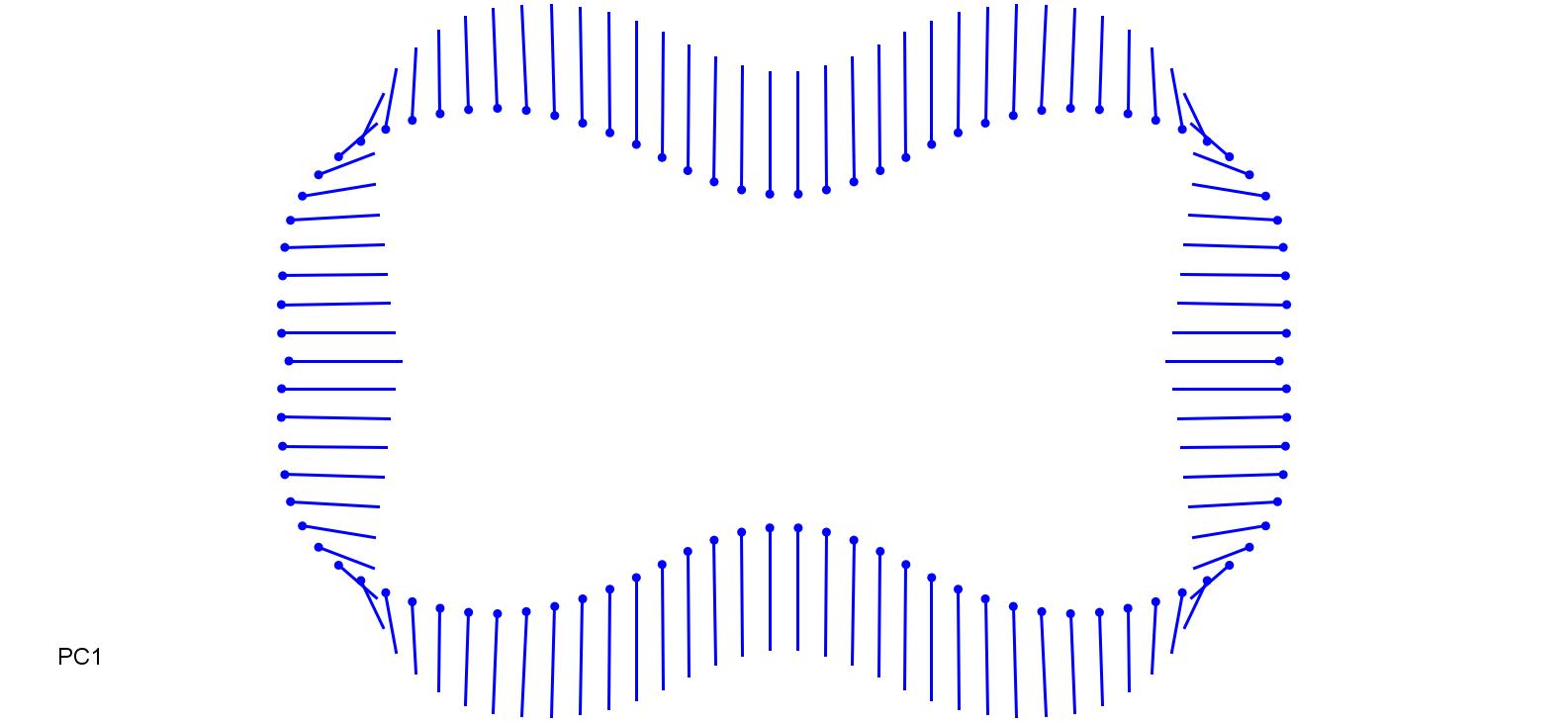

### DigSan-0.2.jpg

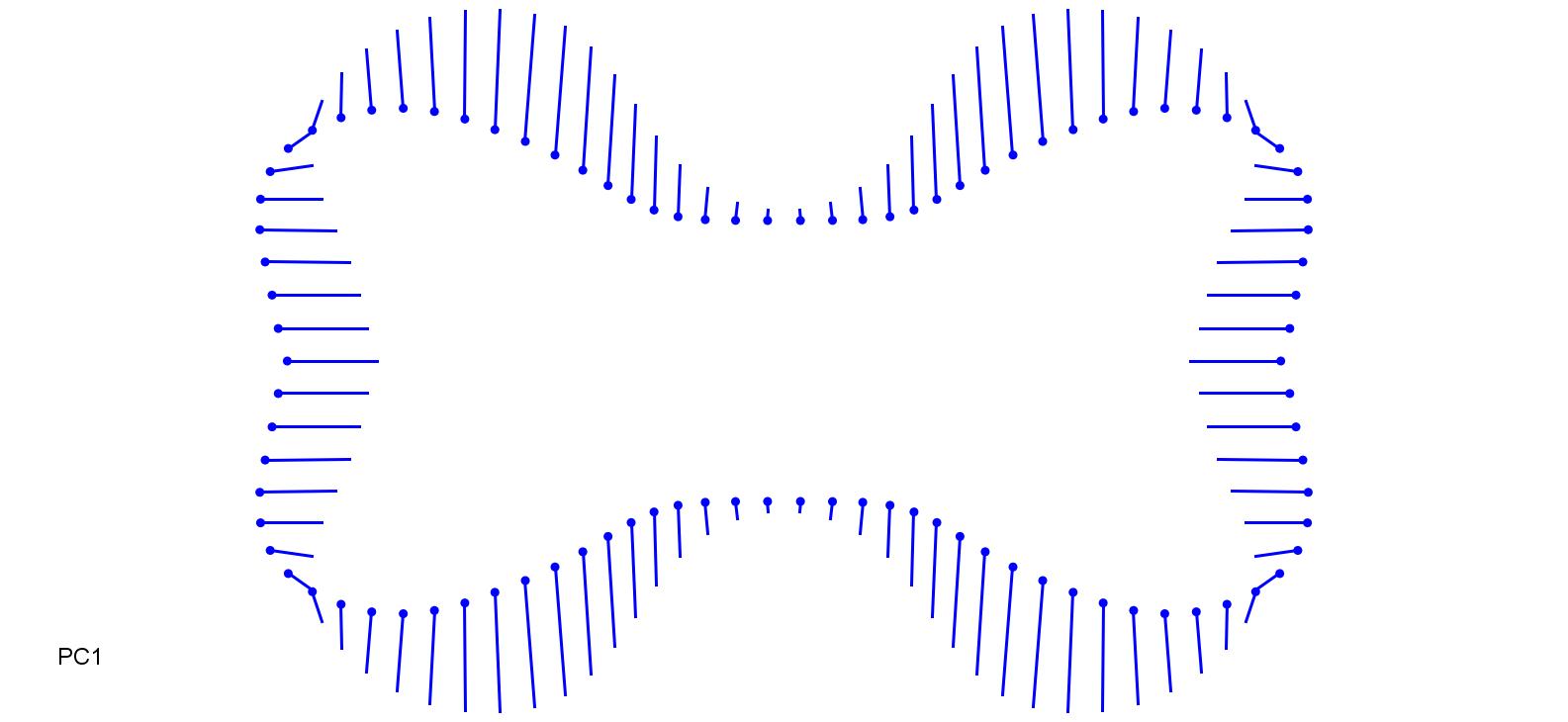

### DigSan_0.25.jpg

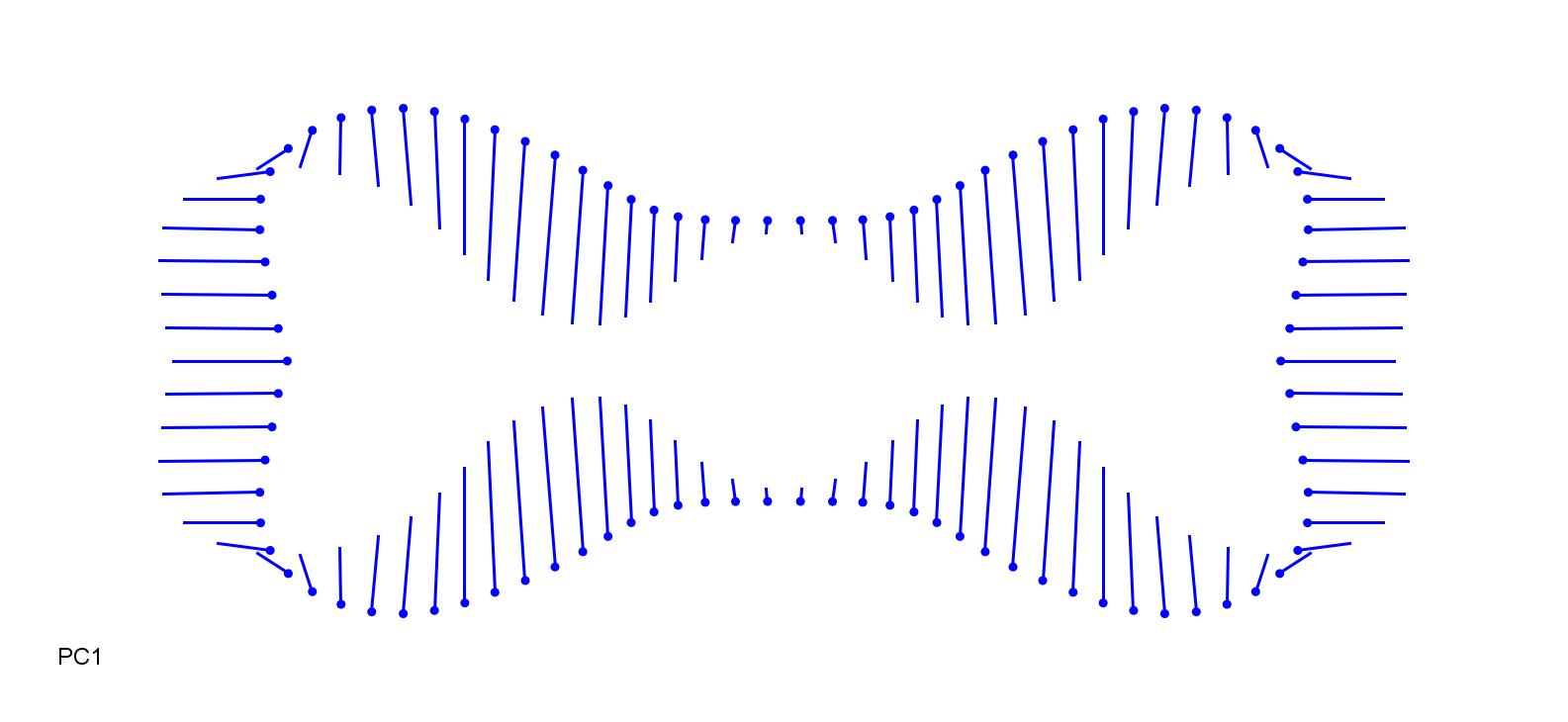

### EchiCrG-0.2.jpg

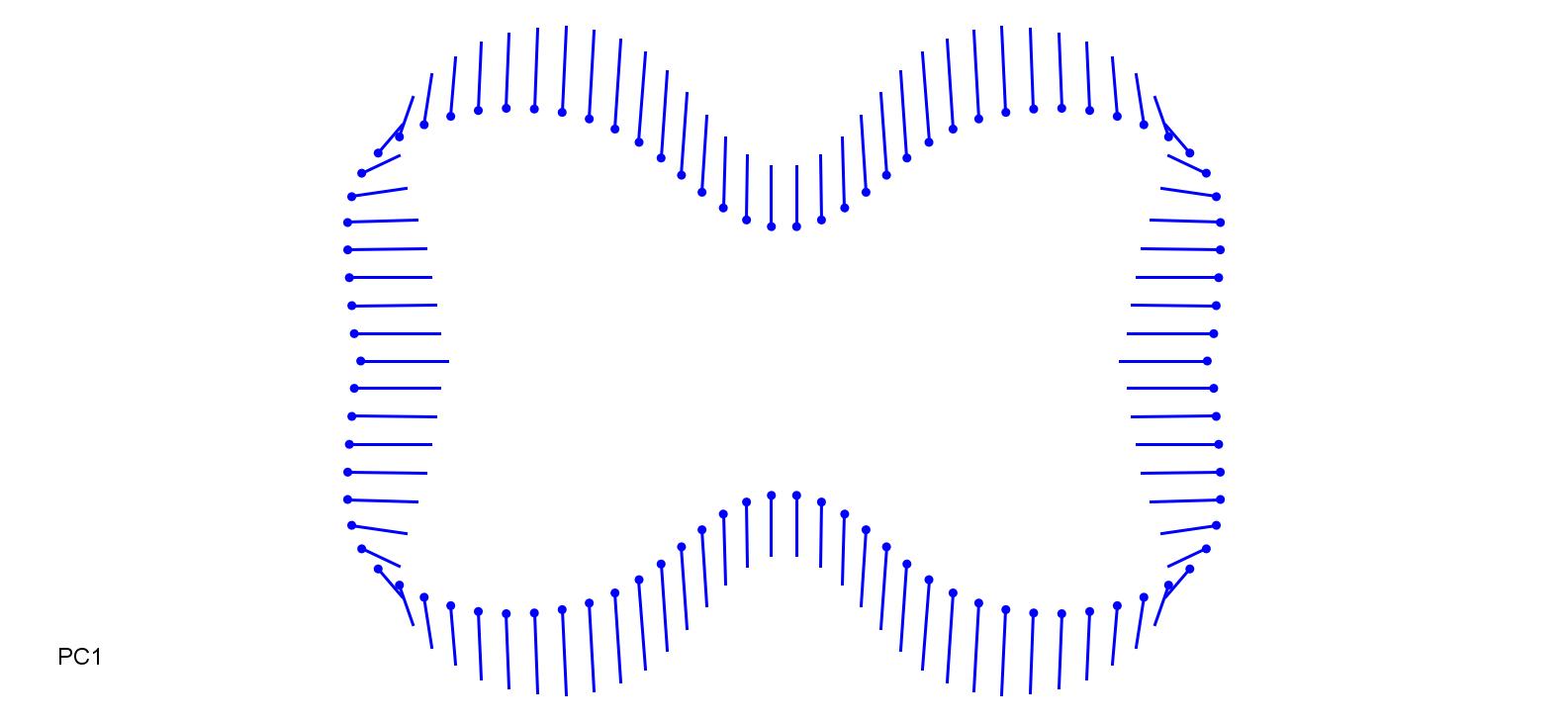

### EchiCrG_0.2.jpg

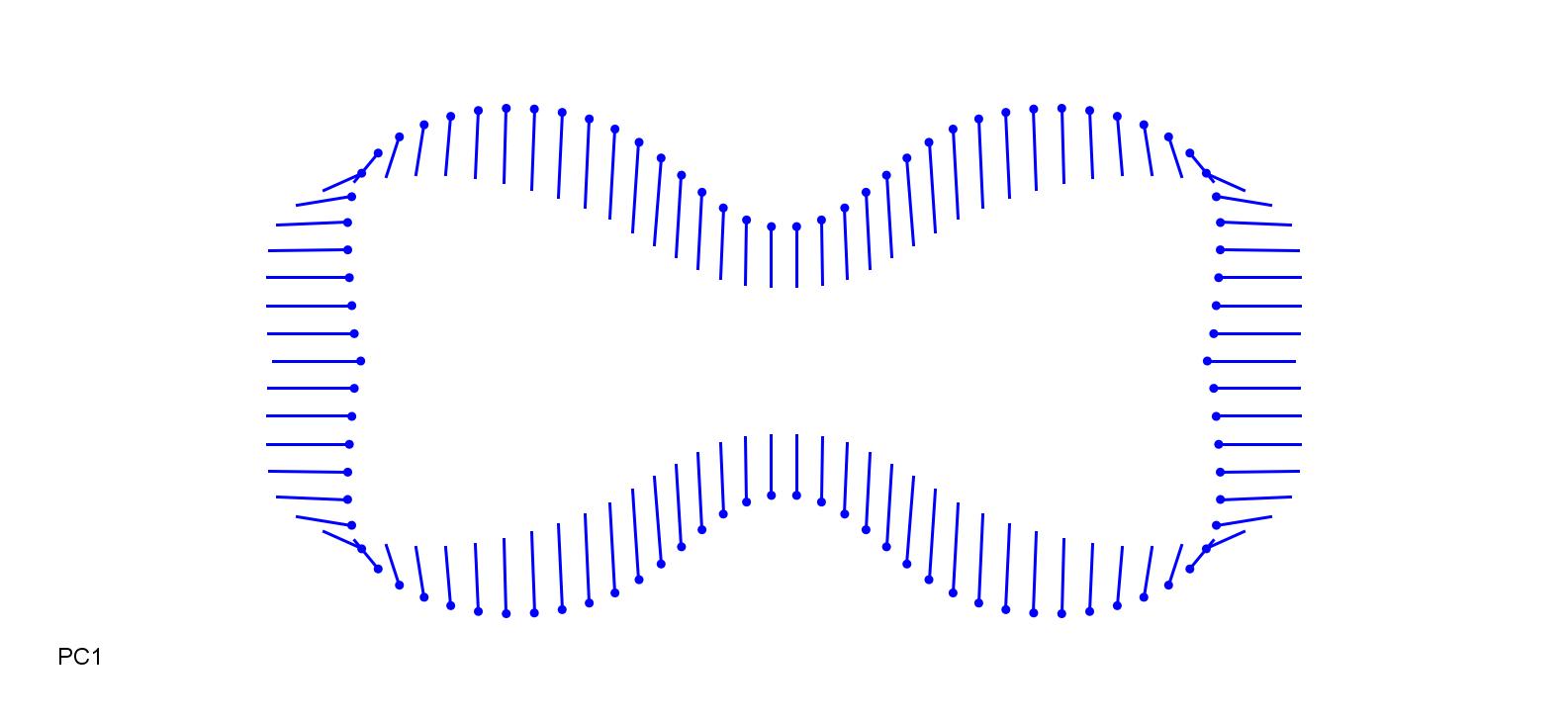

### EhrEre-0.15.jpg

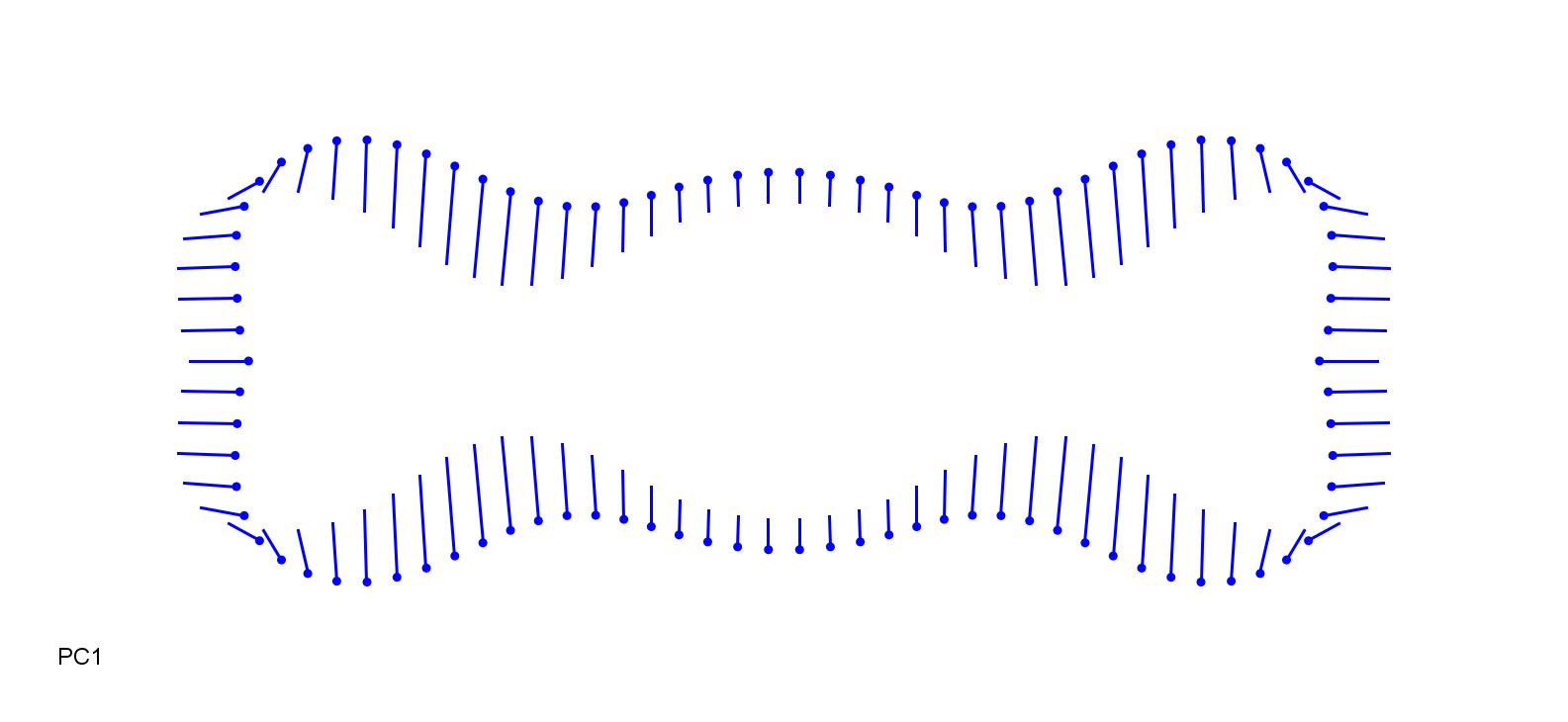

### EhrEre_0.23.jpg

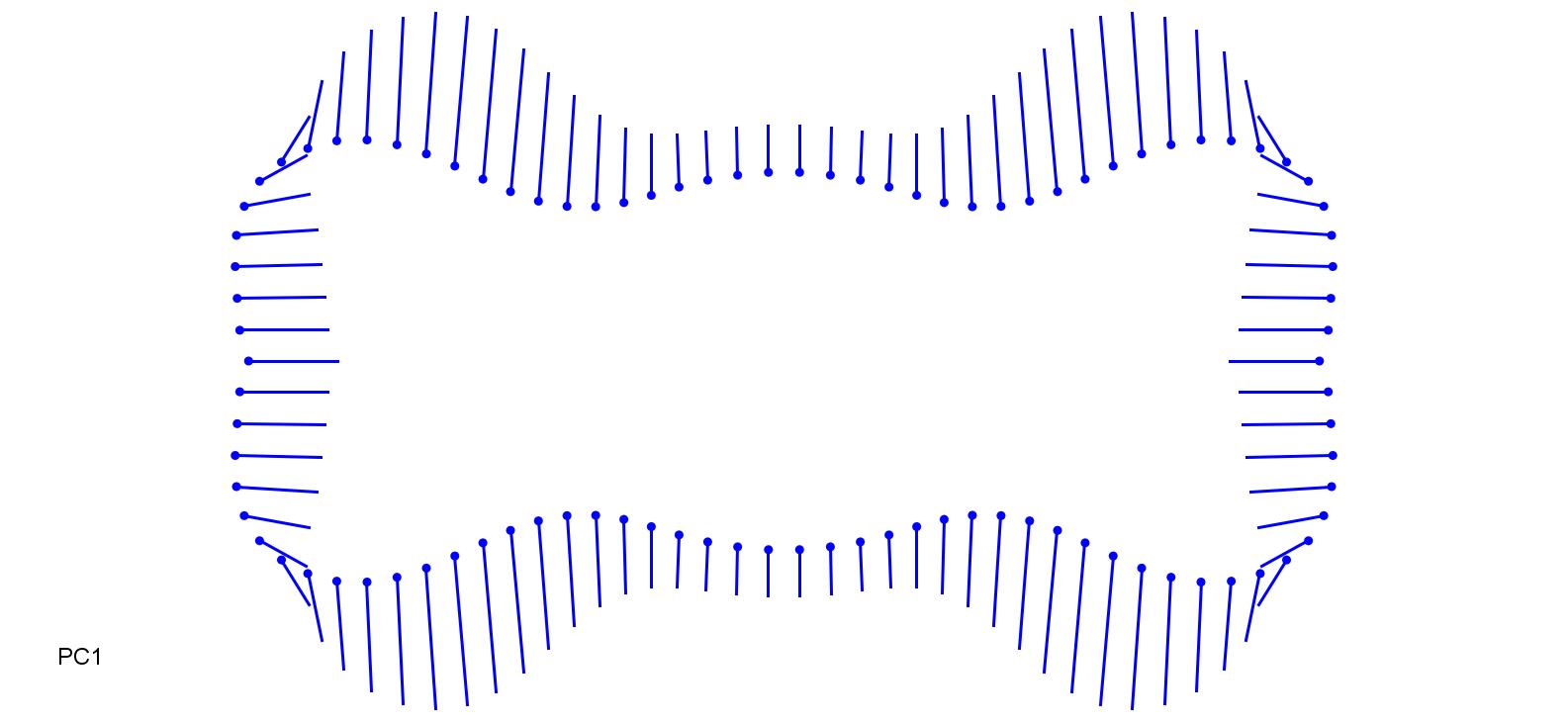

### EnnSco-0.15.jpg

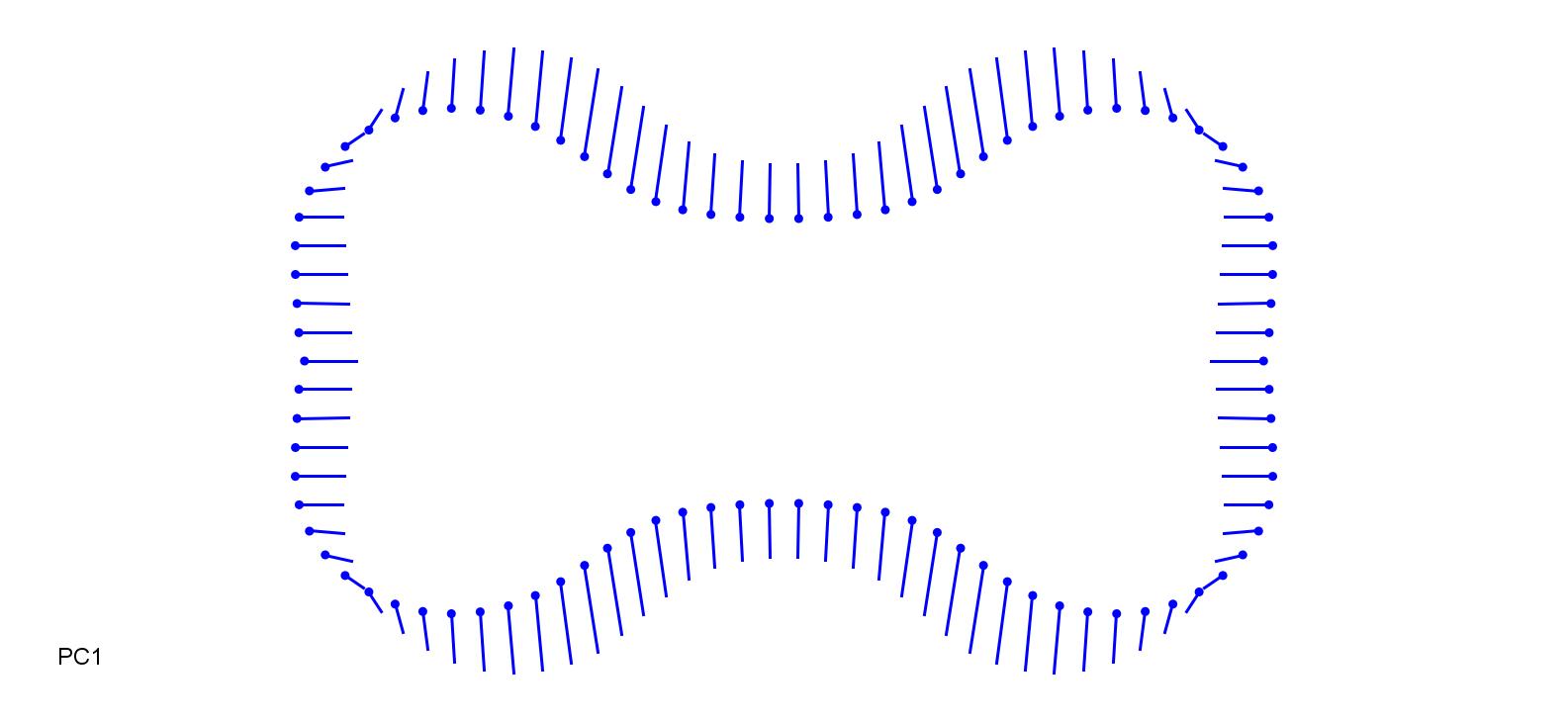

### EnnSco_0.15.jpg

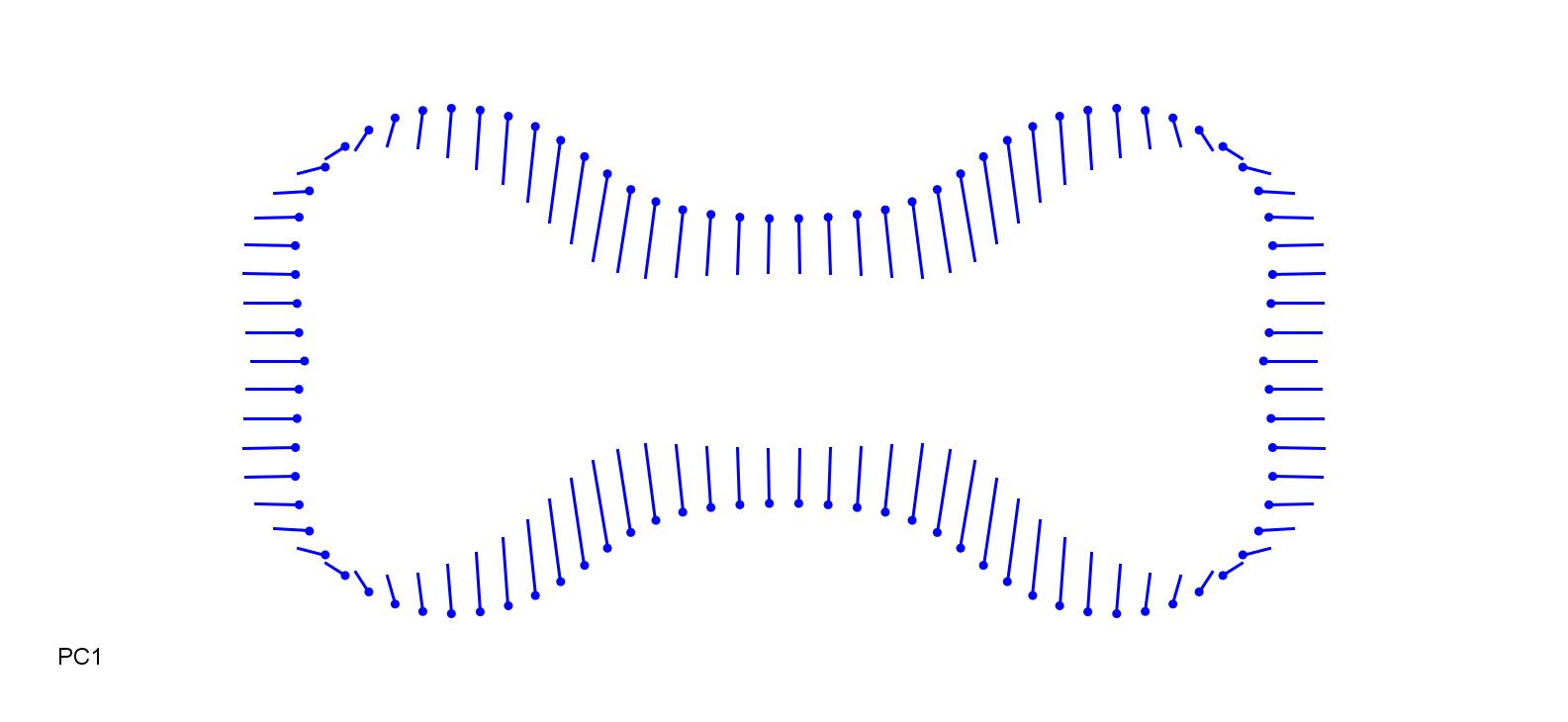

### EraMin-0.3.jpg

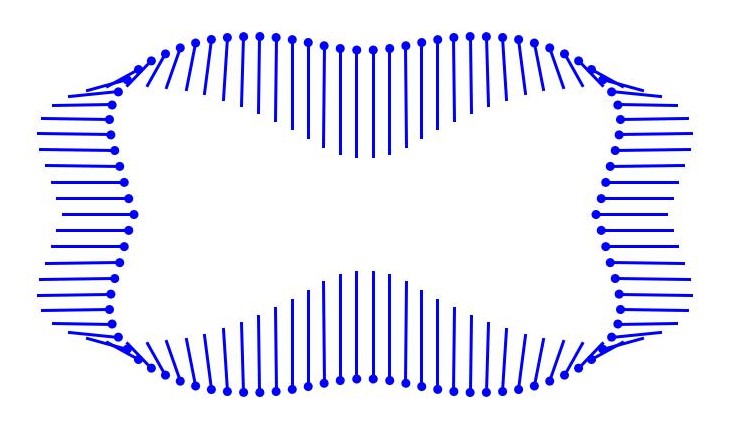

### EraMin_0.3.jpg

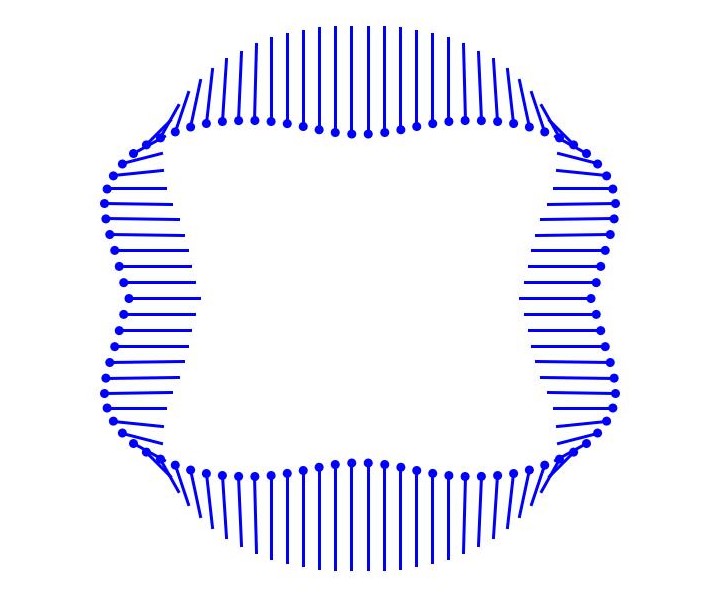

### FesAru-0.18.jpg

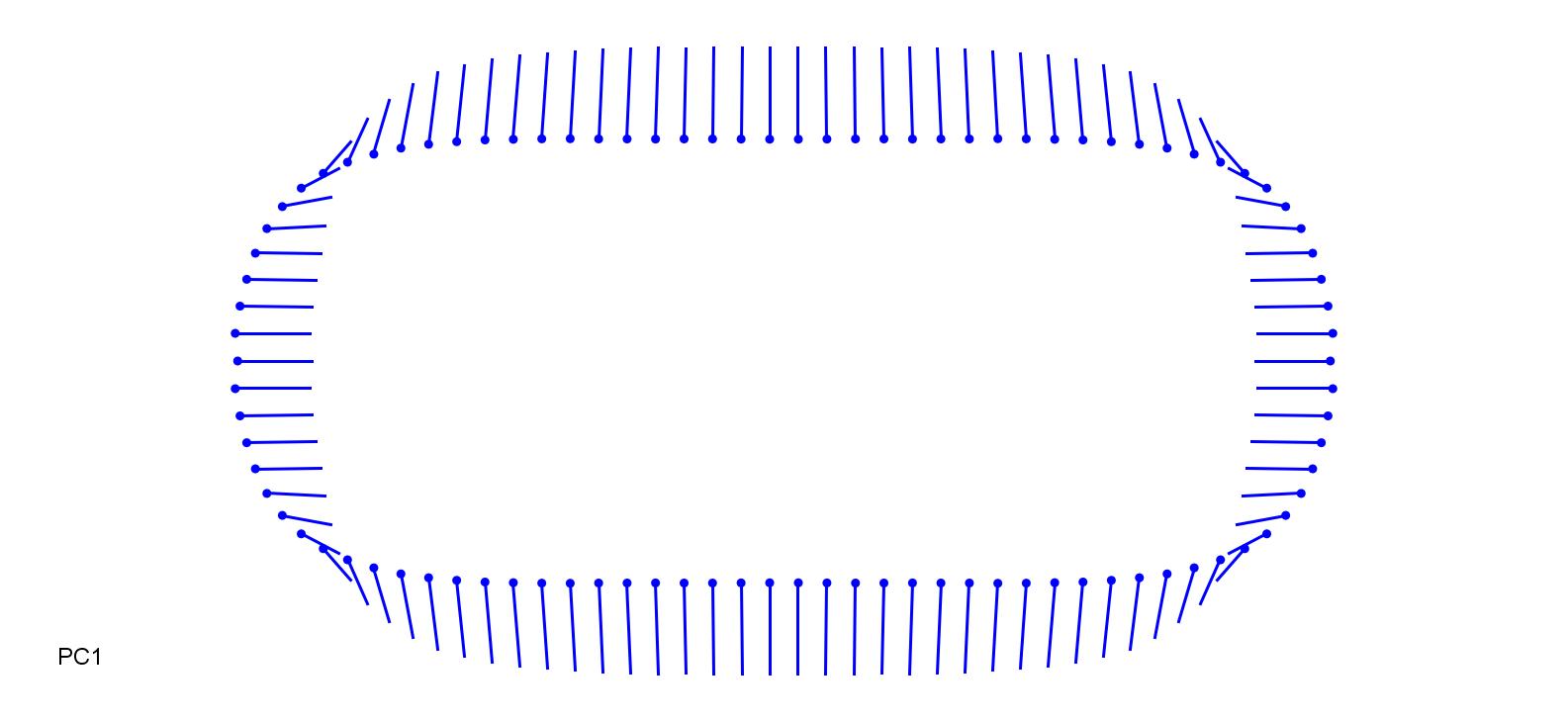

### FesAru_0.18.jpg

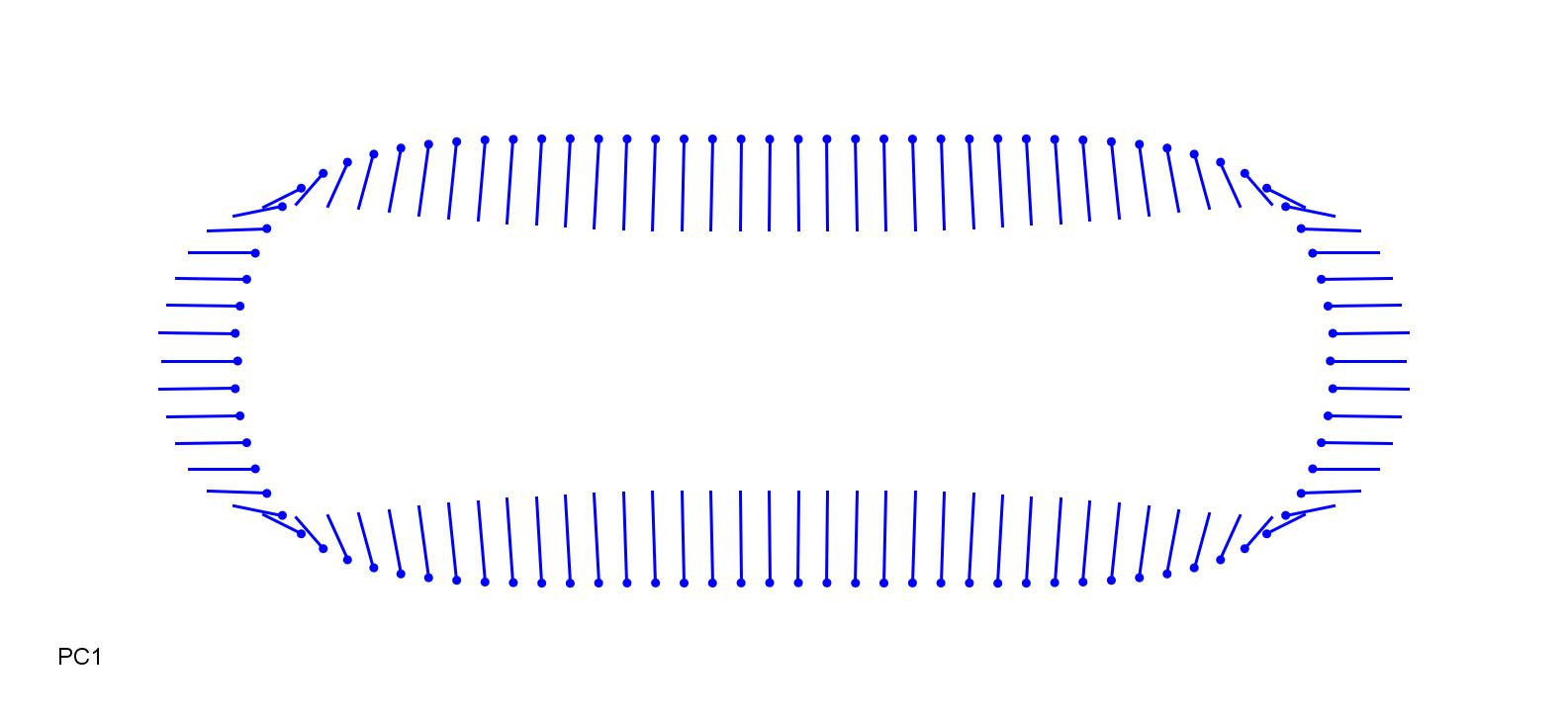

### FesGig-0.25.jpg

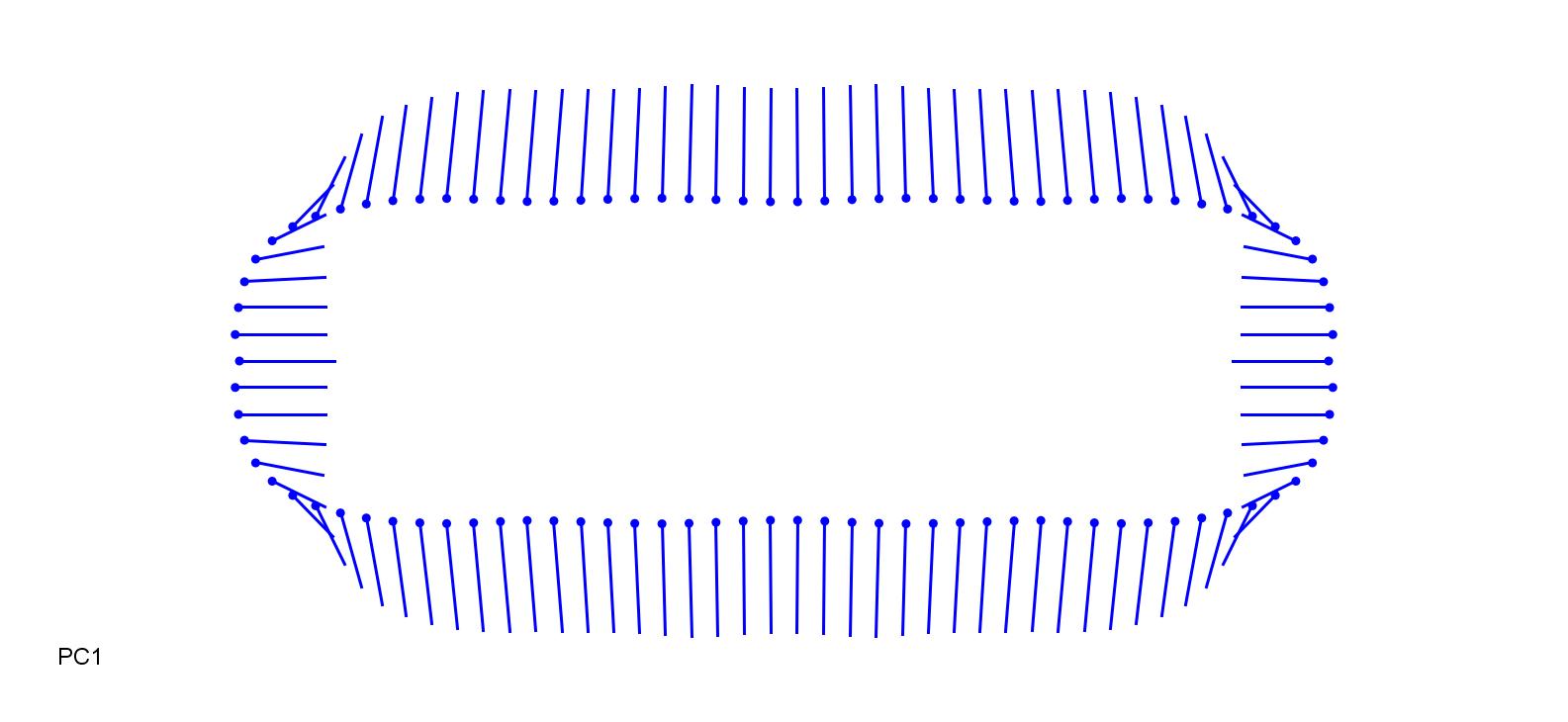

### FesGig_0.25.jpg

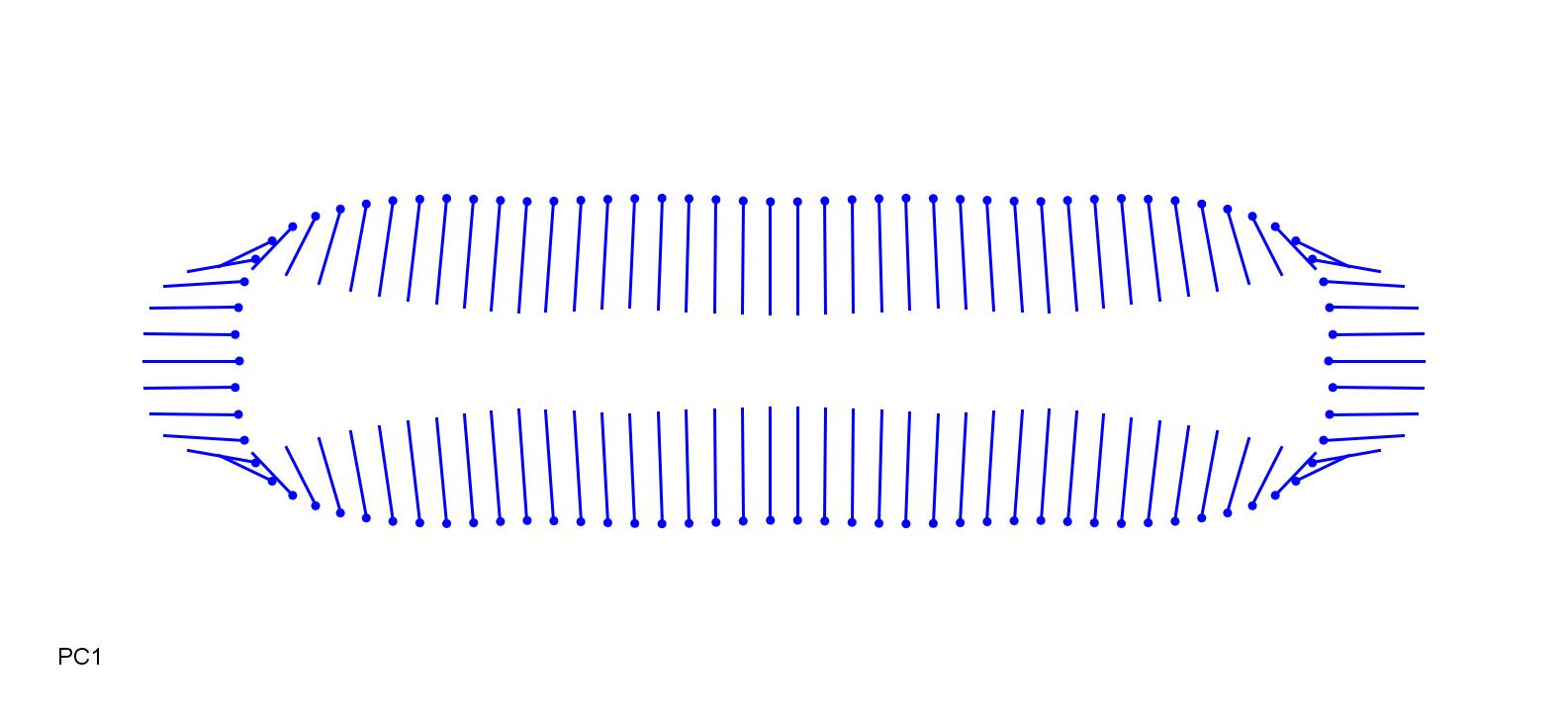

### HacMac-0.2.jpg

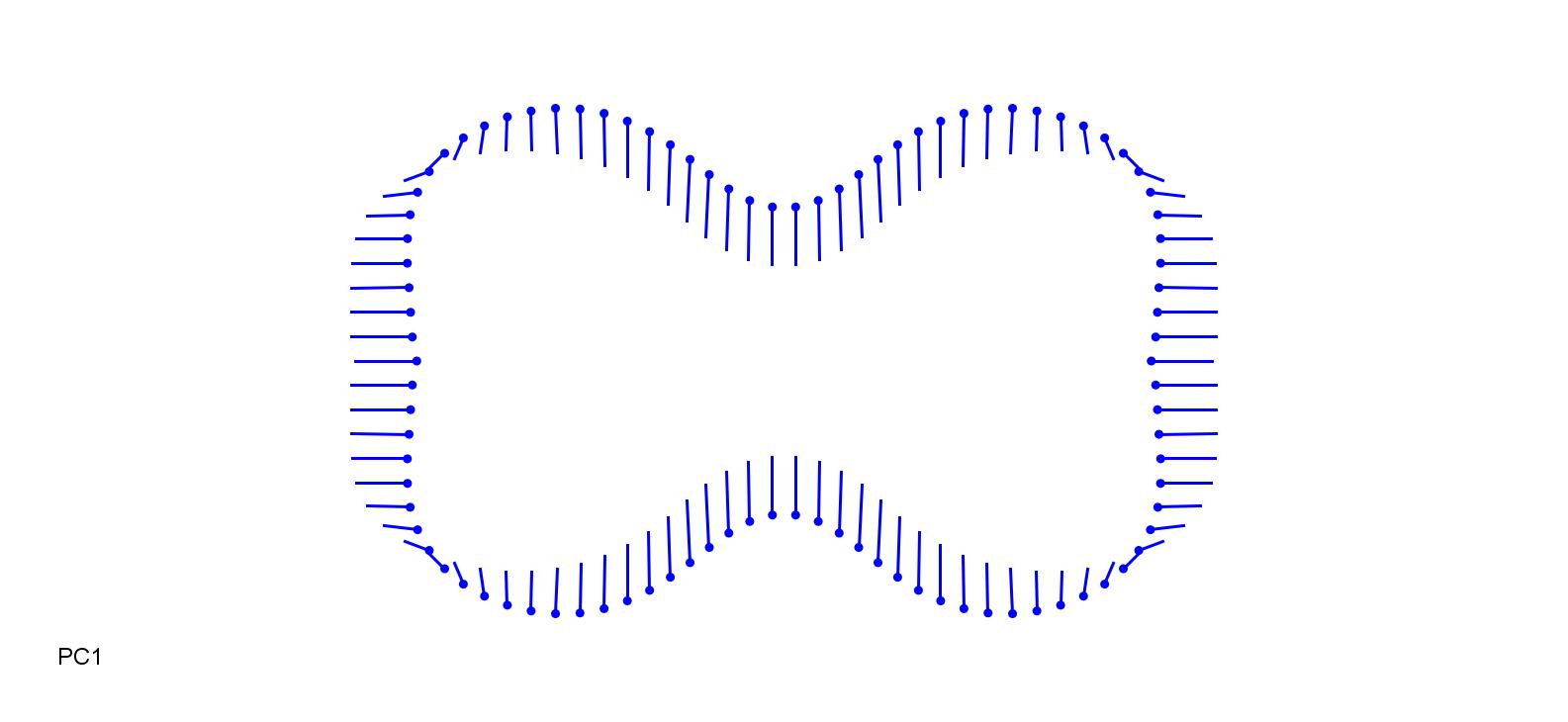

### HacMac_0.2.jpg

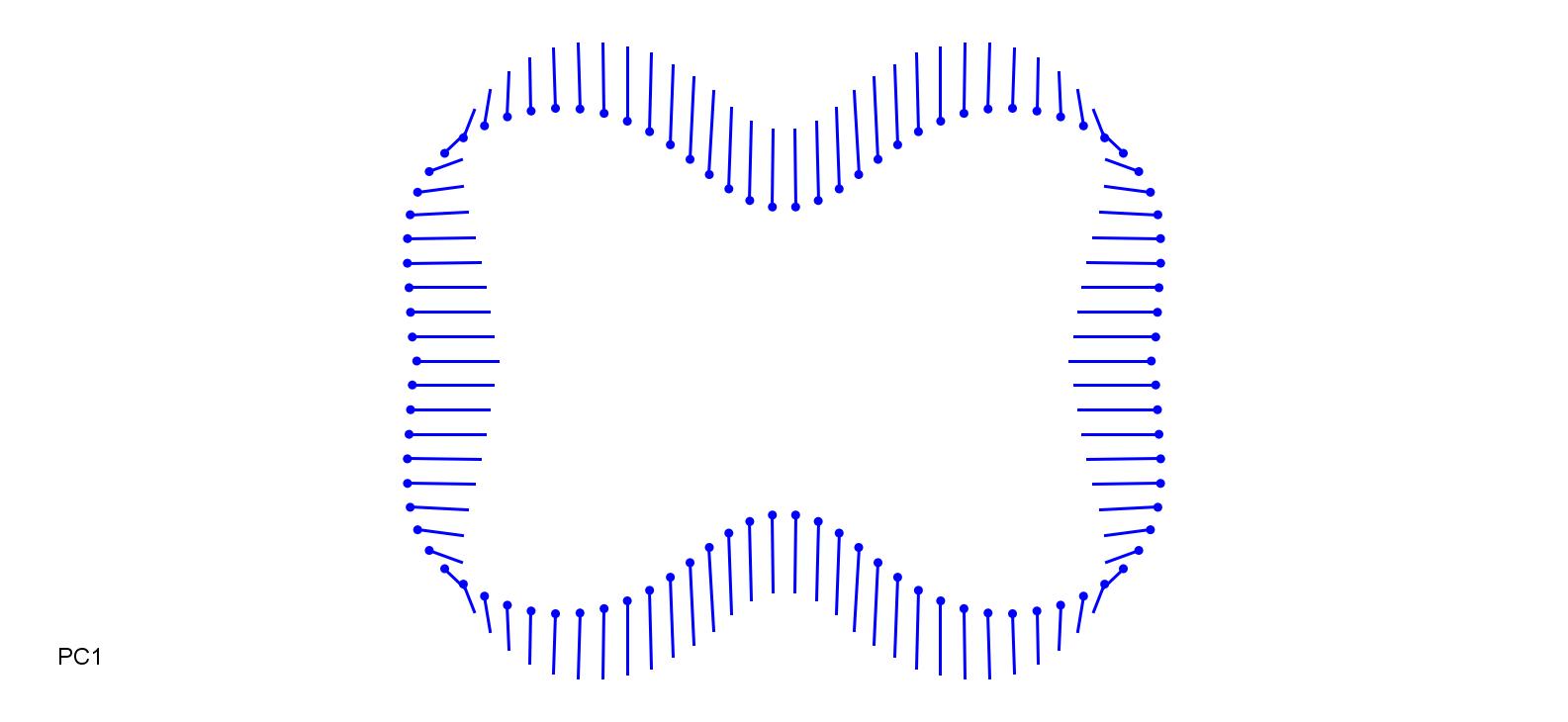

### HelPrat-0.15.jpg

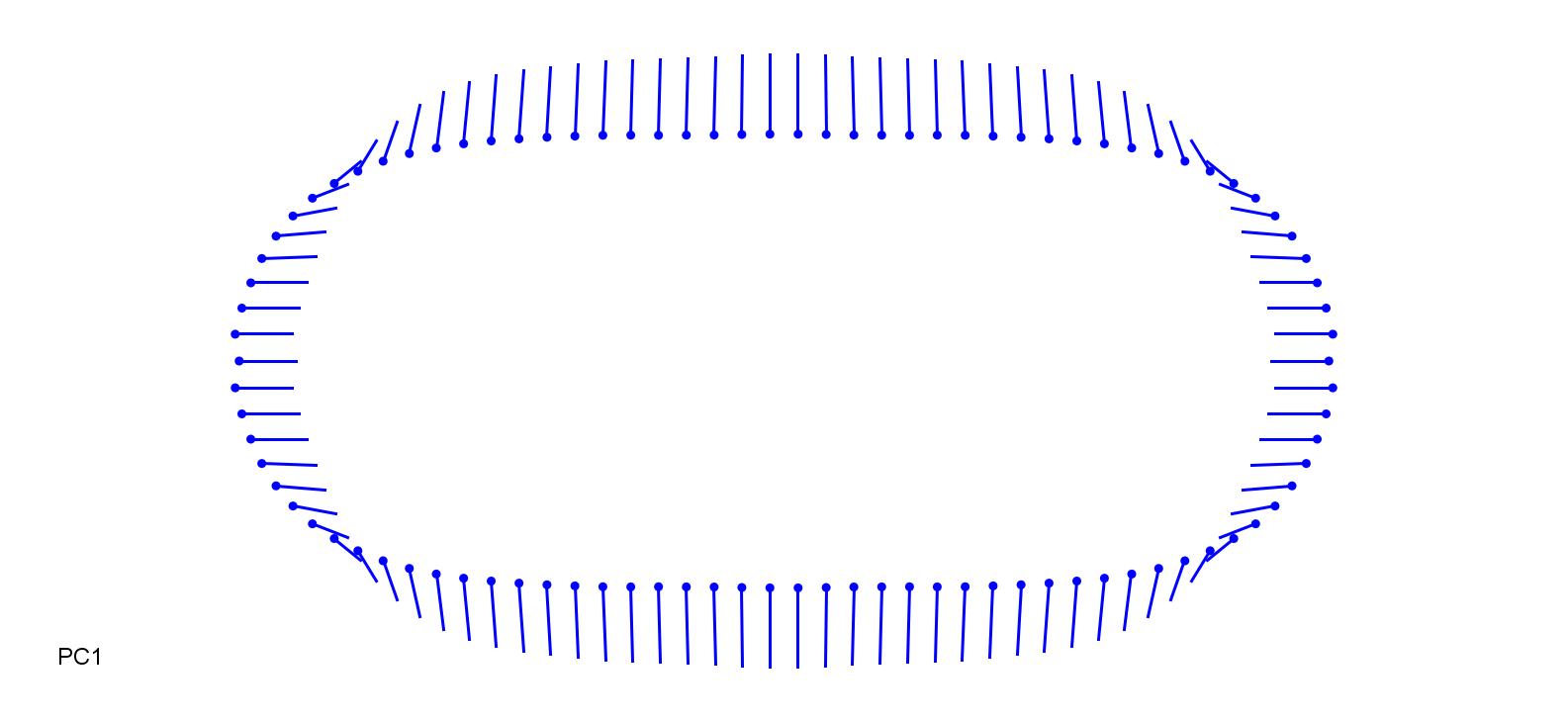

### HelPrat_0.15.jpg

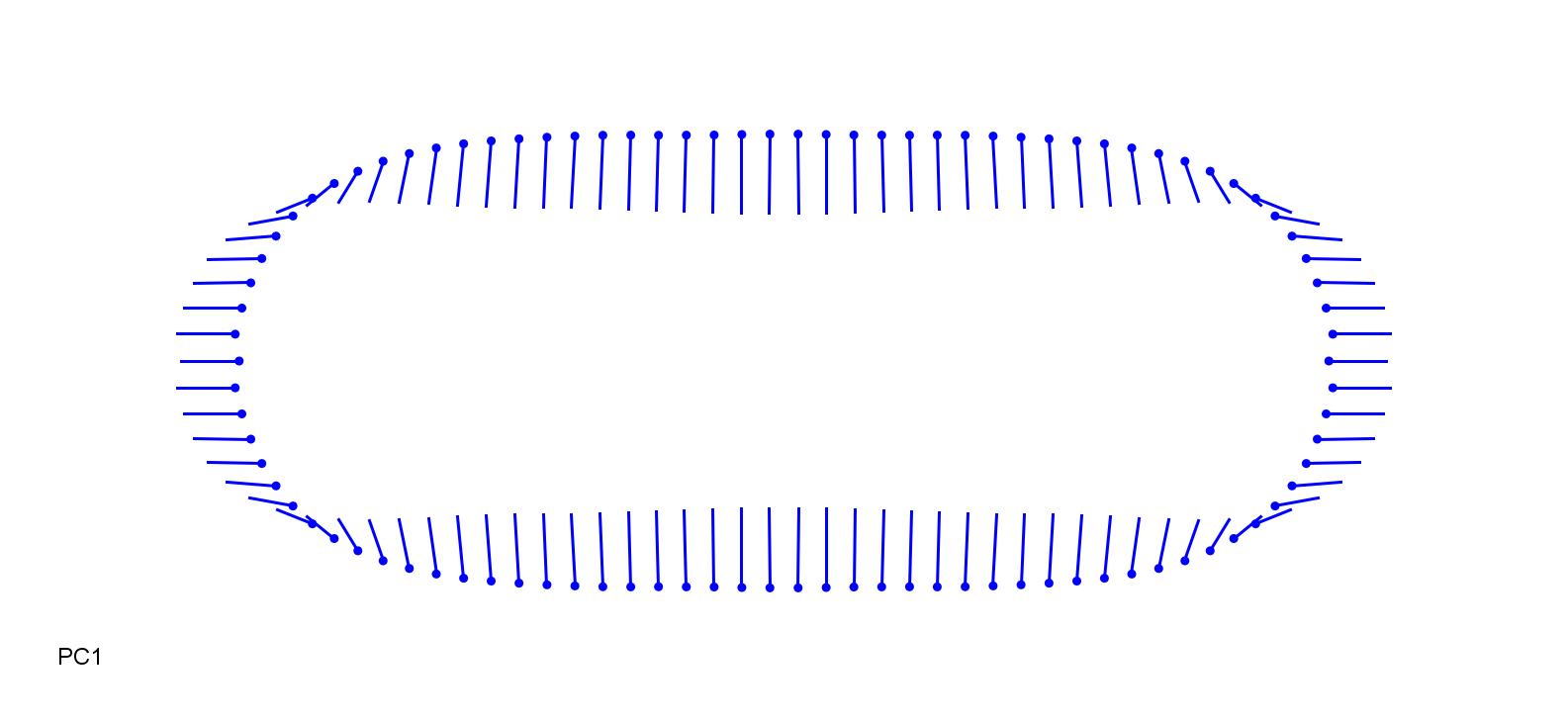

### HelPub-0.2.jpg

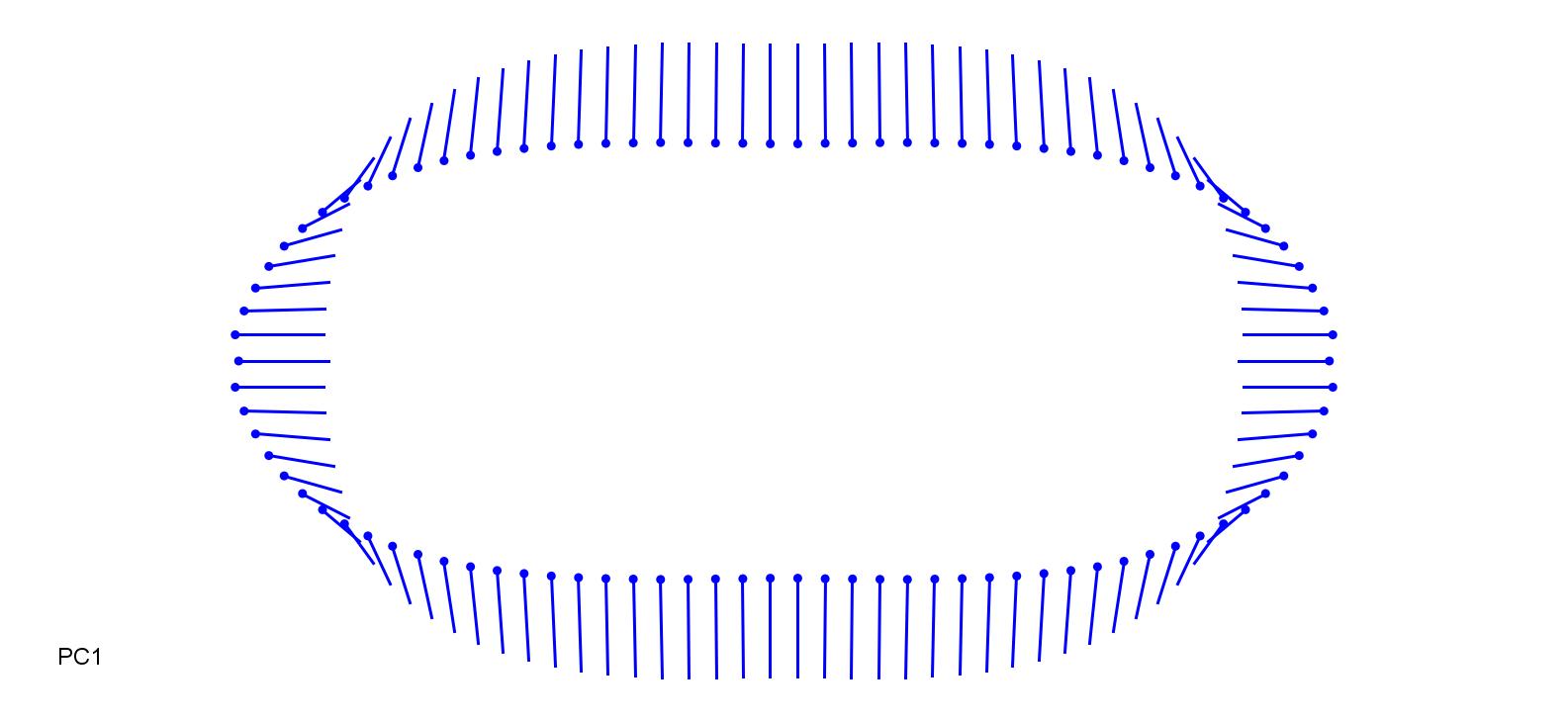

### HelPub_0.2.jpg

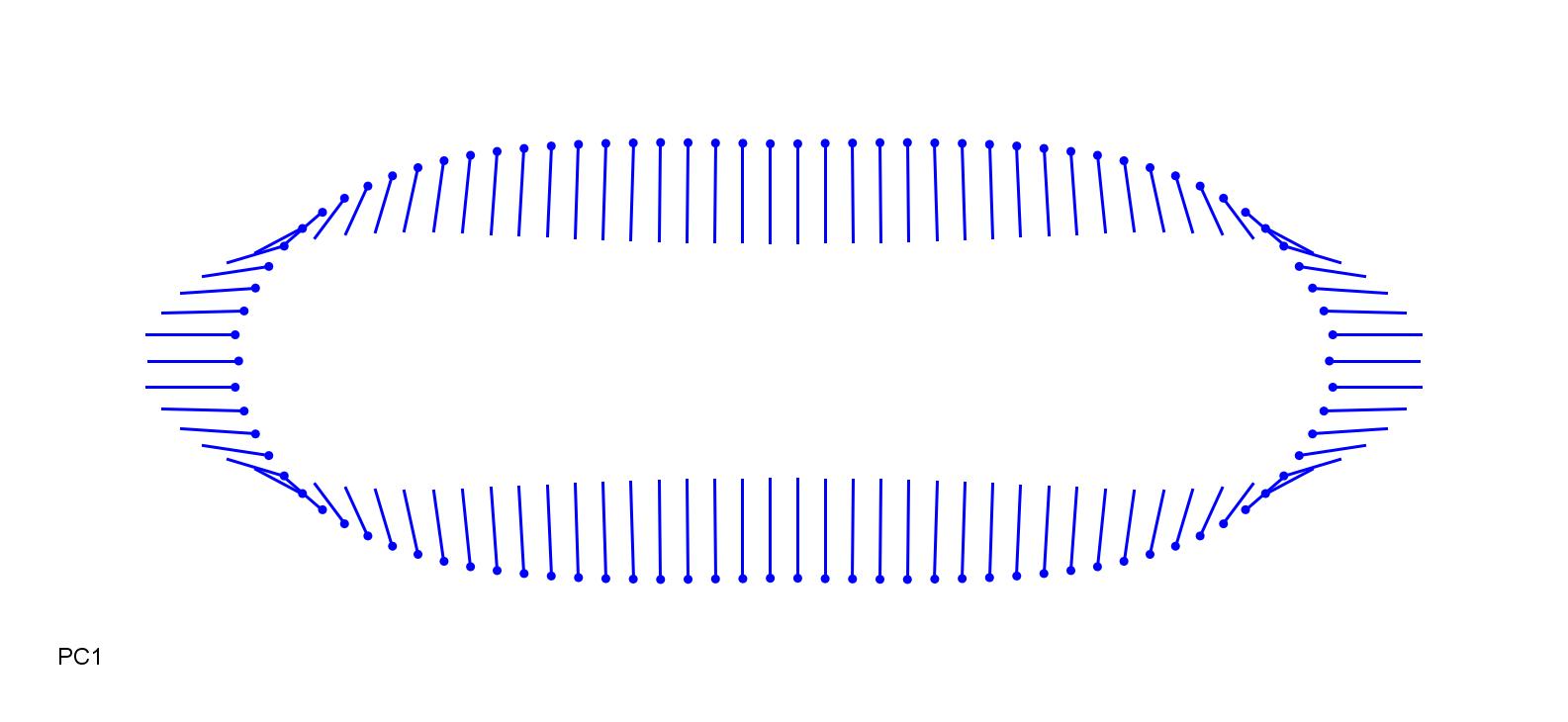

### HolLan-0.2.jpg

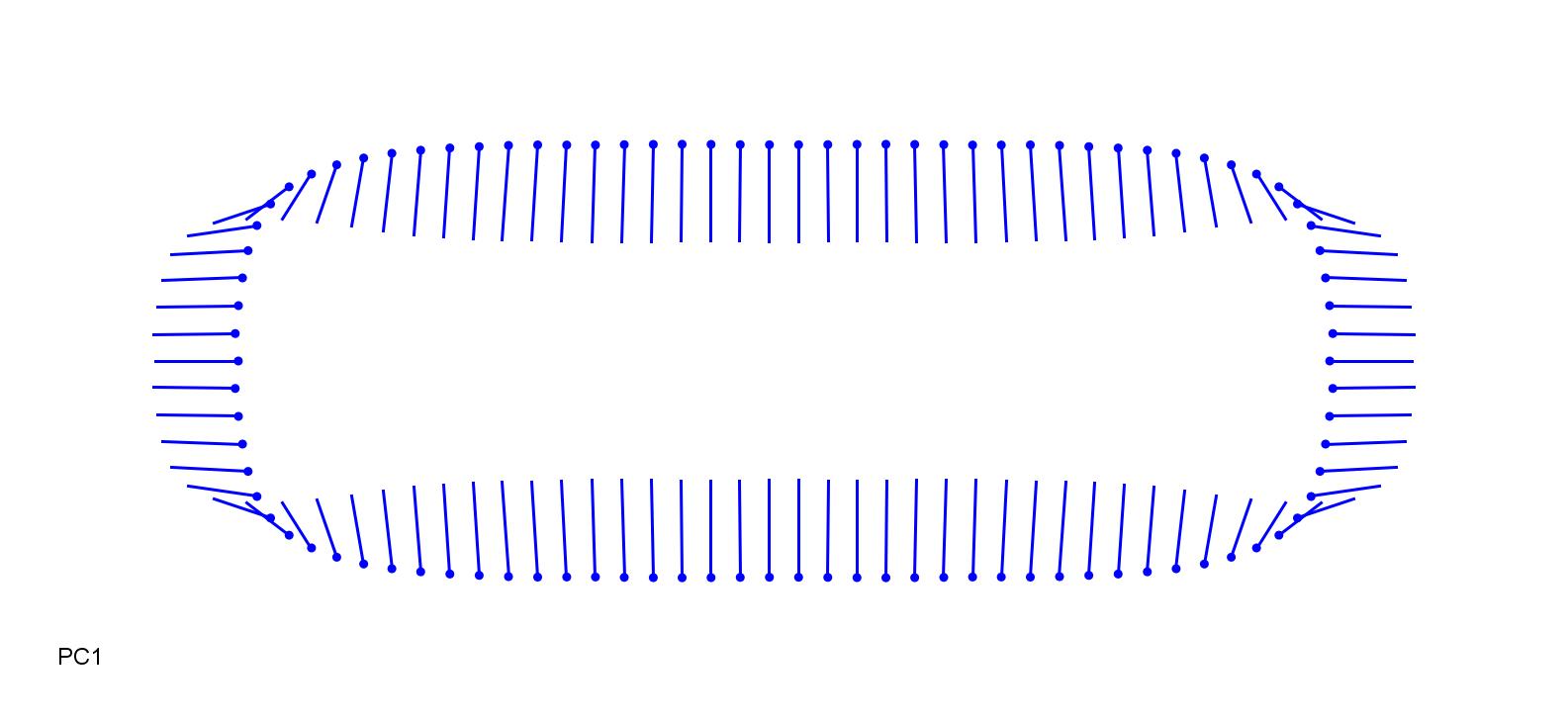

### HolLan_0.25.jpg

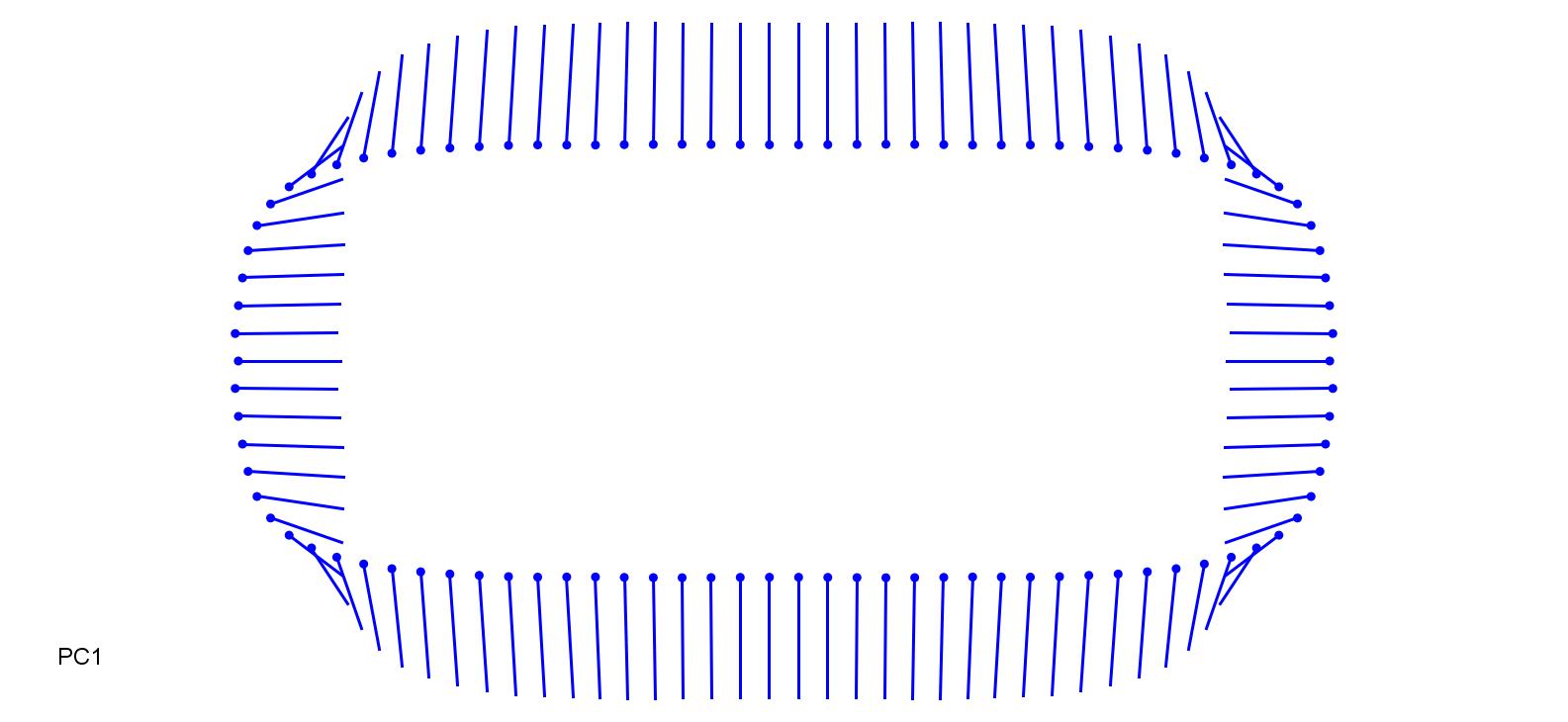

### HolMol-0.2.jpg

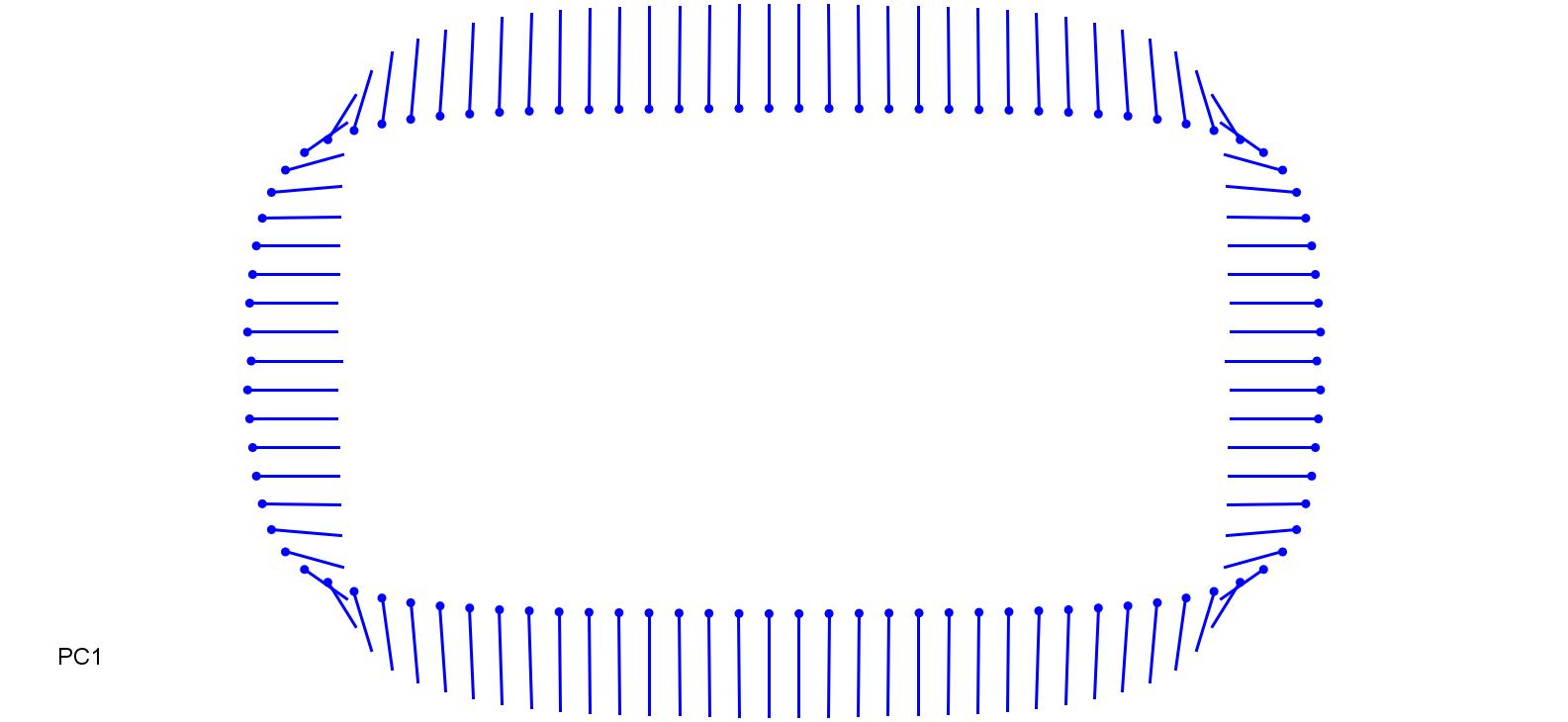

### HolMol_0.2.jpg

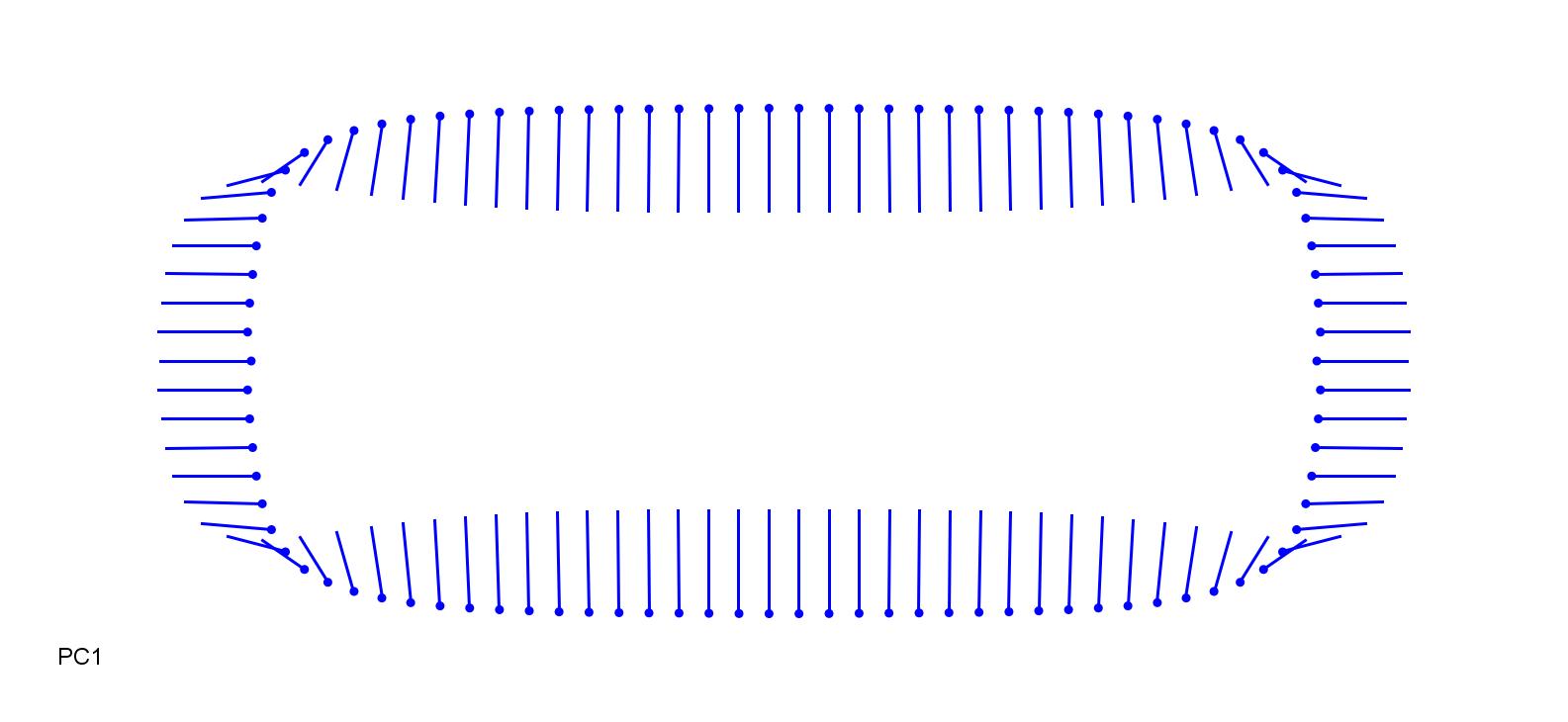

### KoeGla_0.1.jpg

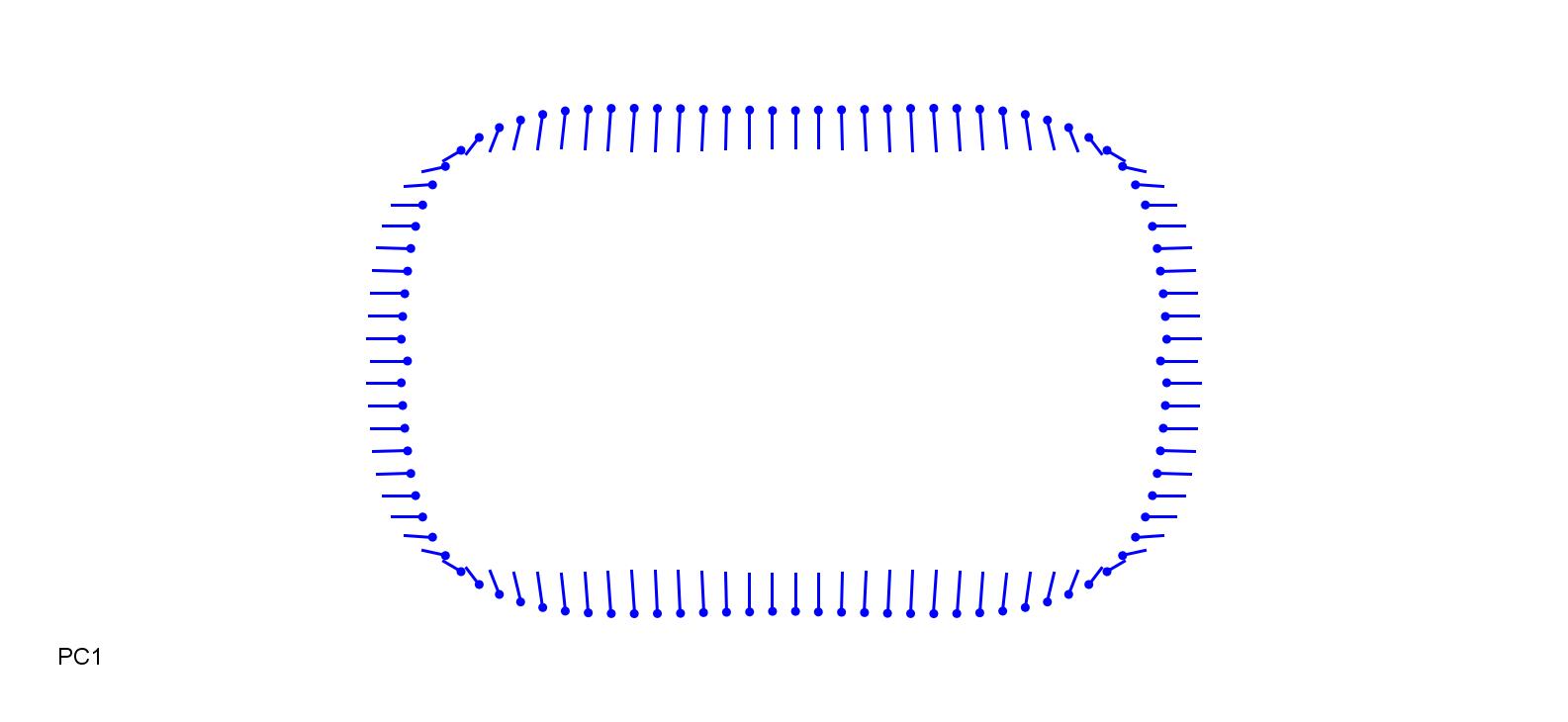

### KoeMac-0.2.jpg

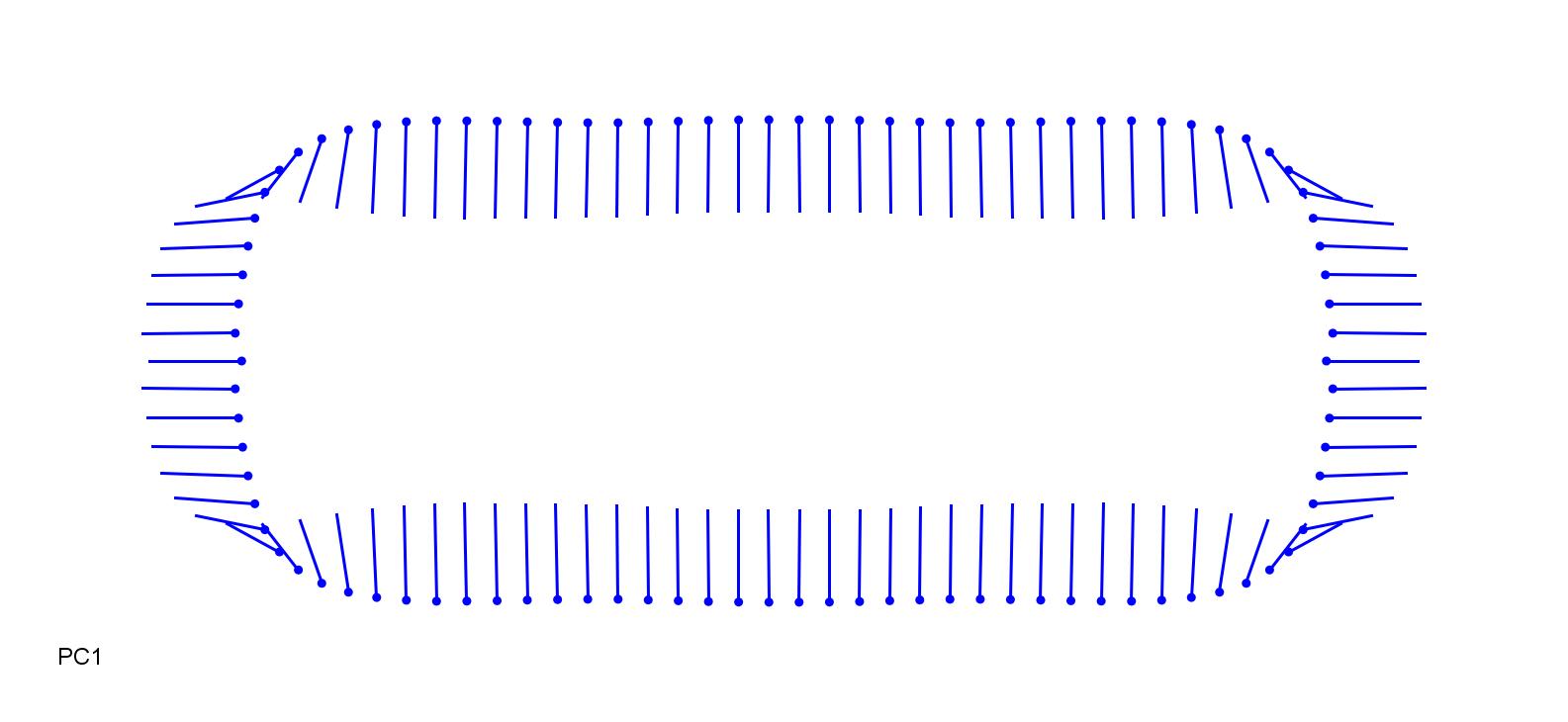

### KoeMac_0.2.jpg

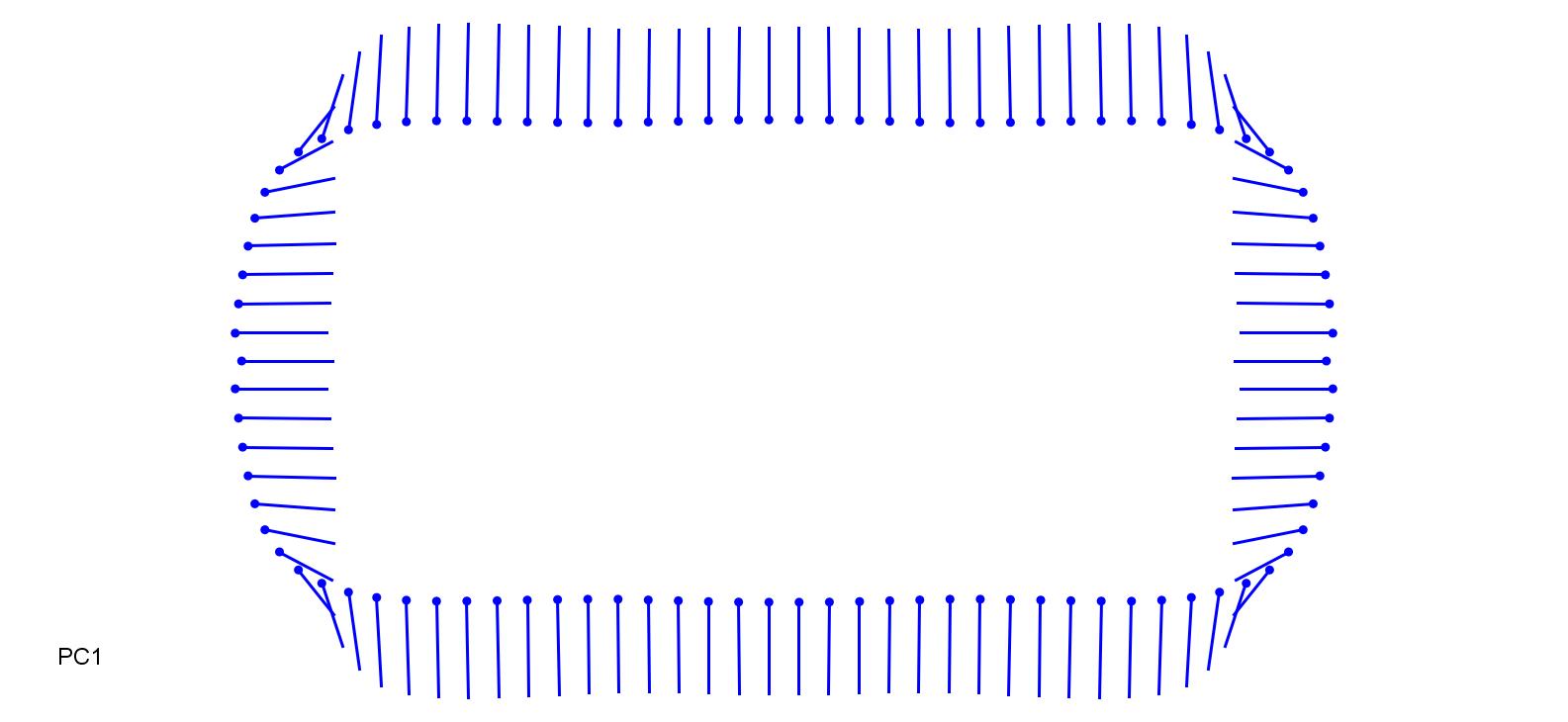

### LeeOry-0.2.jpg

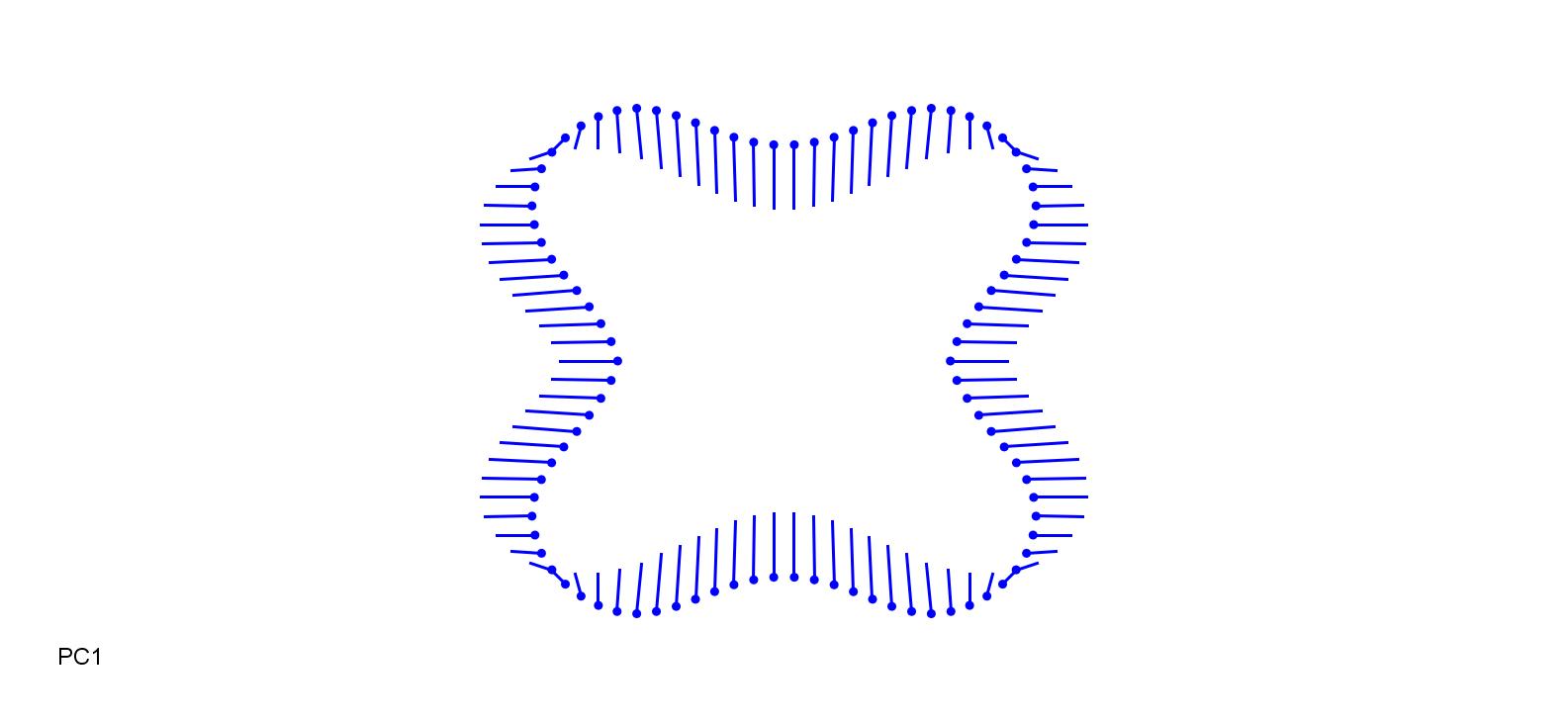

### LeeOry_0.2.jpg

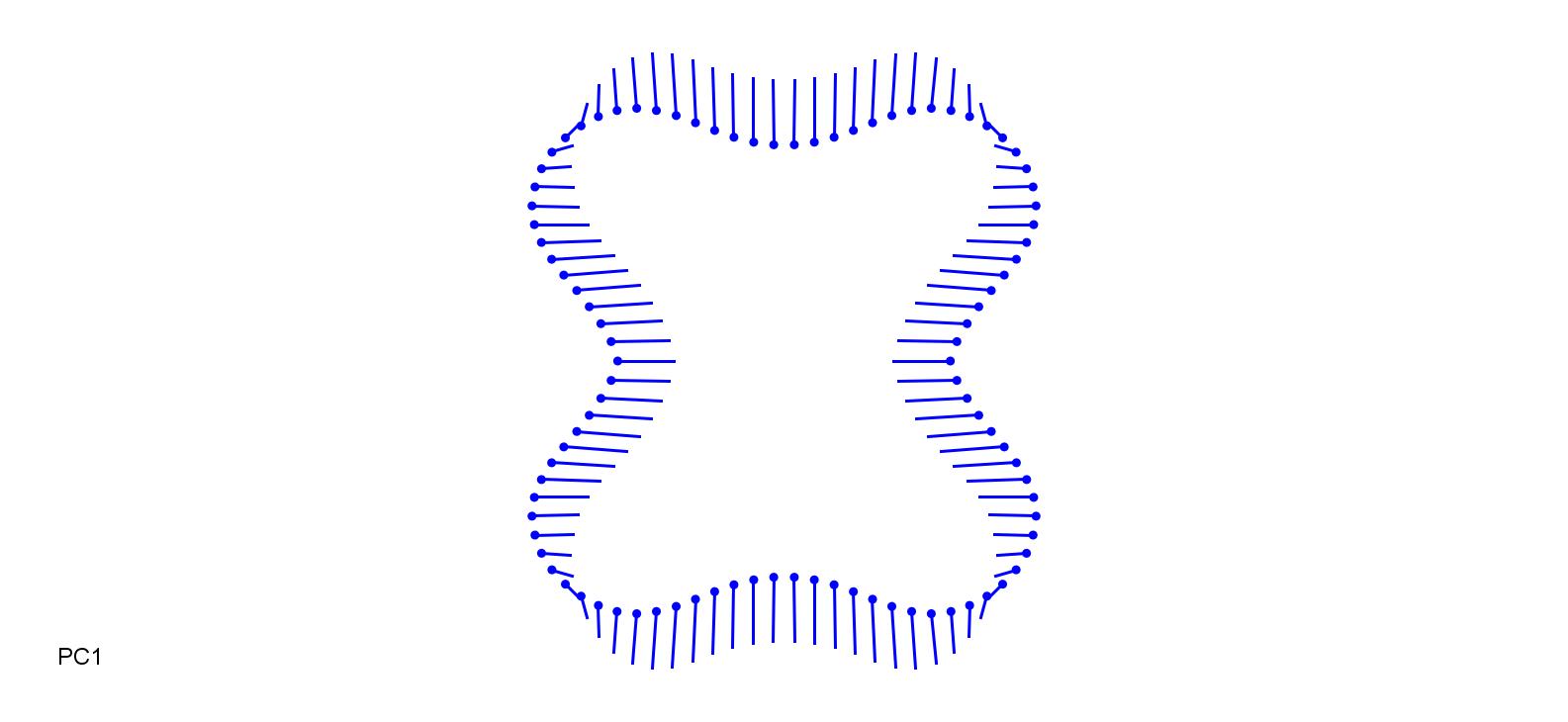

### LolPer-0.25.jpg

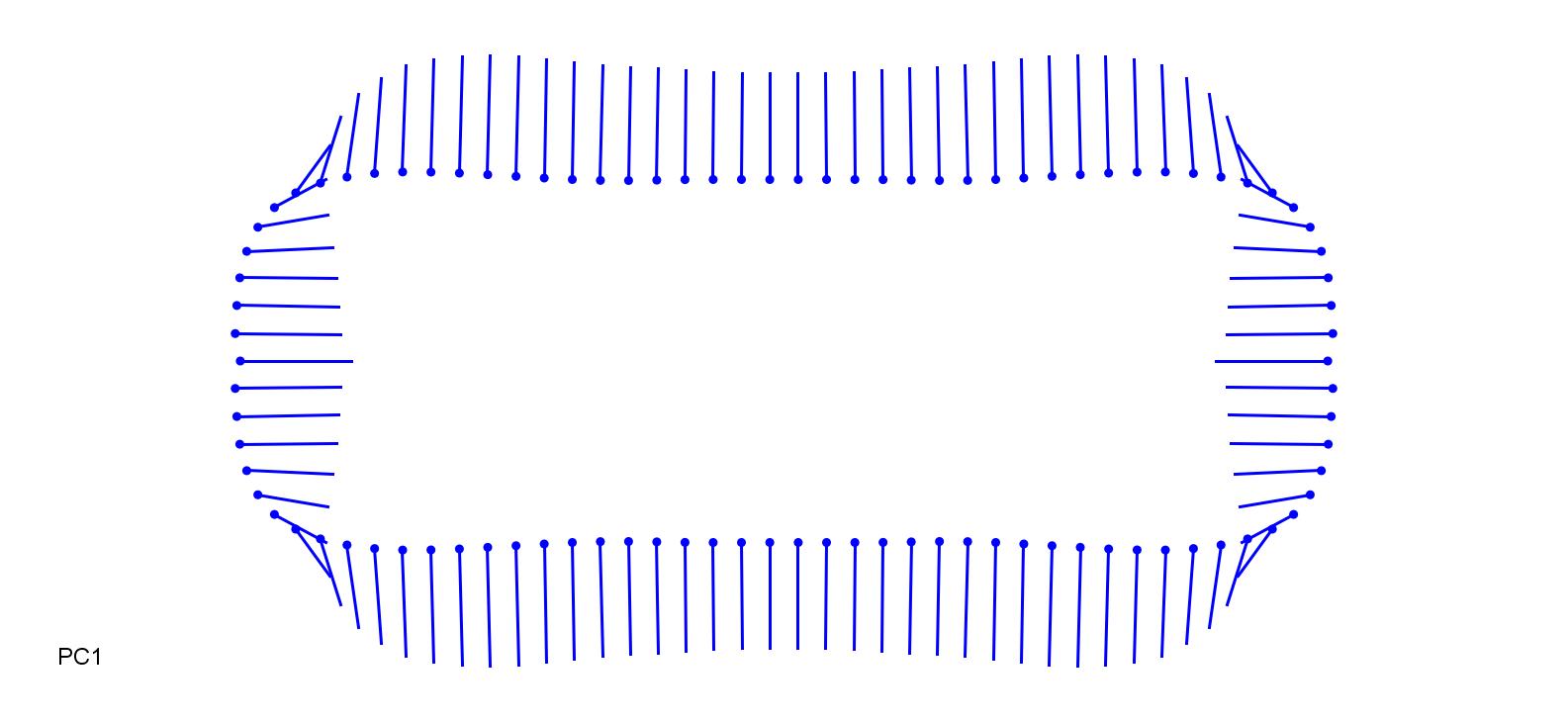

### LolPer_0.15.jpg

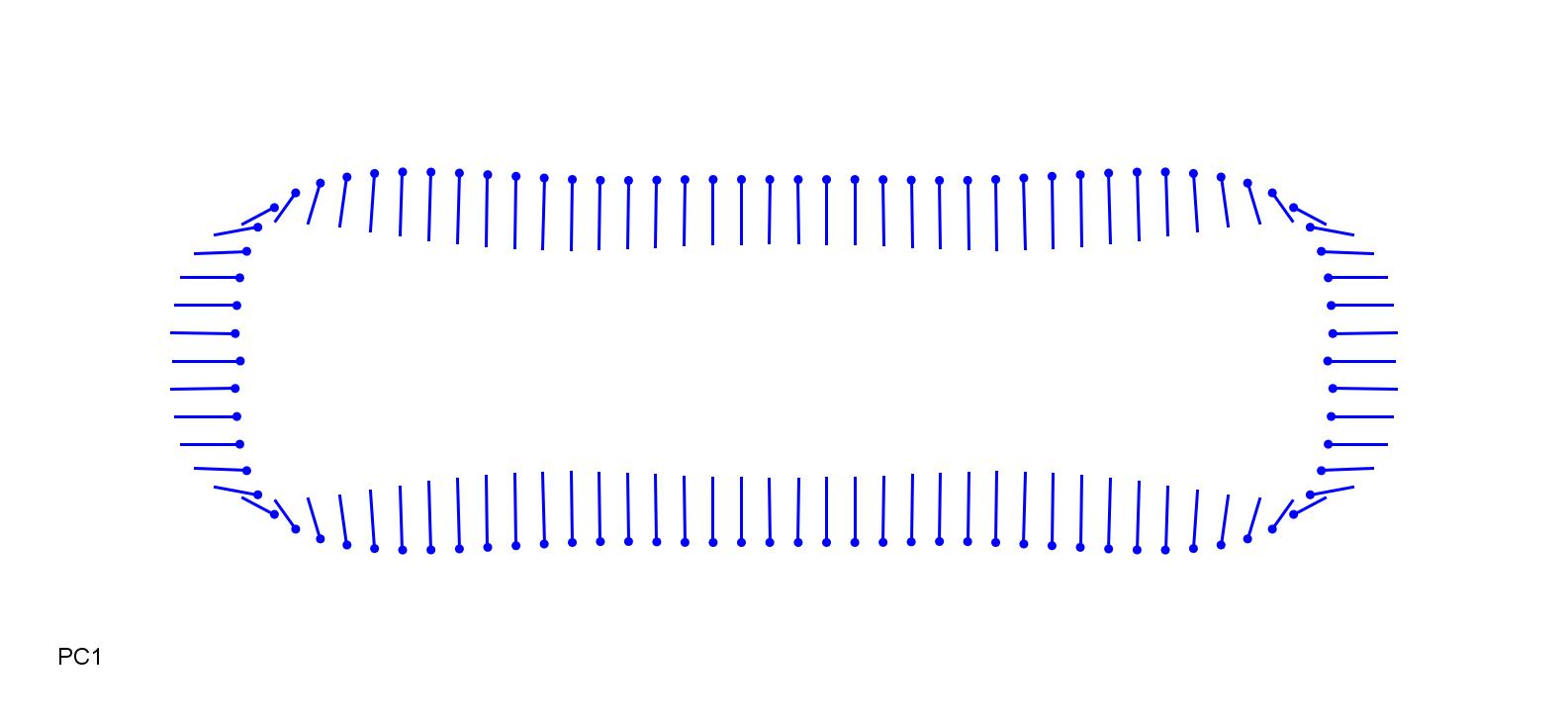

### MelRep-0.2.jpg

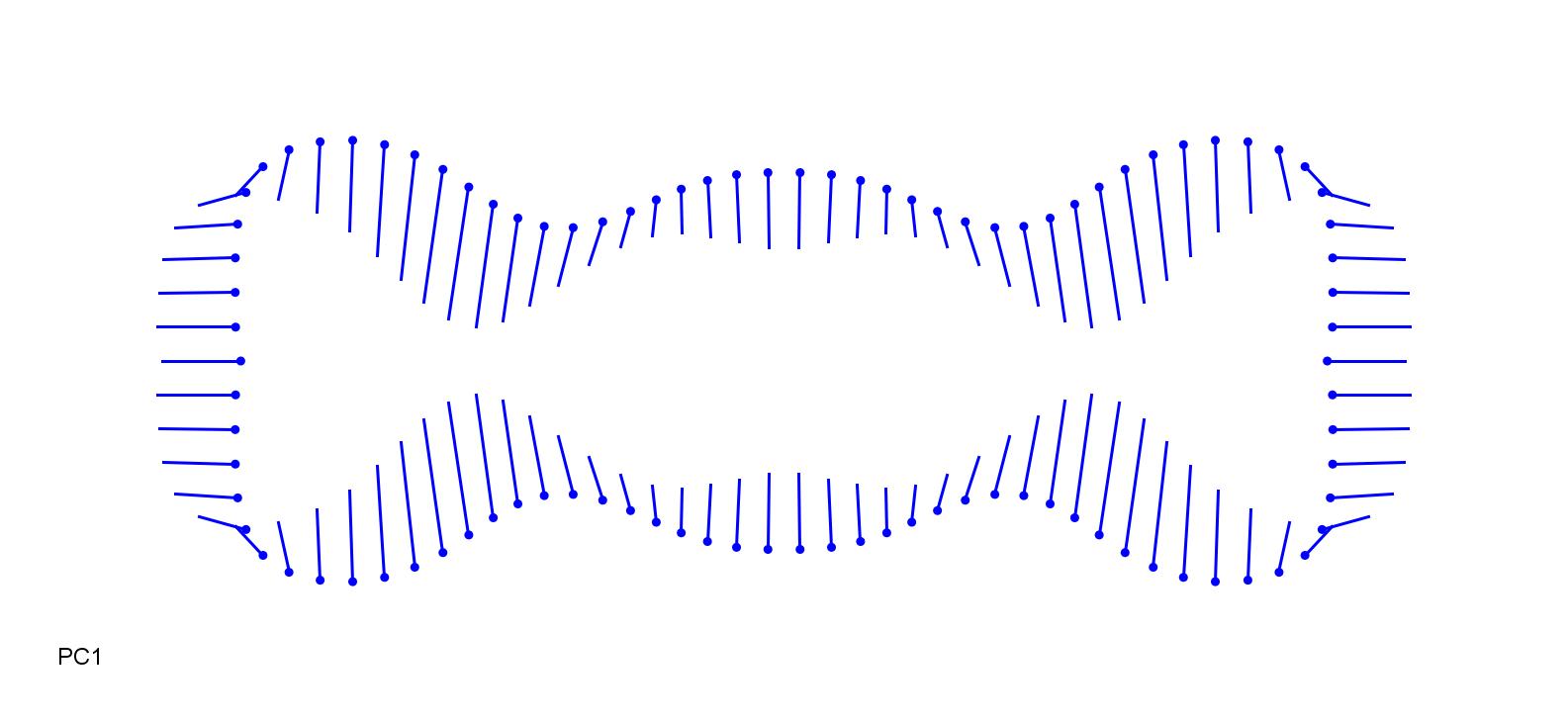

### MelRep_0.2.jpg

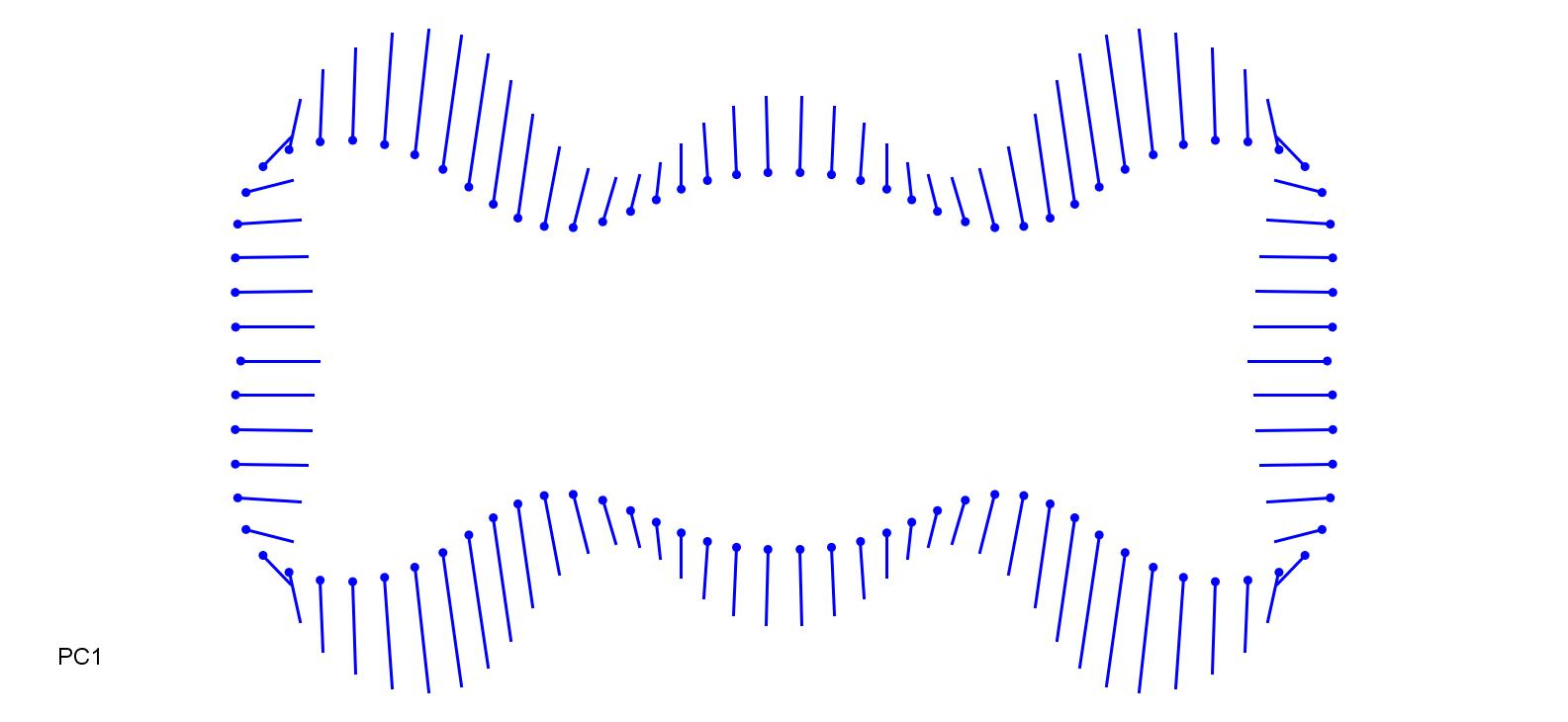

### MilEff-0.2.jpg

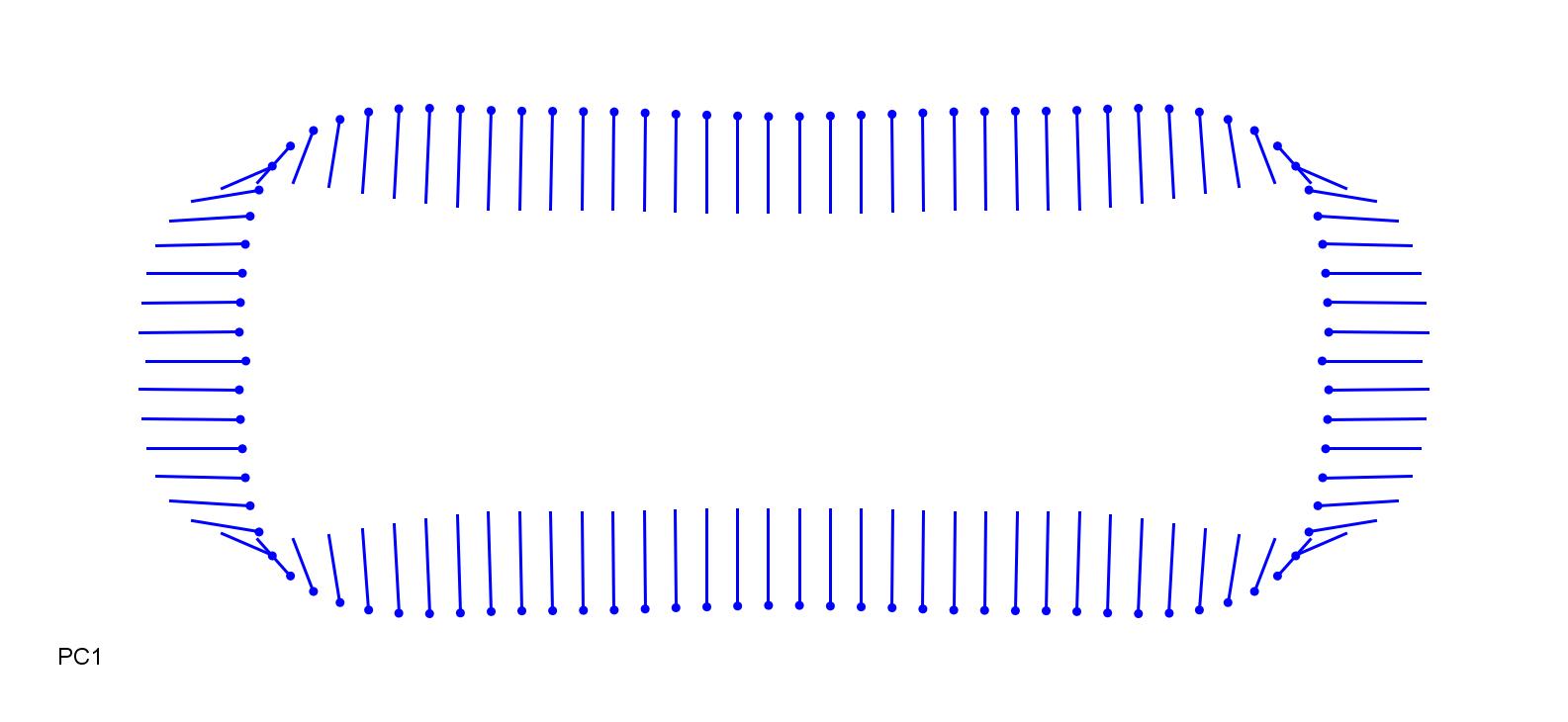

### MilEff_0.2.jpg

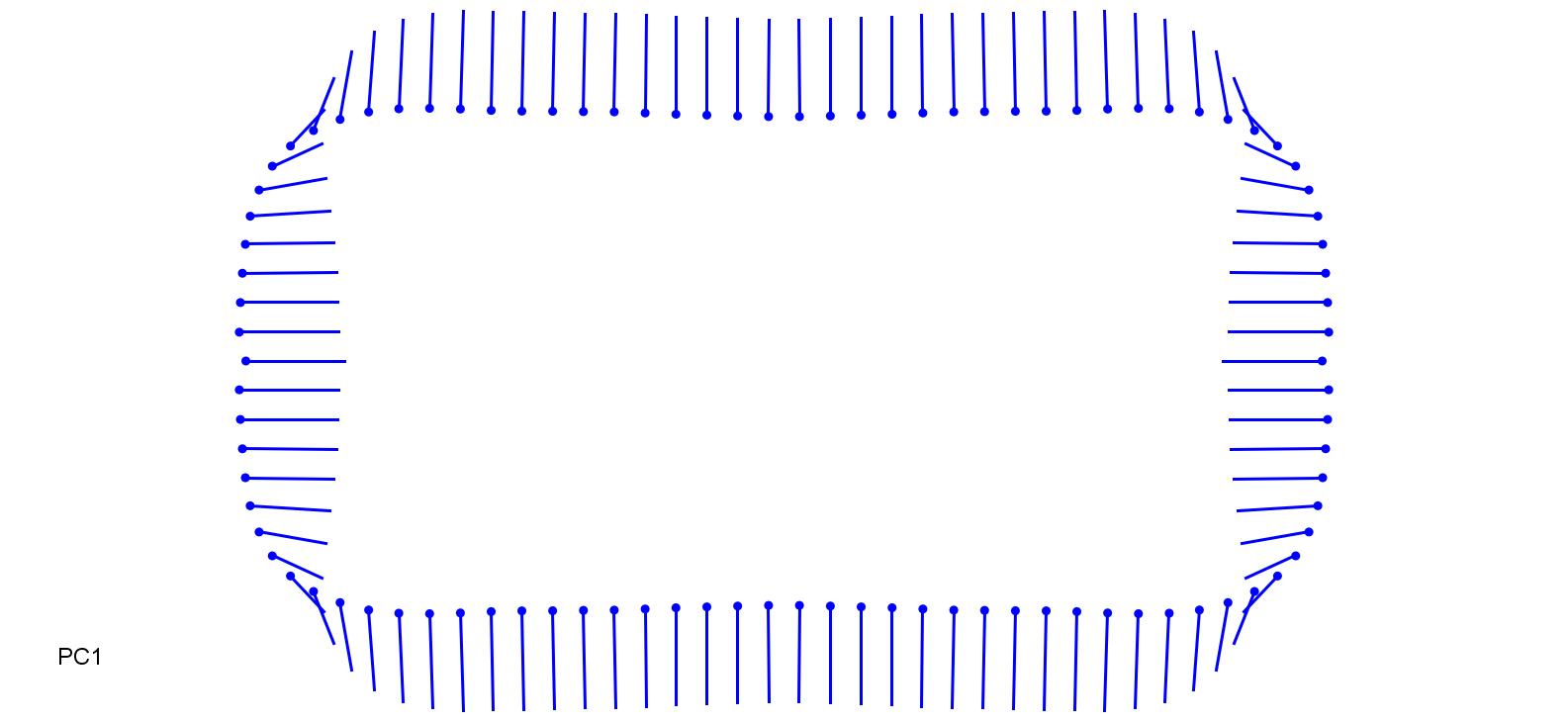

### MolCae_0.2.jpg

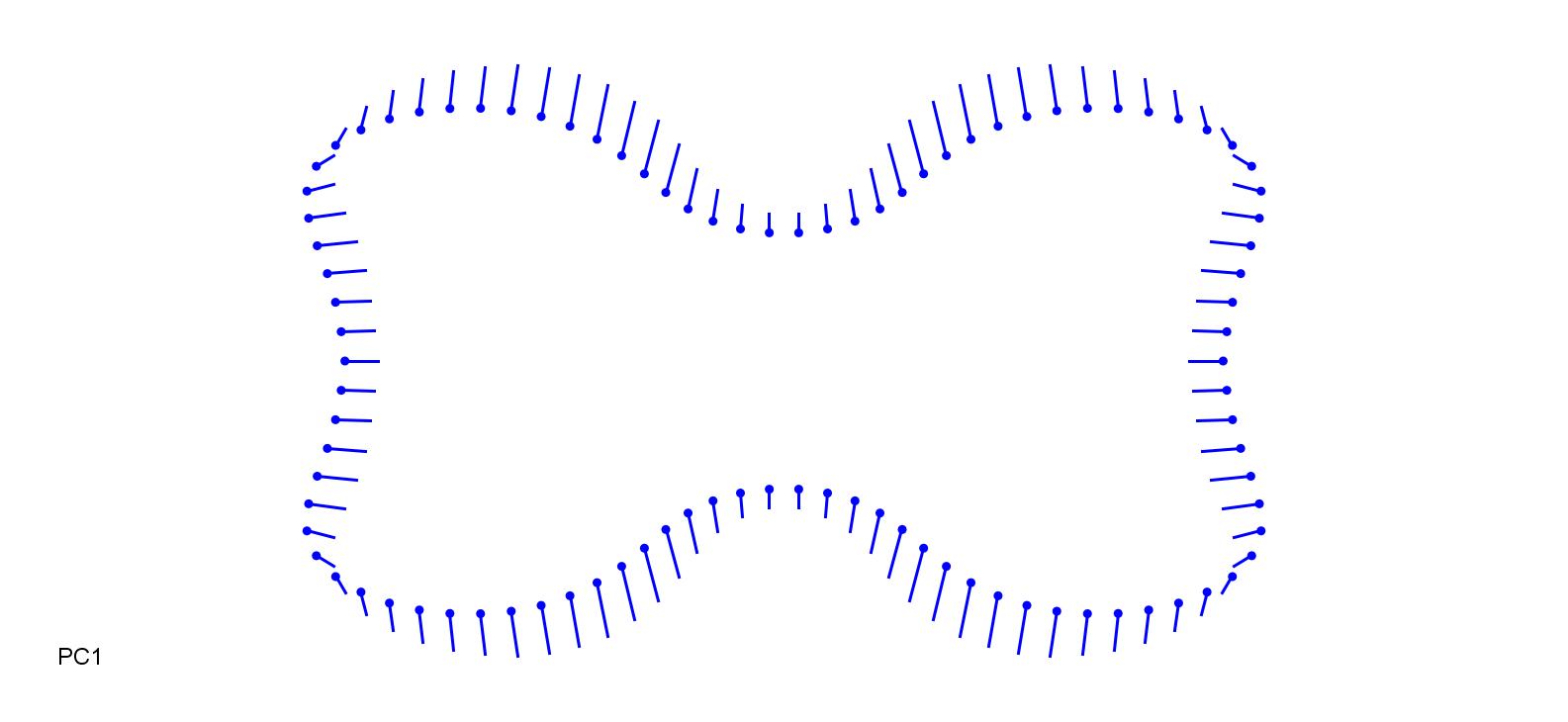

### OrySat-0.1.jpg

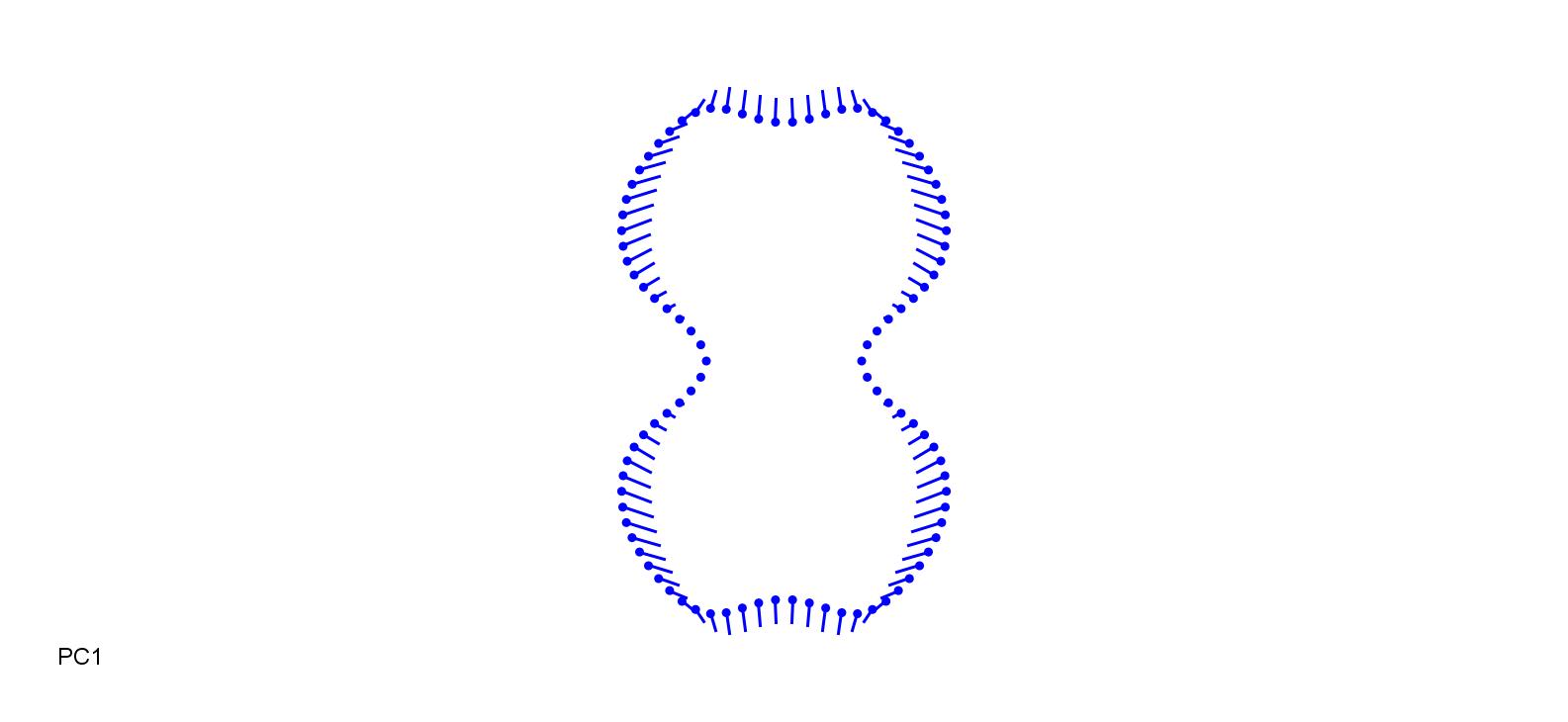

### OrySat_0.13.jpg

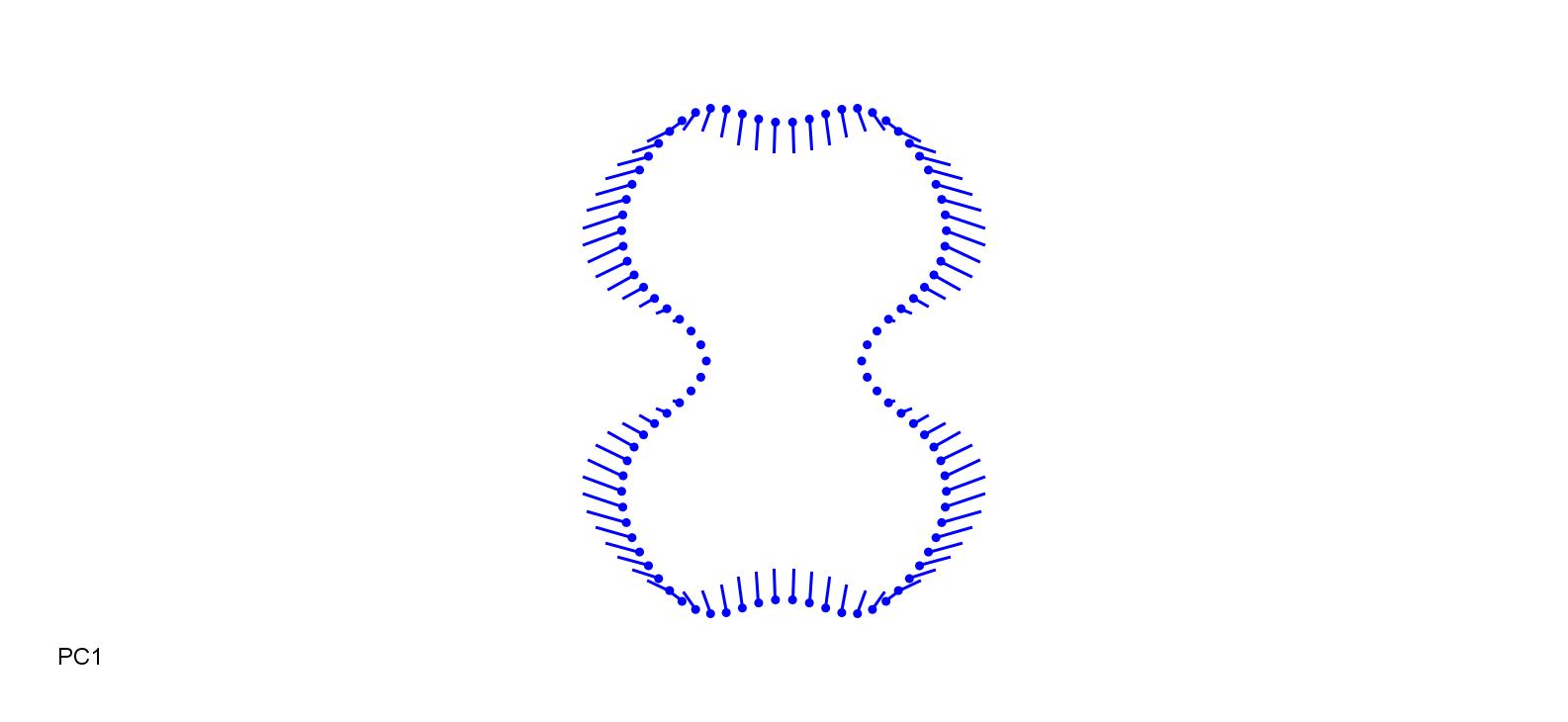

### PhlPrat-0.3.jpg

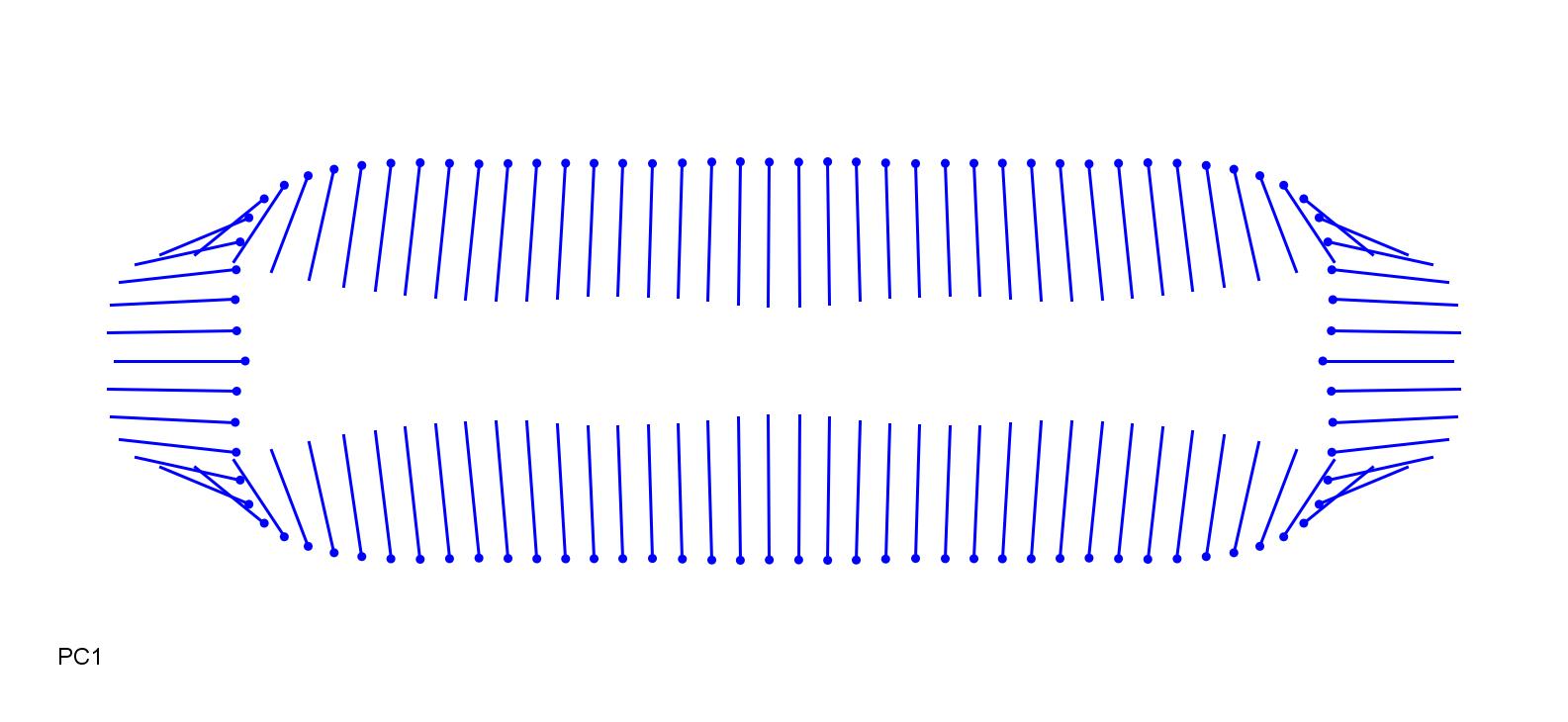

### PhlPrat_0.25.jpg

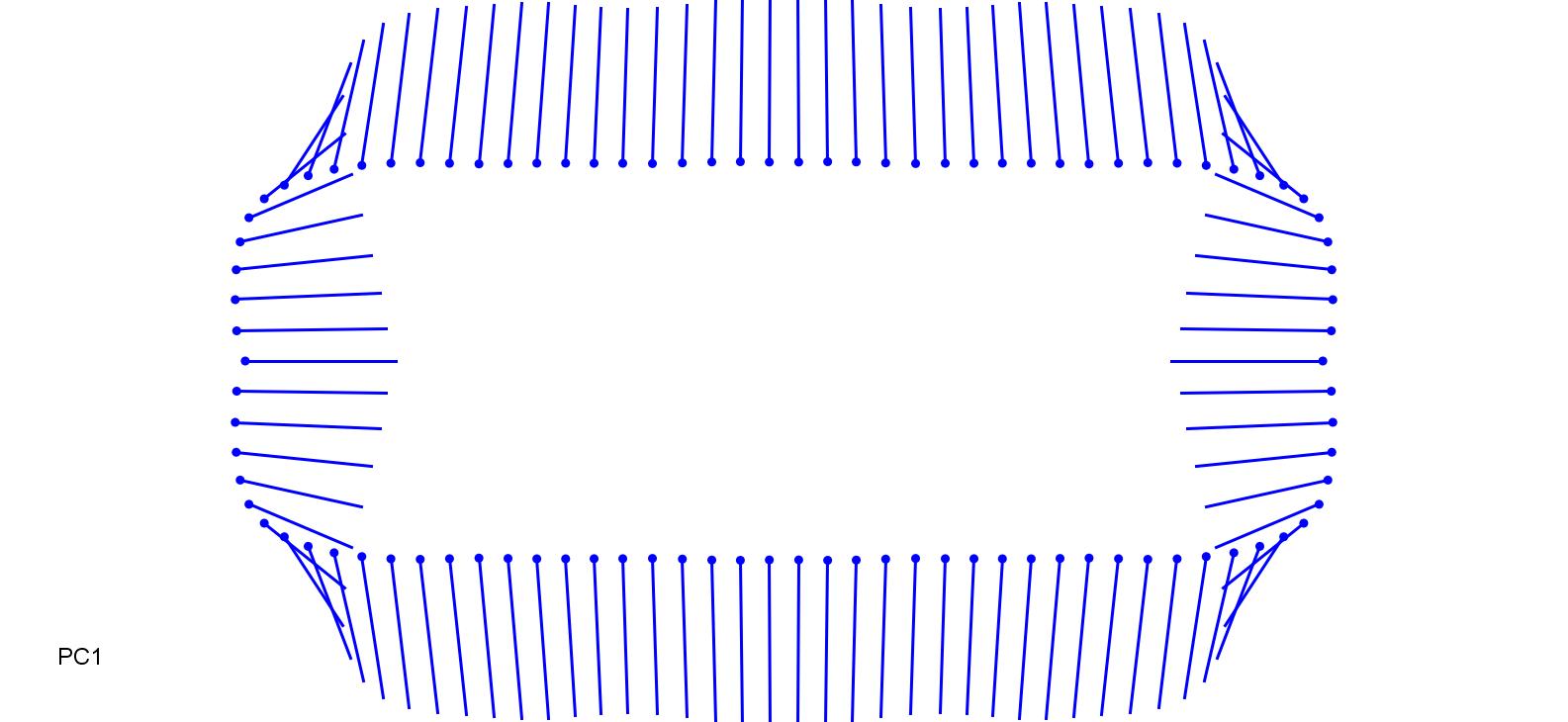

### PhrAus-0.1.jpg

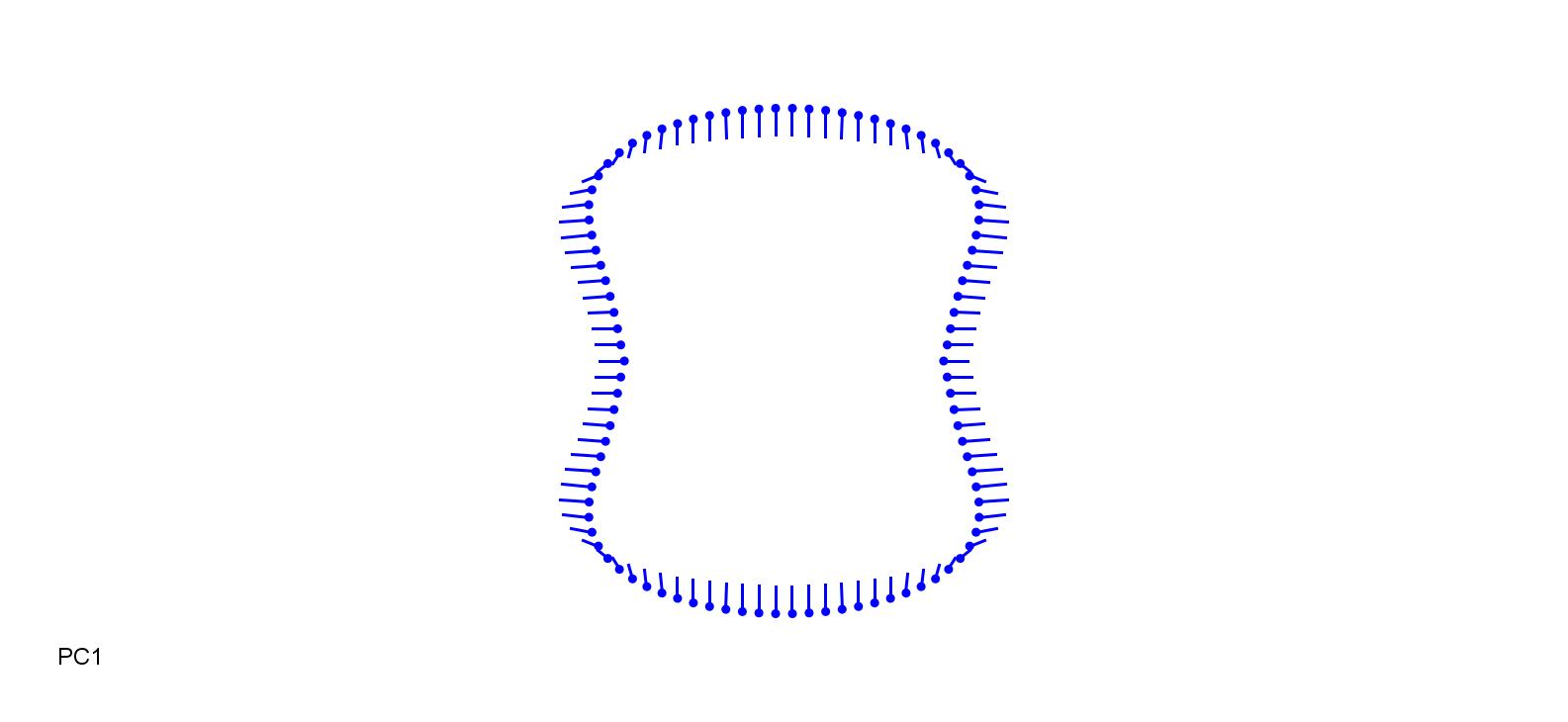

### PhrAus_0.1.jpg

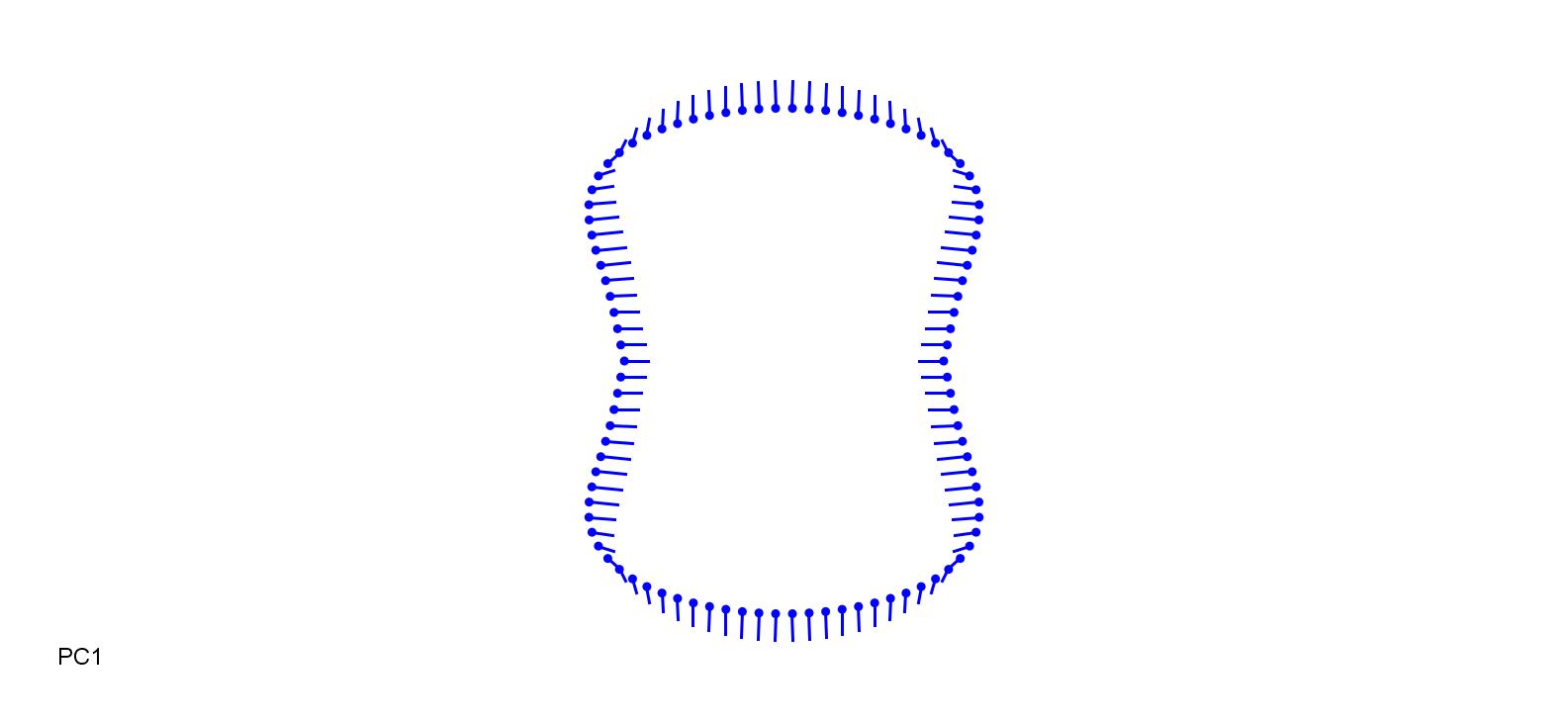

### PhrKar-0.2.jpg

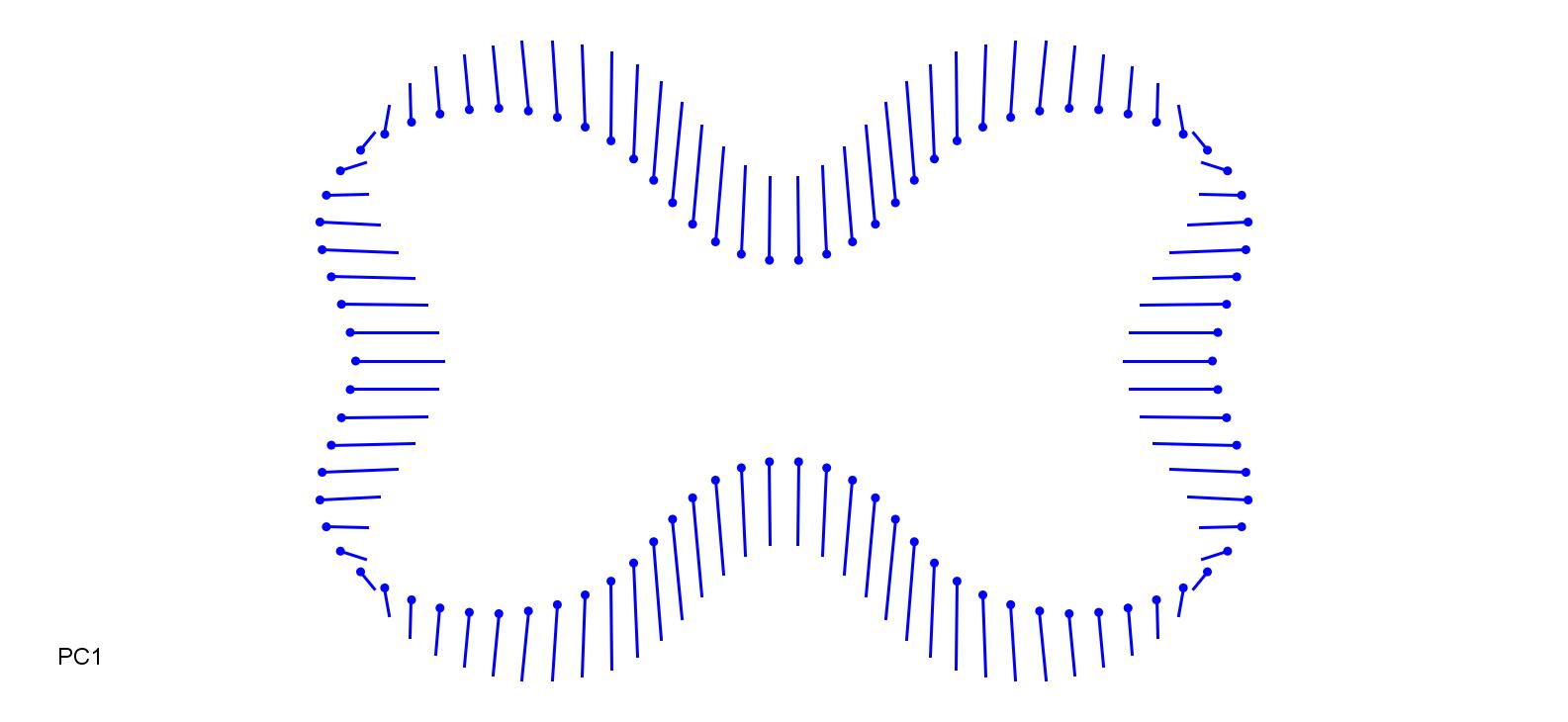

### PhrKar_0.2.jpg

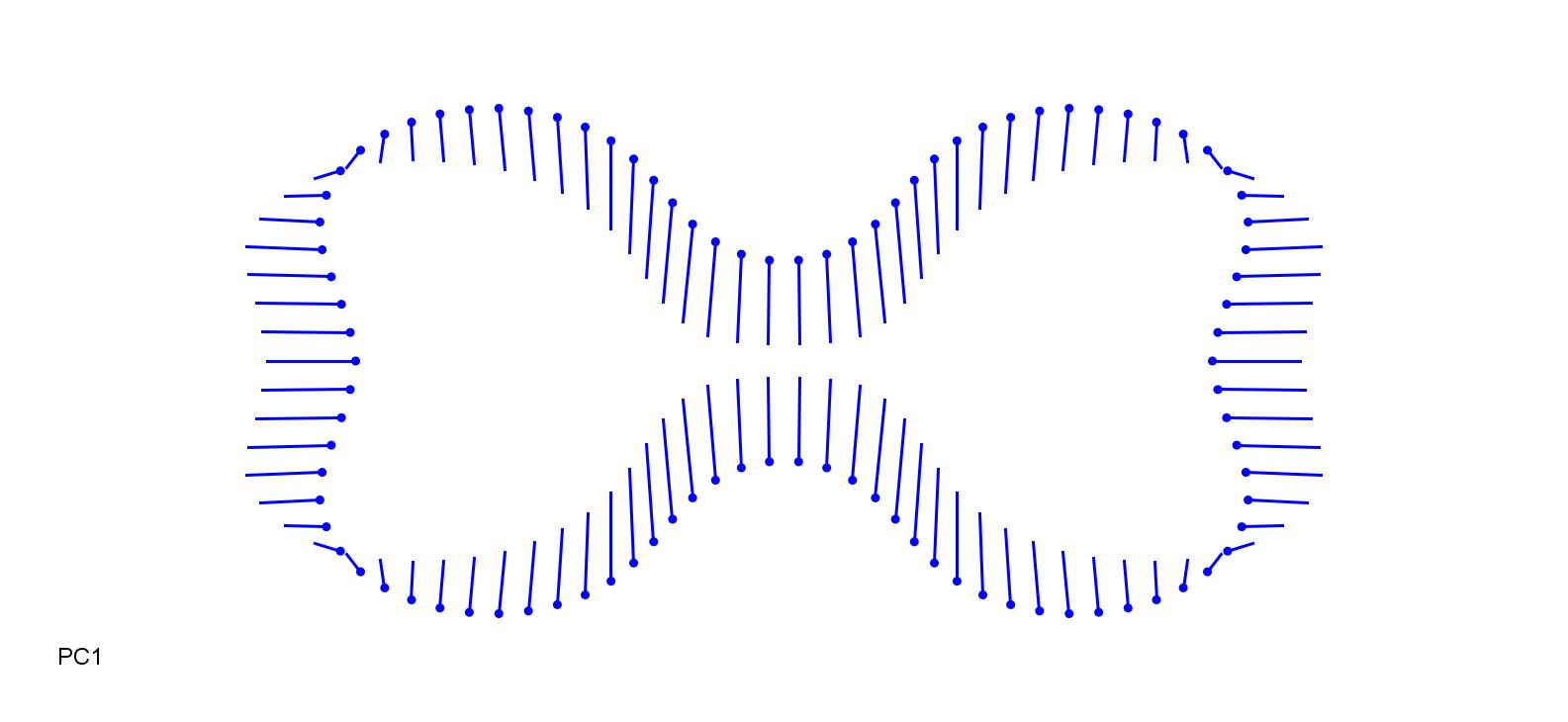

### PoaChai-0.2.jpg

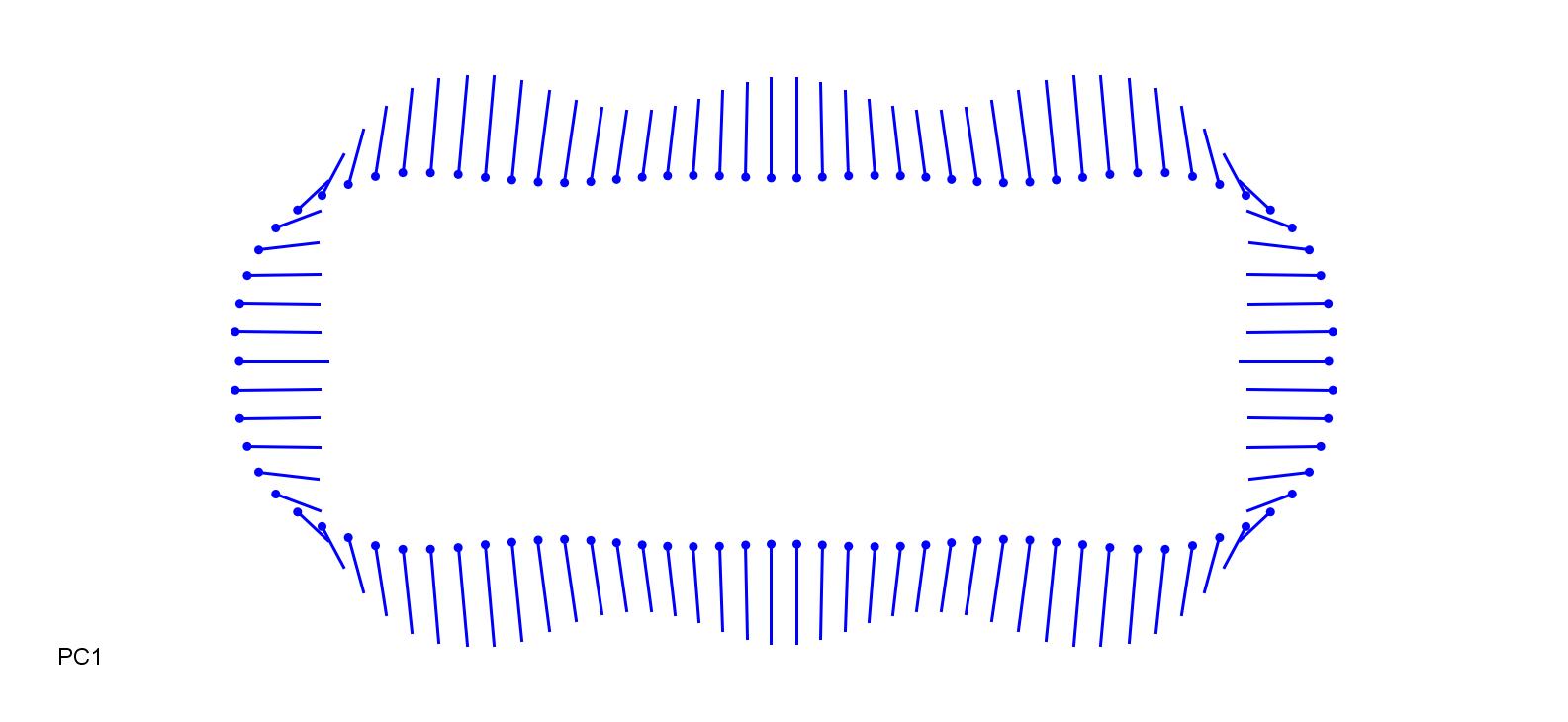

### PoaChai_0.2.jpg

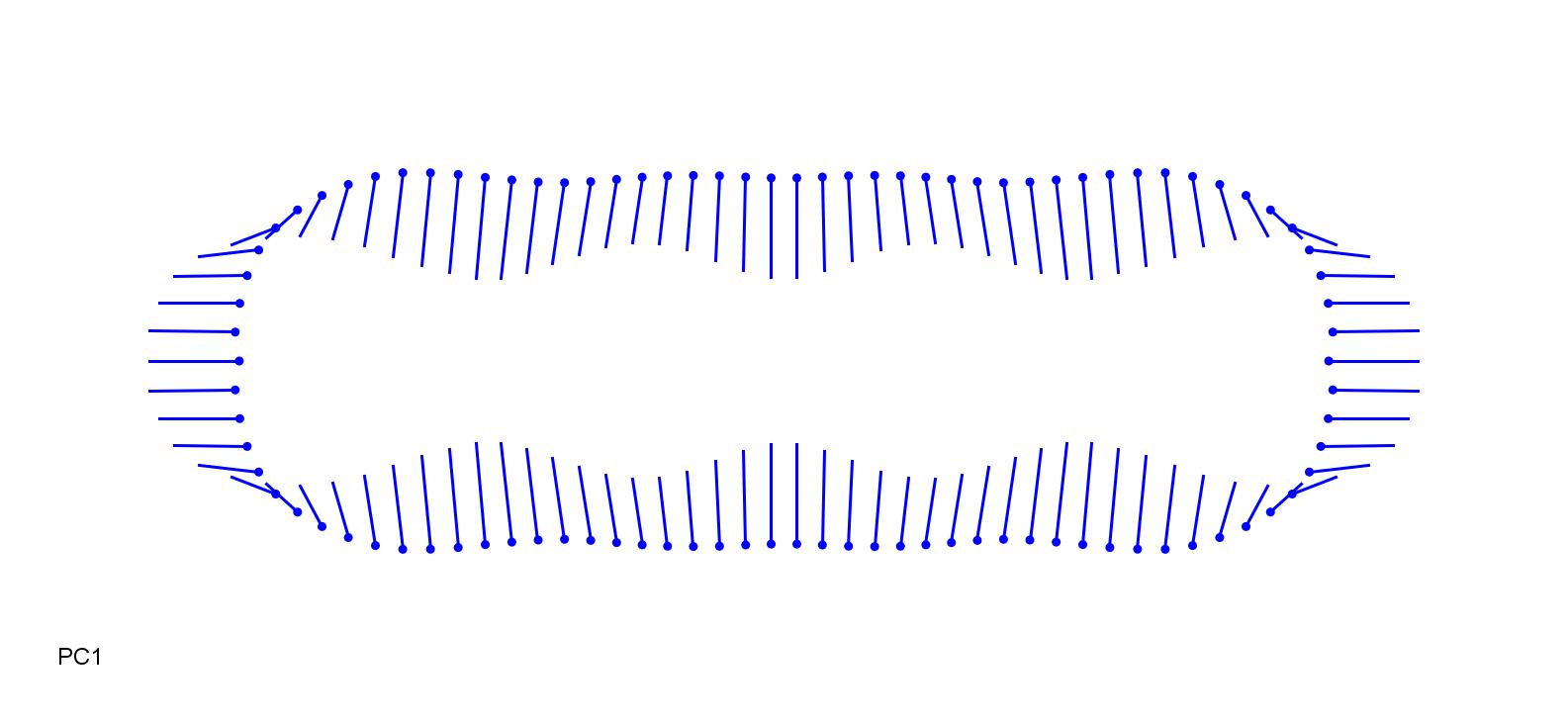

### PoaNem-0.15.jpg

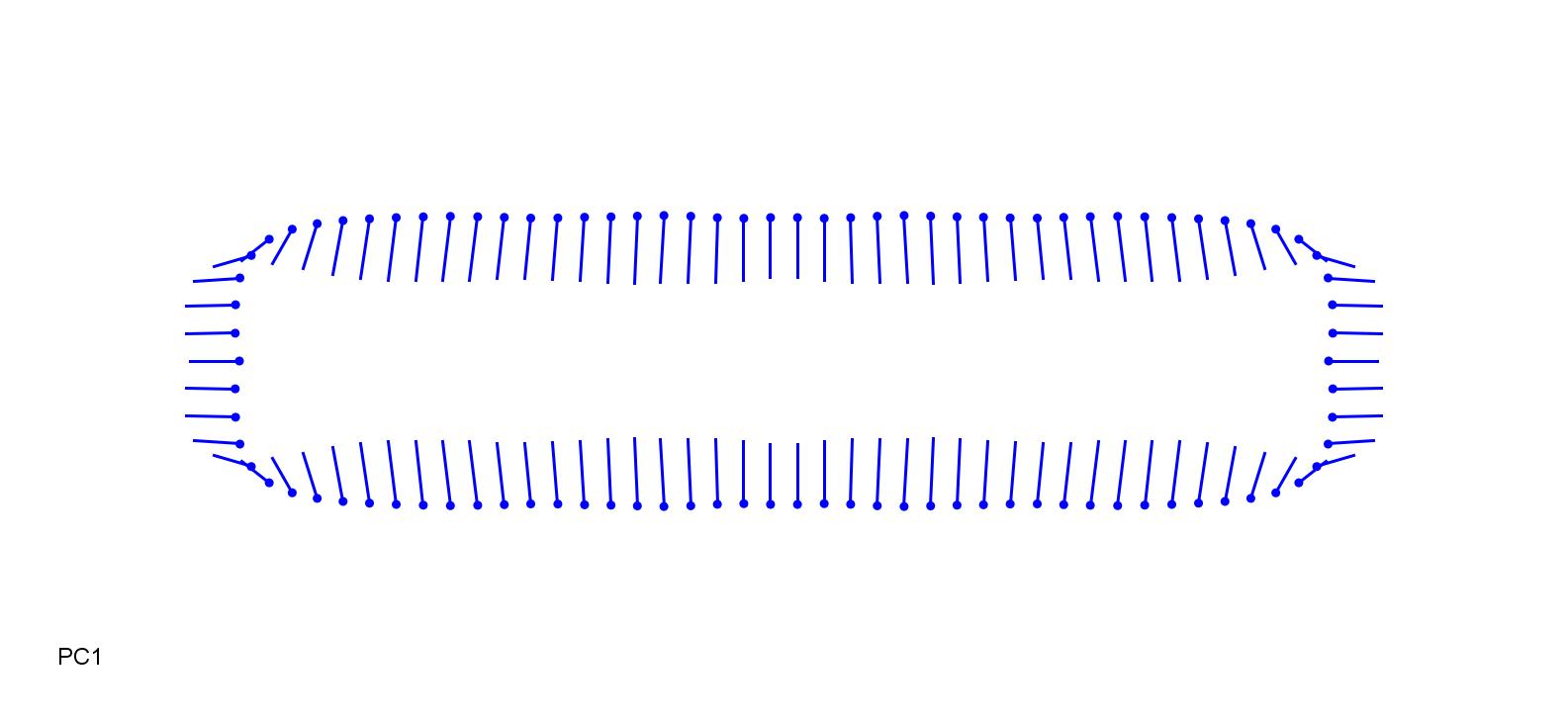

### PoaNem_0.25.jpg

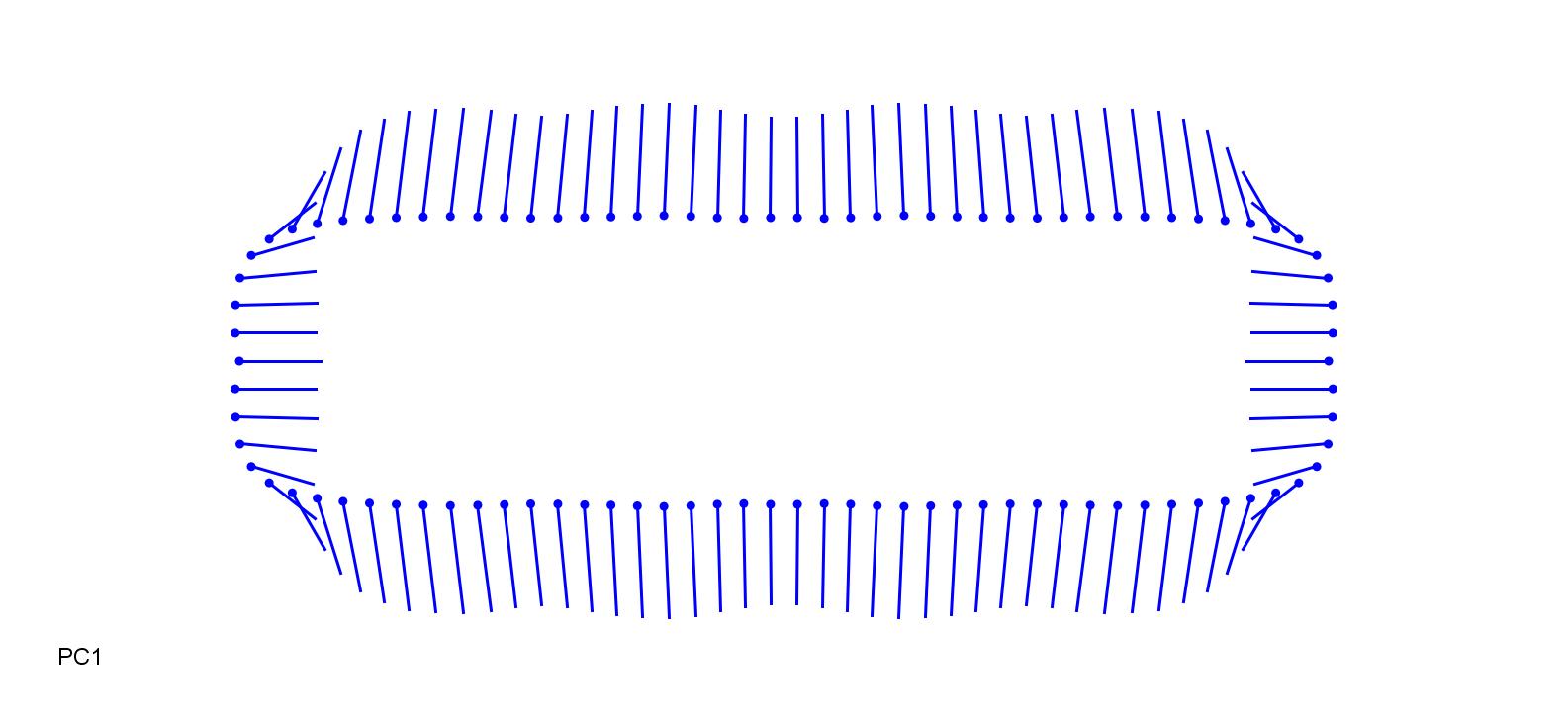

### SchiAra-0.15.jpg

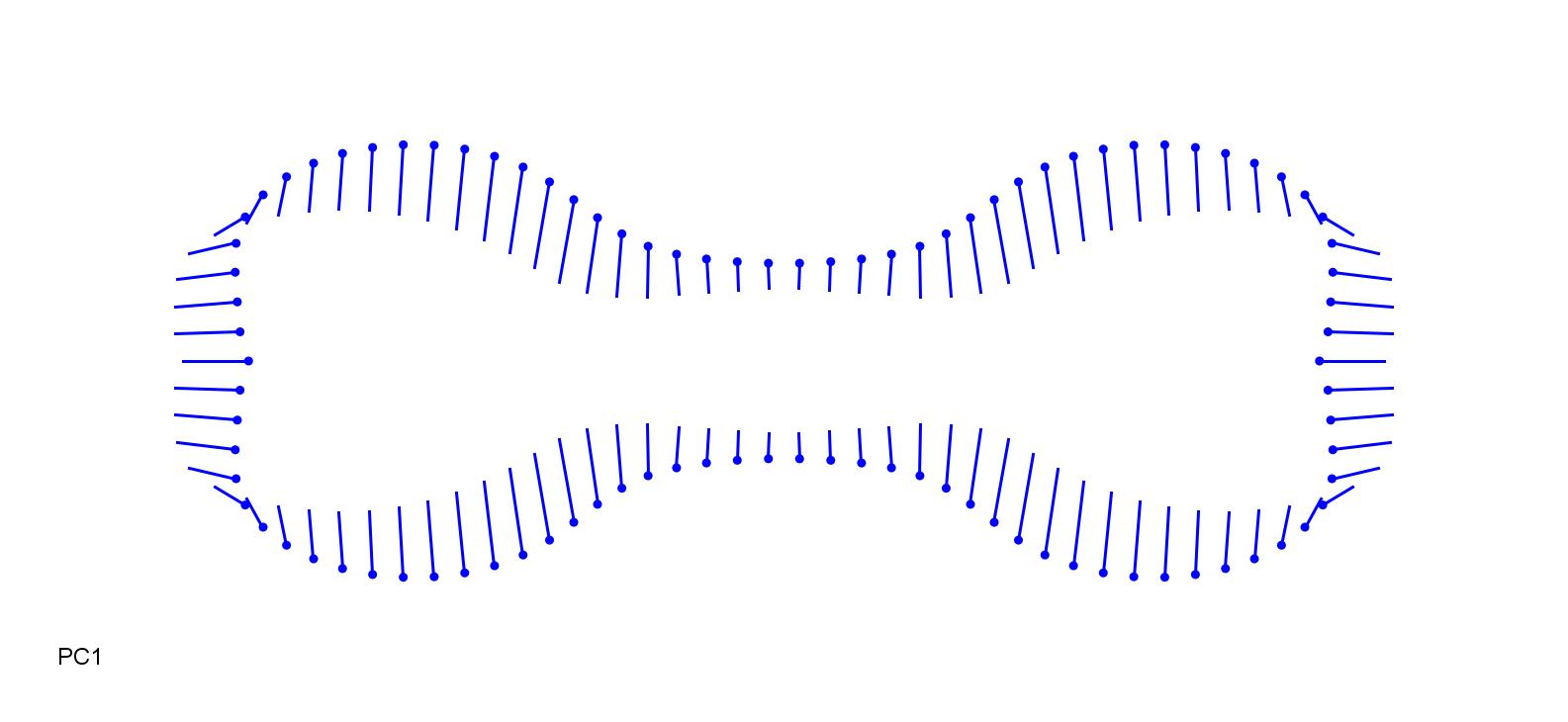

### SchiAra_0.2.jpg

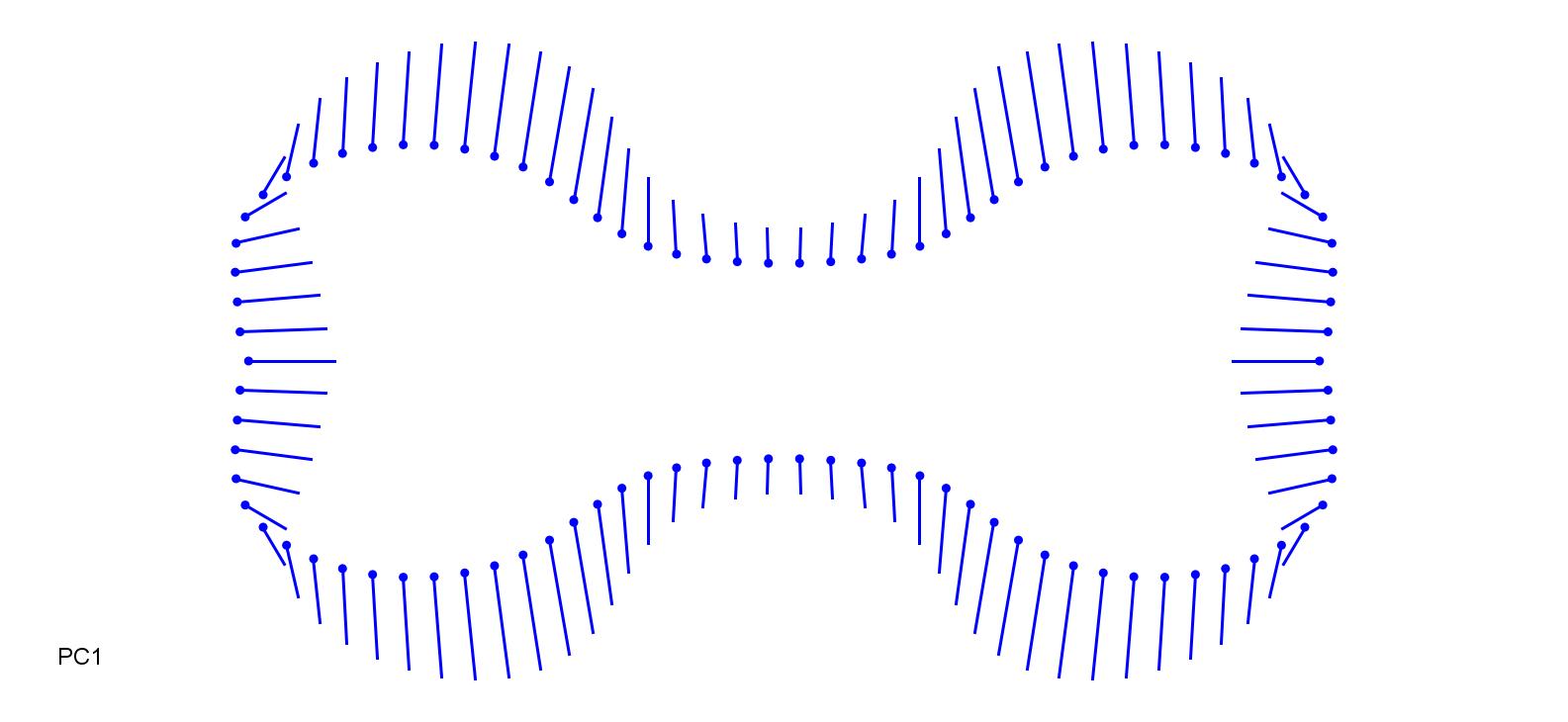

### SchiBar-0.2.jpg

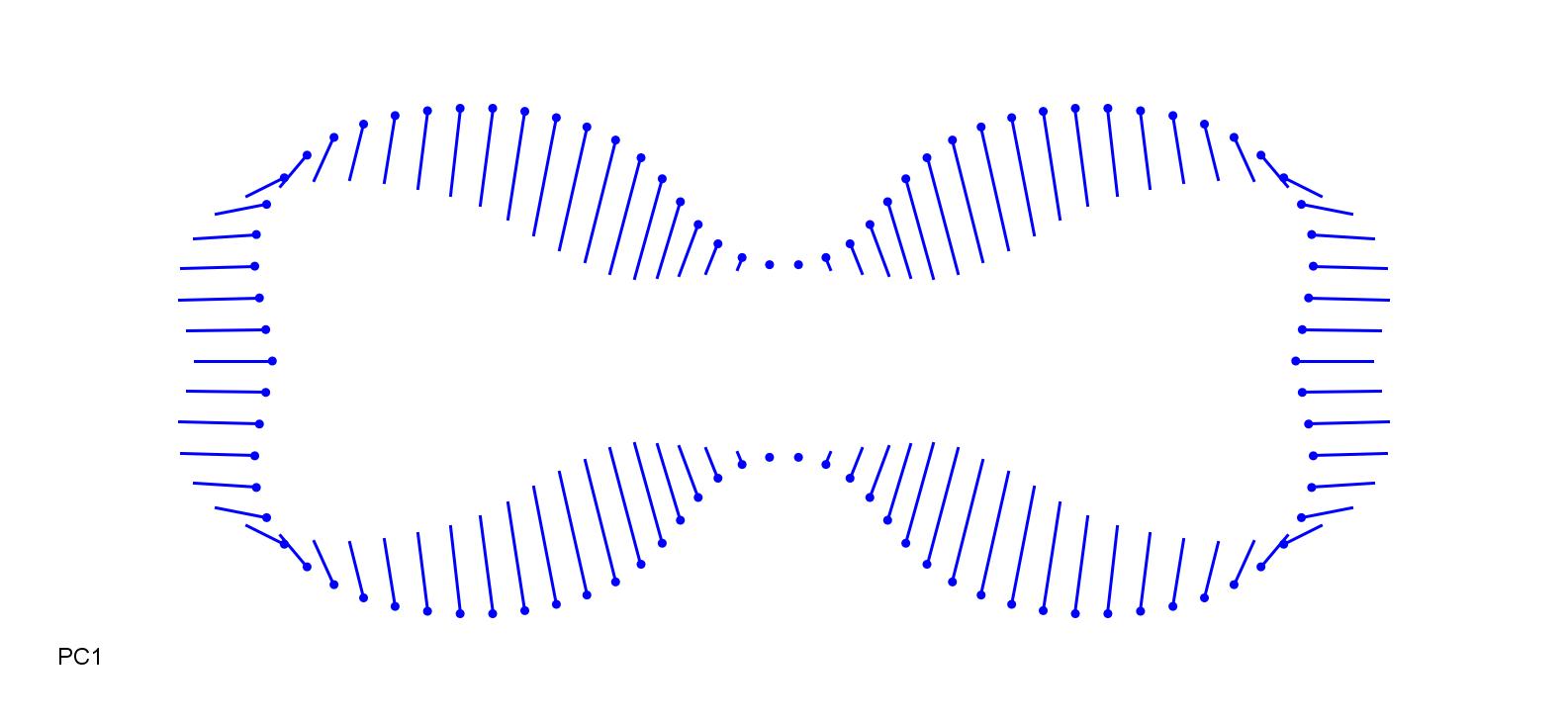

### SchiBar_0.15.jpg

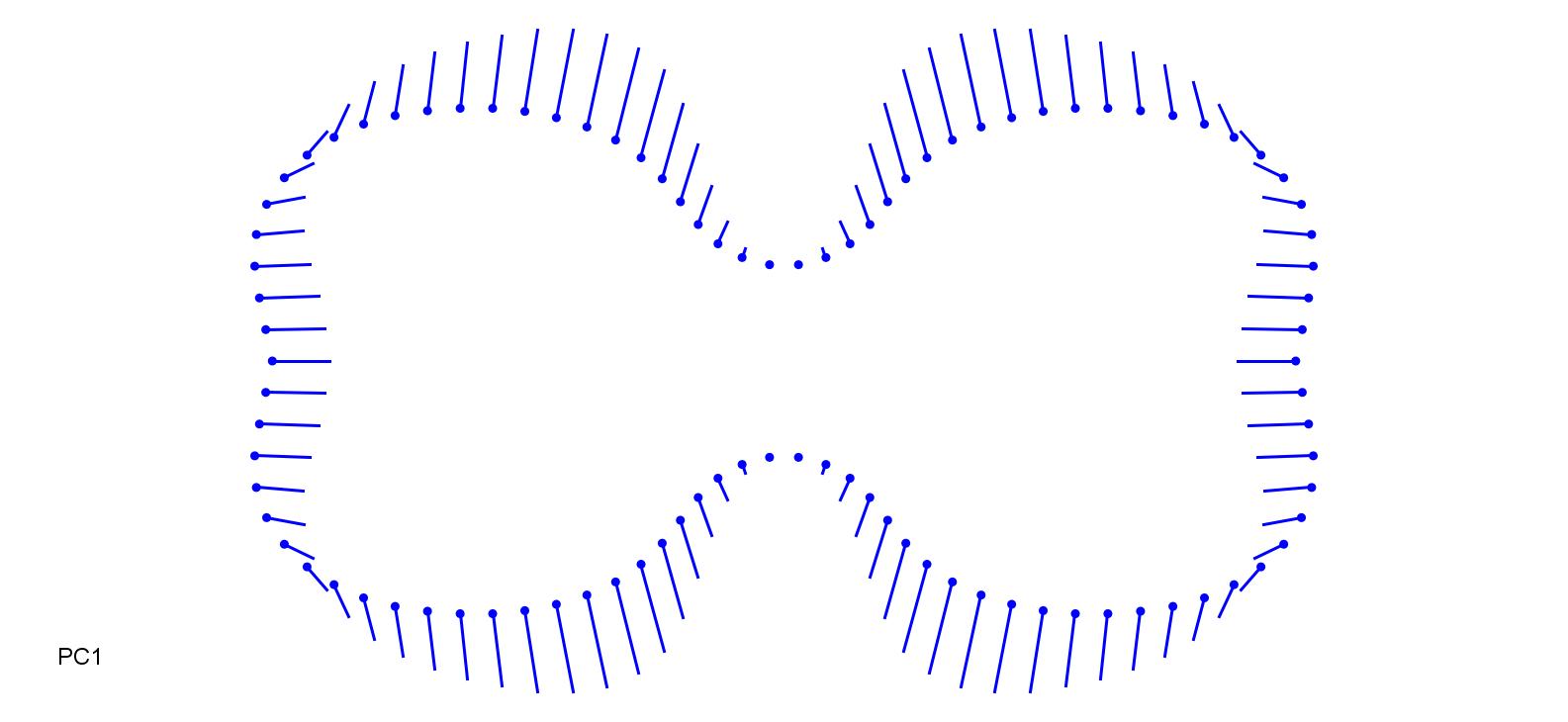

### SchPap-0.2.jpg

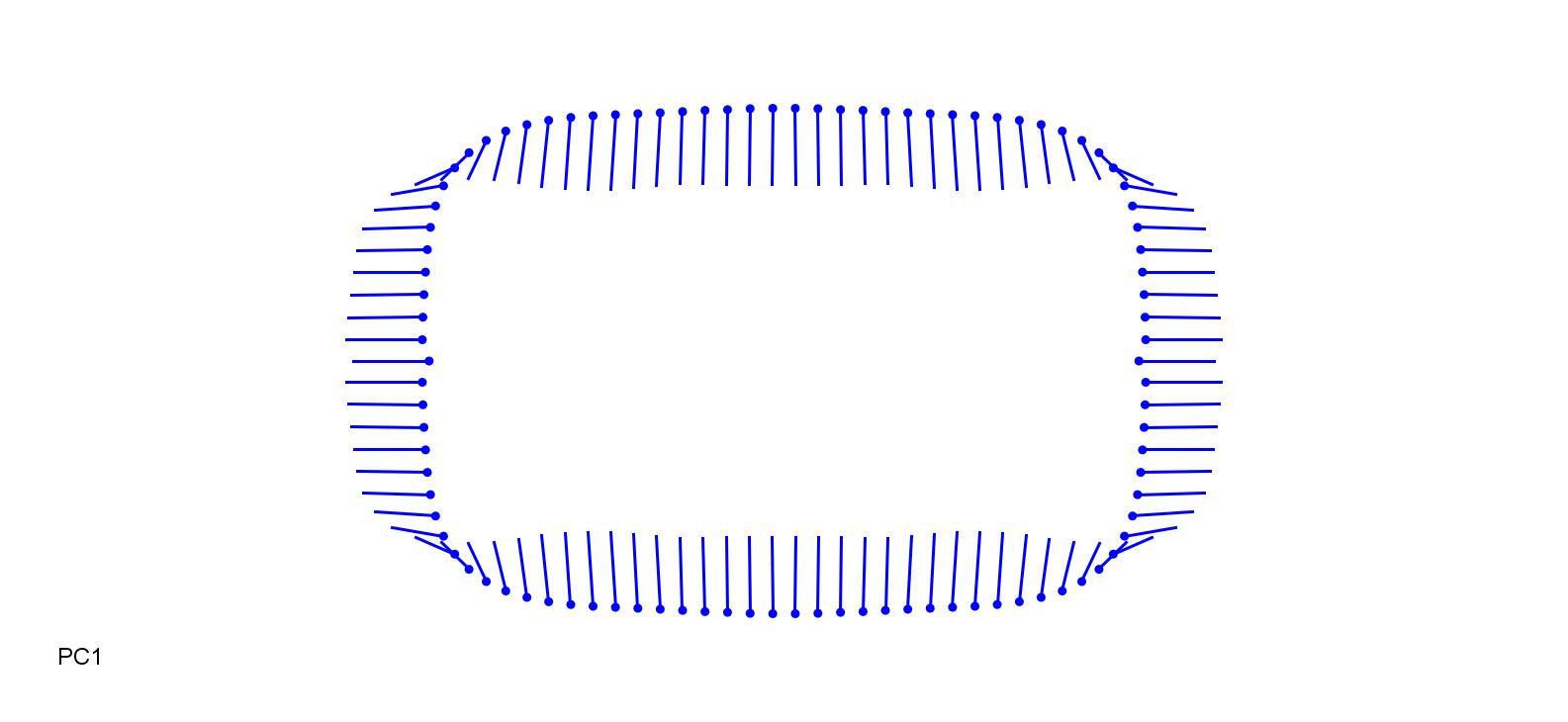

### SchPap_0.2.jpg

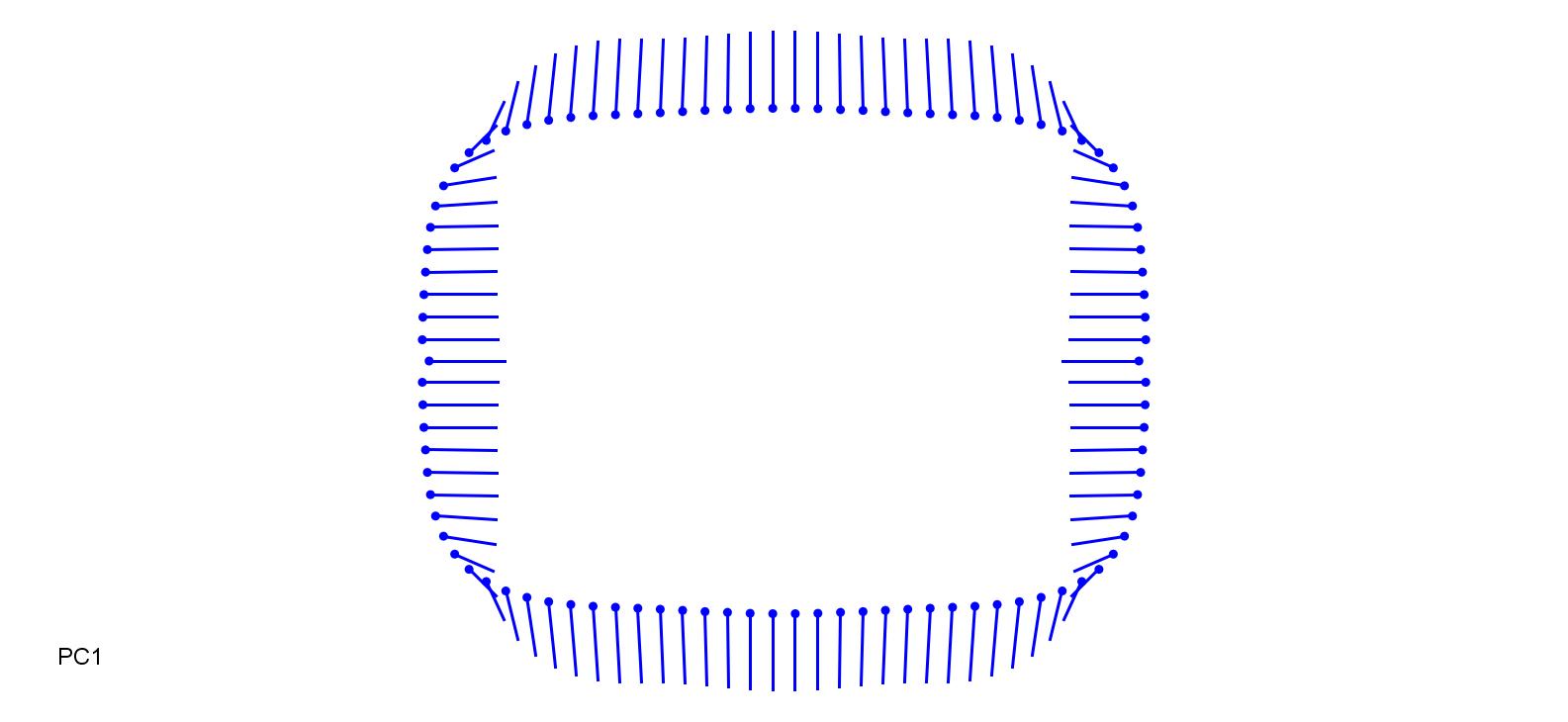

### SesCae-0.2.jpg

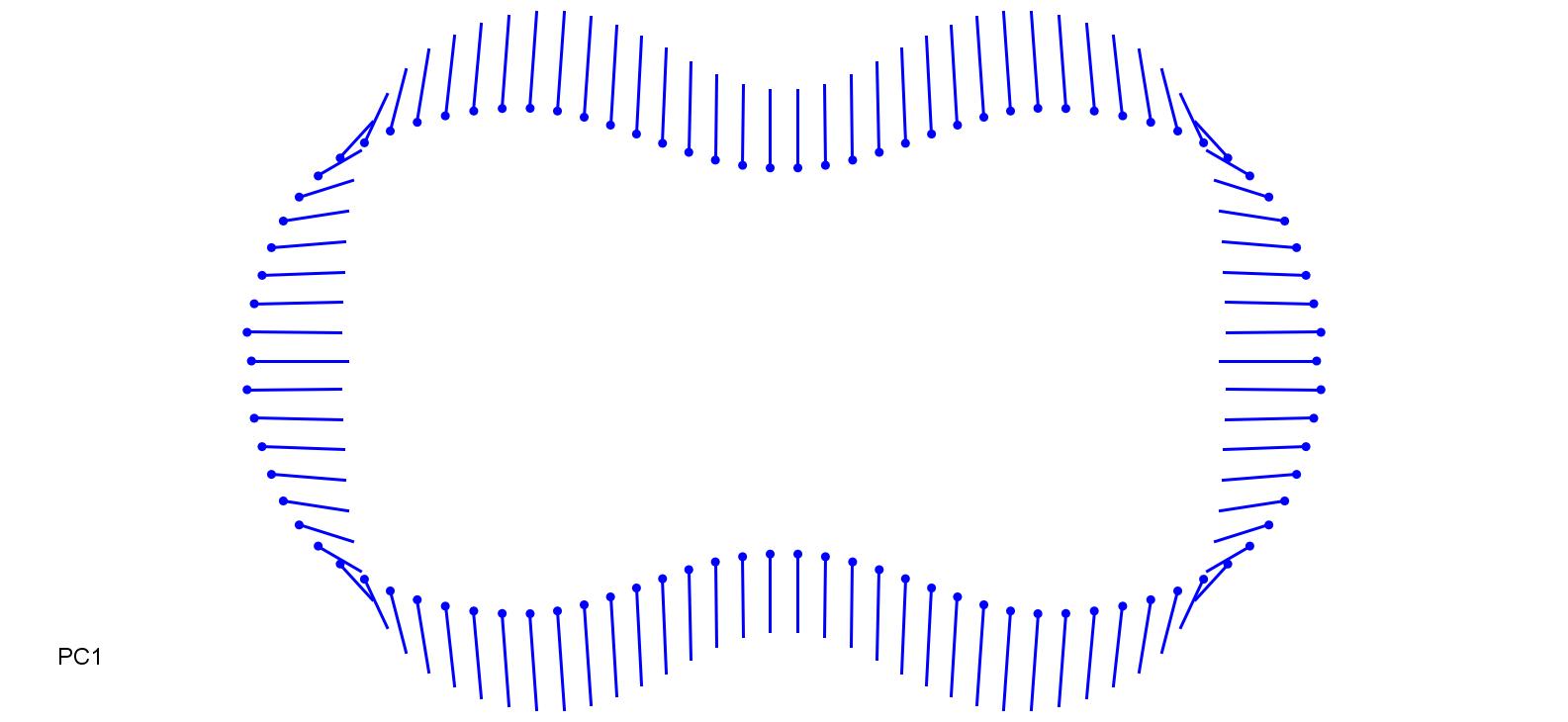

### SesCae_0.15.jpg

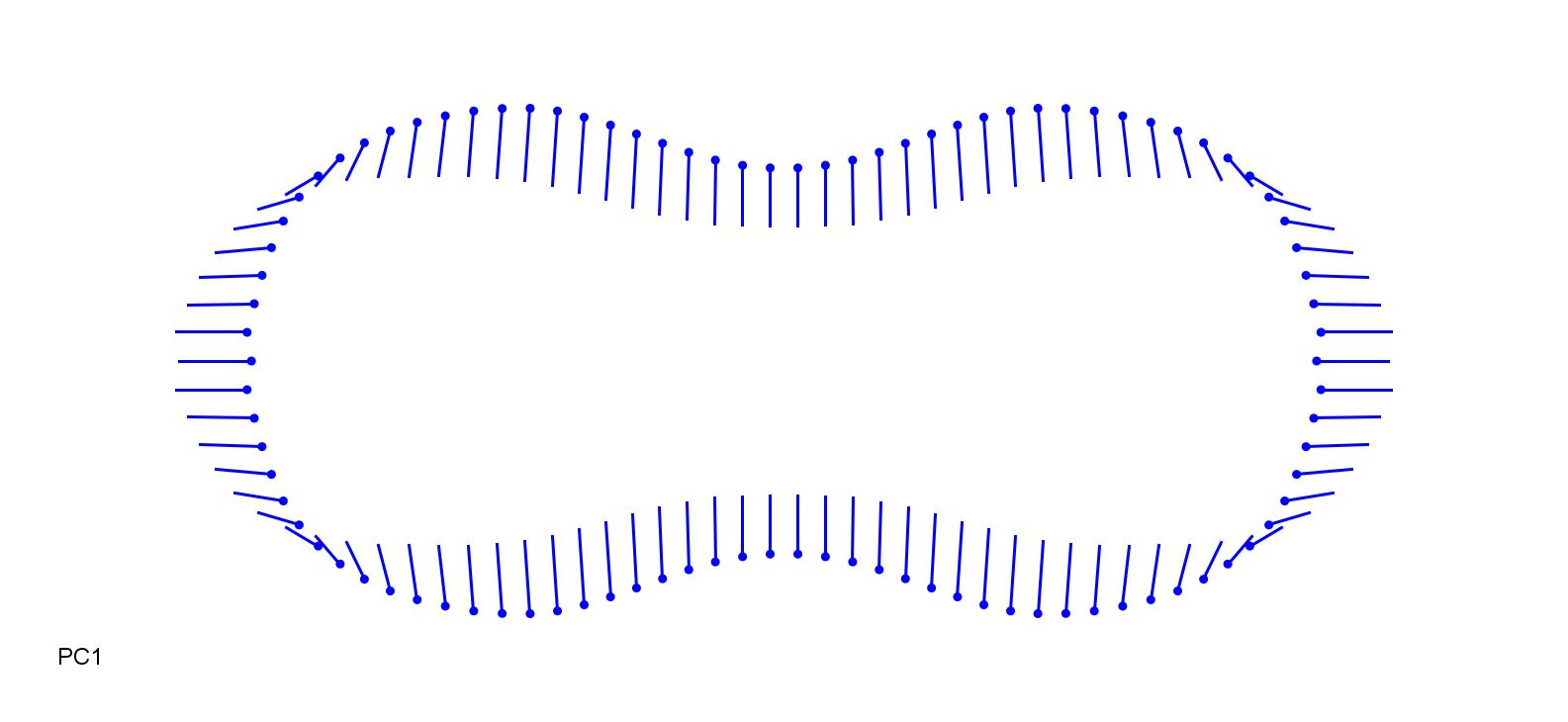

### StiPlu-0.15.jpg

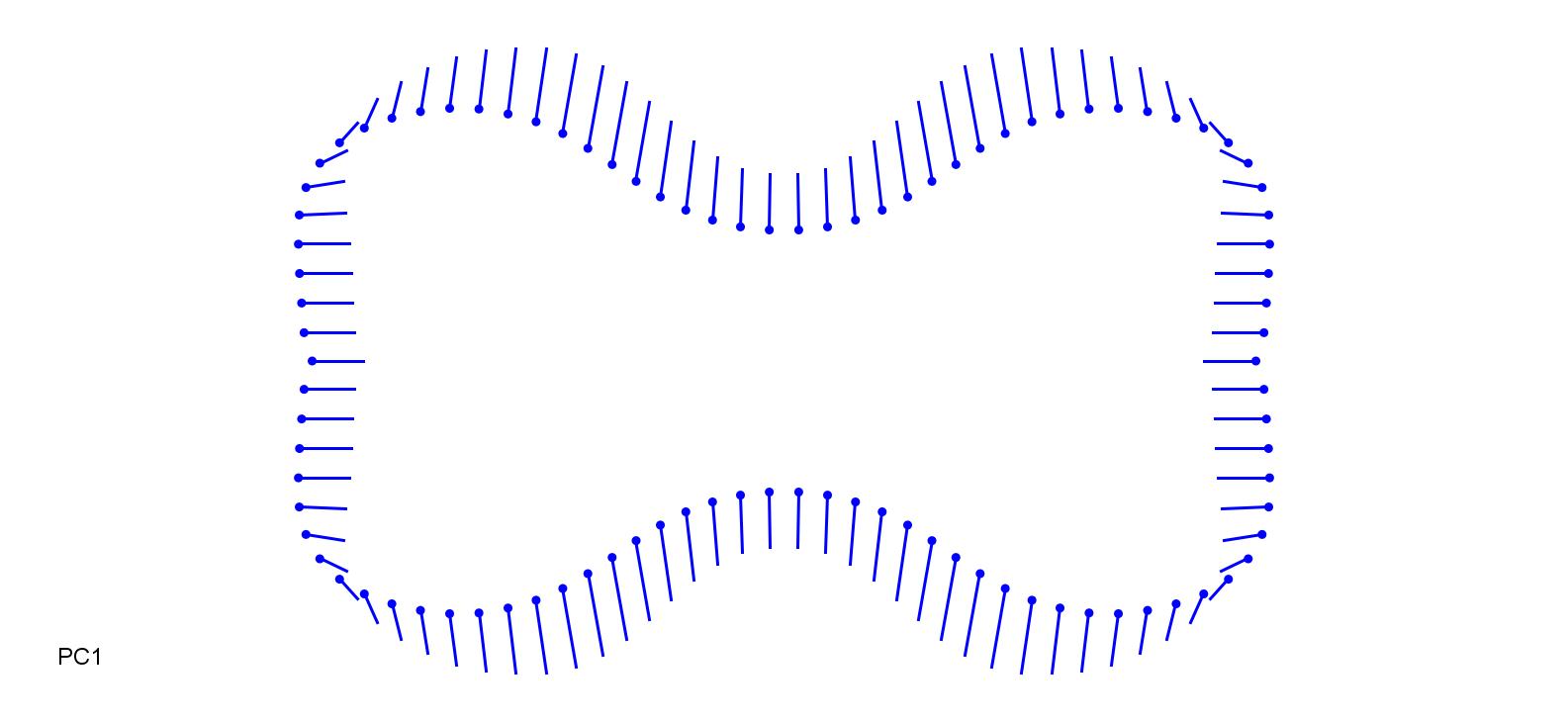

### StiPlu_0.15.jpg

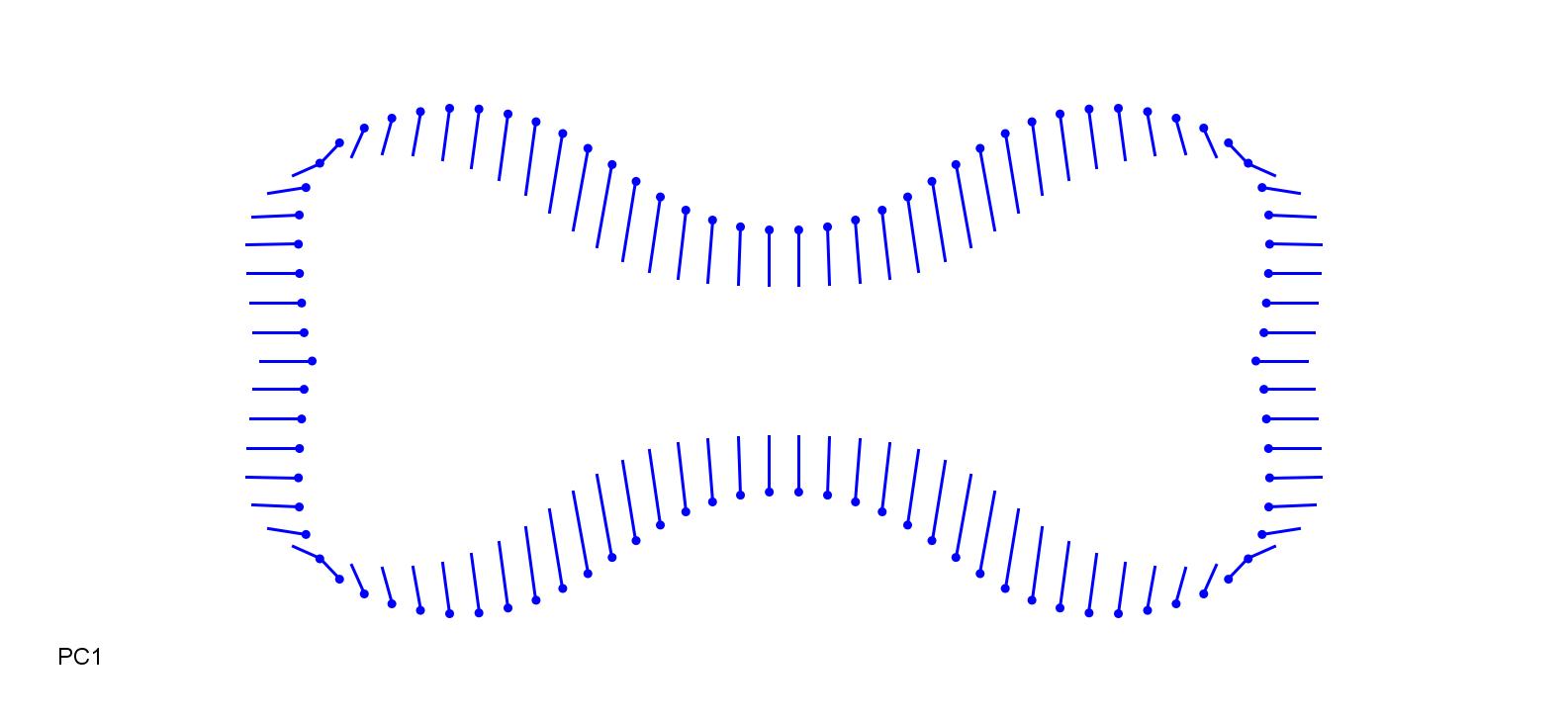

### TriFla-0.3.jpg

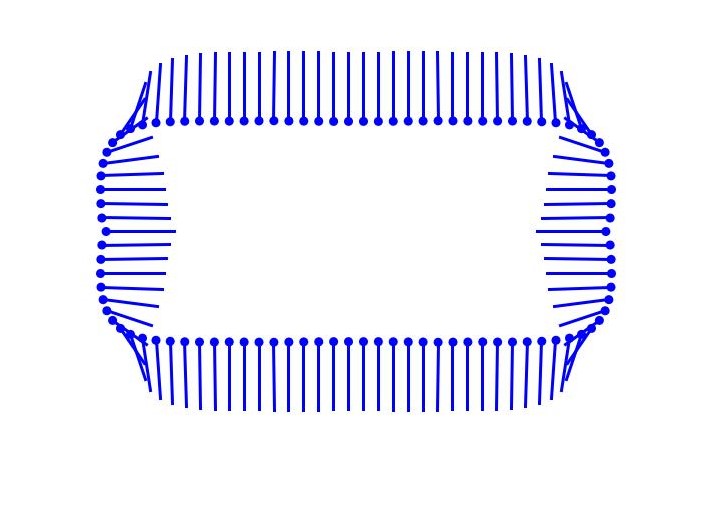

### TriFla_0.3.jpg

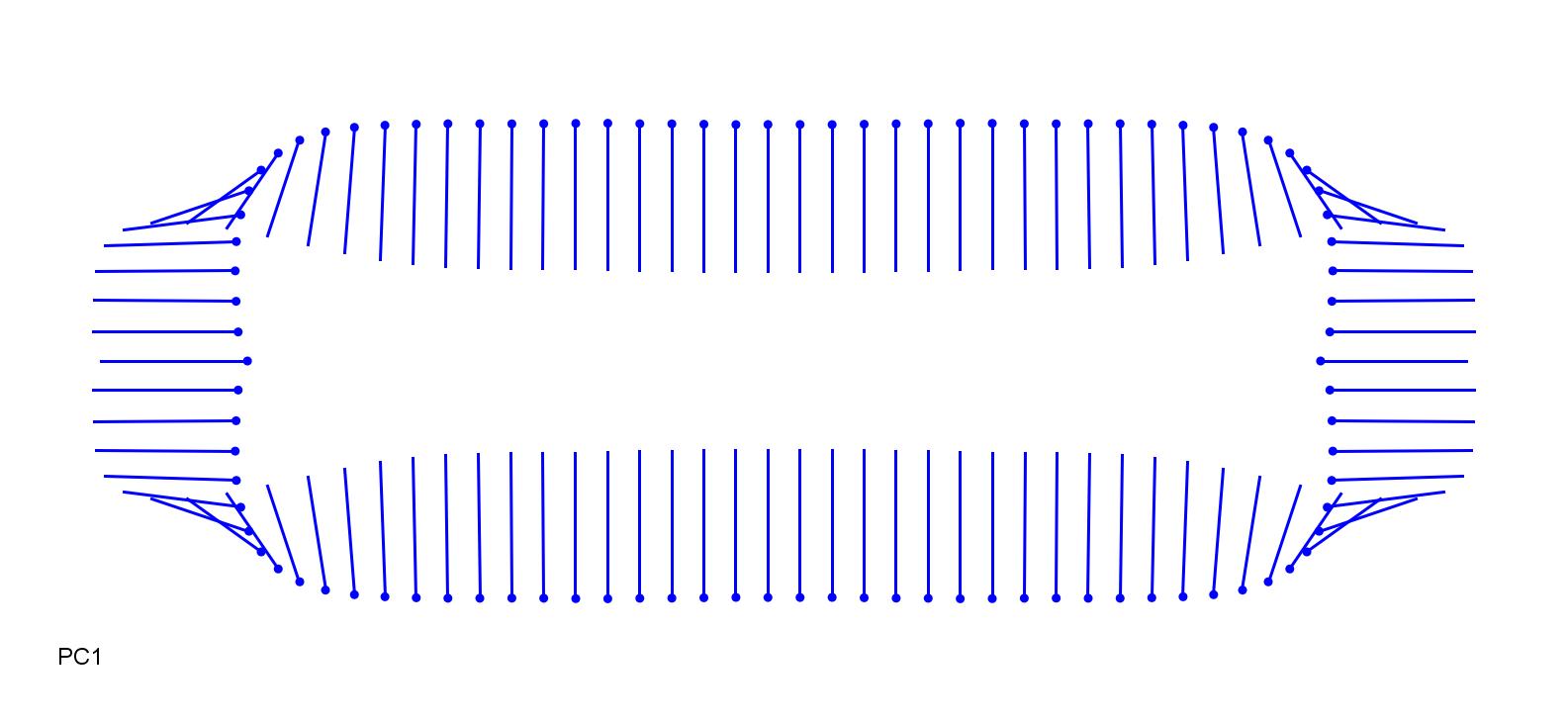

### ZizLat-0.1.jpg

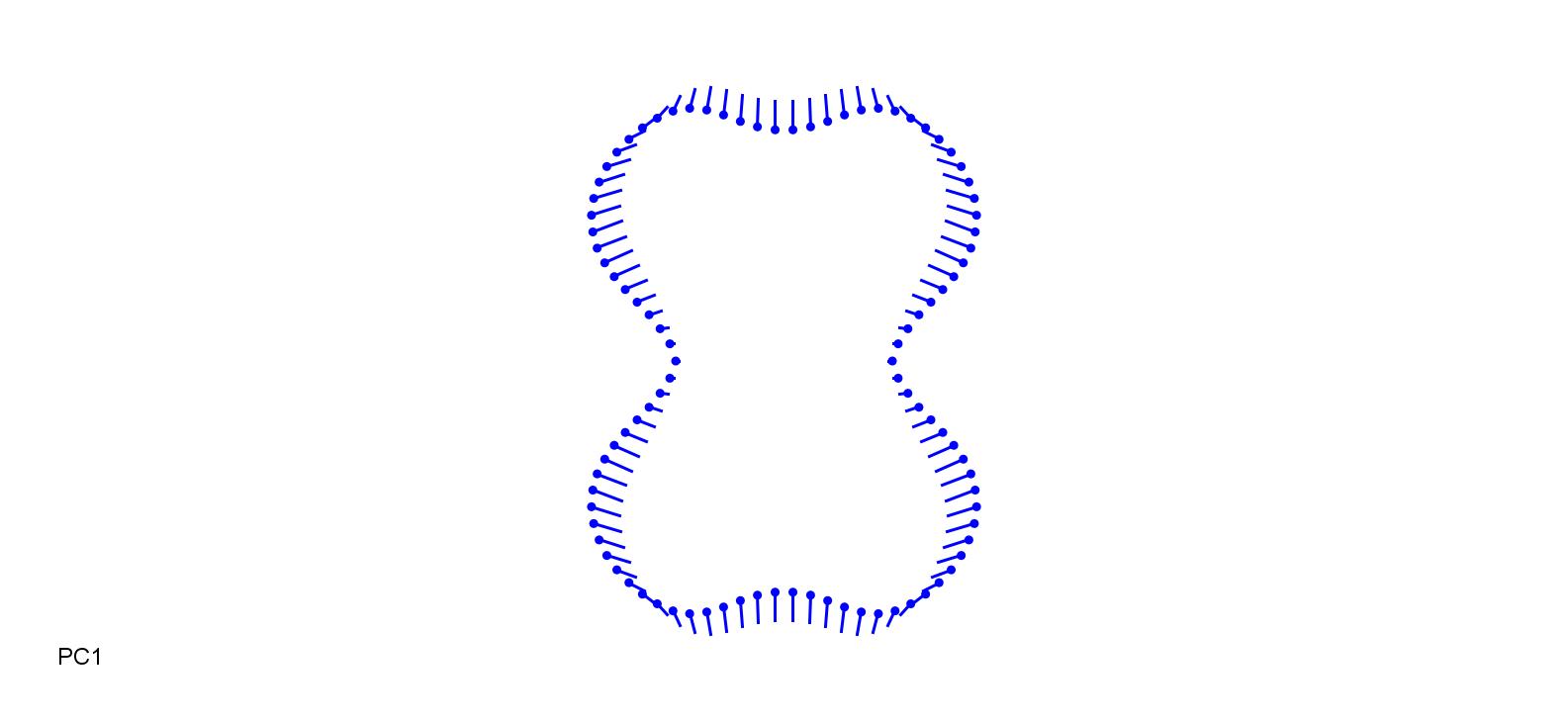

### ZizLat.jpg

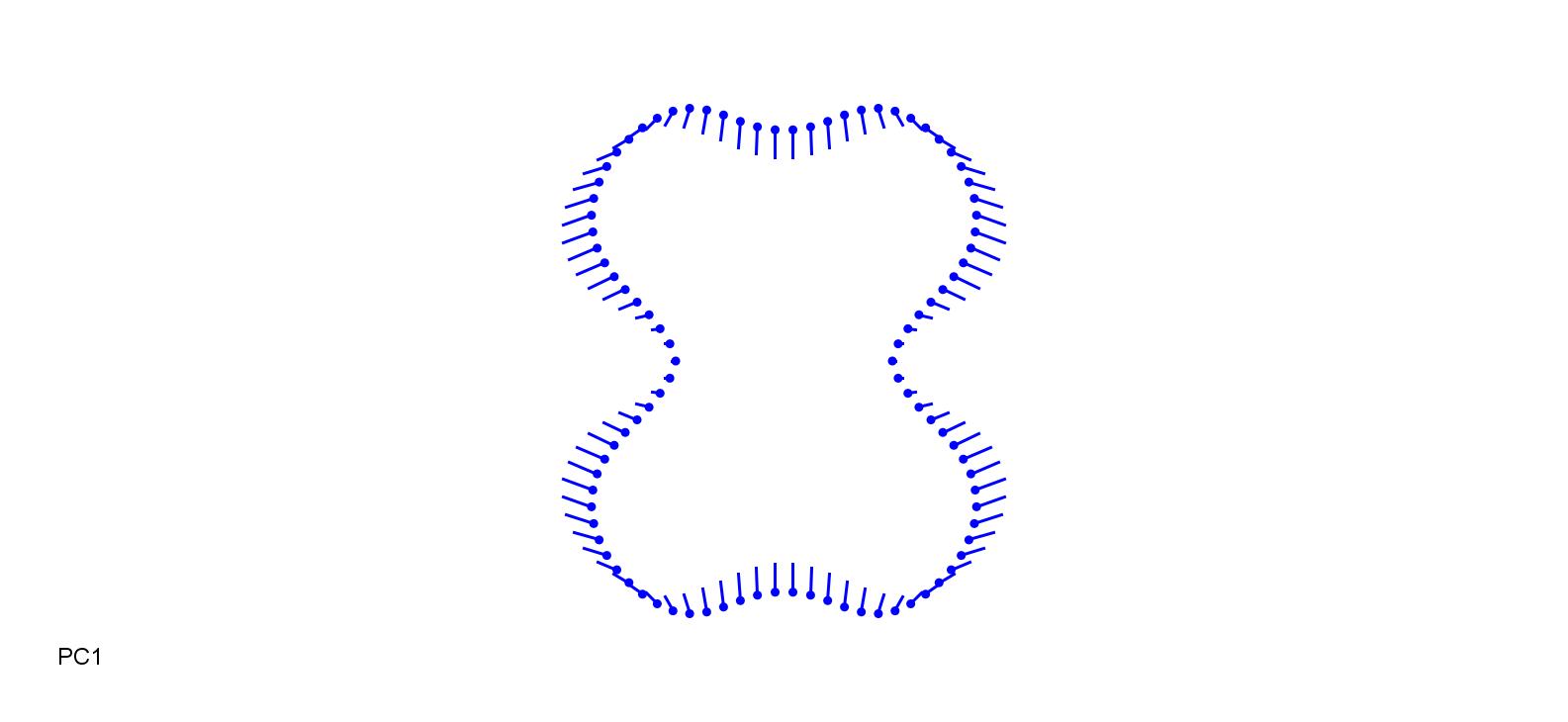
